# Deciphering Ibogaine’s Matrix Pharmacology: Multiple Transporter Modulation at Serotonin Synapses

**DOI:** 10.1101/2025.03.04.641351

**Authors:** Christopher Hwu, Václav Havel, Xavier Westergaard, Adriana M. Mendieta, Inis C. Serrano, Jennifer Hwu, Donna Walther, David Lankri, Tim Luca Selinger, Keer He, Rose Liu, Tyler P. Shern, Steven Sun, Boxuan Ma, Bruno González, Hannah J. Goodman, Mark S. Sonders, Michael H. Baumann, Ignacio Carrera, David Sulzer, Dalibor Sames

## Abstract

Ibogaine is the main psychoactive alkaloid produced by the iboga tree (*Tabernanthe iboga*) that has a unique therapeutic potential across multiple indications, including opioid dependence, substance use disorders, depression, anxiety, posttraumatic stress disorder (PTSD), and traumatic brain injury (TBI). We systematically examined the effects of ibogaine, its main metabolite noribogaine, and a series of iboga analogs at monoamine neurotransmitter transporters, some which have been linked to the oneiric and therapeutic effects of these substances. We report that ibogaine and noribogaine inhibit the transport function of the vesicular monoamine transporter 2 (VMAT2) with sub-micromolar potency in cell-based fluorimetry assays and at individual synaptic vesicle clusters in mouse brain as demonstrated via two-photon microscopy. The iboga compounds also inhibit the plasma membrane monoamine transporters (MATs), prominently including the serotonin transporter (SERT), and a novel iboga target, the organic cation transporter 2 (OCT2). SERT transport inhibition was demonstrated in serotonin axons and soma in the brain and in rat brain synaptosomes, where ibogaine and its analogs did not act as substrate-type serotonin releasers. Noribogaine showed dual inhibition of VMAT2 and SERT with comparable potency, providing an explanatory model for the known neurochemical effects of ibogaine in rodents. Together, the updated profile of the monoamine transporter modulation offers insight into the complexity of the iboga pharmacology, which we termed “matrix pharmacology”. The matrix pharmacology concept is outlined and used to explain why ibogaine and noribogaine do not induce catalepsy, as demonstrated in our study, in contrast to other VMAT2 inhibitors.

**TOC Graphic:** 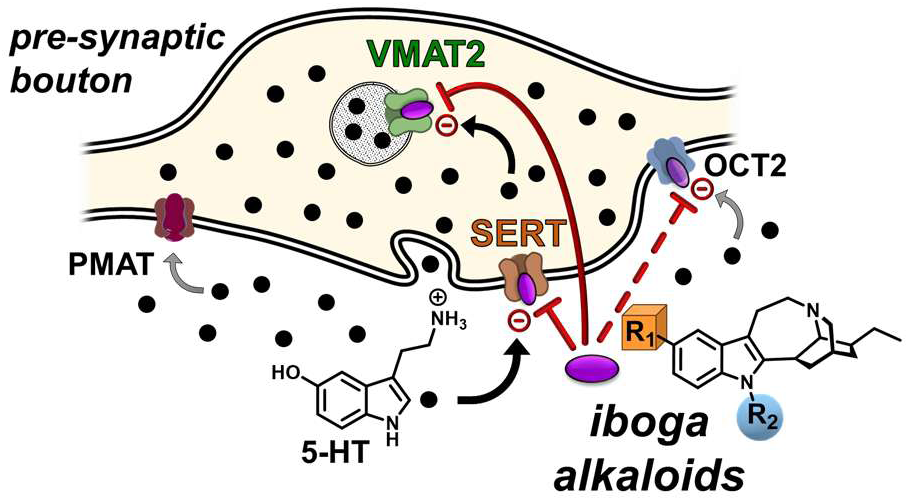

## Introduction

Ibogaine is the main psychoactive principle of the iboga plant (*Tabernanthe iboga*), indigenous to West Central Africa.^1^ The iboga root has been used for centuries to induce visionary states and transpersonal spiritual experiences during ceremonies and has served as a center piece for indigenous religions and spiritual practices.^2^ Outside traditional use, ibogaine and iboga were reported to interrupt dependence on opioids and other reinforcing drugs, including alcohol and cocaine.^3,4^ Through the work of several generations of pioneers, pursued mostly outside the mainstream medical system, ibogaine has provided a new paradigm for treating substance use disorders (SUDs) – albeit a controversial one due to its adverse cardiac effects.^5,6^ This paradigm consists of one or a few treatment sessions with ibogaine, which have acute and long-lasting effects as reported in numerous observational surveys and case studies.^3,7^ Acute effects include dream-like experiences (oneiric effects) that often provide deep psychological insights reduced physiological drug dependence (i.e., large effects in reducing acute withdrawal symptoms). Long-term effects include attenuation of drug craving, and improvement in mood and anxiety symptoms. It may be that ibogaine provides a multi-faceted, global treatment that addresses the core aspects of SUDs.^8,9^ The clinical observations have been supported by numerous preclinical studies showing therapeutic-like effects of ibogaine in rodent models of SUDs and depression.^10,11^

However, administration of ibogaine or iboga materials can lead to rare but severe cardiac adverse effects and sudden death, which have presented substantial obstacles to the development of ibogaine as an FDA-approved medication.^12^ Nevertheless, many individuals desperate for relief from their suffering have sought ibogaine treatment in clinics abroad, a growing trend that has provided pilot clinical observations.^13^

In addition to the anti-addiction effects of ibogaine, its putative therapeutic scope encompasses other psychiatric and neurological conditions, often co-morbid with SUDs, including PTSD and mild traumatic brain injury (mTBI), according to recent surveys of combat veterans with blast exposures.^14,15^ Although the clinical data supporting therapeutic efficacy of ibogaine are almost exclusively based on open label observational studies, the reported effect sizes are notably large across neurological and mental health assessment scales (for example, >90% response rates and 80% remission after a single treatment).^15^ Collectively, the current evidence indicates a trans-diagnostic therapeutic potential of ibogaine rooted in profound neurorestorative effects across biological scales (the molecular, physiological, psychological, and social-spiritual domains).

The emerging therapeutic profile of ibogaine leads to a fundamental question of how such a range of neurorestorative effects may be induced and mediated on a molecular level. Ibogaine is metabolized *in vivo* to form noribogaine, in both humans and rodents, where these two compounds act in tandem due to overlapping pharmacokinetic profiles.^11,16–18^ The reported pharmacological targets and associated effects provide a window on the complexity of signaling pathways that ibogaine and noribogaine engage. The combined action of these two compounds modulates several classes of molecular targets and pathways, including ion channels, neurotransmitter transporters, G protein coupled receptors (GPCRs) and their signaling, and kinase signaling pathways via several modes of action such as ion channel blockade,^19^ monoamine transporter inhibition via an atypical mechanism,^20^ monoamine transporter pharmaco-chaperoning effect,^21,22^ and kinase signaling pathway potentiation.^23^ Interestingly, Interestingly, these wide-ranging effects typically involve low potency drug interactions in the micromolar range (Figure 1).^24–27^

**Figure 1.**
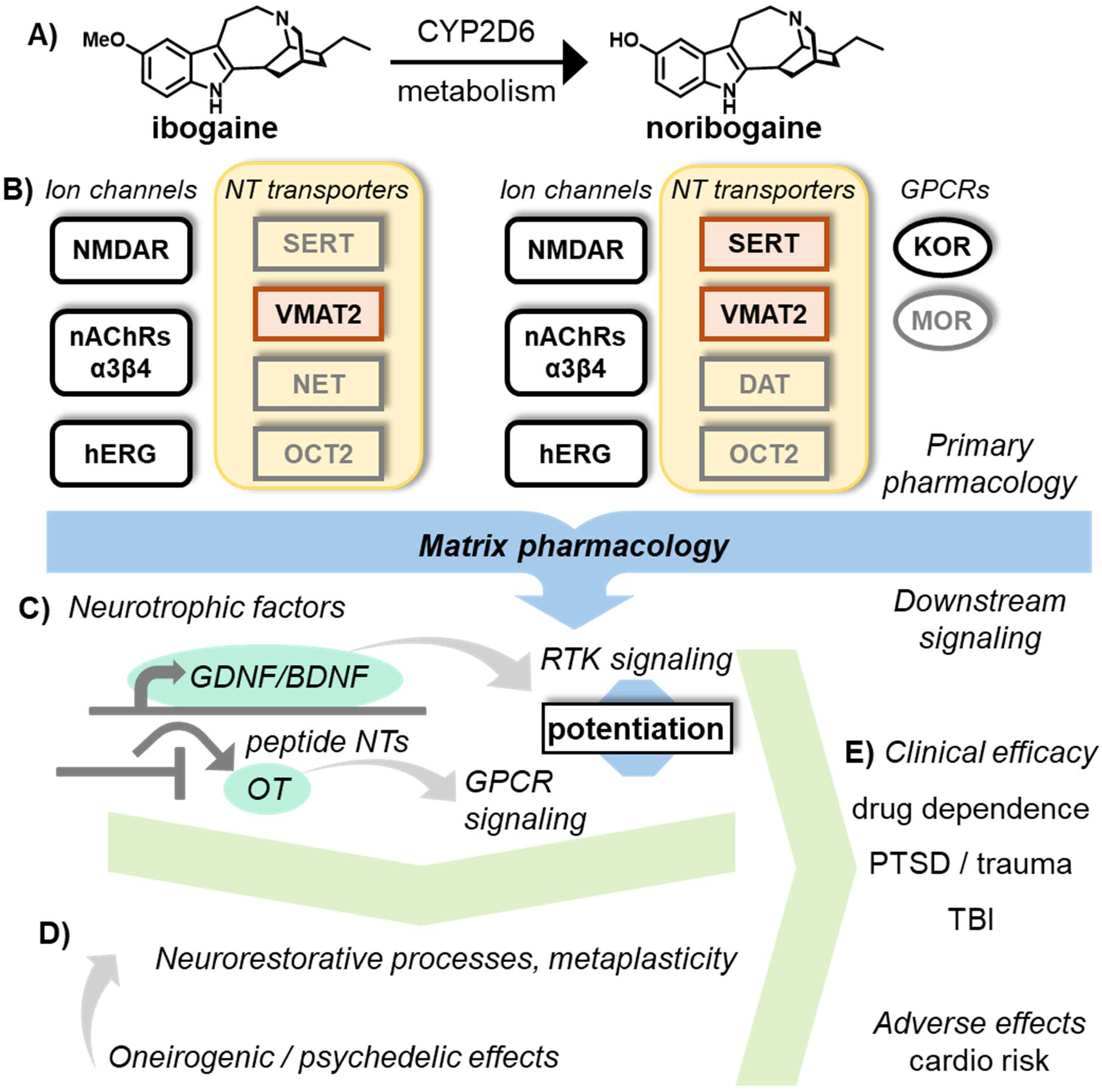
Proposed global model of ibogaine’s pharmacology, downstream molecular signaling, neuro-restorative processes, and clinical therapeutic and adverse effects. (A) Ibogaine is metabolized to noribogaine and both compounds contribute to the complex pharmacology. (B) Examples of known primary molecular targets; the present study focuses on monoamine transporters including the vesicular monoamine transporter 2 (VMAT2) and organic cation transporter 2 (OCT2), disclosed as novel targets in this study. Ibogaine and noribogaine act as weak and modest potency modulators of several targets and signaling pathways, via multiple modes of action, such as blockers of ion channels, atypical inhibitors of monoamine transporters, pharmaco-chaperones of monoamine transporters (SERT and DAT), and potentiators of downstream signaling pathways – a functional profile termed “matrix pharmacology”. (C) The downstream effects include, for example, expression changes of neurotrophic factors such as glial cell derived neurotrophic factor (GDNF) and brain derived neurotrophic factor (BDNF), and potentiation of receptor tyrosine kinase signaling pathways. Oxytocin (OT) signaling has been implicated in mediating reopening of the critical development-like learning states. (D) According to this mechanistic model, the indicated pathways mediate the neuroplasticity and neurorestorative processes, where the oneiric and psychedelic states likely contribute as well. (E) These processes collectively likely mediate the profound therapeutic effects across biological scales.

Here, we introduce the term “matrix pharmacology” – to describe this mechanistic model, defined as modulation of many molecular targets and signaling pathways via multiple modes of action with weak or modest-potency molecular interactions. As the collective data suggests that iboga compounds act throughout the signalome’s information-processing matrix, we wish to differentiate this mechanistic model (matrix pharmacology) from “selective polypharmacology”, which is characterized by potent agonism or antagonism at several targets, such as G protein coupled receptors (GPCRs), as exemplified by LSD.^28,29^ A molecular logic of iboga matrix pharmacology is emerging where iboga compounds do not modulate GPCRs directly (with some exceptions) but rather 1) modulate neurotransmitter dynamics via direct inhibition and modulation of the monoamine transporters, 2) engage ion channels as channel blockers and or antagonists, and 3) potentiate downstream signaling pathways of GPCRs and receptor tyrosine kinases (RTKs).

As downstream effectors activated by the primary pharmacology, iboga compounds modulate gene expression and protein levels of neurotrophic factors in several brain regions with relevance to SUDs and other mental health disorders.^30,31^ Upregulation of myelination markers was reported in morphine-dependent mice after a single ibogaine administration, indicating the possibility of white matter restoration.^32^ It was also suggested that ibogaine may increase oxytocin gene expression,^33^ while another report demonstrated induction of metaplasticity states in the nucleus accumbens (NAc) that sensitize the local synaptic connections to oxytocin, which jointly correlates with reopening a critical learning period for social reward in adult mice (Figure 1).^34^ These results, together with behavioral effects, indicate induction of both specific and broad restorative programs and processes (e.g. reversal of drug dependence phenotype and induction of metaplasticity and neurorestoration states).

In this report, we focus on mapping the actions of iboga compounds at the monoamine transporters, some of which have previously been implicated in ibogaine’s anti-addiction and oneirogenic effects.^20^ The plasma membrane monoamine transporters (MATs), including the dopamine transporter (DAT), norepinephrine transporter (NET), and serotonin transporter (SERT), are essential components of monoamine neurotransmission, as they directly modulate the magnitude and duration of the synaptic activity by rapid removal and recycling of the monoamine neurotransmitters from the extracellular space. They constitute a branch of a solute carriers 6 family (*SLC6*; also known as neurotransmitter sodium symporters, or NSS) that mediate a directional transport coupled to the membrane ion gradient by co-transporting the endogenous monoamine transmitters and sodium ions.^35,36^ DAT and NET are located in the corresponding catecholamine neurons, whereas SERT is expressed in 5-HT neurons, blood-brain barrier (BBB) cells, endochromaffin cells, and blood platelets.^37,38^ MATs are prominent targets of psychoactive substances and medications that inhibit one or more monoamine transporters, including cocaine, methylphenidate, tricyclic antidepressants, and selective serotonin reuptake inhibitors.^39–41^ There are also targets for monoamine releasers, such as amphetamine and MDMA, that act as MAT substrates and induce efflux of native monoamine transmitters via these transporters, often referred to as “reverse transport”.^42^

The functional dynamic range of MAT transporters, described as high-affinity, low-capacity transporters (uptake 1 system) is complemented by the low-affinity, high-capacity transporters (uptake 2 system) comprised of transporters from several different *SLC* families including organic cation transporters 1-3 (OCT1-3, *SLC22A1-3*), plasma membrane monoamine transporter (PMAT, *SLC29A4*), and other more recently identified transporters.^43^ These transporters generally show much lower apparent affinity for biogenic monoamines (as much as two orders of magnitude), but greater maximal transport velocity than MATs, and are driven by the concentration gradient of the substrates (equilibrative transporters). OCT1-3 and PMAT have distinct but often overlapping expression patterns in the CNS on neurons (presynaptic and postsynaptic) and glial cells (Figure 2).^43^

**Figure 2.**
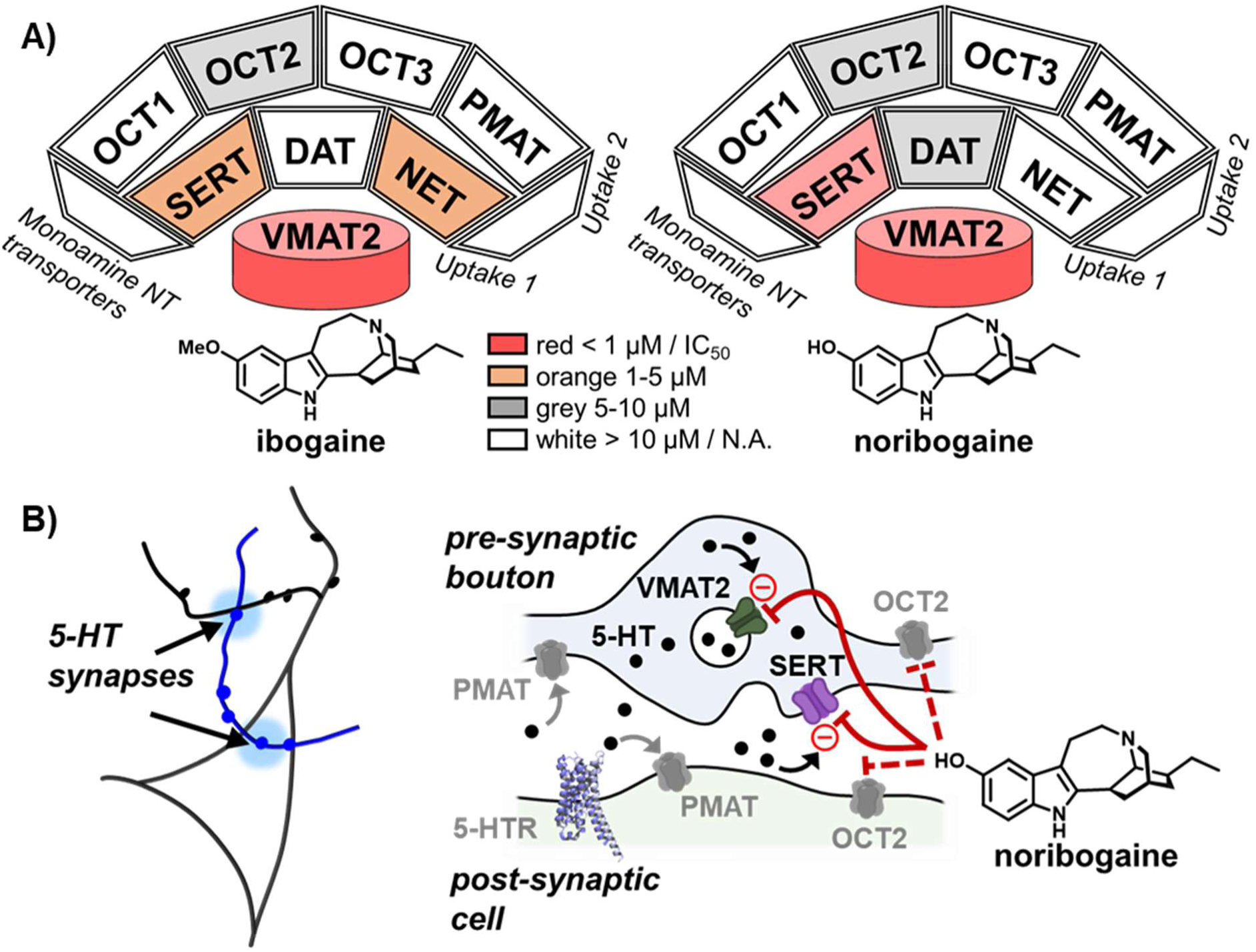
Ibogaine and noribogaine exert dual inhibition of vesicular and plasma membrane transporters. (A) Graphical representation of ibogaine’s (left panel) and noribogaine’s (right panel) inhibitory profile at the vesicular monoamine transporter 2 (VMAT2), uptake 1 and uptake 2 monoamine transporters. Ibogaine inhibits hVMAT2 with submicromolar potency (IC_50, hVMAT2_ = 390 nM) and modest selectivity of around one log unit over hSERT (IC_50, hSERT_ = 2980 nM) and hNET (IC_50, hNET_ = 3670 nM), while having a weak effect at hOCT2 (IC_50, hOCT2_ = 9310 nM). Noribogaine inhibits hVMAT2 (IC_50, hVMAT2_ = 570 nM) and hSERT (IC_50, hSERT_ = 280 nM) with comparable potency. This dual activity is separated by more than one log unit from the effect of noribogaine at hDAT (IC_50, hDAT_ = 6760 nM) and hOCT2 (IC_50, hOCT2_ = 6180 nM). (B) A schematic showing a 5-HT axon (blue line), 5-HT release sites (blue dots) and 5-HT synapses (lighter and larger blue dots) on a pyramidal neuron’s soma and dendrite (left graphic). A pictorial representation of 5-HT synapse and location of VMAT2 on synaptic vesicles, SERT on presynaptic boutons, and uptake 2 transporters OCT2 and PMAT on presynaptic and postsynaptic cells (localization on glial cells is omitted for clarity). Noribogaine has a dual inhibitory effect on VMAT2 and SERT, suggesting complex, non-linear effects on 5-HT and monoamine neurotransmission.

MATs work in tandem with the vesicular monoamine transporters, VMAT1 (*SLC18A1*) and VMAT2 (*SLC18A2*), which load synaptic vesicles or large dense core vesicles with monoamine neurotransmitters as an essential step prior to activity-dependent release via exocytosis (Figure 2B). VMAT1 is expressed in neuroendocrine cells, while VMAT2 is largely found in neurons in the CNS. Both transporters concentrate neurotransmitters in the vesicle lumen, driven by the proton gradient between the cytoplasm and vesicular lumen, via an amine-proton antiport mechanism.^44,45^ The cytosolic monoamines that are not packaged into synaptic vesicles are oxidatively deaminated by the monoamine oxidases. Reserpine and tetrabenazine are classical VMAT2 inhibitors; reserpine (a dual VMAT1 and 2 inhibitor) has been used in the past to treat hypertension and psychosis, while TBZ (and its deuterated analog, selective VMAT2 inhibitors) are prescribed to treat chorea associated with the Huntington’s disease.^46^

The VMAT2 and the plasma membrane uptake 1 and uptake 2 transporters jointly shape the spatial and temporal landscape of monoamine neurotransmission in the CNS and its perturbation by pharmaceutical agents. Biogenic monoamines act as modulatory neurotransmitters via their respective receptors by augmenting or attenuating the excitatory and inhibitory synapses and serve as critical neuroplasticity mediators.^47^

Guided by the framework of matrix pharmacology and the substantial corpus of neurochemical studies with ibogaine and noribogaine in rodents, we hypothesized that there are one or more important molecular targets of iboga alkaloids in the monoamine transporter families, which have not been examined in detail previously (the “iboga X project”). The effects of ibogaine or noribogaine on monoamine neurotransmitter dynamics *in vivo* are complex; for example, the brain tissue analysis in mice and rats showed, in terms of total tissue concentrations, a decrease in dopamine and increase in the dopamine metabolite, DOPAC (dihydroxyphenylacetic acid), which is a tell-tale sign of VMAT2 inhibition.^20^ Weak binding affinities of ibogaine and noribogaine at VMAT2 in human brain striatal homogenates (ibogaine, IC_50_ = 14.6 μM; and noribogaine, IC_50_ = 29.5 μM) have been reported, supporting the hypothesis that VMAT2 may be directly inhibited by the iboga compounds.^48^ Ibogaine and noribogaine were also reported to acutely raise extracellular 5-HT levels in the Nucleus accumbens (NAc), while the total tissue content analysis revealed minimal or no 5-HT decrease and a more pronounced decrease of the 5-HT metabolite, 5-HIAA (5-hydroxyindolylacetic acid), consistent with SERT inhibition.^20^ These measurements indicate a complex monoamine modulation scenario and provides a rationale to systematically examine monoamine transporters, including VMAT2 and the uptake 1 and uptake 2 plasma membrane monoamine transporters.

Our previous work revealed that even small structural perturbations of the iboga system can have large effects on *in vitro* and *in vivo* pharmacology, as demonstrated by the oxa-iboga class of compounds (benzofuran-based iboga analogs).^31^ We therefore examined a focused SAR space of the ibogamine core in terms of the monoamine transporter activity.

## Results

### Ibogaine and noribogaine inhibit the function of multiple monoamine transporters

To examine the effect of ibogaine and noribogaine on human VMAT2 (hVMAT2), we used HEK293 (human embryonic kidney 293) cells stably transfected with hVMAT2 (hVMAT2-HEK cells) and a VMAT2 fluorescent substrate, FFN206 (Figures S2).^49^ For uptake 1 plasma membrane monoamine transporters, we used APP+ [4-(4-dimethylamino)phenyl-1-methylpyridinium] as the validated fluorescent substrate and the corresponding stably transfected HEK cells (hSERT-HEK, hDAT-HEK, and hNET-HEK). These cell-based assays provide an optical readout for the fluorescent substrate uptake (fluorescence units) mediated by the selected transporter.^50^ The extent of inhibition induced by the tested compounds is quantified by the difference in the fluorescence signal between the vehicle and the iboga compound treatments (Figures S1, S4, and S5). Similar protocols were developed for the uptake 2 transporters, OCT1-3 and PMAT, using APP+ (OCT1 and PMAT) and ASP+ (4-[4-(dimethylamino)styryl]-*N*-methylpyridium; OCT2-3) as fluorescent substrates (Figures S6-S9). Established transporter inhibitors, including indatraline (DAT; Figure S4), reboxetine (NET; Figure S5), imipramine (SERT; Figures 3 and S1), tetrabenazine (VMAT2; Figures 3 and S2), and decynium-22 (OCT1-3 and PMAT; Figures S6-S9) were used for each transporter as positive controls.^51^

**Figure 3.**
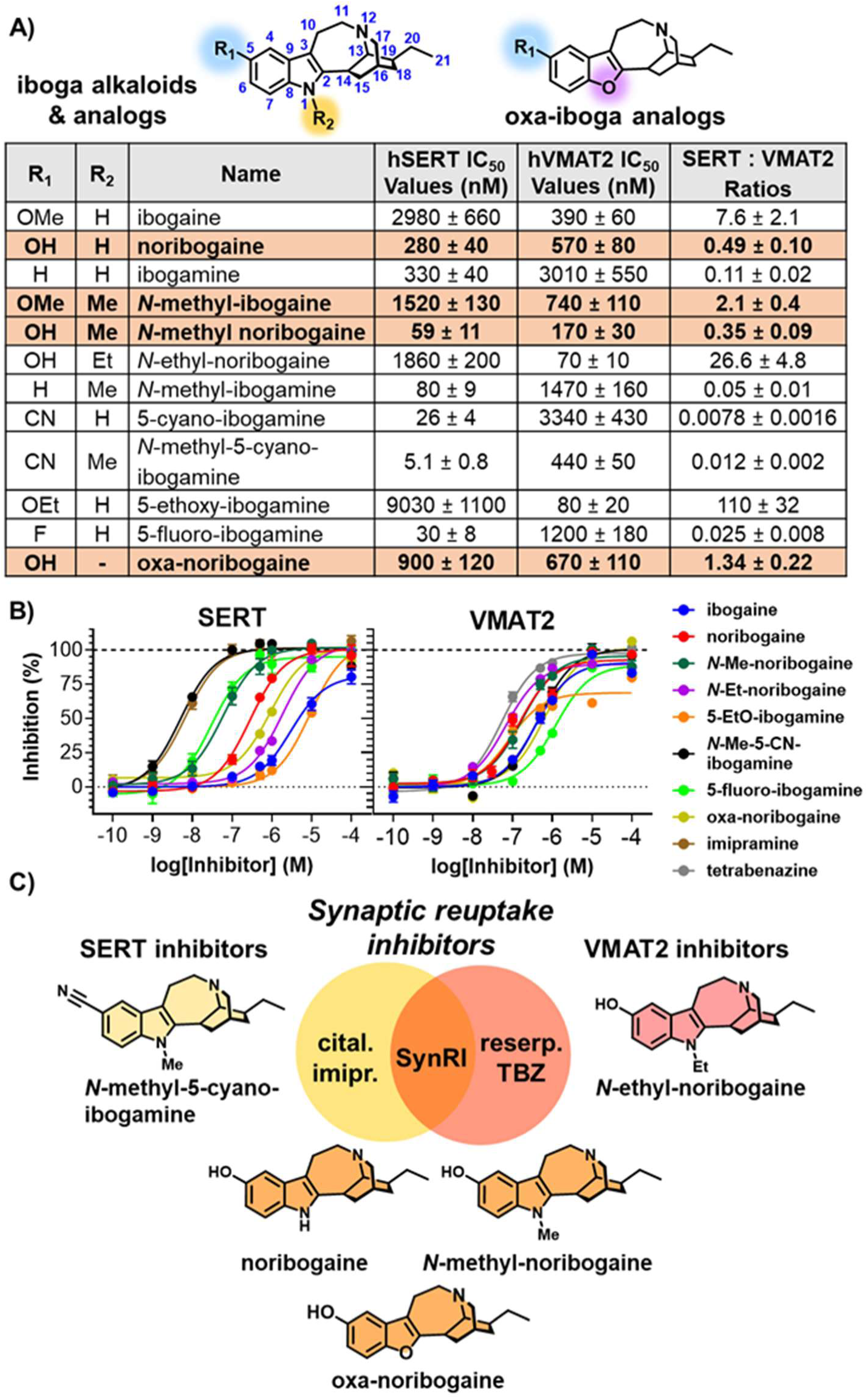
Iboga compounds define a new class of pharmacological agents, termed “synaptic reuptake inhibitors” or “SynRIs”. (A) Structure-activity relationship (SAR) examination of two positions of ibogamine and oxa-ibogamine (indicated on top, the 5-position in blue, the 1-position in yellow and purple) with respect to VMAT2 and SERT transport. We introduce a new numbering system in line with the numbering of simple tryptamines and indoles. The table provides IC_50_ values for transport inhibition obtained by fluorimetry based on inhibition of uptake of fluorescent substrates, APP+ for hSERT and FFN206 for hVMAT2, *n* = 4 biological replicates, mean ± SEM. Brick red shading highlights SynRI compounds with balanced inhibitory profiles; i.e., with IC_50_ values for SERT and VMAT2 that fall within a three-fold range (one-half log). (B) Dose-response curves for control inhibitor (for hSERT: imipramine and for hVMAT2: tetrabenazine) and selected compounds in hSERT-HEK cells (left panel) and hVMAT2-HEK cells (right panel). The y-axis shows normalized inhibition of fluorescent substrate uptake (uninhibited – inhibited) ± SEM, derived from four separate experiments (*n* = 4). (C) Graphical representation of the dual inhibitory activity at SERT and VMAT2 exhibited by noribogaine, *N*-methyl-ibogaine, *N*-methyl-noribogaine, and oxa-noribogaine. Minor modifications of the iboga structure yield a range of activities in the SERT and VMAT2 continuum including potent and selective SERT inhibitors and VMAT2 inhibitors, as well as balanced SynRI compounds.

We found that both ibogaine and noribogaine inhibit hVMAT2 transport with comparable potency (IC_50, ibogaine, hVMAT2_ = 390 nM, IC_50, noribogaine, hVMAT2_ = 570 nM, Figure 2A, Figure 3, and Figure S2). The values indicate more potent inhibition effect than those previously reported for hVMAT2 binding in a radioligand displacement assay in post-mortem human brain (IC_50, ibogaine, striatum_ = 14.6 μM and IC_50, noribogaine, striatum_ = 29.5 μM).^48^ Ibogaine shows approximately a one-log higher affinity for hVMAT2 over hSERT (IC_50, hSERT_ = 2980 nM) and NET (IC_50, hNET_ = 3670 nM) (Figures 3, S1, and S5). Noribogaine inhibits hVMAT2 (IC_50, hVMAT2_ = 570 nM) and SERT (IC_50, hSERT_ = 280 nM) with comparable potency, which represents an unusual pharmacological profile (Figures 2-3, and S1-S2). The VMAT2 and SERT dual inhibition is over 10-fold more potent than the effect of noribogaine at DAT (IC_50, DAT_ = 6760 nM) (Figure S4). In terms of uptake 2 transporters, we found that ibogaine (IC_50, OCT2_ = 9310 nM) and noribogaine (IC_50, OCT2_ = 6180 nM) inhibit transport by hOCT2 with weak but measurable potency, while showing no inhibition below 10 μM at other uptake 2 transporters examined (Figures 2A and S6-S9).

### Synaptic reuptake inhibitors (SynRIs) are defined by dual inhibition of the vesicular monoamine transporter 2 and a plasma membrane monoamine transporter with comparable inhibitory potencies

The inhibition of both VMAT2 and a plasma membrane monoamine transporter is relatively uncommon in the pharmacopeia. The potent VMAT2 inhibitors such as reserpine or tetrabenazine (TBZ) are selective for VMAT2 versus plasma membrane monoamine transporters (>1000-fold potency differences for reserpine and >70-fold for TBZ, Figure S10),^52^ while potent SERT inhibitors such as imipramine and fluoxetine are selective for SERT over VMAT2 (typically >100-fold, Figure S10).^53–56^ Although there are reports that suggest the possibility of cross-activity between the MATs and VMAT2 for the classical compounds,^55,57^ VMAT2 inhibitors do not block MATs in brain tissue or *in vivo* at concentration exposures that lead to vesicular depletion.^52,58^ We therefore propose a new class of pharmacological agents - termed “synaptic reuptake inhibitors” or “SynRIs” – as agents exerting dual inhibition of VMAT2 and SERT (or the other plasma membrane monoamine transporters including DAT or NET) with comparable inhibitory potencies (within a half log in IC_50_ values, Figures 3, S1-S2, and S4-S5). The dual transporter inhibition profile suggests complex effects at 5-HT synapses and release sites, as well as other monoamine synapses, as noribogaine directly modulates the synaptic vesicular, cytoplasmic, and extracellular pools of 5-HT (Figure 2B). Some of the *in vivo* neurochemical measurements reported previously are consistent with this pharmacological profile (see Introduction and Discussion).

There is a precedent for VMAT2 inhibitors that also block DAT transport with comparable potencies, as exemplified by deoxygenated lobeline analogs.^59^ *S*-amphetamine has comparable potency as a [^3^H] 5-HT releaser via SERT in rat synaptosomes (EC_50_ = 1.8 µM) and uptake blocker at VMAT2 (IC_50_ = 3.3 µM) in crude vesicular rat brain fractions; whereas its releasing potency was reported to be much greater at NET and DAT in synaptosomes (EC_50, NET_ = 7.1 nM and EC_50, DAT_ = 24.8 nM, respectively).^42,60^

### Synthesis of iboga alkaloids and their analogs from natural voacangine

We next examined the effect of substitution in the 1-position and 5-position (indole substituents) on the balance between the VMAT2 and SERT inhibition (we propose a new, more logical iboga system numbering as shown in Figure 3A). These two structural positions are readily modifiable via late-stage functionalization of naturally derived iboga alkaloids (Scheme 1), with the exception of (±)-oxa-noribogaine and (-)-fluoro-ibogamine analogs, which required a total synthesis as reported previously.^23,31,61,62,63^ We do not use *Tabernanthe iboga* materials for analog synthesis for eco-cultural reasons (to avoid overharvesting this resource); instead, we used natural (-)-voacangine obtained by extraction from the root bark of *Voacanga Africana* (Scheme 1).^31,64^ Voacangine was converted to desmethyl voacangine, using ethanethiol and boron tribromide, and its subsequent hydrolysis and decarboxylation yielded noribogaine. We used noribogaine as the starting point to generate a number of readily accessible derivatives, by taking advantage of reactive sites on the indole core.

**Scheme 1.**
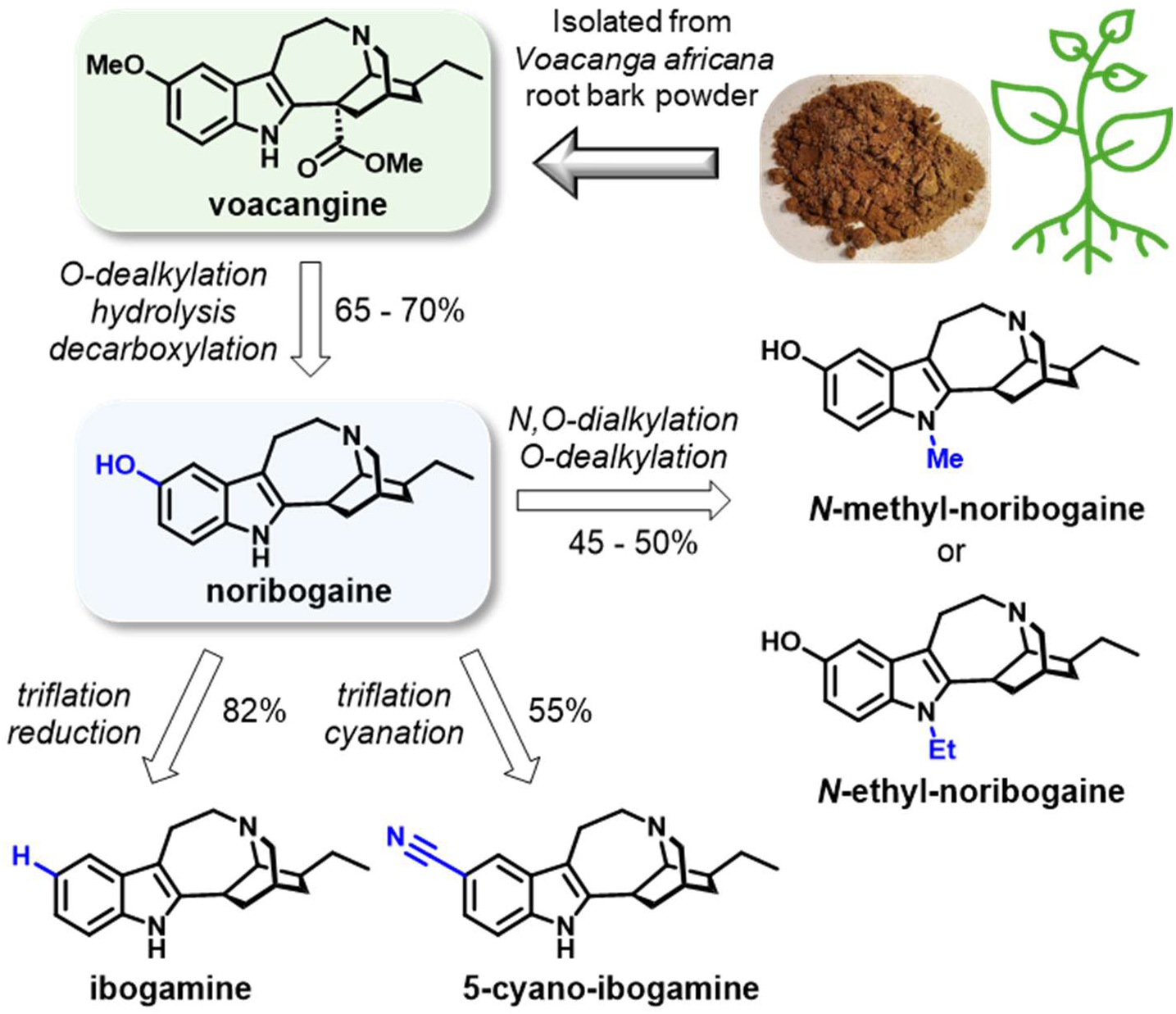
Synthesis of iboga analogs via late-stage functionalization of natural voacangine. Noribogaine, accessible semi-synthetically from voacangine isolated from root bark of *Voacanga Africana*, serves as a versatile intermediate for preparation of analogs with amplified and targeted transporter inhibitory profiles. Detailed synthetic procedures are provided in supplementary information.

The phenol moiety can be derivatized selectively under mild basic conditions to obtain alkoxy analogs (with alkyl halide, carbonate) or the reactive triflate intermediate [by bis(trifluoromethanesulfonyl)aniline, tertiary amine base). 5-Triflate-ibogamine can be further transformed by palladium-mediated hydrogenation into ibogamine,^65^ a substantial component of the total iboga alkaloid content of *Tabernanthe iboga*, but not produced by *Voacanga africana* plants in sufficient quantity to support cost-effective direct isolation. The nitrile group is a prominent feature of the selective SERT inhibitor citalopram, as such we hypothesized that this group could have a dramatic effect on ibogamine’s SERT inhibition. Aryl halides and pseudohalides, like triflates can be transformed into nitriles by a widely used palladium-mediated reaction using a suitable nitrile source (zinc cyanide), carried at elevated temperatures (80 – 160°C). However, we found that a prolonged reaction time and high excess of reagents were required to achieve a desirable level of conversion at a lower temperature range required to limit degradation.

Indole nitrogen is rapidly alkylated in good yields when treated with alkyl halide in dimethylsulfoxide in the presence of potassium hydroxide. Milder alkylation conditions (metal carbonate in hot acetonitrile) can be used to *N*-alkylate more labile analogs, that would not tolerate the harsher KOH/DMSO conditions. For analogs where restoring the free phenol group was desirable, the alkoxy group was cleaved using ethanethiol and boron tribromide.

Oxa-noribogaine was prepared and characterized as a racemate by a total synthesis as reported previously,^31^ while (-)-5- and (-)-6-fluoro-ibogamine analogs were prepared from a chromatographically resolved isoquinuclidine intermediate to match the natural iboga stereochemistry.

### Structure-activity relationship (SAR) of iboga alkaloids at hVMAT2 and hSERT

Removal of the methyl group in the 5-position transforms VMAT2-prefering activity of ibogaine to a balanced inhibitory profile of noribogaine, by increasing the SERT potency 10-fold (Figures 2-3 and S1-S2). Excision of the entire 5-methoxy group of ibogaine, rendering another natural iboga alkaloid, ibogamine, inverts the ratio of the two activities by shifting the potency about 10-fold at each target, giving a SERT-preferring pharmacological profile (IC_50, hSERT_ = 330 nM, IC_50, hVMAT2_ = 3010 nM; Figures 2-3 and S1-S2). In contrast, extending the steric bulk of the methoxy to ethoxy group in the same position, accomplished by synthesizing 5-ethoxyibogamine, affords a potent VMAT2 inhibitor with more than 100-fold selectivity over SERT (IC_50,hVMAT2_ = 80 nM, IC_50,SERT_ = 9030 nM, Figures 3A-C and S1-S2). In further contrast, placing a nitrile group in the 5-position gives 5-cyano-ibogamine, which is a potent SERT inhibitor with high selectivity over VMAT2 (IC_50, hSERT_ = 26 nM, IC_50, hVMAT2_ = 3340 nM, Figures 3A-B and S1-S2). Thus, the 5-position has a strong effect on both transporters and the observed relative inhibitory potencies. The hydroxy or alkoxy group is important for the observed sub-micromolar inhibitory potency at hVMAT2, as the corresponding 5-fluoroibogamine analogs and ibogamines (H in the 5-position) are weak hVMAT2 inhibitors (Figure 3A and Figure S2).

### With respect to hSERT activity, the nitrile and fluorine 5-substituents give potent, low nanomolar hSERT inhibitors (Figure 3A and Figure S1)

*N*-alkylation of the indole nitrogen (1-position) can also have robust SAR effects. For example, *N*-methylation of noribogaine resulted in increasing the potency two- to three-fold at both targets, giving a SynRI ligand, *N*-methyl-noribogaine (IC_50, hVMAT2_ = 59 nM, IC_50, SERT_ = 170 nM, Figures 3A-C and S1-S2). Conversely, *N*-ethyl-noribogaine is a potent hVMAT2 compound with good selectivity over hSERT (IC_50,hVMAT2_ = 70 nM, IC_50,SERT_ = 1860 nM, Figures 3A-C and S1-S2). *N*-methylation of the potent hSERT inhibitor 5-cyano-ibogamine further increases the inhibitory potency at hSERT, yielding a compound with low nanomolar potency at hSERT (IC_50, SERT_ = 5 nM, IC_50, hVMAT2_ = 440 nM; Figures 3A-B and S1-S2). Replacing the nitrogen with oxygen, rendering the oxa-iboga system, as exemplified by oxa-noribogaine (IC_50, SERT_ = 900 nM, IC_50, hVMAT2_ = 670 nM; Figures 3A-C and S1-S2) results in a nearly perfect balance between hSERT and hVMAT2 due to a modest decrease of the hSERT potency versus noribogaine.

We found that hOCT2 has its own SAR trends. Removal of the 5-methoxy group or *N*-methylation of ibogaine, yielding ibogamine (IC_50, hOCT2_ = 2.0 μM) and *N*-methyl-ibogaine (IC_50, hOCT2_ = 2.3 μM), respectively, gives the most potent compounds of the series at hOCT2 in the range of low micromolar inhibitory potencies (Figure S6).

The combined SAR at these two positions enabled us to generate a series of iboga compounds with a broad range of activities, including potent hVMAT2 or hSERT inhibitors, balanced SynRI compounds (Figure 3C), and compounds with varying relative potencies at these two targets. The rest of monoamine transporters were also examined for each of the featured compounds (Figures S1-S2 and S4-S9). The data discussed above was obtained by fluorimetry of the transfected cells using a plate reader. We confirmed these results with epifluorescence and confocal fluorescence microscopy, which showed that the fluorescence signals originate from intracellular localization of the fluorescent substrates, which is dependent on the function of relevant transporters (Figure S3).

### Iboga alkaloids inhibit uptake but do not induce release of serotonin (5-HT) in brain synaptosomes

We next examined the effect of iboga compounds on monoamine transporters natively expressed in rat brain tissue preparations. For this purpose, synaptosomes have been used with much success enabling measurements of both uptake and release of radiolabeled neurotransmitters; namely, tritiated dopamine, [^3^H]DA, norepinephrine, [^3^H]NE, and serotonin, [^3^H]5-HT.^42^ Synaptosomes are prepared by fractionation of brain tissue. Caudate tissue was used for DAT assays whereas whole brain minus caudate and cerebellum was used for NET and SERT assays. Synaptosomes preserve much of the molecular machinery and physiological functions of presynaptic boutons, such as respiration, membrane potential, depolarization-induced exocytosis, neurotransmitter uptake, and pharmacologically induced monoamine efflux.^66^

We examined a set of iboga compounds on [^3^H]5-HT uptake in synaptosomes, in the presence of DAT and NET inhibitors, effectively examining the function of native rat SERT (rSERT, Table 1).^67^ Noribogaine was about 10-fold more potent than ibogaine in this assay, consistent with the literature and the inhibitory potency values determined in the optical assays with hSERT-transfected cells as detailed above (ibogaine: IC_50, rSERT_ = 3037 nM versus IC_50, hSERT_ = 2980 nM; noribogaine: IC_50, rSERT_ = 326 nM versus IC_50, hSERT_ = 280 nM, Tables 1 and Figure S12). Both sets of values are similar to those reported previously in rat synaptosomes (ibogaine: IC_50, rSERT_ = 3150 nM, noribogaine: IC_50, rSERT_ = 330 nM).^17^ Close similarities of the obtained values are noteworthy considering the differences between these two assays. For the other iboga analogs, the values between the synaptosome and fluorescence assays also matched well, except for *N*-ethyl-noribogaine where the inhibition of [^3^H]5-HT uptake in synaptosomes was more than 20-fold more potent than the fluorescent SERT substrate inhibition in transfected cells (IC_50, rSERT_ = 80 nM versus IC_50, hSERT_ = 1860 nM, Table 1B and Figure S12). The reasons for the large difference in potency for this specific compound are unclear; however, we note that *N*-ethyl-noribogaine differs from the rest of tested compounds by its high potency of VMAT2 inhibition. The greater relative potency of *N*-ethyl-noribogaine in synaptosomes versus transfected cells was also found at NET and DAT (Figure S12). This leads to a speculation that the increased potency at MATs may be due to a functional coupling between the VMAT2 and MAT transporters in the native system but not in the singly transfected cell lines, rather than species differences (rat versus human). The source of this discrepancy is unclear.

**Table 1.** Effect of iboga alkaloids on SERT-mediated uptake and release. (A) Iboga analogs inhibit SERT-mediated [^3^H]5-HT uptake (*n* = 3) in rat-derived brain synaptosomes, but do not initiate its release (*n* = 3). (B) Comparison of inhibitory activity determined for iboga analogs in cell-based fluorescent uptake assay (*n* = 4) with values observed in rat brain synaptosomes presented on panel A.

| A) | Compound | [<sup>3</sup>H]5-HT Uptake Inhibition<br>SERT IC<sub>50</sub> (nM) Values | [<sup>3</sup>H]5-HT Release via SERT<br>EC<sub>50</sub> (nM) Values (E<sub>max</sub> %) |
| --- | --- | --- | --- |
|  | ibogaine | 3037.0 ± 291.0 | inactive |
|  | noribogaine | 326.1 ± 47.6 |  |
|  | ibogamine | 554.1 ± 49.0 |  |
|  | N-methyl noribogaine | 13.6 ± 1.2 |  |
|  | N-ethyl-noribogaine | 80.1 ± 7.2 |  |
|  | 5-cyano-ibogamine | 65.2 ± 5.4 |  |
|  | N-methyl-5-cyano-ibogamine | 9.8 ± 0.8 |  |
|  | cocaine | 295.7 ± 29.5 | not tested |
|  | amphetamine | not tested | 3336.0 ± 445.0 (91.0%) |

**Table 1. Effect of iboga alkaloids on SERT-mediated uptake and release.**
| B) | Compound | SERT Transport Inhibition IC<sub>50</sub> Values (nM) |  |
| --- | --- | --- | --- |
|  |  | Uptake of [<sup>3</sup>H]5-HT | Uptake of APP+ |
|  | ibogaine | 3037 | 2980 |
|  | noribogaine | 326 | 280 |
|  | ibogamine | 554 | 330 |
|  | N-methyl noribogaine | 14 | 59 |
|  | N-ethyl-noribogaine | 80 | 1860 |
|  | 5-cyano-ibogamine | 65 | 26 |
|  | N-methyl-5-cyano-ibogamine | 10 | 5 |
|  | cocaine | 296 | not tested |

Generally, transporter substrates differ from non-transported inhibitors by stimulating the efflux of pre-accumulated substrates (“trans-acceleration” in the terminology of W. Stein).^68^ Synaptosomes allow compounds to be assayed for promoting the release of preloaded [^3^H]5-HT, which would be indicative of acting as SERT substrates. A previous study showed that ibogaine did not release [^3^H]5-HT from synaptosomes, but noribogaine was not tested.^69^ Considering the ability of ibogaine and noribogaine to stabilize conformations of SERT distinct from those induced by SSRIs, and their SERT pharmacochaperone activity,^20–22^ we re-examined this question with a wider set of iboga analogs (Table 1 and Figure S12). No [^3^H]5-HT release was observed for ibogaine, noribogaine, or ibogamine; nor for any of the synthetic iboga analogs tested. This contrasts with the robust [^3^H]5-HT release by amphetamine, a drug shown to be a SERT substrate by numerous lines of evidence (positive control, Table 1 and Figure S12).^42^

The ratio of binding to uptake inhibitory constants at SERT is low (∼ 1) which also supports the model where ibogaine and noribogaine are non-transported SERT inhibitors. In contrast, transporter substrates typically exhibit large ratios (10s-100s) due to markedly weaker potency in ligand binding displacement assays relative to the potency obtained in uptake inhibition assays.^17^

### Imaging SERT and VMAT2 inhibition by iboga alkaloids with subcellular resolution in mouse brain

We next set out to examine SERT inhibition by the iboga compounds in acutely sectioned living brain slices in several brain regions containing different parts of serotonin neurons (cell bodies, dendrites, and axons). For this purpose, we used the recently reported molecular probe, SERTlight, which is a selective fluorescent SERT substrate that labels 5-HT neurons in mouse brain with good morphological fidelity in a SERT-dependent manner (Figure 4A).^50^ This imaging probe enables visualization of SERT function in the context of 5-HT neuronal morphology by incubating brain slices with SERTlight (10 μM, 30 min incubation), or injecting SERTlight into the brain in living animals, and acquiring images inside tissue via two-photon fluorescence microscopy. The cell bodies and dendrites were imaged in the dorsal raphe (DR), the location of a majority of 5-HT neuronal cell bodies that project throughout the forebrain (Figure 4A, F). The 5-HT axonal projections appearing as punctate strings were imaged in the substantia nigra pars reticulata (SNpr) and dorsal striatum (DS), regions with relatively high density of 5-HT axons, and somatosensory cortex (SS, or barrel cortex), that has sparse 5-HT axonal innervation (Figure 4A-E). In the 5-HT axons in these regions, citalopram (2 μM, pre- and co-incubation with SERTlight) used as a positive control resulted in complete suppression of SERTlight labeling, consistent with *in situ* axonal SERT inhibition. In contrast, ibogaine (2 μM) had no effect on SERTlight labeling using the same concentration and incubation protocol as for citalopram, consistent with a relatively weak, micromolar inhibitory effect of ibogaine as determined in the cell assays and synaptosomes (Figure 4B, C, E). On the other hand, noribogaine (2 μM) showed a complete inhibition of SERTlight signal in the axons, in line with its sub-micromolar SERT inhibition potency (Figure 4B, C, E, F). Similarly, the potent SERT inhibitor, 5-cyano-noribogaine (2 μM), fully inhibited SERTlight uptake; and a qualitative dose-response study indicated a marked suppression of SERTlight fluorescent labeling at 200 nM concentration of this iboga analog (Figure 4D). These results indicate that noribogaine, and its more potent analog, 5-cyano-noribogaine, can inhibit axonal SERT function in native brain tissue in a concentration range that matches the brain exposures of free drug at behaviorally active doses (> 2 μM, see the pharmacokinetic studies below).

**Figure 4.**
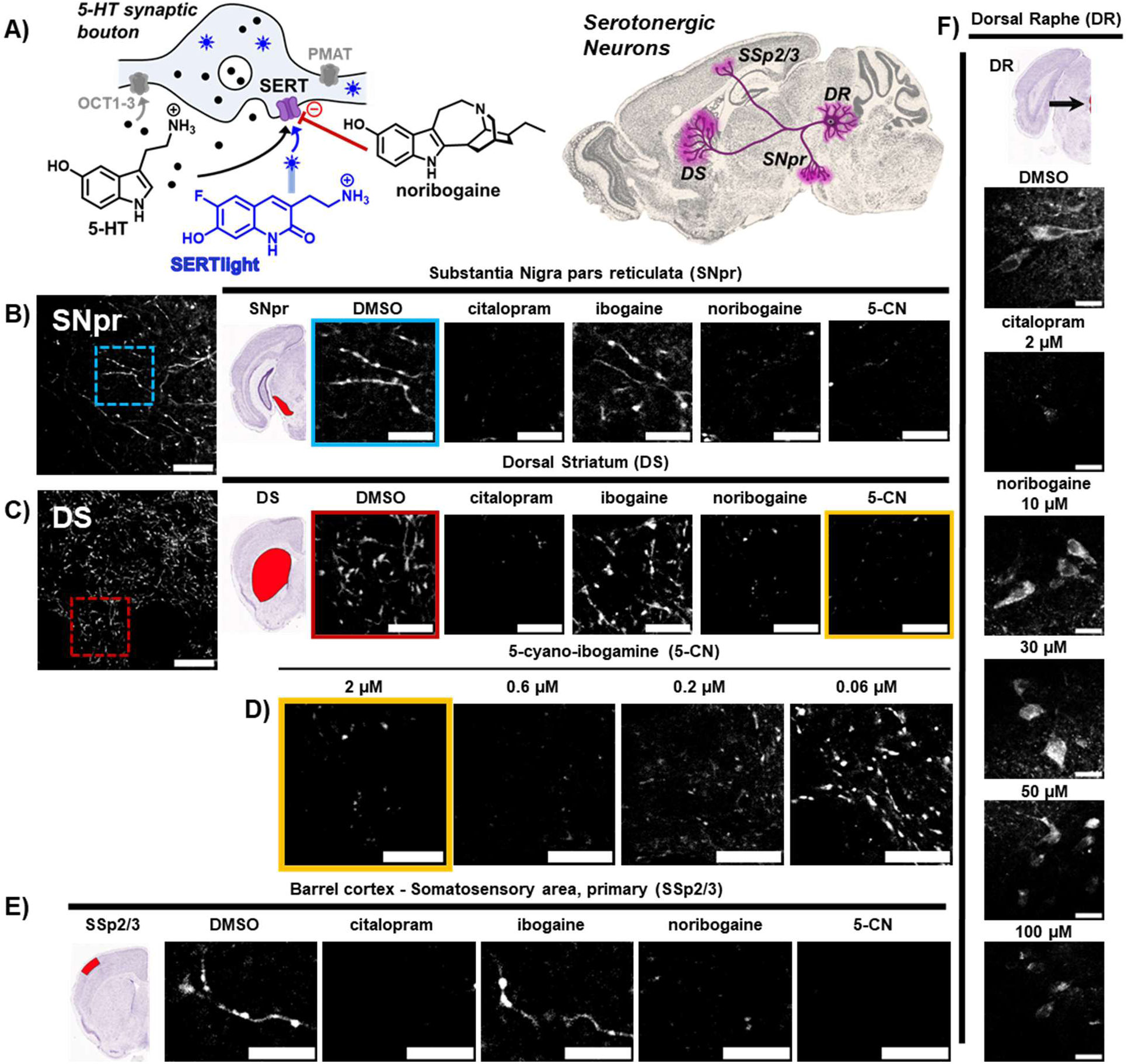
Inhibition of SERT in axons, cell bodies, and dendrites of 5-HT neurons in mouse brain visualized by SERTlight and two-photon imaging. (A) Left panel: A schematic graphic of the concept of the imaging method. A 5-HT axonal bouton and release site is shown featuring SERT in the plasma membrane, which transports 5-HT and SERTlight (as the fluorescent substrate) to yield fluorescently labeled 5-HT neurons. Inhibition of SERT with noribogaine or other iboga analogs diminishes the labeling of 5-HT neurons. Examples of the other transporters examined in this study (OCT2 and PMAT)are also shown. Right panel: A schematic representation of a sagittal slice of mouse brain (Allen Brain Atlas) showing the brain areas chosen for imaging, including dorsal raphe (DR), the seat of 5-HT cell bodies and dendrites; substantia nigra pars reticulata (SNpr) and dorsal striatum (DS), the regions with relatively dense 5-HT axonal innervation; and somatosensory cortex layers 2/3 (SSp2/3), a location with relatively sparse 5-HT axons. (B) Left, zoomed-out microscopy image in SNpr (Scale bar: 20 μm) showing axonal strings and the location of the inset image shown right (Scale bar: 7 μm). In-between the microscopy images, mouse brain atlas image highlighting the SNpr region (Bregma: 1.8 mm, Allen Brain Institute). For the inset images to the right, SERTlight (10 μM for 30 minutes) accumulates in 5-HT axons in WT mice. Co-incubation of acute mouse brain slices with the SERT inhibitor citalopram (2 μM for 30 minutes) and SERTlight (10 μM for 30 minutes) inhibited SERTlight uptake (positive control). Co-incubation of acute mouse brain slices with ibogaine (2 μM for 30 minutes) and SERTlight (10 μM for 30 minutes) did not alter probe uptake into 5-HT axons, but co-incubation with noribogaine (2 μM for 30 minutes) or 5-cyano-ibogamine (2 μM for 30 minutes) ablated SERTlight probe uptake. (C) Left, zoomed-out microscopy image in DS (Scale bar: 20 μm) showing axonal strings and the location of the inset image shown to the right (Scale bar: 7 μm). In-between images, a mouse brain atlas representation highlighting the dorsal striatum (DS) in red (Bregma: -3.3 mm, Allen Brain Institute). For the inset images to the right, the same treatment conditions as in B above. (D) Two-photon images in DS showing the dose-response of co-incubating SERTlight (10 μM for 30 minutes) with 5-cyano-ibogamine. Marked labeling inhibition is observed at 0.2 μM concentration of 5-cyano-ibogamine. (E) Left, mouse brain atlas image highlighting the barrel cortex (SSp2/3) region containing sparse 5-HT axonal projections (Bregma: -1.0 mm, Allen Institute). Images to right (scale bar: 7 μm), the same treatment conditions as in B above. (F) Vertical organization of panels, top: mouse brain atlas image highlighting the small DR region (Bregma: -4.5 mm, Allen Brain Institute). Image below shows SERTlight staining of 5-HT neuronal cell bodies, the larger objects with darker nuclei, and punctate staining of dendrites. From top to bottom, co-incubation of SERTlight with citalopram (positive control) and increasing concentrations of noribogaine (10, 30, 50, and 100 μM) shows weak inhibitory potency of noribogaine, but not citalopram, in this brain region. For all panels, representative images from *n* = 3 mice, three slices per mouse per region.

In the DR, the somatodendritic labeling pattern of SERTlight was completely inhibited by citalopram (2 μM); however, noribogaine showed no effect at 2 μM and 10 μM concentrations, with the latter concentration being five-fold higher than a concentration effecting complete SERTlight labeling suppression in 5-HT axons (Figure 4F). A dose-response study demonstrated that even 30 μM concentration of noribogaine had no apparent effect on the somatodendritic SERT function. A clear inhibition was only achieved at 100 μM concentration of noribogaine (Figure 4F). Thus, in semi-quantitative terms, there is more than 50-fold difference in the inhibitory potency of noribogaine between the axonal and somato-dendritic brain regions. A simple explanation that the cell bodies and dendrites of 5-HT neurons express other transporter(s), which take up SERTlight but are noribogaine-insensitive, was eliminated by showing that: 1) there was no SERTlight somatodendritic labeling in SERT knock-out (SERT-KO) mice, 2) SERTlight was not a substrate for DAT, NET, or any of the uptake 2 transporters.^50^ Instead, our results indicate that noribogaine may have the ability to “read” a different functional status of the SERT molecules in cell bodies versus axons, embodying essentially an “axon-selective” or “brain-region-selective” SERT inhibitor.

We next examined the effect of ibogaine, noribogaine, and 5-cyano-ibogamine on mouse VMAT2 (mVMAT2) function in acute mouse brain slices using the fluorescent VMAT2 substrate, FFN200. We previously demonstrated that FFN200 labels synaptic vesicle clusters in a mVMAT2-dependent manner in the monoaminergic axons in the striatum (Figure 5A).^70^ FFN200 affords a highly dense punctate labeling pattern which is attenuated by VMAT2 inhibitors, such as dihydrotetrabenazine (dhTBZ, 2 μM), (Figure 5b). In addition to a qualitative assessment, the punctate pattern was quantified (see Methods for details) and confirmed large inhibitory effects by ibogaine and noribogaine (each at 2 μM), consistent with their sub-micromolar potency at VMAT2. In comparison, the weaker VMAT2 inhibitor, 5-cyano-ibogamine (2 μM), showed no inhibitory effects at the equivalent concentration. These outcomes support the mechanistic model in which ibogaine and noribogaine directly inhibit VMAT2, an intracellular molecular target expressed on monoamine synaptic vesicles, and thus are expected to limit the synaptic vesicular neurotransmitter pools by drug exposures reached by administering biologically active doses (> 2 μM, after 10 mg/kg, see pharmacokinetics results below).

**Figure 5.**
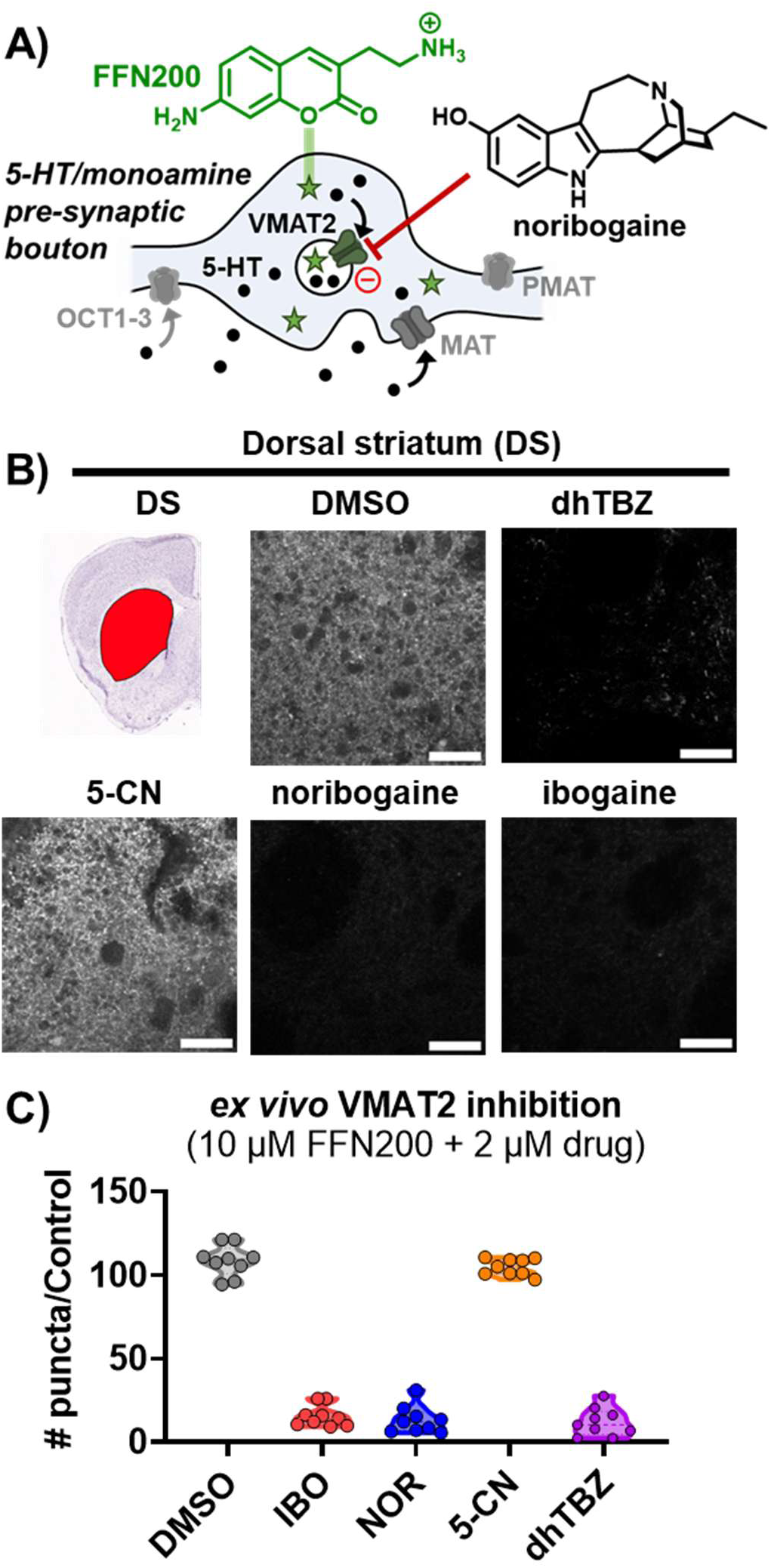
Inhibition of VMAT2 by iboga alkaloids visualized with resolution of individual synaptic vesicle clusters of monoamine axons in the brain. (A) A graphical sketch of the imaging method visualizing VMAT2 transport in the brain. A monoaminergic axonal bouton and release site (in either 5-HT, DA, or NE axons) where intracellular neurotransmitter or FFN200 (as a fluorescent substrate) are transported into the synaptic vesicles. The puncta in the microscopy images represent individual axonal vesicle clusters. (B) Left panel, mouse brain atlas image highlighting the dorsal striatum (DS) region containing high density of DA, but also 5-HT and NE neuronal axonal projections (Bregma: -3.3 mm, Allen Brain Institute). Top middle panel (vehicle, DMSO), incubation of acute mouse brain slices with FFN200 (10 μM for 30 minutes) leads to probe accumulation in VMAT2-expressing vesical clusters in WT mice giving a dense punctate pattern imaged by two-photon microscopy. Top right panel (dhTBZ), co-incubation of acute mouse brain slices with FFN200 (10 μM for 30 minutes) and the VMAT2 inhibitor dhTBZ (2 μM for 30 minutes) inhibited the labeling pattern (positive control). Bottom right and middle panels: co-incubation of FFN200 (10 μM for 30 minutes) with ibogaine (2 μM for 30 minutes) or noribogaine (2 μM for 30 minutes) completely inhibited FFN200 uptake. Bottom left panel: coincubation with 5-cyano-ibogamine (2 μM for 30 minutes) under the same conditions as above had no effect on labeling pattern of FFN200. (C) Quantification of the number of puncta per image under different conditions. Ex./em. for FFN200: 740 nm/460±25 nm. Scale bar for images: 20 μm. Representative images from *n* = 3 mice, three slices per mouse.

The results obtained with SERTlight and FFN200 support a model in which noribogaine modulates both SERT and VMAT2 in the 5-HT presynaptic boutons, as well as VMAT2 in the DA and NE presynaptic boutons and release sites at relevant brain exposures (Figure 2B).

### Ibogaine and noribogaine do not induce catalepsy despite their VMAT2 inhibitory activity

VMAT2 inhibitors at sufficient doses induce catalepsy in humans and animals, characterized by an immobile position and lack of volitional motor behavior, a consequence of dopamine depletion in the striatum and basal ganglia. We used a simple bar test to determine catalepsy in mice where the animals are situated in the instrument in a rearing position with the front paws placed on an elevated bar (Figure 6A). Under vehicle control conditions, animals come off the bar immediately (< 2 – 4 seconds), whereas the cataleptic state is determined by a time threshold (30 – 60 s) of remaining with both paws on the bar.^71,72^ In this manner, catalepsy in mice can be readily differentiated from sedation, which is a non-specific readout of suppression of spontaneous locomotor behavior as measured by placing animals in a novel arena (open field, OF, test, (Figure 6A)).^73^

**Figure 6.**
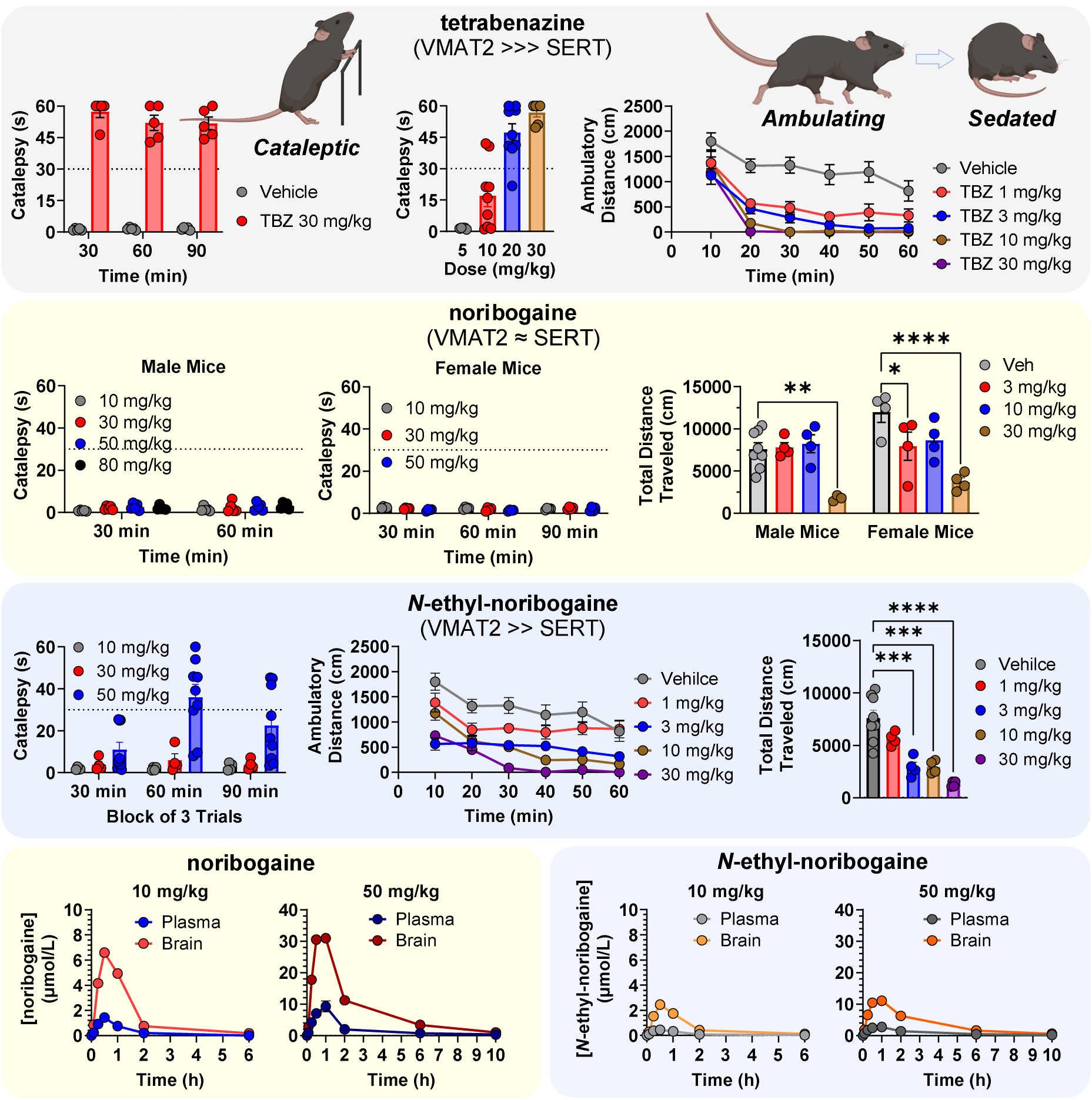
Ibogaine and noribogaine do not show behavioral signs of monoamine depletion. (A) Tetrabenazine acutely induces catalepsy as evidenced by the catalepsy bar test for over 90 min (*n* = 5), with the severity following a dose-response trend (*n* = 5 - 9). This diminishes also the observed locomotor activity in an open-field test (*n* = 4 - 8). (B) Noribogaine administration does not result in catalepsy in male (*n* = 5), nor female mice (*n* = 4), but produce sedation-like behavior resulting in diminished ambulatory behavior (*n* = 4 - 8), presented as total distance traveled over 60 min. (C) High doses of *N*-ethyl-noribogaine induces catalepsy (*n* = 4 - 9) and suppresses locomotor activity in an open field (*n* = 4 - 8). (D) Noribogaine administration rapidly produces high, freely available brain concentration levels. *N*-ethyl-noribogaine is no less brain penetrant, but significantly less available compared to noribogaine due to its high nonspecific brain tissue homogenate binding. Pharmacokinetic data and derived parameters are available in the supplementary information file (Figure S11). Compounds were administered by s.c. injection. Data were analyzed using one-way ANOVA followed by Dunnett’s post hoc test. All values represented as mean ± SEM, ****p < 0.0001, ***p < 0.001, **p < 0.01, *p < 0.05.

We used TBZ, an established selective VMAT2 inhibitor, as a positive control. As expected, TBZ induced both sedation in the open field test and catalepsy in the bar test in a dose-dependent manner (Figure 6A). In contrast, noribogaine did not induce catalepsy even at high doses (30, 50 and 80 mg/kg, s.c.), whereas the locomotion was strongly suppressed already at 30 mg/kg in both male and female mice (Figure 6B). Similarly, ibogaine did not induce catalepsy while suppressing locomotion, but the assessment of catalepsy at higher doses (30 mg/kg and higher) was prevented by strong tremors, a recognized effect of ibogaine and several other natural iboga alkaloids (Figures S13-S15).^74^ After administration of high doses of noribogaine or other iboga compounds, we did not observe a loss of righting reflex (LORR), indicating that the heavy sedation and catalepsy states were not confounded by anesthesia (Figure 7).^75–77^

**Figure 7.**
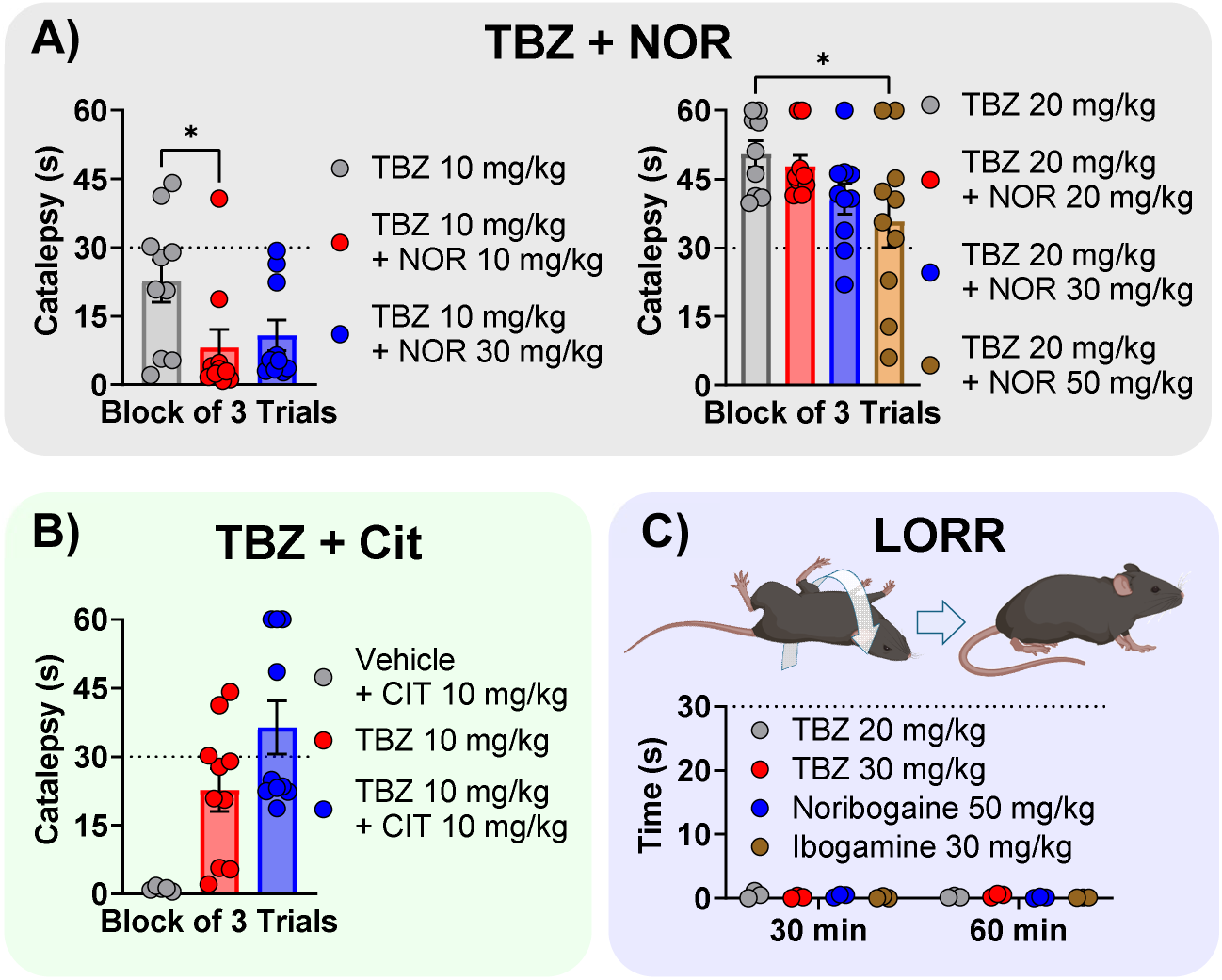
Ibogaine and noribogaine attenuate tetrabenazine-induced catalepsy. (A) Effect of noribogaine administration on TBZ-induced catalepsy at sub-threshold (10 mg/kg) and full effect (20 mg/kg) doses (all groups n = 10). (B) Citalopram potentiated catalepsy effect of TBZ 10 mg/kg (n = 10, except veh+CIT n = 5). (C) Loss of righting reflex (LORR) was assessed one-hour post injection (n = 3). Data analyzed using one-way ANOVA followed by Dunnett’s post hoc test. All values represented as mean ± SEM, *p < 0.05.

The potent VMAT2 inhibitor, *N*-ethyl-noribogaine, also showed a lack of catalepsy up to 30 mg/kg, while this dose completely suppressed locomotion. However, at an even higher dose of 50 mg/kg, catalepsy was induced at the 60-minute mark, while trending toward cataleptic states at the 30 and 90 minutes post drug administration (Figure 6C). The difference in cataleptic effects between noribogaine and *N*-ethyl-noribogaine is not due to a relatively lower brain penetration by noribogaine, as both compounds produce comparable total brain concentrations (Figure S11). However, due to significantly higher nonspecific protein retention of *N*-ethyl-noribogaine (both in plasma and tissue), noribogaine at the same dose shows a nearly three-fold greater estimated free drug exposure in the brain, in terms of both maximum concentration (C_max_) and total drug exposure (AUC, area under the curve, Figure S11) (given 50 mg/kg of each compound, s.c.: C_max, noribogaine_ = 31 μM versus C_max, *N*-Et-noribogaine_ = 11 μM (Figure 6D)). Thus, the greater VMAT2 inhibitory potency of *N*-ethyl-noribogaine (IC_50, hVMAT2_ = 70 nM) versus noribogaine IC_50, hVMAT2_ = 570 nM), as determined *in vitro*, likely underlies the observed differences in catalepsy *in vivo*.

The observation that *N*-ethyl-noribogaine, a potent VMAT2 inhibitor with high brain bioavailability, requires high doses and brain concentrations to induce catalepsy suggests that the iboga matrix pharmacology, the complex pharmacological background of iboga alkaloids, exerts a protective effect against monoamine tissue depletion and catalepsy. To test this hypothesis, we examined the effect of noribogaine on TBZ-induced catalepsy in mice. First, we used a sub-threshold dose of TBZ to be able to observe effects in either direction, namely the suppression or amplification of catalepsy (10 mg/kg TBZ). Indeed, we found that noribogaine alleviated the cataleptic activity of TBZ (Figure 7A). At a higher dose of TBZ (20 mg/kg) that induces full expression of catalepsy, noribogaine still showed a dose-dependent trend, and a statistically significant effect at a high dose (50 mg/kg), toward attenuating the TBZ-induced cataleptic effects. Ibogaine had a similar attenuating effect even at low dose (3 mg/kg, Figure S13). Thus, the catalepsy protection hypothesis of iboga compounds is supported by three points: 1) noribogaine does not induce catalepsy by itself even at high non-toxic doses, 2) noribogaine and ibogaine attenuates TBZ-induced catalepsy, and 3) *N*-ethyl-noribogaine (a potent VMAT2 blocker) only induces catalepsy at high doses and brain concentrations (Figure S14). For comparison, we also examined the effect of a selective SERT inhibitor (citalopram, 10 mg/kg, S.C.) on a sub-threshold TBZ dose, which showed an opposite effect to that of noribogaine, leading to amplification of TBZ’s cataleptic effects (Figure 7B). This observation is consistent with previous reports in rodents in which SSRIs increased the catalepsy and other motor effects such as oral tremors induced by TBZ or haloperidol;^71,78^ and humans in which SSRIs have been reported to exacerbate the motor deficits in a segment of Parkinson’s patients.^79^ Our results indicate that the SERT blockade and transient elevation of 5-HT induced by noribogaine in the NAc and striatum cannot explain its catalepsy mitigating effects. Instead, we propose that there is a functional interplay between VMAT2 inhibition and the iboga matrix pharmacology that moderates the monoamine levels and or monoamine receptor signaling *in vivo*. Previous neurochemistry studies showed that ibogaine and noribogaine induced a transient decrease of extracellular dopamine concentration in the striatum, as well as a strong tissue reduction of dopamine throughout the brain.^20^ However, at these doses, ibogaine (e.g., 40 mg/kg) did not induce a complete dopamine depletion and catalepsy. To our knowledge, catalepsy had not been previously described for ibogaine or noribogaine in rodents. In humans, high doses of ibogaine (> 10 mg/kg, P.O.) can induce ataxia, but not catalepsy.^18^ Anecdotally, very high doses of ibogaine (> 20 mg/kg) can lead to anesthesia-like states, but catalepsy has not been reported.^8^

## Discussion

In this report, we examine the pharmacological profile of iboga alkaloids at monoamine neurotransmitter transporters, with a focus on the parent compound, ibogaine, its main metabolite, noribogaine, and a small series of iboga analogs. We find that ibogaine and noribogaine exhibit a complex monoamine modulatory profile by inhibiting three or more transporters within a pharmacologically relevant concentration range.

Specifically, we demonstrated that ibogaine and noribogaine inhibit the transport function of the vesicular transporter VMAT2 with sub-micromolar potency, using a fluorescent VMAT2 substrate (FFN206) in cell-based assays. We also demonstrated the inhibition of VMAT2 by ibogaine and noribogaine in synaptic vesicle clusters of intact monoamine axons in mouse brain, using two-photon microscopy imaging. Thus, we provide evidence for VMAT2 transport inhibition by iboga alkaloids and the potency range of this effect (sub-micromolar to low micromolar, Figure 5) in two experimental systems. Ibogaine (IC_50, hVMAT2_ = 390 nM) and noribogaine (IC_50, hVMAT2_ = 570 nM) have similar inhibitory potency at VMAT2. We did not elucidate the exact mechanism of this inhibitory effect, as it could result from the iboga alkaloids acting as 1) VMAT2 non-substrate inhibitors, 2) substrate inhibitors, or 3) lipophilic bases that diminish the vesicular pH gradient. The third option is less likely for ibogaine and noribogaine since these two alkaloids possess markedly different lipophilicity while exerting comparable potencies on the uptake of VMAT2 fluorescent substrates. However, the base effect can contribute to the greater potency of more lipophilic *N*-ethyl-noribogaine and 10-ethoxy-ibogamine, in a similar manner as proposed for amphetamine, which functions, at least in part, as a lipophilic base at synaptic vesicles.^80^ This question will be addressed in future studies.

We also examined the effect of iboga compounds on two types of monoamine plasma membrane transporters, the uptake 1 and uptake 2 transporters. In line with previous results reported in the literature,^20^ we show that noribogaine is about 10-fold more potent than ibogaine at inhibiting hSERT as determined using the SERT fluorescent substrate APP+ in cell-based assays. Noribogaine has a comparable inhibitory potency at hSERT (IC_50, hSERT_ = 280 nM) and hVMAT2 (IC_50, hSERT_ = 570 nM, Figures 3 and S1), exhibiting an uncommon pharmacological profile of dual inhibition of these two critical transporters at 5-HT synapses and release sites. We proposed the term “synaptic reuptake inhibitors” or “SynRIs” to highlight this interesting mechanistic feature and contrast it to the well-known VMAT2 inhibitors and SSRIs (Figures 3 and S10). The inhibition of OCT2, identified here as a novel molecular target of iboga alkaloids, may be pharmacologically relevant *in vivo,* considering the estimated free brain concentrations of noribogaine based on the pharmacokinetic studies reported here (C_max_ ∼ 6.6 μM after 10 mg/kg, and 31 μM after 50 mg/kg, s.c., Figures 6 and S11), and previous reports in humans (C_max_ ∼ low micromolar after 10 mg/kg of oral ibogaine in opioid-dependent individuals).^18^ OCT2 certainly needs to be entered into the iboga matrix of known targets, particularly for SAR studies with new analogs.^81^

The presented SAR study showed that the relative potencies at SERT and VMAT2 can be tuned over a broad range, identifying analogs with a balanced SynRI profile (e.g., noribogaine, oxa-noribogaine), dominant VMAT2 (*N*-ethyl-noribogaine), and dominant SERT (5-cyano-ibogamine) (Figures 3 and S1-S2). We also determined that none of the iboga compounds - with varying SERT inhibitory potency - induce release of [^3^H] 5-HT in synaptosomes, confirming the non-substrate inhibitory effect at SERT, consistent with a previous report on ibogaine (Figure S12).^69^ The structural dimensions of the iboga compounds, in which the aliphatic amino group of the tryptamine unit is embedded in the bulky isoquinuclidine ring system, are too large for the transport to take place through SERT on a reasonable time scale, in contrast to the smaller known SERT substrates and 5-HT releasers. However, a recent report claimed a partial [^3^H] 5-HT release (E_max_∼30%) induced by noribogaine in SERT-transfected cells.^63^ With regard to *in vivo* neurochemistry, although there is a single report claiming a 5-HT release induced by ibogaine in nucleus accumbens,^82^ other studies described results consistent with a SERT non-substrate inhibition profile.^17,20^

We further probed the effect of iboga alkaloids on SERT transport in acute mouse brain tissue with subcellular spatial resolution enabled by the state-of-the-art imaging probe, SERTlight, and two-photon microscopy.^50^ While noribogaine (2 μM) completely inhibited the axonal SERT in three different brain regions, as expected, it was far less potent at inhibiting SERT in 5-HT neuronal cell bodies and dendritic compartments (> 50-fold difference; Figure 4). While this finding was surprising, there is a precedent for different binding potencies of noribogaine at VMAT2 in distinct human brain regions (IC_50, binding, striatum_ = 29.5 μM versus IC_50, binding, cortex_ = 5 μM).^48^ It has been previously demonstrated that ibogaine is an atypical SERT inhibitor (non-competitive inhibition) that binds and stabilizes SERT in inward-open conformational states, in contrast to cocaine-based blockers or SSRIs that bias the SERT conformational dynamics toward outward-facing conformations.^83,84^ It has been proposed that the pharmaco-chaperone effect of iboga compounds at SERT (along with DAT and likely NET) is related to the stabilization of inward-open conformational states sampled by the transporter in its folding and trafficking pathway.^85^ Inspired by the ibogaine’s mode of action, more potent SERT pharmacochaperones were generated and conformation-selective SERT ligands were developed via computational methods.^22,86^ On this background of rich iboga SERT pharmacology, we speculate that noribogaine “detects” different SERT conformational or functional states in the brain, which may be related to post-translational modifications and or complexation dynamics with ancillary proteins, transporter-associated chaperones, or cytoskeleton proteins, as part of the local membrane composition and cellular functional states.^87,88^

Considering the findings of this study and prior literature, a complex picture of monoamine neurotransmission modulation by ibogaine and noribogaine emerges (Figure 2). The iboga compounds inhibit monoamine transporters known to play critical roles in monoamine neurotransmission, including the synaptic vesicle transporter VMAT2, plasma membrane monoamine transporters of uptake 1 family (most notably SERT), and the uptake 2 transporter OCT2, expressed on neuronal and glial cells. The dual or multiple site modulation predicts complex, non-linear effects on neurotransmission, especially when considering functional couplings between the different neurotransmitter pools (i.e. vesicular, cytoplasmic, and extracellular) and the complex mutual interactions between 5-HT and catecholamines.^89–91^

The updated monoamine transporter profile of the iboga compounds provides a new explanatory model for the questions posed previously based on the extensive body of neurochemical literature: namely, why do iboga compounds have such a profound effect on dopamine metabolism (a strong dopamine decrease and metabolite DOPAC increase in total tissue content in most brain regions), while the effect on 5-HT metabolism is modest (no or small decrease in 5-HT and a modest decrease of metabolite 5-HIAA).^20^ The direct inhibition of VMAT2 has an important role in the explanatory hypothesis; while ibogaine and noribogaine act as VMAT2 transport inhibitors *in vitro* and *in vivo*, they do not appear to inhibit DAT *in vivo* (*in vitro* inhibition potency at DAT is more than one log unit weaker compared to VMAT2; Figure S4). Thus, the effect of the iboga compounds on dopamine metabolism and neurotransmission is dominated by the VMAT2 inhibitory effect and largely the absence of DAT inhibition. This model is also consistent with the effect of ibogaine and noribogaine on extracellular dopamine: generally, a decrease in striatum, and no effect or a decrease in NAc, as well as attenuation of cocaine-induced dopamine increase in NAc and striatum.^20,92^ The lack of DAT inhibition by iboga compounds *in vivo*, and VMAT2-mediated depletion of vesicular dopamine pools explains the behavioral effects in which ibogaine decreases the ambulatory stimulation by cocaine, an effect dependent on the vesicular release of dopamine via exocytosis.^20^ Thus, ibogaine and noribogaine as VMAT2 inhibitors limit the accumulation of DA into synaptic vesicles, resulting in increased metabolism of dopamine and the neurochemical signature of VMAT2 inhibitors (decreased tissue dopamine and increased dopamine metabolites, Figure 5).

In contrast, at 5-HT axonal release sites, noribogaine acts as a SynRI, a dual inhibitor of VMAT2 and SERT. In this manner, the re-uptake of 5-HT released via activity-dependent exocytosis is limited by SERT inhibition (and potentially OCT2 inhibition), which protects 5-HT from extensive metabolism within 5-HT neurons. VMAT2 inhibition likely contributes to the temporal profile and firing pattern of 5-HT transmission (transient increase of extracellular 5-HT in NAc and striatum) post ibogaine or noribogaine administration (Figure 5).^58^

We propose that the complexity of iboga’s modulation of monoamine neurotransmission provides clues and insights about the global picture of the iboga matrix pharmacology. The present results do not suggest non-discriminate inhibition or activation of many random targets by iboga substances, but rather modulation of a combination of molecular targets that are functionally coupled in critical and highly tuned processes, such as monoamine neurotransmission (e.g., SERT/OCT2 and VMAT2, Figure 2). Furthermore, the effects at certain individual targets are atypical revealing higher order molecular functions. The prime exhibit is SERT, a pivotal element of serotonergic transmission, in which several modes of action of the iboga compounds have been reported: including the acute inhibition of SERT via distinct conformational states, pharmaco-chaperone effect (the latter effects may modulate the long-term function of SERT), and now the axon specific inhibitory effects (which may lead to targeted 5-HT circuit modulation). Hence, we extrapolate the existing evidence and speculate that the iboga matrix pharmacology may have a deep sophisticated logic based on unique molecular effects at specific targets and complex combinations of molecular targets and interventions.

A previous study reported displacement of tritiated vesamicol by ibogaine in human brain (IC_50_ ≤ 10 μM), suggesting inhibition of the vesicular acetylcholine transporter (VAChT),^48^ and hence, a likely effect on the vesicular pools and neurotransmission of acetylcholine. This effect – coupled with the well-established action of iboga compounds as antagonists of several nicotinic acetylcholine receptors (nAChRs) – indicates multi-pronged modulation of the cholinergic neurotransmission.^93–96^ The inhibition of NMDAR by the iboga compounds as channel blockers is also well documented and adds the glutamate neurotransmission to the list of critical neurotransmission posts modulated by the iboga alkaloids.^19^ Finally, we reported previously that iboga compounds potentiate kinase signaling pathways activated by the fibroblast growth factor receptors (FGFRs), suggesting modulation of downstream effectors of receptor tyrosine kinases and neurotrophic factors, together with the reported re-casting of neurotrophin expression levels.^23,30^

Based on these observations and extrapolations, *we put forth a theory that predicts that the molecular targets or signaling processes reported for the iboga compounds represent a fragment or a preview of the full pharmacological profile of unprecedented complexity – which we termed “matrix pharmacology”*. We defined this pharmacological category as modulation of multitudes of molecular targets (often targets that are functionally coupled), via multiple modes of action, and via weak potency molecular interactions that permeate the entire information and bioenergy processing matrix. We also propose that this new concept, or a new pharmacological category, is essential for the interpretation and re-interpretation of the experimental results in the iboga space. For example, selective single-target perturbations ought not to be considered in isolation, but in context of the iboga matrix pharmacology. This is a challenging task that will ultimately require computational models, to interpret and potentially predict experimental outcomes of modifications of the iboga matrix pharmacology.

The iboga analogs with a single dominant target are useful tools for stepwise probing and deciphering the iboga matrix mechanisms. We recently introduced the oxa-iboga compounds (benzofuran iboga analogs) that have KOR as the dominant target (greater than one log more potent at KOR versus at other examined targets). These analogs show atypical signaling and behavioral characteristics (e.g. lack of aversion or pro-depressive effects in mice) in contrast to standard KOR agonists, which we ascribed, at least in part, to the interactions of KOR with the iboga matrix pharmacology.^31^ In the present article, we introduce additional examples of such iboga analogs with one or two dominant targets: for example, 5-cyano-ibogamine as a potent SERT inhibitor, and *N*-ethyl-noribogaine as an analog with dominant VMAT2 (based on the human receptor profile, or dominant VMAT2/SERT activity based on rat synaptosomes, Figures 3 and S1-S2 and Table 1).

In this context, the conceptual framework of matrix pharmacology may be useful for explaining why ibogaine or noribogaine do not induce catalepsy even at high doses, in contrast to the standard VMAT2 inhibitors. To this end, we examined the effect of varying VMAT2 potency within the iboga system on mouse behavior. We confirmed that catalepsy can be readily induced in mice by TBZ, a potent VMAT2 inhibitor, as determined by using a bar test (Figure 6). Unlike the TBZ positive control, noribogaine did not show any catalepsy even at very high doses (50 and 80 mg/kg, s.c.) where the 50 mg/kg dose resulted in > 20 μM estimated maximum concentration of free drug in the brain (Figure 6D), which is 10-times greater than the concentration effecting complete inhibition of VMAT2 transport in striatal brain slices (2 μM, Figure 5). Examining *N*-ethyl-noribogaine, a compound with high VMAT2 potency, catalepsy was not induced by 10 and 30 mg/kg doses, but only by the top administered dose of 50 mg/kg (Figure S14). Remarkably, a nanomolar VMAT2 inhibitor, *N*-ethyl-noribogaine (IC_50, hVMAT2_ = 70 nM *in vitro*), did not induce catalepsy at the estimated maximum brain exposure of 7,000 nM of free drug (30 mg/kg dose). The SynRI profile, the dual VMAT2-SERT inhibition cannot explain these findings as citalopram had the opposite effect, increasing the catalepsy of TBZ (Figure 7). We therefore propose that the background iboga matrix pharmacology mitigates or integrates the consequences of the VMAT2 inhibition effect. This hypothesis was further supported by demonstrating that noribogaine attenuated the cataleptic effects of TBZ.

The effect of VMAT2 inhibitors on dopamine levels is well established, as discussed above, leading to dose-dependent dopamine depletion and catalepsy.^97^ Simple explanations based on known effects that mitigate those of TBZ, such as monoamine oxidases inhibition, DAT blockade, or direct dopamine receptor agonism do not apply in this case, as ibogaine and noribogaine also buffer the opposite effect of a transient dopamine increase induced by cocaine (DAT inhibitor) or opioids (indirect circuit mechanism).^20,98^ Hence, evidence points to a complex multi-factorial mechanism that restrains both neurochemical extremes – high and low dopamine levels – and the behavioral consequences as demonstrated by attenuating the effects of both DAT and VMAT2 inhibition. Our discussion here focuses on the acute (drug-on) neurochemical and behavioral effects, which likely shape the long-term sequels of iboga compounds.^30^ While much work lies ahead on the path toward elucidation of the full complexity and sophistication of iboga matrix pharmacology, the observed mitigation of behavioral consequences of VMAT2 inhibition is an important example.

## Conclusions

In summary, the present study brings two major contributions. First, we provide a comprehensive map of monoamine transporters as primary pharmacological targets of the iboga alkaloids. For this task, we used a combination of *in vitro* and *ex vivo* experimental methods, including the state-of-the-art optical imaging of transporter function in brain tissue, to provide the first direct evidence for VMAT2 and OCT2 inhibition. Second, we propose the concept of iboga matrix pharmacology to advance the understanding of molecular mechanisms, interpretation of results including the therapeutic and psycho-spiritual effects, and development of novel iboga analogs. Our results support the proposed theory that ibogaine and iboga alkaloids are molecular entities operating via a novel, unprecedented mechanism that likely underlies their remarkable therapeutic and healing effects.

In the reports of past ethnographers and ethnobotanists, we find descriptions of a spectrum of effects induced by consumption of the iboga root bark by the indigenous groups in Gabon; for example, central stimulant (at low doses) or intense psychedelic and visionary (at high doses) activities. Iboga was characterized as a substance with a major social impact, serving as a central, unifying agent to local communities.^2,99^ They speak of iboga’s ability to transform one’s subjective views of self and the world, release individuals and social groups from the trauma and difficult living circumstances of post colonialism, and restore clarity and calm amidst a rapidly changing world and culture around them – in essence describing, in our interpretation, a unique psycho-social restorative “technology”. More recently and largely outside Africa, reports suggest profound therapeutic and healing capabilities of ibogaine across therapeutic indications. These include substance use disorders (with opioid, cocaine and alcohol), depression and anxiety, PTSD, mTBI, and potentially chronic traumatic encephalopathy (CTE).^3,4,14,15,100,101^ The recent narratives from volunteers across the world, recounting their subjective experiences during and post ibogaine treatment, are in conceptual themes similar to the conclusions of the ethnographic studies focused on the African indigenous communities carried out in the 1950-1960’s that describe personality reformatting, worldview resetting, and psychosocial restorative effects.^8,102^ If the past and contemporary reports are confirmed in rigorous studies and safe use of ibogaine and iboga analogs, then these substances would constitute a unique class of therapeutics and molecular technologies. The present study adds a few new cornerstones to the molecular puzzle of the iboga pharmacology and puts forth a conceptual framework for the mechanistic complexity it presents.

## Associated Content

### Supporting Information

The Supporting Information is available free of charge at the following link.

Experimental details including *in vitro* functional assays, rat brain synaptosome radioligand uptake and release, and behavioral catalepsy and locomotor activity evaluation protocols, data analysis, and results, synthetic schemes and protocols, and NMR spectra.

## Author Information

### Corresponding Authors

**Dalibor Sames** - Department of Chemistry and The Zuckerman Mind Brain Behavior Institute, Columbia University, New York, New York 10027, United States;

### Authors

**Christopher Hwu** - Department of Chemistry, Columbia University, New York, New York 10027, United States.

**Václav Havel** - Department of Chemistry, Columbia University, New York, New York 10027, United States.

**Xavier Westergaard** - Department of Biological Sciences, Columbia University, New York, New York 10027, United States;

Department of Psychiatry, Columbia University Irving Medical Center, New York, New York 10032, United States.

**Adriana M. Mendieta** - Department of Chemistry, Columbia University, New York, New York 10027, United States.

**Inis C. Serrano** - Department of Chemistry, Columbia University, New York, New York 10027, United States.

**Jennifer Hwu** - Department of Chemistry, Columbia University, New York, New York 10027, United States.

**Donna Walther** - Designer Drug Research Unit, Intramural Research Program, National Institute on Drug Abuse, National Institutes of Health, Baltimore, Maryland 21224, United States.

**David Lankri** - Department of Chemistry, Columbia University, New York, New York 10027, United States.

**Tim Luca Selinger** - Department of Chemistry, Columbia University, New York, New York 10027, United States.

**Keer He** - Department of Chemistry, Columbia University, New York, New York 10027, United States; orcid.org/0009-0005-2006-4970.

**Rose Liu** - Department of Biology, Barnard College, New York, New York 10027, United States.

**Tyler P. Shern** - Department of Chemistry, Columbia University, New York, New York 10027, United States.

**Steven Sun** - Department of Chemistry, Columbia University, New York, New York 10027, United States.

**Boxuan Ma** - Department of Biological Sciences, Columbia University, New York, New York 10027, United States.

**Bruno González** - Department of Chemistry, Columbia University, New York, New York 10027, United States.

**Hannah J. Goodman** - Department of Chemistry, Columbia University, New York, New York 10027, United States.

**Mark S. Sonders** - Division of Molecular Therapeutics, New York State Psychiatric Institute, New York, New York 10032, United States.

**Michael H. Baumann** - Designer Drug Research Unit, Intramural Research Program, National Institute on Drug Abuse, National Institutes of Health, Baltimore, Maryland 21224, United States.

**Ignacio Carrera** - Laboratorio de Síntesis Orgánica, Departamento de Química Orgánica, Facultad de Química, Universidad de la República, Montevideo, Uruguay 11200.

**David Sulzer** - Department of Psychiatry, Department of Neurology, Department of Molecular Pharmacology and Therapeutics, Columbia University Irving Medical Center, New York, New York 10032, United States;

Division of Molecular Therapeutics, New York State Psychiatric Institute, New York, New York 10032, United States.

Complete contact information is available at the following link.

### Author Contributions

^†^ C.H., V.H., and X.W. contributed equally to this work.

### Notes

A. D. Sames, V.H., C.H., D.L., I.C.S and B.G. are inventors on iboga-related patents. D. Sames is a co-founder of Gilgamesh Pharmaceuticals. The other authors declare no competing financial interest.

## Acknowledgments

This work was supported by The G Harold & Leila Y Mathers Charitable Foundation (MF- 2107-01880, Iboga X Project, D.Sames. and V.H.), and the National Institute on Drug Abuse (NIDA) (R01-DA050613 to D. Sames) of the National Institutes of Health (NIH), and the NIDA Intramural Research Program (NIDA ZIADA00052-17, M.H.B.). We would like to thank the following philanthropists for the timely support of the Sames iboga research program: Monica Winsor (Trustee of the William H. Donner Foundation), Jeffrey C. Walker (Walker Family Foundation), Austin Hearst (The Austin & Gabriela Hearst Foundation), Scott and Kristin Edmondson (Scott and Kristin Edmondson Charitable Gift Fund), Zaki Manian, Joshua Mailman (The Joshua Mailman Foundation), and an anonymous funder. This work was also supported in part by the National Institutes of Mental Health (NIMH) of the National Institutes of Health (NIH), grant R01MH108186 (awarded to D. Sames and D. Sulzer), and R01DA07418 and the Freedom Together Foundation (awarded to D. Sulzer). Research reported in this publication was supported by the Office of The Director, National Institutes of Health of the National Institutes of Health under Award Number S10OD026749. The content is solely the responsibility of the authors and does not necessarily represent the official views of the National Institutes of Health. The authors thank Dr. Gary W. Miller and Mr. Joshua M. Bradner of the Department of Environmental Health Sciences of Columbia University Mailman School of Public Health for their gift of hVMAT2-HEK cell cultures that were instrumental to this work, and Dr. Daniel Wacker and Ms. Audrey Warren of the Departments of Pharmacological Sciences, Neuroscience, and Genetics and Genomic Sciences of Icahn School of Medicine at Mount Sinai for their gift of hOCT1-3-HEK and hPMAT-HEK cell cultures.

Illustrations presented on Figures 6, 7, S13, S14, and S15 were created from icons obtained from Biorender.com (Publication License *GM27SS5YWS)*.

