## Supplementary Information for "Deciphering Ibogaine’s Matrix Pharmacology: Multiple Transporter Modulation at Serotonin Synapses"

### Table of Contents


**A. Supplementary Figures Referenced in the Main Text.** 3

**B. Synthesis.** 30

1. Materials and Methods. 30
2. Synthetic Protocols. 30

**C. NMR and MS Spectra.** 45

**D. Materials and Methods.** 85

1. Cell Culture Maintenance and Experimental Preparations. 85
2. Determination of $IC_{50}$ Metrics of Ibogaine Derivatives at hDAT, hNET, and hSERT Through APP+ Uptake Blockage. 86
3. Determination of $IC_{50}$ Metrics of Ibogaine Derivatives at hVMAT2 Through FFN206 Uptake Blockage. 86
4. Determination of $IC_{50}$ Metrics of Ibogaine Derivatives at hOCT1 and hPMAT Through APP+ Uptake Blockage. 87
5. Determination of $IC_{50}$ Metrics of Ibogaine Derivatives at hOCT2 and hOCT3 Through ASP+ Uptake Blockage. 87
6. Determination of $IC_{50}$ Metrics of Ibogaine Derivatives at hDAT, hNET, and hSERT Through Tritiated Monoamine Uptake Blockage Using Rat Brain Synaptosomes. 87
7. Determination of $EC_{50}$ Metrics of Ibogaine Derivatives at hDAT, hNET, and hSERT Through Tritiated 5-HT Release By SERT Using Rat Brain Synaptosomes. 88
8. One-Photon Epifluorescence Microscopy Imaging of Respective Uptake in hSERT-HEK and hVMAT2-HEK Cell Cultures. 88
9. One-Photon Confocal Microscopy Imaging of Fluorescence Uptake in hSERT-HEK and hVMAT2-HEK Cells. 89
10. Determination of SERT Inhibition by Ibogaine Derivatives in Acute Murine Brain Slices. 89
11. Determination of VMAT2 Inhibition by Ibogaine Derivatives in Acute Murine Brain Slices. 90
12. Determination of Pharmacokinetic Parameters in Mice (Bienta Enamine Biological Services). 91
13. Animal Behavioral Studies and Usage. 93

**E. References.** 94

### A. Supplementary Figures Referenced in the Main Text.

**a**

| Name | hSERT $IC_{50}$ Values ± SEM (nM) |
|---|---|
| Imipramine | 6.5 ± 1.5 |
| Ibogaine | 2980 ± 660 |
| Noribogaine | 280 ± 40 |
| Ibogamine | 330 ± 40 |
| *N*-methyl-ibogaine | 1520 ± 130 |
| *N*-methyl-noribogaine | 59 ± 11 |
| *N*-ethyl-noribogaine | 1860 ± 200 |
| *N*-methyl-ibogamine | 80 ± 9 |
| 5-cyano-ibogamine | 26 ± 4 |
| *N*-methyl-5-cyano-ibogamine | 5.1 ± 0.8 |
| *N*-ethyl-5-cyano-ibogamine | 61 ± 11 |
| *N*-methyl-5-fluoro-ibogamine | 61 ± 13 |
| *N*-ethyl-5-fluoro-ibogamine | 36 ± 8 |
| *N*-methyl-6-fluoro-ibogamine | 510 ± 72 |
| *N*-ethyl-6-fluoro-ibogamine | 6.6 ± 1.7 |
| 5-ethoxy-ibogamine | 9030 ± 1100 |
| 5-fluoro-ibogamine | 30 ± 8 |
| 6-fluoro-ibogamine | 77 ± 19 |

**b**

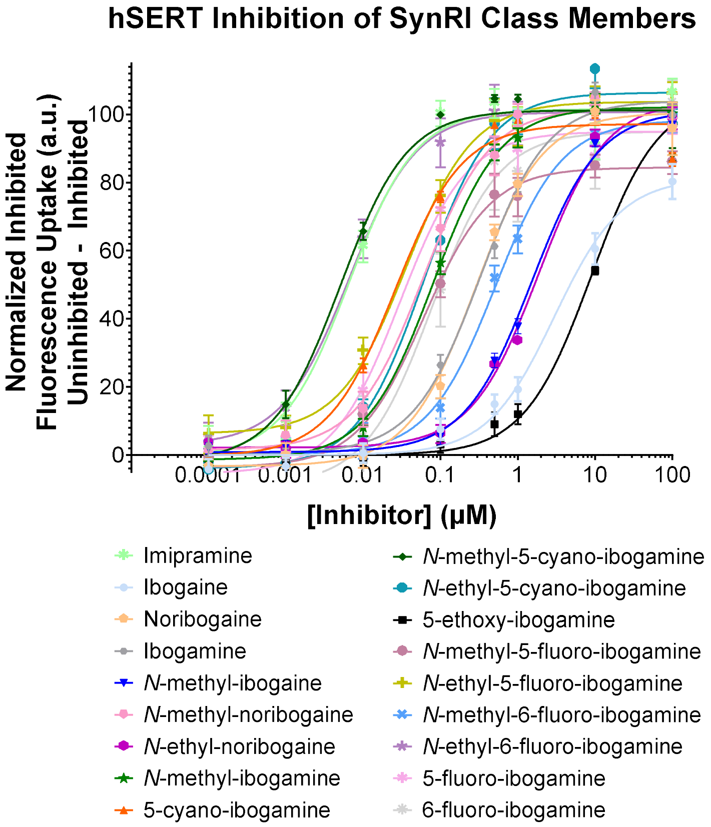

**Figure S1.** hSERT $IC_{50}$ values ± SEM for SynRI iboga alkaloids. (a) Quantification of $IC_{50}$ values of imipramine (a potent SERT inhibitor)[1] and SynRI iboga alkaloids at hSERT using APP+ (final concentration: 1.1 μM)[2] as the substrate for respective functional inhibition experiments. Potent iboga compounds are highlighted in light red. (b) Graphical data is presented as normalized inhibited fluorescence uptake (uninhibited – inhibited) ± SEM and derived from four separate experiments.

**a**

| Name | hVMAT2 $IC_{50}$ Values ± SEM (nM) |
| --- | --- |
| Tetrabenazine | 52 ± 6 |
| Ibogaine | 390 ± 60 |
| Noribogaine | 570 ± 80 |
| Ibogamine | 3010 ± 550 |
| *N*-methyl-ibogaine | 740 ± 110 |
| *N*-methyl-noribogaine | 170 ± 30 |
| *N*-ethyl-noribogaine | 70 ± 10 |
| *N*-methyl-ibogamine | 1470 ± 160 |
| 5-cyano-ibogamine | 3340 ± 430 |
| *N*-methyl-5-cyano-ibogamine | 440 ± 50 |
| *N*-ethyl-5-cyano-ibogamine | 600 ± 95 |
| *N*-methyl-5-fluoro-ibogamine | 5060 ± 1580 |
| *N*-ethyl-5-fluoro-ibogamine | 1720 ± 420 |
| *N*-methyl-6-fluoro-ibogamine | 1730 ± 520 |
| *N*-ethyl-6-fluoro-ibogamine | 1920 ± 570 |
| 5-ethoxy-ibogamine | 80 ± 20 |
| 5-fluoro-ibogamine | 1200 ± 180 |
| 6-fluoro-ibogamine | 1100 ± 170 |

**b**

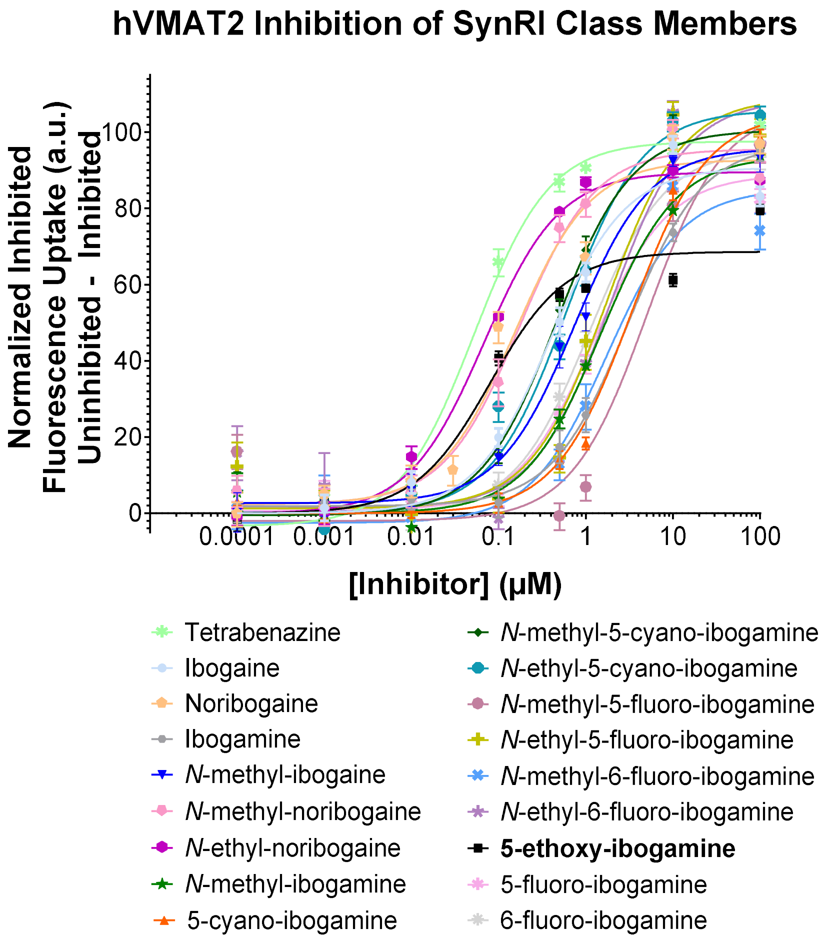

**Figure S2.** hVMAT2 $IC_{50}$ values $\pm$ SEM for iboga alkaloids. (a) Quantification of $IC_{50}$ values of tetrabenazine (a potent and selective VMAT2 inhibitor)[3] and iboga alkaloids at hVMAT2 using FFN206 (final concentration: 0.75 μM) as the substrate for respective functional inhibition experiments. Potent iboga compounds are highlighted in light red. (b) Graphical data is presented as normalized inhibited fluorescence uptake (uninhibited – inhibited) $\pm$ SEM and derived from four separate experiments.

### *Epifluorescence Microscopy*

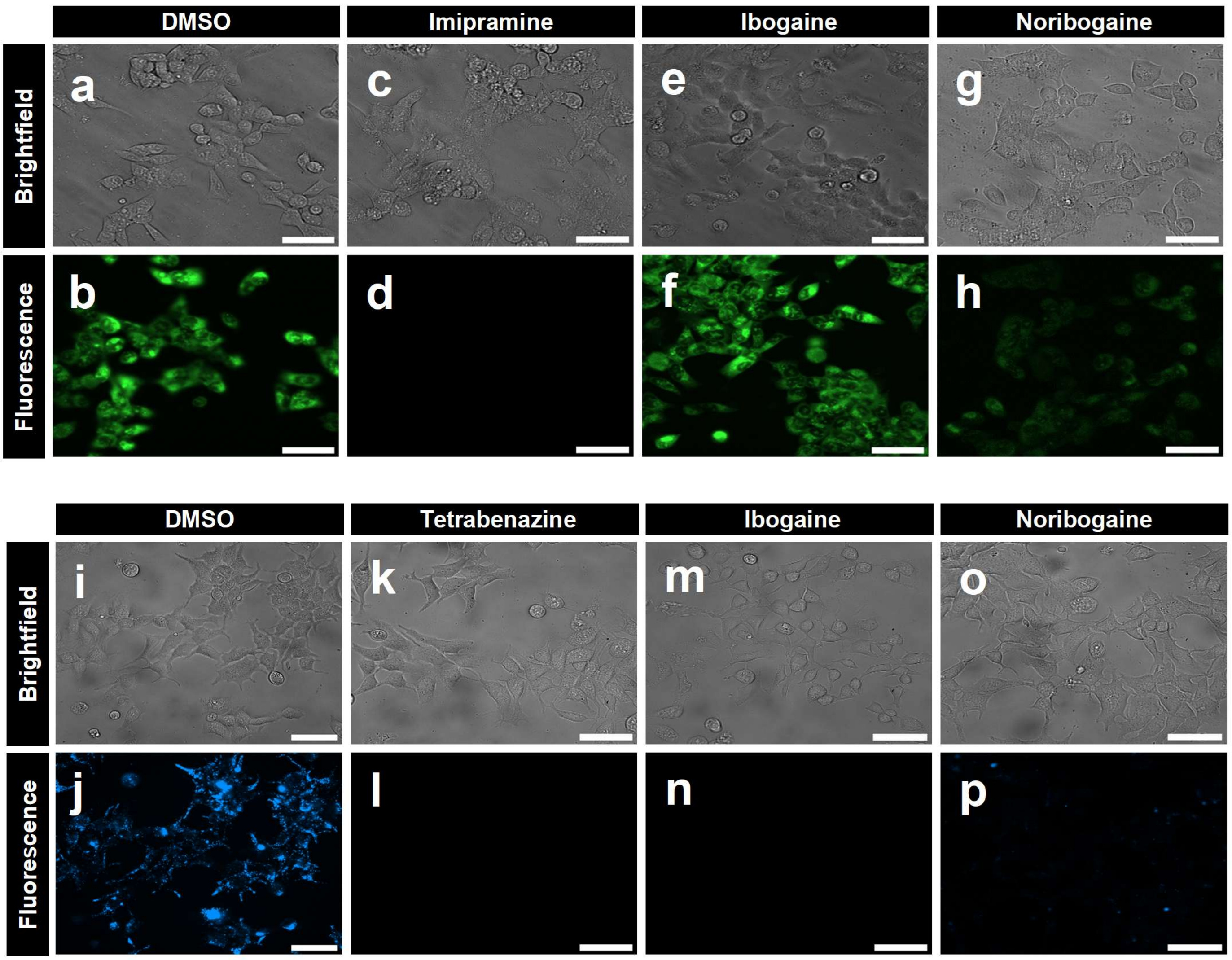


### *Confocal Microscopy*

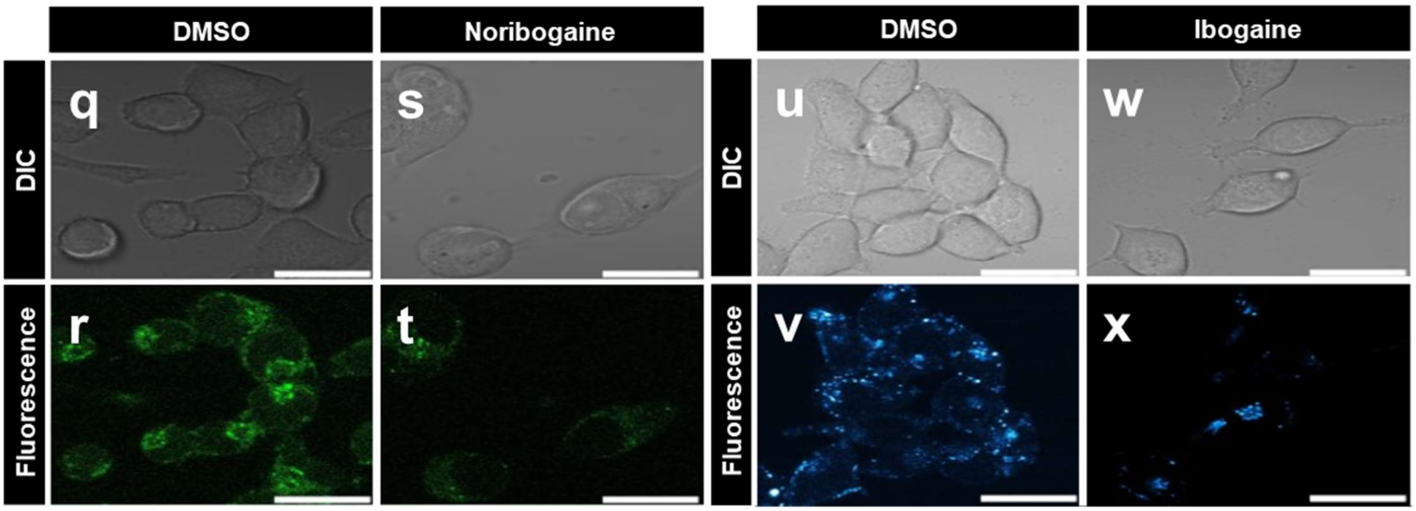

**Figure S3**. Fluorescence microscopy imaging of hSERT-HEK and hVMAT2-HEK cells using appropriate fluorescent substrates in the presence of ibogaine or noribogaine. ***Top panel***: bright field microscopy images of hSERT-HEK cells incubated in the absence or presence of iboga compound (2 μM, 60 minutes) including (e) ibogaine or (g) noribogaine. After the addition of the fluorescence substrate, APP+ (2 μM, five minutes), fluorescence microscopy images showed that uptake was blocked when previously treated with (h) noribogaine. Uptake of APP+ was unhindered with the prior treatment of hSERT-HEK cells with either vehicle (DMSO; b) or (f) ibogaine. (c,d) Imipramine, a potent SERT inhibitor, was utilized as a positive control of uptake inhibition. ***Middle panel***: bright field microscopy images of hVMAT2-HEK cells incubated with either a (i) vehicle (DMSO) or an iboga alkaloid (2 μM, 30 minutes) including (m) ibogaine or (o) noribogaine. After the addition of the fluorescence substrate, FFN206 (20 μM, 120 minutes), fluorescence microscopy images showed lack of uptake in hVMAT2-HEK cell cultures when preincubated with (n) ibogaine or (p) noribogaine, comparable to cell cultures that were incubated with (l) tetrabenazine, a potent selective VMAT2 inhibitor. Scale bar: 25 μm. ***Bottom panel***: Differential Contrast Interference (DIC) and fluorescence images taken by confocal microscopy in either hSERT-HEK or hVMAT2-HEK cells for further qualitative assessment of inhibition. (q,r) Incubation of vehicle (DMSO, 60 minutes) followed by a 30-minute co-incubation with SERTLight (10 μM) demonstrate high intracellular accumulation in hSERT-HEK. Fluorescent uptake was diminished in the presence of (s,t) noribogaine (2 μM, 60 minutes) in comparison to vehicle. Probe fluorescence was observed in hVMAT2-HEK cells pretreated with (u,v) vehicle (DMSO, 30 minutes) followed by treatment with FFN206 (20 μM, 120 minutes). When (w,x) ibogaine (2 μM, 30 minutes) is applied before FFN206 (20 μM, 120 minutes), fluorescence output was reduced. Scale bars for all images (three images per well in triplicate wells) taken at 63× objective: 25 μm.

**a**

| Name | hDAT $IC_{50}$ Values ± SEM (nM) |
|---|---|
| Indatraline | 5.3 ± 1.1 |
| Ibogaine | 12650 ± 2990 |
| Noribogaine | 6760 ± 2440 |
| Ibogamine | 1050 ± 160 |
| *N*-methyl-ibogaine | 710 ± 70 |
| *N*-methyl-noribogaine | 770 ± 100 |
| *N*-ethyl-noribogaine | > 30000 |
| *N*-methyl-ibogamine | 2870 ± 220 |
| 5-cyano-ibogamine | 2330 ± 130 |
| *N*-methyl-5-cyano-ibogamine | 1840 ± 130 |
| *N*-ethyl-5-cyano-ibogamine | 1940 ± 200 |
| *N*-methyl-5-fluoro-ibogamine | 3380 ± 380 |
| *N*-ethyl-5-fluoro-ibogamine | 6770 ± 760 |
| *N*-methyl-6-fluoro-ibogamine | > 30000 |
| *N*-ethyl-6-fluoro-ibogamine | 1010 ± 80 |
| 5-ethoxy-ibogamine | 5290 ± 640 |
| 5-fluoro-ibogamine | 860 ± 110 |
| 6-fluoro-ibogamine | 3080 ± 520 |

**b**

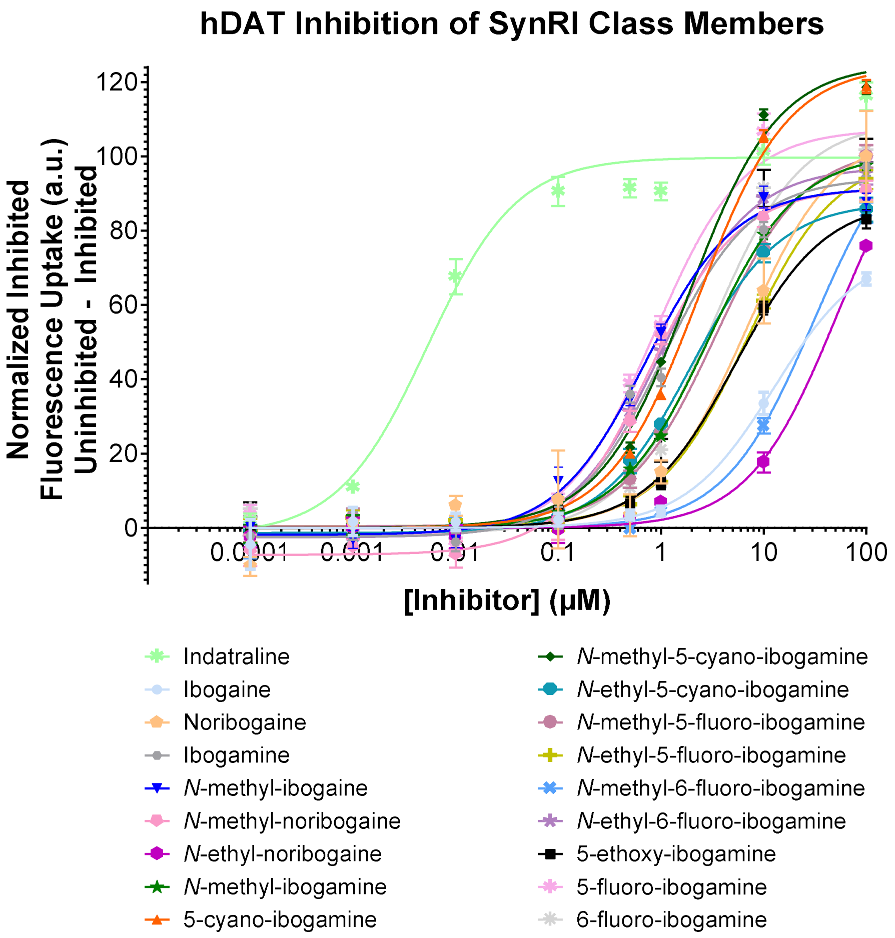

**Figure S4.** hDAT $IC_{50}$ values ± SEM for SynRI iboga alkaloids. (a) Quantification of $IC_{50}$ values of indatraline (a potent DAT inhibitor)[5] and SynRI iboga alkaloids at hDAT using APP+ (final concentration: 1.1 μM)[2] as the substrate for respective functional inhibition experiments. (b) Graphical data is presented as normalized inhibited fluorescence uptake (uninhibited – inhibited) ± SEM and derived from four separate experiments.

**a**

| Name | hNET $IC_{50}$ Values ± SEM (nM) |
|---|---|
| Reboxetine | 3.9 ± 1.3 |
| Ibogaine | 3670 ± 1080 |
| Noribogaine | 19310 ± 2890 |
| Ibogamine | 5020 ± 1180 |
| *N*-methyl-ibogaine | 1800 ± 440 |
| *N*-methyl-noribogaine | 1220 ± 250 |
| *N*-ethyl-noribogaine | 1000 ± 180 |
| *N*-methyl-ibogamine | 8370 ± 1220 |
| 5-cyano-ibogamine | 1670 ± 200 |
| *N*-methyl-5-cyano-ibogamine | 2580 ± 450 |
| *N*-ethyl-5-cyano-ibogamine | 590 ± 60 |
| *N*-methyl-5-fluoro-ibogamine | 2240 ± 290 |
| *N*-ethyl-5-fluoro-ibogamine | 660 ± 60 |
| *N*-methyl-6-fluoro-ibogamine | 7170 ± 1420 |
| *N*-ethyl-6-fluoro-ibogamine | 4990 ± 620 |
| 5-ethoxy-ibogamine | 3820 ± 1150 |
| 5-fluoro-ibogamine | 14170 ± 2100 |
| 6-fluoro-ibogamine | 6290 ± 2510 |

**b**

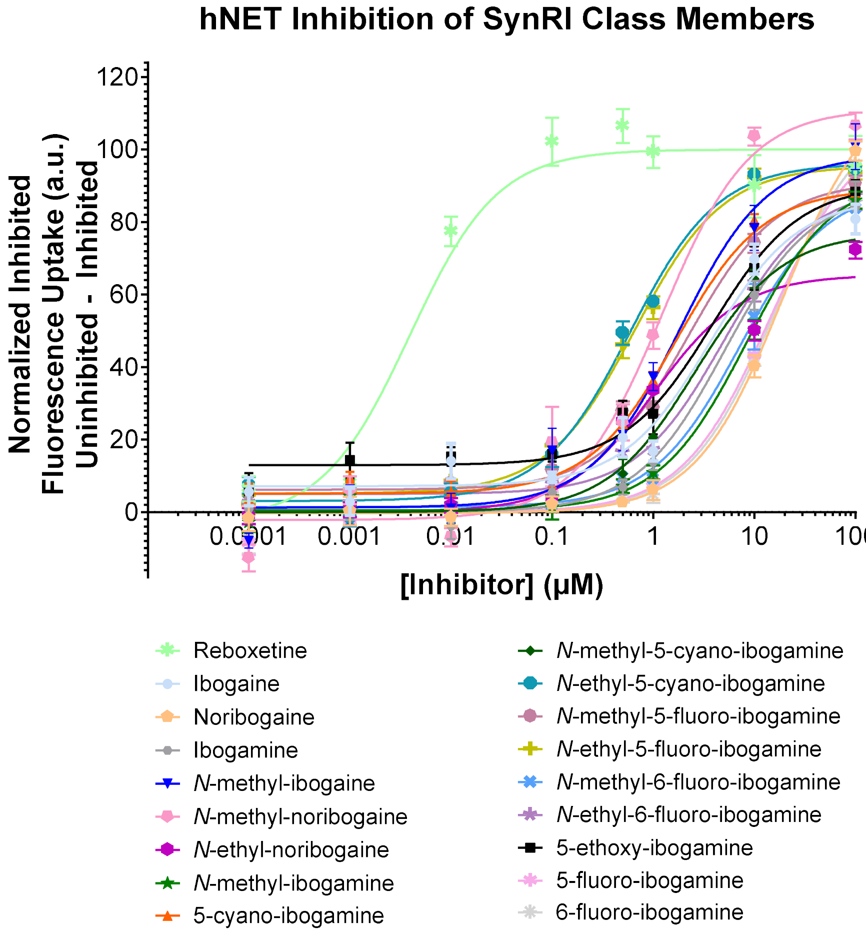

**Figure S5.** hNET $IC_{50}$ values ± SEM for SynRI iboga alkaloids. (a) Quantification of $IC_{50}$ values of reboxetine (a potent NET inhibitor)[6] and SynRI iboga alkaloids at hNET using APP+ (final concentration: 1.1 μM)[2] as the substrate for respective functional inhibition experiments. (b) Graphical data is presented as normalized inhibited fluorescence uptake (uninhibited – inhibited) ± SEM and derived from four separate experiments.

**a**

| Name | hOCT1 $IC_{50}$ Values ± SEM (nM) |
|---|---|
| Decynium-22 | 1420 ± 359 |
| Ibogaine | 22240 ± 5970 |
| Noribogaine | > 30000 |
| Ibogamine | 6190 ± 1350 |
| *N*-methyl-ibogaine | 23530 ± 14450 |
| *N*-methyl-noribogaine | 15040 ± 10920 |
| *N*-ethyl-noribogaine | > 30000 |
| *N*-methyl-ibogamine | > 30000 |
| 5-cyano-ibogamine | 11560 ± 3180 |
| *N*-methyl-5-cyano-ibogamine | > 30000 |
| *N*-ethyl-5-cyano-ibogamine | 12200 ± 5640 |
| *N*-methyl-5-fluoro-ibogamine | 12360 ± 3030 |
| *N*-ethyl-5-fluoro-ibogamine | 5240 ± 1290 |
| *N*-methyl-6-fluoro-ibogamine | 10530 ± 4150 |
| *N*-ethyl-6-fluoro-ibogamine | 14530 ± 3190 |
| 5-ethoxy-ibogamine | 12690 ± 3620 |

**b**

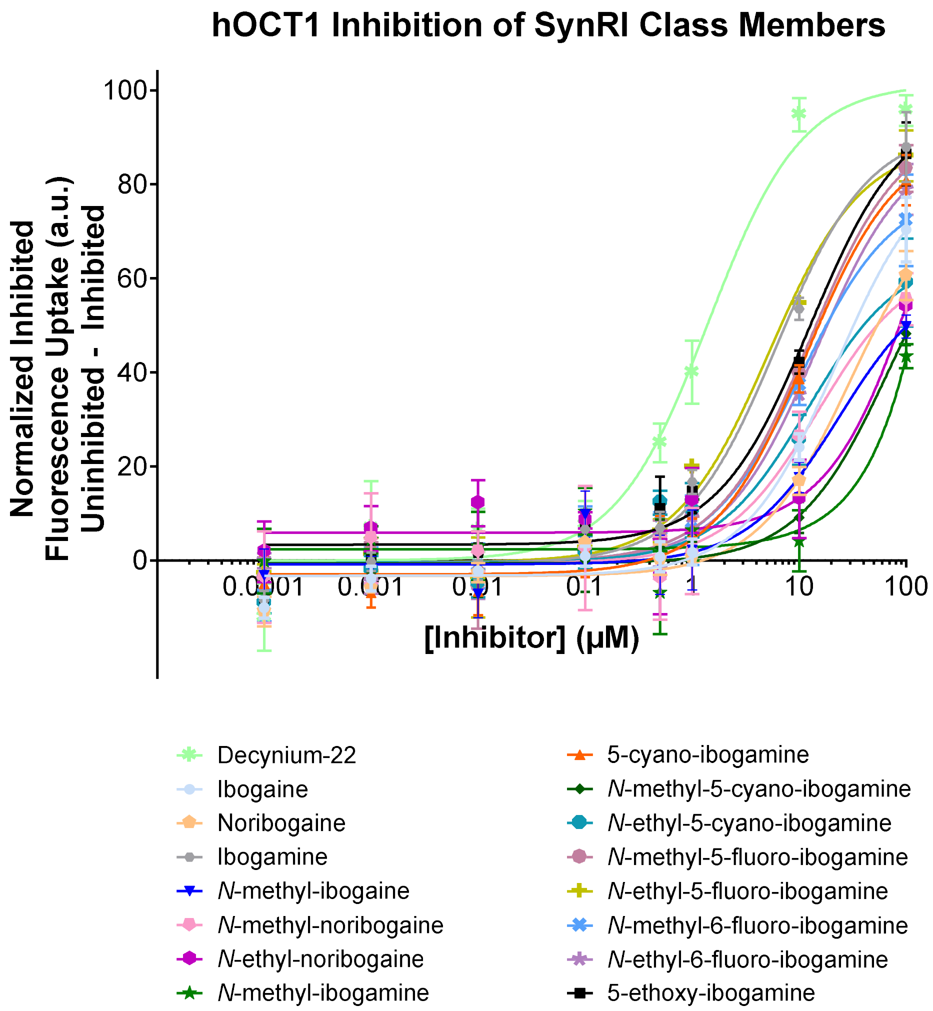

**Figure S6.** hOCT1 $IC_{50}$ values $\pm$ SEM for SynRI iboga alkaloids. (a) Quantification of $IC_{50}$ values of decynium-22 (a hOCT1 inhibitor)[7] and SynRI iboga alkaloids at hOCT1 using APP+ (final concentration: 12.5 μM)[7] as the substrate for respective functional inhibition experiments. (b) Graphical data is presented as normalized inhibited fluorescence uptake (uninhibited – inhibited) $\pm$ SEM and derived from four separate experiments.

**a**

| Name | hOCT2 $IC_{50}$ Values ± SEM (nM) |
|---|---|
| Decynium-22 | 29 ± 8 |
| Ibogaine | 9310 ± 2030 |
| Noribogaine | 6180 ± 1240 |
| Ibogamine | 2050 ± 330 |
| *N*-methyl-ibogaine | 2340 ± 470 |
| *N*-methyl-noribogaine | 5090 ± 1340 |
| *N*-ethyl-noribogaine | 12220 ± 1450 |
| *N*-methyl-ibogamine | 3370 ± 740 |
| 5-cyano-ibogamine | 3450 ± 650 |
| *N*-methyl-5-cyano-ibogamine | 5980 ± 1270 |
| *N*-ethyl-5-cyano-ibogamine | 400 ± 60 |
| *N*-methyl-5-fluoro-ibogamine | 1580 ± 180 |
| *N*-ethyl-5-fluoro-ibogamine | 910 ± 100 |
| *N*-methyl-6-fluoro-ibogamine | 450 ± 70 |
| *N*-ethyl-6-fluoro-ibogamine | 1970 ± 170 |
| 5-ethoxy-ibogamine | 7010 ± 1430 |
| 5-fluoro-ibogamine | 350 ± 70 |
| 6-fluoro-ibogamine | 500 ± 90 |

**b**

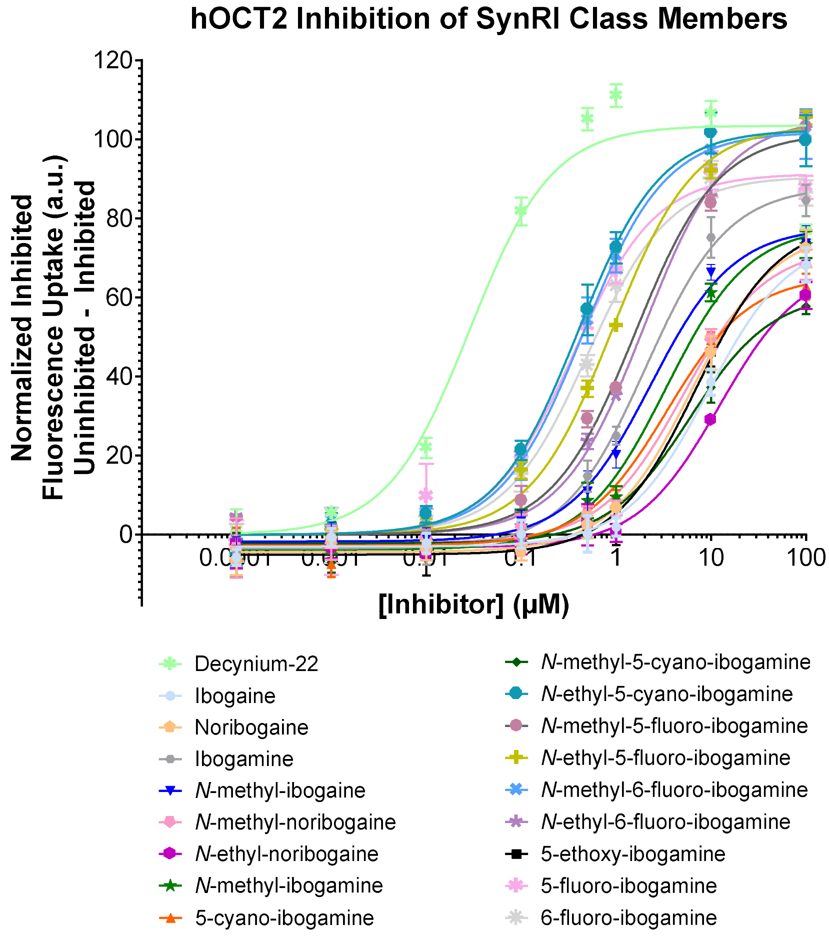

**Figure S7.** hOCT2 $IC_{50}$ values ± SEM for SynRI iboga alkaloids. (a) Quantification of $IC_{50}$ values of decynium-22 (a hOCT2 inhibitor)[7] and SynRI iboga alkaloids at hOCT2 using ASP+ (final concentration: 5 μM)[7] as the substrate for respective functional inhibition experiments. Potent compounds are highlighted in light red. (b) Graphical data is presented as normalized inhibited fluorescence uptake (uninhibited – inhibited) ± SEM and derived from four separate experiments.

**a**

| Name | hOCT3 $IC_{50}$ Values ± SEM (nM) |
|---|---|
| Decynium-22 | 340 ± 104 |
| Ibogaine | > 30000 |
| Noribogaine | > 30000 |
| Ibogamine | > 30000 |
| *N*-methyl-ibogaine | > 30000 |
| *N*-methyl-noribogaine | > 30000 |
| *N*-ethyl-noribogaine | > 30000 |
| *N*-methyl-ibogamine | > 30000 |
| 5-cyano-ibogamine | > 30000 |
| *N*-methyl-5-cyano-ibogamine | No Activity |
| *N*-ethyl-5-cyano-ibogamine | > 30000 |
| *N*-methyl-5-fluoro-ibogamine | > 30000 |
| *N*-ethyl-5-fluoro-ibogamine | 21100 ± 4500 |
| *N*-methyl-6-fluoro-ibogamine | > 30000 |
| *N*-ethyl-6-fluoro-ibogamine | > 30000 |
| 5-ethoxy-ibogamine | > 30000 |

**b**

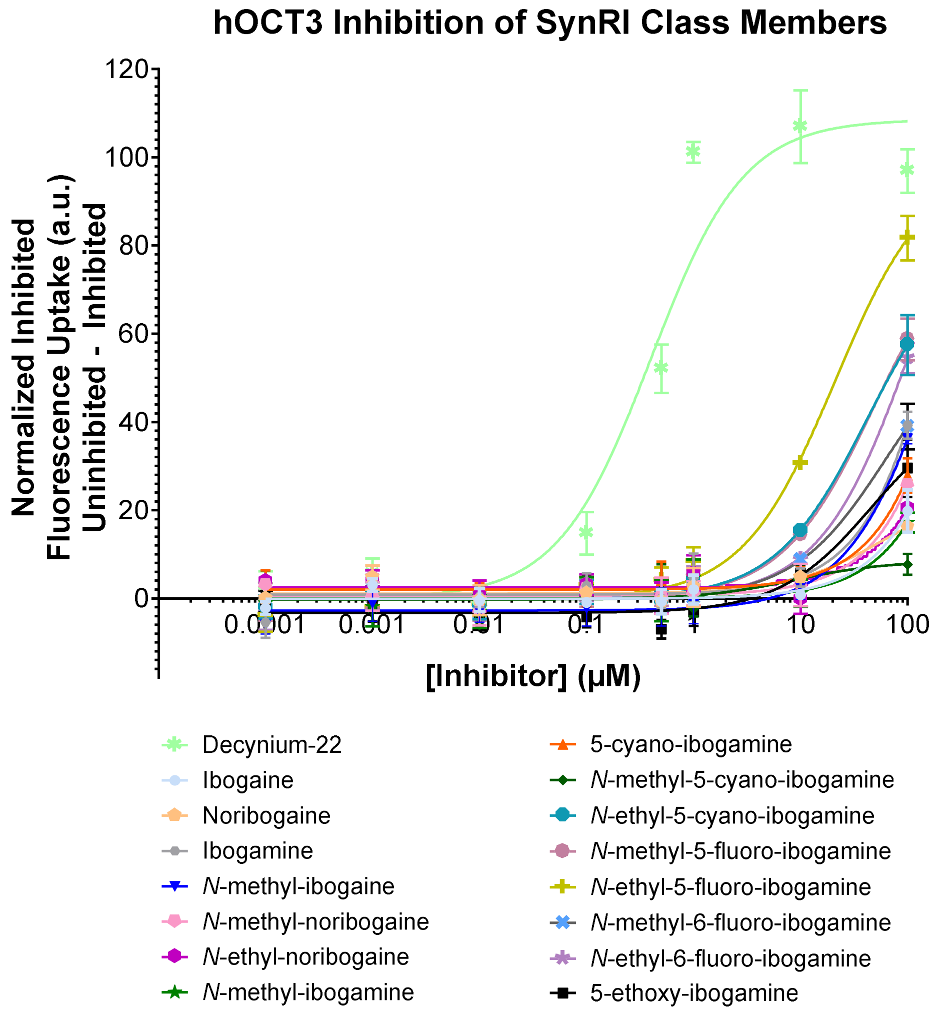

**Figure S8.** hOCT3 $IC_{50}$ values ± SEM for SynRI iboga alkaloids. (a) Quantification of $IC_{50}$ values of decynium-22 (a hOCT3 inhibitor)[7] and SynRI iboga alkaloids at hOCT3 using ASP+ (final concentration: 5 μM)[7] as the substrate for respective functional inhibition experiments. (b) Graphical data is presented as normalized inhibited fluorescence uptake (uninhibited – inhibited) ± SEM and derived from four separate experiments.

**a**

| Name | hPMAT $IC_{50}$ Values ± SEM (nM) |
|---|---|
| Decynium-22 | 362 ± 39 |
| Ibogaine | > 30000 |
| Noribogaine | No Activity |
| Ibogamine | > 30000 |
| *N*-methyl-ibogaine | 16140 ± 8680 |
| *N*-methyl-noribogaine | 13370 ± 8810 |
| *N*-ethyl-noribogaine | > 30000 |
| *N*-methyl-ibogamine | No Activity |
| 5-cyano-ibogamine | > 30000 |
| *N*-methyl-5-cyano-ibogamine | No Activity |
| *N*-ethyl-5-cyano-ibogamine | > 30000 |
| *N*-methyl-5-fluoro-ibogamine | 27450 ± 14250 |
| *N*-ethyl-5-fluoro-ibogamine | 24860 ± 13790 |
| *N*-methyl-6-fluoro-ibogamine | 14470 ± 3470 |
| *N*-ethyl-6-fluoro-ibogamine | 16470 ± 5160 |
| 5-ethoxy-ibogamine | > 30000 |

**b**

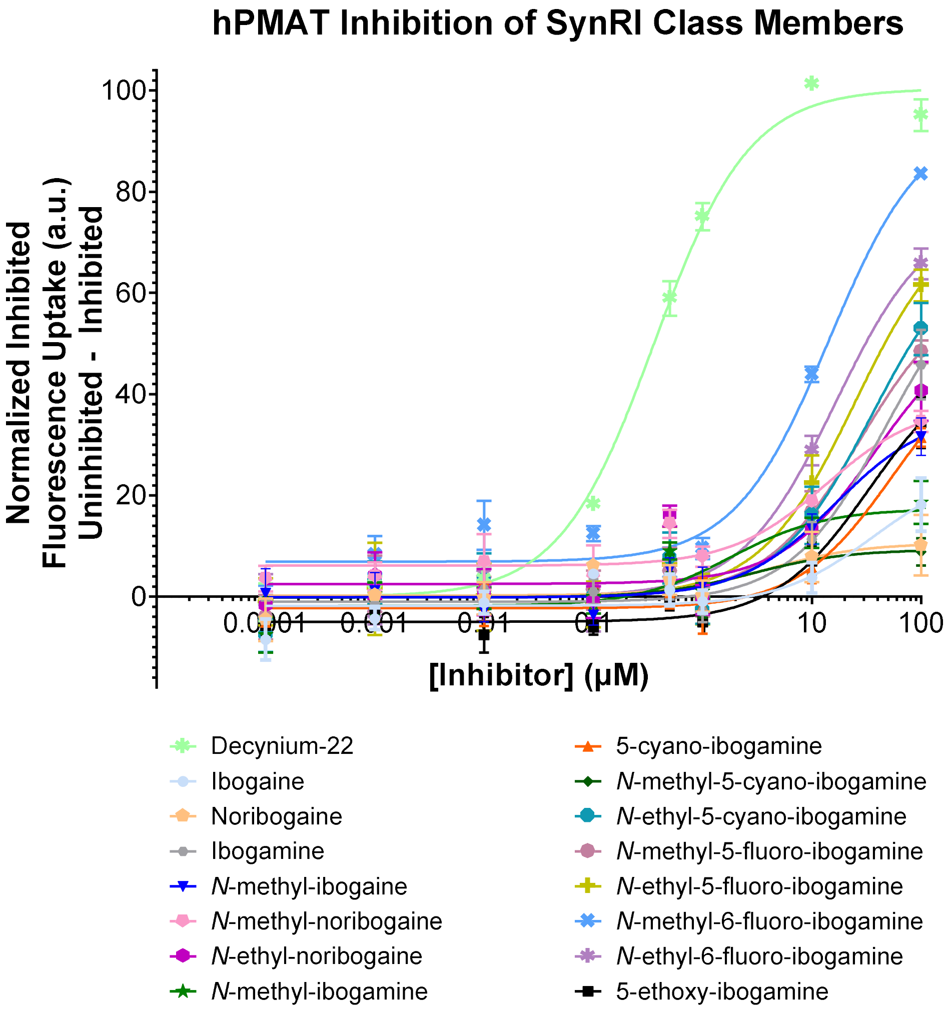

**Figure S9.** hPMAT $IC_{50}$ values ± SEM for SynRI iboga alkaloids. (a) Quantification of $IC_{50}$ values of decynium-22 (a hPMAT inhibitor)[8] and SynRI iboga alkaloids at hPMAT using APP+ (final concentration: 12.5 μM)[7,9] as the substrate for respective functional inhibition experiments. (b) Graphical data is presented as normalized inhibited fluorescence uptake (uninhibited – inhibited) ± SEM and derived from four separate experiments.

| Pharmacological Agent | hDAT *IC₅₀* Values ± SEM (nM) | hNET *IC₅₀* Values ± SEM (nM) | hSERT *IC₅₀* Values ± SEM (nM) | hVMAT2 *IC₅₀* Values ± SEM (nM) |
|---|---|---|---|---|
| Imipramine | 16740 ± 4660 | 510 ± 140 | 6.5 ± 1.5 | 10480 ± 880 |
| Fluoxetine | 7120 ± 1860 | 27840 ± 9810 | 23 ± 3 | 2850 ± 400 |
| Reserpine | No Biological Activity | > 30000 | No Biological Activity | 11 ± 2 |
| Tetrabenazine | > 30000 | 3810 ± 1890 | 21980 ± 9530 | 52 ± 6 |

**Figure S10**. Monoamine transporter inhibition by imipramine, fluoxetine, reserpine, and tetrabenazine. Gray-shaded metrics indicate increased potency of the pharmacological agent at the said transporter. $IC_{50}$ values ± SEM for each pharmacological agent at each of the four Uptake 1 monoamine transporters are displayed and derived from four separate experiments.

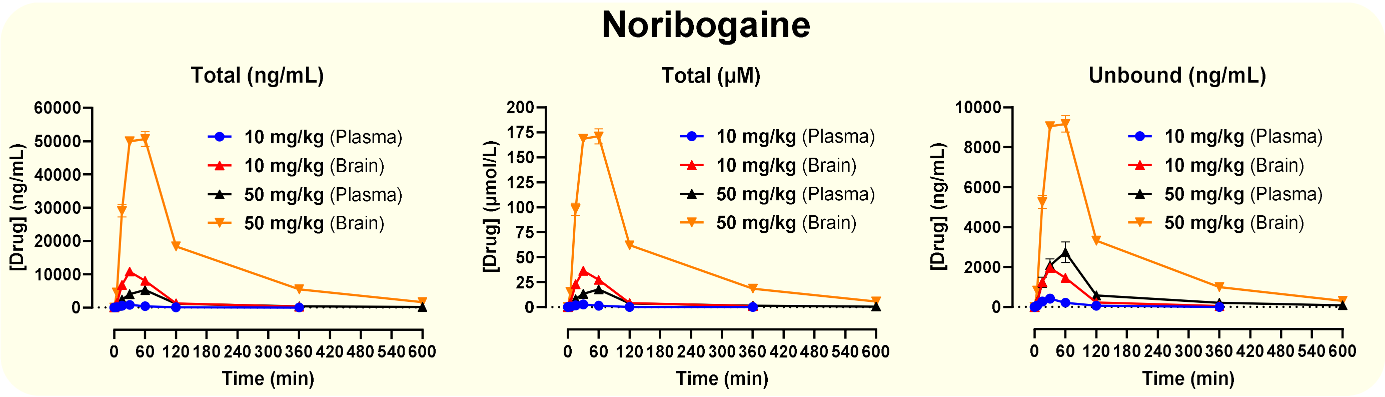


Plasma protein (bound 48%, unbound 52%) and brain tissue (bound 82%, unbound 18%) fractions

| Time (min) | Dose 10 mg/kg | | | | | | Dose 50 mg/kg | | | | | |
|---|---|---|---|---|---|---|---|---|---|---|---|---|
| | Total Plasma (ng/mL) | | | Total Brain (ng/g) | | | Total Plasma (ng/mL) | | | Total Brain (ng/g) | | |
| 5 | 136 | 161 | 93 | 1242 | 1861 | 1036 | 479 | 693 | 382 | 4426 | 6208 | 3011 |
| 15 | 492 | 455 | 623 | 6776 | 5869 | 7801 | 1528 | 2404 | 3207 | 25856 | 32242 | 29040 |
| 30 | 736 | 939 | 770 | 10034 | 11468 | 10891 | 3167 | 3626 | 5219 | 49594 | 50769 | 49593 |
| 60 | 462 | 401 | 412 | 8508 | 7986 | 7727 | 5398 | 6956 | 3495 | 55165 | 48590 | 48241 |
| 180 | 79 | 134 | 113 | 860* | 1414 | 1410 | 1041 | 1127 | 1194 | 16254 | 19135 | 19720 |
| 360 | 0 | 0 | 23* | 214 | 242 | 369 | 342 | 507 | 392 | 5221 | 6827 | 4437 |
| 600 | / | / | / | / | / | / | 137 | 215 | 177 | 1334 | 2114 | 1543 |
| | Total Plasma (μM) | | | Total Brain (μM) | | | Total Plasma (μM) | | | Total Brain (μM) | | |
| 5 | 0.46 | 0.54 | 0.31 | 4.19 | 6.28 | 3.50 | 1.62 | 2.34 | 1.29 | 14.93 | 20.94 | 10.16 |
| 15 | 1.66 | 1.54 | 2.10 | 22.86 | 19.80 | 26.32 | 5.15 | 8.11 | 10.82 | 87.23 | 108.78 | 97.97 |
| 30 | 2.48 | 3.17 | 2.60 | 33.85 | 38.69 | 36.74 | 10.69 | 12.23 | 17.61 | 167.32 | 171.28 | 167.31 |
| 60 | 1.56 | 1.35 | 1.39 | 28.70 | 26.94 | 26.07 | 18.21 | 23.47 | 11.79 | 186.11 | 163.93 | 162.75 |
| 180 | 0.27 | 0.45 | 0.38 | 2.90 | 4.77 | 4.76 | 3.51 | 3.80 | 4.03 | 54.84 | 64.56 | 66.53 |
| 360 | 0.00 | 0.00 | 0.08 | 0.72 | 0.82 | 1.25 | 1.15 | 1.71 | 1.32 | 17.61 | 23.03 | 14.97 |
| 600 | / | / | / | / | / | / | 0.46 | 0.73 | 0.60 | 4.50 | 7.13 | 5.20 |

*Grubbs' outlier test: Significant outlier. $P < 0.05$*

| *Matrix* | Plasma (10 mg/kg) | | Brain (10 mg/kg) | | Plasma (50 mg/kg) | | Brain (50 mg/kg) | |
|---|---|---|---|---|---|---|---|---|
| *$T_{1/2}$ (min)* | 54 | | 61 | | 159 | | 122 | |
| *$T_{max}$ (min)* | 30 | | 30 | | 60 | | 60 | |
| **$C_{max}$** | | | | | | | | |
| *Total* | 815 | 2.75 | 10800 | 36.4 | 5280 | 17.8 | 50700 | 171.1 |
| *Unbound* | 424 | 1.43 | 1955 | 6.59 | 2746 | 9.26 | 9177 | 31.0 |
| *(unit)* | *ng/mL* | *μM* | *ng/g* | *μM* | *ng/mL* | *μM* | *ng/g* | *μM* |
| **$AUC_{0\rightarrow t\ hr}$ ($AUC_{last}$)** | | | | | | | | |
| *Total* | 1070 | 3.6 | 19667 | 66.3 | 13267 | 44.8 | 157000 | 529.7 |
| *Unbound* | 556 | 1.9 | 3560 | 12.0 | 6899 | 23.3 | 28417 | 95.9 |
| *Unit* | *ng × hr/mL* | *μmol × hr/L* | *ng × hr/g* | *μmol × hr/L* | *ng × hr/mL* | *μmol × hr/L* | *ng × hr/g* | *μmol × hr/L* |

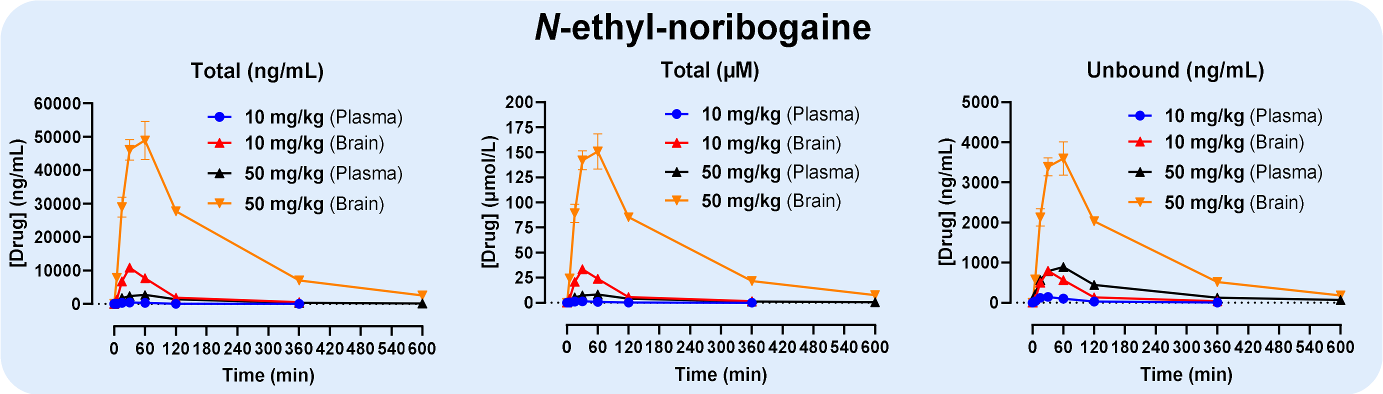


Plasma protein (bound 67%, unbound 33%) and brain tissue (bound 93%, unbound 7%) fractions

| Time (min) | Dose 10 mg/kg | | | | | | Dose 50 mg/kg | | | | | |
|---|---|---|---|---|---|---|---|---|---|---|---|---|
| | Total Plasma (ng/mL) | | | Total Brain (ng/g) | | | Total Plasma (ng/mL) | | | Total Brain (ng/g) | | |
| 5 | 123 | 89 | 94 | 1466 | 1441 | 982 | 703 | 678 | 523 | 8021 | 9502 | 6104 |
| 15 | 303 | 336 | 472 | 5692 | 6235 | 8404 | 1996 | 1375 | 1802 | 32103 | 22996 | 31753 |
| 30 | 511 | 384 | 441 | 12460 | 9655 | 10392 | 2095 | 2799 | 2268 | 41561 | 51881 | 44743 |
| 60 | 264 | 314 | 376 | 6680 | 8073 | 8416 | 2511 | 2932 | 2648 | 39026 | 58741 | 49040 |
| 180 | 64 | 102 | 105 | 1455 | 2053 | 2007 | 1347 | 1559 | 1096 | 29361 | 27141 | 26538 |
| 360 | 26 | 20 | 33 | 580 | 379 | 561 | 460 | 382 | 296 | 7187 | 8341 | 5513 |
| 600 | / | / | / | / | / | / | 174 | 239 | 185 | 2428 | 2598 | 2487 |
| | Total Plasma (µM) | | | Total Brain (µM) | | | Total Plasma (µM) | | | Total Brain (µM) | | |
| 5 | 0.00 | 0.00 | 0.00 | 0.00 | 0.00 | 0.00 | 2.17 | 2.09 | 1.61 | 24.72 | 29.29 | 18.81 |
| 15 | 0.38 | 0.28 | 0.29 | 0.38 | 0.28 | 0.29 | 6.15 | 4.24 | 5.56 | 98.94 | 70.87 | 97.86 |
| 30 | 0.93 | 1.04 | 1.46 | 0.93 | 1.04 | 1.46 | 6.46 | 8.63 | 6.99 | 128.09 | 159.90 | 137.90 |
| 60 | 1.57 | 1.18 | 1.36 | 1.57 | 1.18 | 1.36 | 7.74 | 9.04 | 8.16 | 120.27 | 181.04 | 151.14 |
| 180 | 0.81 | 0.97 | 1.16 | 0.81 | 0.97 | 1.16 | 4.15 | 4.80 | 3.38 | 90.49 | 83.65 | 81.79 |
| 360 | 0.20 | 0.31 | 0.32 | 0.20 | 0.31 | 0.32 | 1.42 | 1.18 | 0.91 | 22.15 | 25.71 | 16.99 |
| 600 | / | / | / | / | / | / | 0.54 | 0.74 | 0.57 | 7.48 | 8.01 | 7.66 |

| *Matrix* | Plasma (10 mg/kg) | Brain (10 mg/kg) | Plasma (50 mg/kg) | Brain (50 mg/kg) |
|---|---|---|---|---|
| *$T_{1/2}$ (min)* | 81 | 74 | 141 | 122 |
| *$T_{max}$ (min)* | 30 | 30 | 60 | 60 |

| | $C_{max}$ | | | | | | | |
|---|---|---|---|---|---|---|---|---|
| *Total* | 445 | 1.37 | 10800 | 33.3 | 2700 | 8.32 | 48900 | 150.7 |
| *Unbound* | 147 | 0.45 | 794 | 2.45 | 891 | 2.75 | 3594 | 11.1 |
| *(unit)* | *ng/mL* | *µM* | *ng/g* | *µM* | *ng/mL* | *µM* | *ng/g* | *µM* |
| | $AUC_{0\to t\ hr}$ ($AUC_{last}$) | | | | | | | |
| *Total* | 920 | 2.8 | 20667 | 63.7 | 9767 | 30.1 | 185000 | 570.2 |
| *Unbound* | 304 | 0.9 | 1519 | 4.7 | 3223 | 9.9 | 13598 | 41.9 |
| *Unit* | *ng × hr/mL* | *µmol × hr/L* | *ng × hr/g* | *µmol × hr/L* | *ng × hr/mL* | *µmol × hr/L* | *ng × hr/g* | *µmol × hr/L* |

**Figure S11.** Pharmacokinetics data collected in male C57BL/6J mice. Determined total drug concentrations and pharmacokinetic parameters were recalculated using rat (male Sprague Dawley) protein binding data to convert to unbound (free) drug fraction.

**a**

| Name | Inhibition of [$^3$H]DA Uptake at DAT $IC_{50}$ ± SD (nM) | Inhibition of [$^3$H]NE Uptake at NET $IC_{50}$ ± SD (nM) | Inhibition of [$^3$H]5HT Uptake at SERT $IC_{50}$ ± SD (nM) | DAT : NET Ratio | DAT : SERT Ratio |
|---|---|---|---|---|---|
| Ibogaine | > 10000 | > 10000 | 3037.0 ± 291.0 | ~ 1 | > 0.304 |
| Noribogaine | > 10000 | 3857.0 ± 288.0 | 326.1 ± 47.6 | > 0.386 | > 0.033 |
| Ibogamine | > 10000 | 4527.0 ± 434.0 | 554.1 ± 49.0 | > 0.453 | > 0.055 |
| *N*-methyl-noribogaine | 4738.0 ± 350.0 | 432.3 ± 49.2 | 13.6 ± 1.2 | 0.090 | 0.003 |
| *N*-ethyl-noribogaine | > 10000 | 81.8 ± 5.8 | 80.1 ± 7.2 | > 0.008 | > 0.008 |
| 5-cyano-ibogamine | > 10000 | 984.5 ± 168.0 | 65.2 ± 5.4 | > 0.098 | > 0.006 |
| *N*-methyl-5-cyano-ibogamine | 9779.0 ± 1390.0 | 440.3 ± 47.4 | 9.8 ± 0.8 | 0.045 | 0.001 |
| Cocaine | 195.6 ± 16.0 | 362.1 ± 46.0 | 295.7 ± 29.5 | 1.856 | 1.512 |

**b**

| Name | [$^3$H]5HT Release by SERT | |
|---|---|---|
| | $EC_{50}$ ± SD (nM) | [% $E_{max}$] |
| Ibogaine | Inactive | Inactive |
| Noribogaine | Inactive | Inactive |
| Ibogamine | Inactive | Inactive |
| *N*-methyl-noribogaine | Inactive | Inactive |
| *N*-ethyl-noribogaine | Inactive | Inactive |
| 5-cyano-ibogamine | Inactive | Inactive |
| *N*-methyl-5-cyano-ibogamine | Inactive | Inactive |
| Amphetamine | 3336.0 ± 445.0 | 91.043 |

**c**

| Name | Inhibition of [$^3$H]DA Uptake at DAT $IC_{50}$ (nM) | Functional APP+ Inhibition at DAT $IC_{50}$ (nM) | Inhibition of [$^3$H]NE Uptake at NET $IC_{50}$ (nM) | Functional APP+ Inhibition at NET $IC_{50}$ (nM) | Inhibition of [$^3$H]5HT Uptake at SERT $IC_{50}$ (nM) | Functional APP+ Inhibition at SERT $IC_{50}$ (nM) |
|---|---|---|---|---|---|---|
| Ibogaine | > 10000 | 12650 | > 10000 | 3670 | 3037.0 | 2980 |
| Noribogaine | > 10000 | 6760 | 3857.0 | 19310 | 326.1 | 280 |
| Ibogamine | > 10000 | 1050 | 4527.0 | 5020 | 554.1 | 330 |
| *N*-methyl-noribogaine | 4738.0 | 770 | 432.3 | 1220 | 13.6 | 59 |
| *N*-ethyl-noribogaine | > 10000 | > 30000 | 81.8 | 1000 | 80.1 | 1860 |
| 5-cyano-ibogamine | > 10000 | 2330 | 984.5 | 1670 | 65.2 | 26 |
| *N*-methyl-5-cyano-ibogamine | 9779.0 | 1840 | 440.3 | 2580 | 9.8 | 5.1 |

**d**

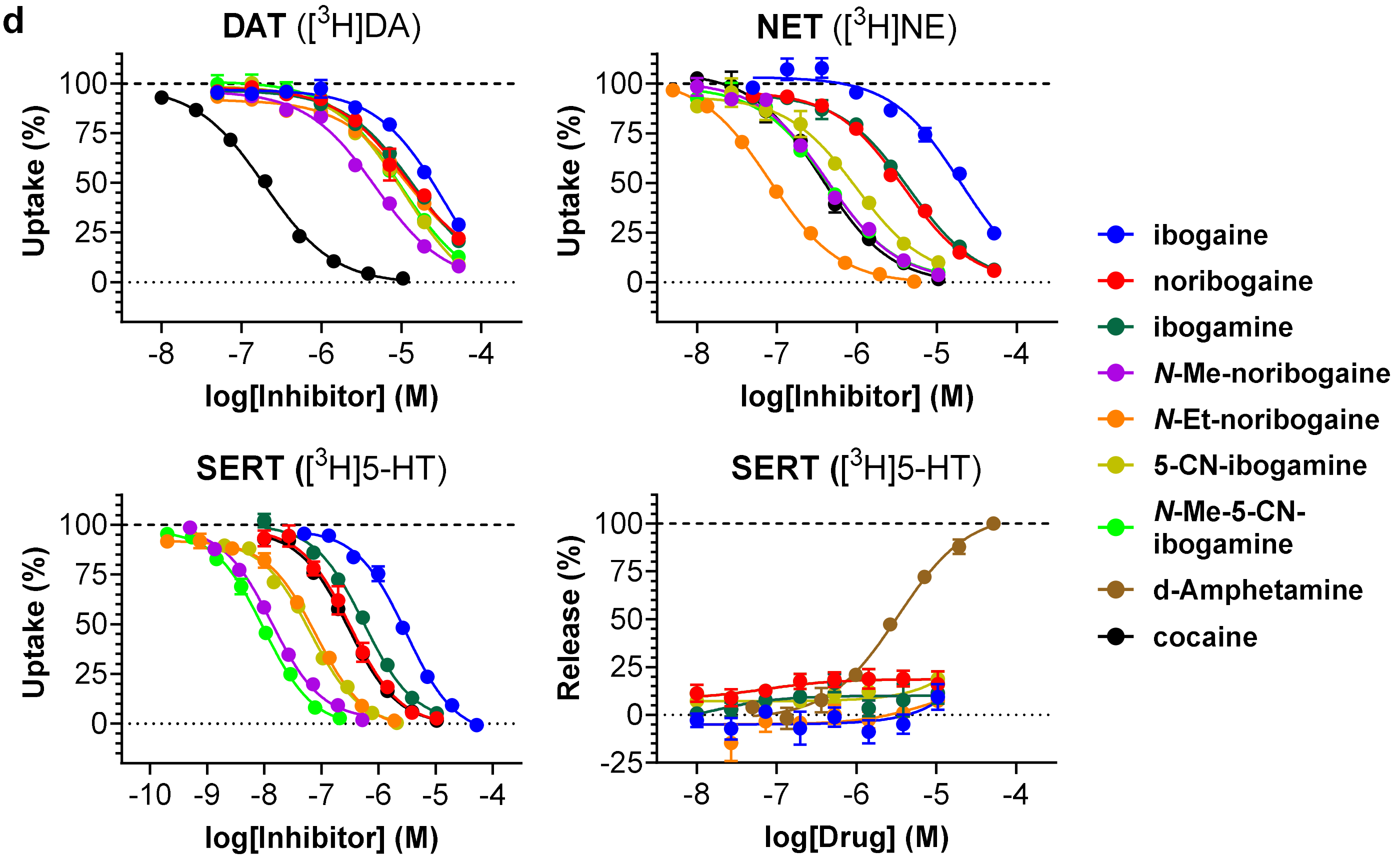

**Figure S12**. Effects of ibogaine analogs on tritiated monoamine uptake and release in rat brain synaptosomes. (a) Quantification of $IC_{50}$ values of ibogaine and respective analogs at DAT, NET, and SERT upon measurement of inhibited tritiated monoamine uptake in rat brain synaptosomes. Values are presented as inhibited tritiated monoamine uptake $\pm$ SD and derived from three separate experiments. Additionally, DAT : NET and DAT : SERT ratios are calculated for each tested compound. (b) Quantification of $EC_{50}$ values of ibogaine and respective analogs at SERT upon measurement of tritiated 5-HT released by SERT in rat brain synaptosomes. Values are presented as tritiated 5-HT release $\pm$ SD and derived from three separate experiments. (c) Side-by-side comparison of inhibition metrics derived from inhibition of tritiated monoamine uptake in rat striatal synaptosomes with those derived from drug inhibition of APP+ uptake by transfected hDAT, hNET, and hSERT expressed in HEK cells. Note that inhibition values from functional APP+ inhibition were averaged from four separate experiments. (d) Dose-response curves for inhibition of [$^3$H]DA, [$^3$H]NE and [$^3$H]5-HT uptake. Iboga derivatives do not induce release of [$^3$H]5-HT via SERT.

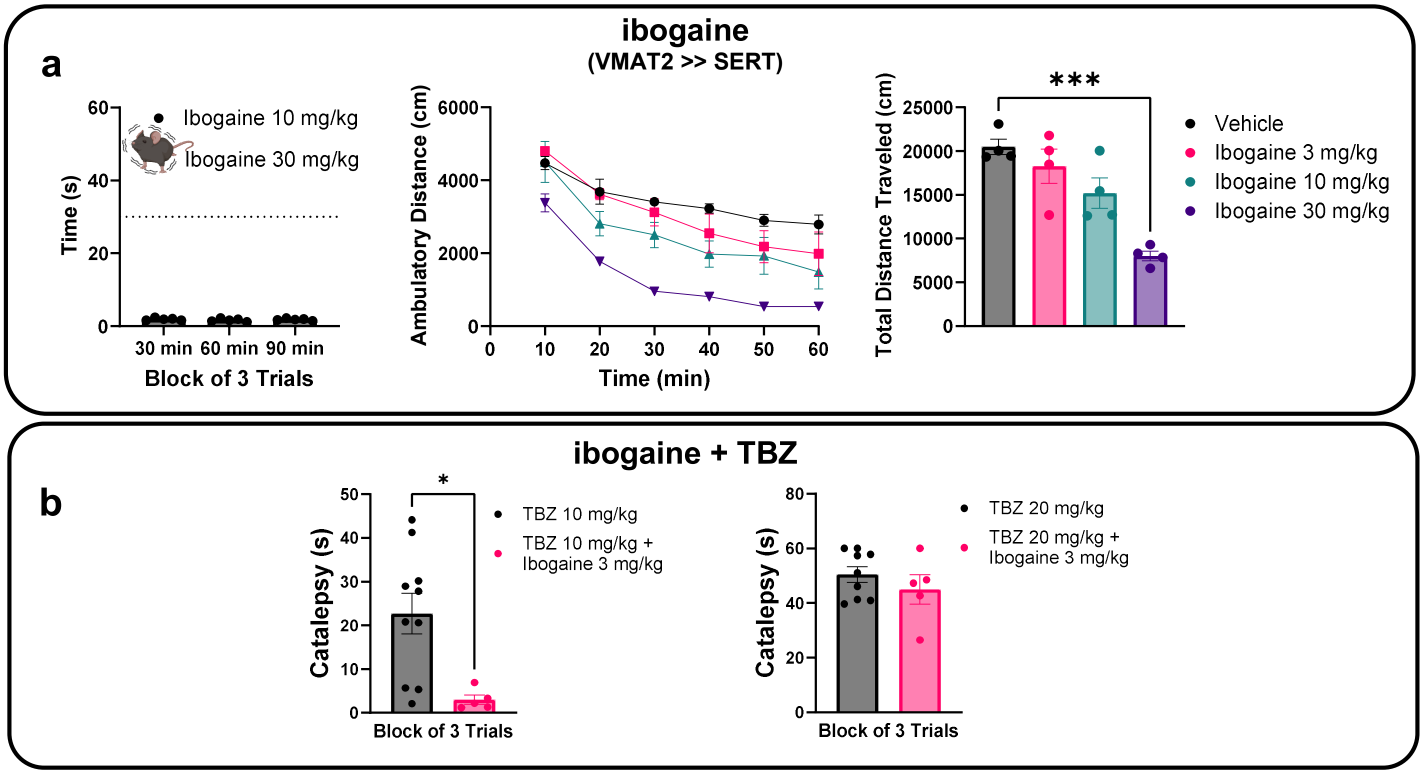


**Figure S13. Ibogaine Effect in Catalepsy Bar Test and Locomotion.** (a) Evaluation of catalepsy and locomotor activity induced by ibogaine. Catalepsy was assessed via the bar test at 30-, 60-, and 90-minutes post subcutaneous injection ($n$ = 5 per group). Locomotor activity was measured in an open field for one hour ($n$ = 4 per group). Data analyzed using one-way ANOVA followed by Dunnett's post hoc test, with comparisons made against the vehicle group (b) Ibogaine attenuated the effects of TBZ at 10 mg/kg (P = 0.0119) but had minimal impact at 20 mg/kg. Data are presented as mean ± SEM, with group sizes ranging from 4 to 10 (n = 4–10). Statistical significance was determined using unpaired t-tests. p values indicate comparisons between TBZ-treated and TBZ + ibogaine-treated groups. The mouse icon indicates the dosages at which the drug induced tremors (catalepsy was not assessed at this dose as tremorgenic activity is a confounding factor). All values represented as mean ± SEM, ***p < 0.001, **p < 0.01, *p < 0.05

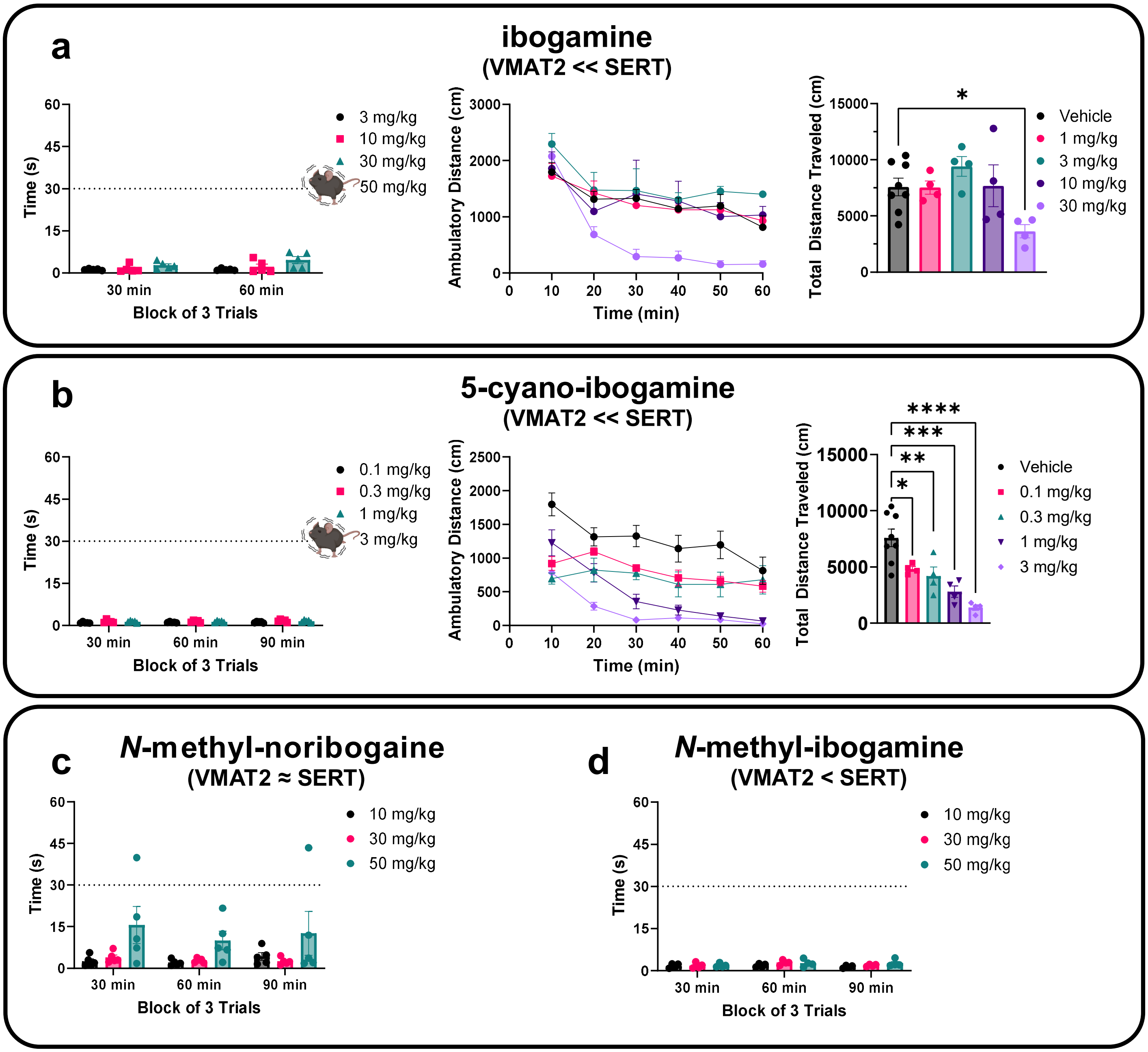


**Figure S14.** Evaluation of catalepsy and locomotor activity induced by (a) ibogamine and (b) 5-cyano-ibogamine. Catalepsy was assessed via the bar test 30, 60, 90 minutes post subcutaneous injection ($n$ = 4 – 9 / group). Locomotor activity was measured in an open field for one hour ($n$ = 4 – 8 / group). (c) Evaluation of catalepsy induced by *N*-methyl-noribogaine and (d) *N*-methyl-ibogamine at 30, 60, 90 min post injection. The mouse icon indicates the dosages at which the drug induced tremors (catalepsy was not assessed at this dose as tremorgenic activity is a confounding factor). Data was analyzed using one-way ANOVA followed by Dunnett's post hoc test, with comparisons made against the vehicle group. All values represented as average ± SEM, ****p < 0.0001, ***p < 0.001, **p < 0.01, *p < 0.05.

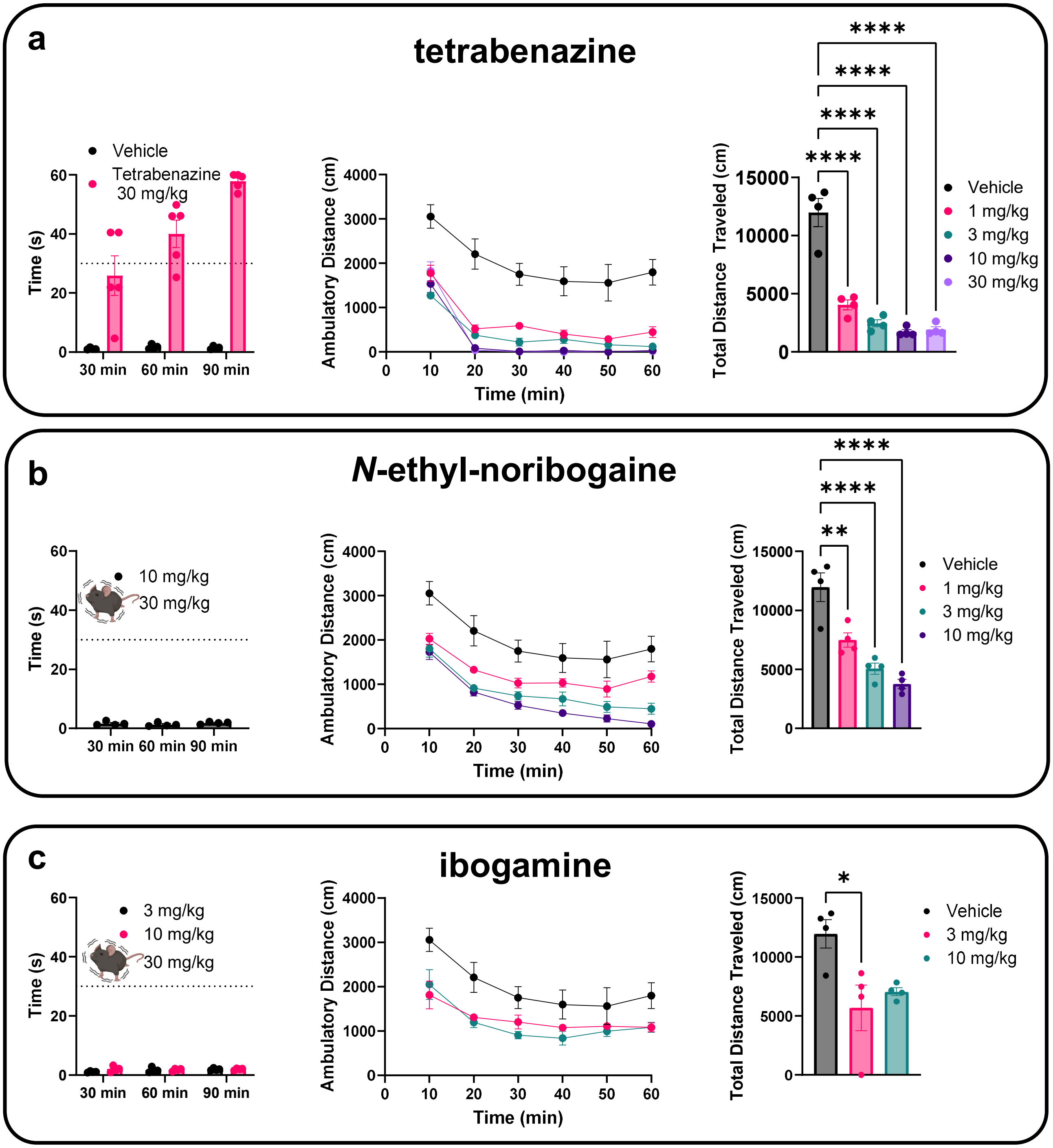


**Figure S15.** Evaluation of catalepsy and locomotor activity induced by (a) tetrabenazine, (b) *N*-ethyl-noribogaine, and (c) ibogamine in female mice. Catalepsy was assessed via the bar test at 30, 60, 90 minutes post subcutaneous injection ($n$ = 4 – 9 / group). The mouse icon indicates the dosages at which the drug induced tremors (catalepsy was not assessed at this dose as tremorgenic activity is a confounding factor). Locomotor activity was measured in an open field for one hour ($n$ = 4 / group). All values represented as mean ± SEM, ****p < 0.0001, ***p < 0.001, **p < 0.01, *p < 0.05.

| Assay | Ligand | Species | % Inh. |
|---|---|---|---|
| Cholinesterase, Acetyl, ACES | Acetylthiocholine | Human | 28 |
| Cyclooxygenase COX-1 | Arachidonic acid | Human | 1 |
| Cyclooxygenase COX-2 | Arachidonic Acid | Human | -7 |
| Monoamine Oxidase MAO-A | Kynuramine | Human | 5 |
| Phosphodiesterase PDE3A | FAM-cAMP | Human | -7 |
| Phosphodiesterase PDE4D2 | FAM-cAMP | Human | 1 |
| Protein Tyrosine Kinase, LCK | Poly(Glu:Tyr) | Human | 7 |
| Adenosine A2A | [$^3$H]CGS-21680 | Human | 9 |
| **Adrenergic alpha1A** | **[$^3$H]Prazosin** | **Human** | **51** |
| Adrenergic alpha2A | [$^3$H]Rauwolscine | Human | 2 |
| Adrenergic beta1 | [$^{125}$I]Cyanopindolol | Human | -13 |
| Adrenergic beta2 | [$^3$H]CGP-12177 | Human | -6 |
| Norepinephrine Transporter (NET) | [$^{125}$I]RTI-55 | Human | 23 |
| Androgen (Testosterone) | [$^3$H]Methyltrienolone | Human | 41 |
| Calcium Channel L-Type, Dihydropyridine | [$^3$H]Nitrendipine | Rat (cerebral cortex) | -4 |
| Cannabinoid CB1 | [$^3$H]SR141716A | Human | -5 |
| Cannabinoid CB2 | [$^3$H]WIN-55,212-2 | Human | -11 |
| Cholecystokinin CCK1 (CCKA) | [$^{125}$I]CCK-8 | Human | 2 |
| Dopamine D1 | [$^3$H]SCH-23390 | Human | 7 |
| Dopamine D2S | [$^3$H]Spiperone | Human | 9 |
| Dopamine Transporter (DAT) | [$^{125}$I]RTI-55 | Human | 37 |
| Endothelin ETA | [$^{125}$I]Endothelin-1 | Human | 3 |
| GABAA, Flunitrazepam, Central | [$^3$H]Flunitrazepam | Rat (brain no cerebellum) | -5 |
| Glucocorticoid | [$^3$H]Dexamethasone | Human | 1 |
| Glutamate, NMDA, Agonism | [$^3$H]CGP-39653 | Rat | 0 |
| Histamine H1 | [$^3$H]Pyrilamine | Human | 10 |
| Histamine H2 | [$^{125}$I]Aminopotentidine | Human | 10 |
| Muscarinic M1 | [$^3$H]*N*-Methylscopolamine | Human | 11 |
| Muscarinic M2 | [$^3$H]*N*-Methylscopolamine | Human | 3 |
| Muscarinic M3 | [$^3$H]*N*-Methylscopolamine | Human | -3 |
| Opiate delta1 (OP1, DOP) | [$^3$H]Naltrindole | Human | -3 |
| Opiate kappa (OP2, KOP) | [$^3$H]Diprenorphine | Human | 13 |
| Opiate mu (OP3, MOP) | [$^3$H]Diprenorphine | Human | 17 |
| Potassium Channel (KA) | [$^{125}$I]alpha-Dendrotoxin | Rat | -4 |
| **Potassium Channel hERG** | **[$^3$H]Dofetilide** | **Human** | **76** |
| 5-HT$_{1A}$ | [$^3$H]8-OH-DPAT | Human | -11 |
| 5-HT$_{1B}$ | [$^3$H]GR125743 | Human | -5 |
| 5-HT$_{2A}$ | [$^3$H]Ketanserin | Human | 21 |
| 5-HT$_{2B}$ | [$^3$H]Lysergic acid diethylamide | Human | -5 |
| 5-HT$_3$ | [$^3$H]GR-65630 | Human | 10 |
| Serotonin Transporter (SERT) | [$^3$H]Paroxetine | Human | 1 |
| Sodium Channel, Site 2 | [$^3$H]Batrachotoxinin | Rat | 43 |
| Vasopressin V1A | [$^{125}$I]PhenylacetylTyr(Me) PheGlnAsnArgProArgTyr | Human | -19 |
| Nicotinic Acetylcholine alpha4beta2 | [$^3$H]Cytisine | Human | 2 |

**Figure S16**. Results of SafetyScreen44 (commercial screening assay **PP241**, Eurofins Panlabs Discovery Services) of Ibogaine Hydrochloride at 10 μM.
https://www.eurofinsdiscovery.com/catalog/safetyscreen44-panel-tw/PP241

### B. Synthesis.

#### 1. Materials and Methods.

**General Considerations.** Reagents and solvents were obtained from commercial sources (Fisher Scientific or Sigma Aldrich) and were used without further purification unless otherwise stated. The enantio-enriched isoquinuclidine used for respective compounds was resolved from the racemic exo-isoquinuclidine by WuXi AppTec (Shanghai) Co. Ltd. Reactions were monitored by TLC using solvent mixtures appropriate to each reaction. Column chromatography was performed on silica gel (40 – 63 μm) or basic alumina (60 – 200 μm). For preparative TLC, glass plates coated with a 1 mm silica layer were used. Nuclear magnetic resonance spectra were recorded on Bruker 400 or 500 MHz instruments, as indicated. Chemical shifts are reported as δ values in ppm referenced to $CDCl_3$ ($^1H$ NMR = 7.26 and $^{13}C$ NMR = 77.16), methanol-$d_4$ ($^1H$ NMR = 3.31 and $^{13}C$ NMR = 49.00) or water-$d_2$ ($^1H$ NMR = 4.79). Multiplicity is indicated as follows: s (singlet); d (doublet); t (triplet); q (quartet); p (pentet); dd (doublet of doublets); td (triplet of doublets); dt (doublet of triplets); dq (doublet of quartets); ddd (doublet of doublet of doublets); ddt (doublet of doublet of triplets); m (multiplet); br (broad). All carbon peaks are rounded to one decimal place unless such rounding would cause two close peaks to become identical; in these cases, two decimal places are retained. Low-resolution mass spectra were recorded on an Advion CMS quadrupole instrument (ionization mode: APCI+ or ESI+) or Agilent GC-MS (ionization mode: EI). High-resolution mass spectra (HRMS) were acquired on a high-resolution Waters XEVO G2-XS QToF mass spectrometer equipped with a UPC2 SFC inlet, on-board fluidics and an ESI probe.

#### 2. Synthetic Protocols.

**Noribogaine hydrochloride.**

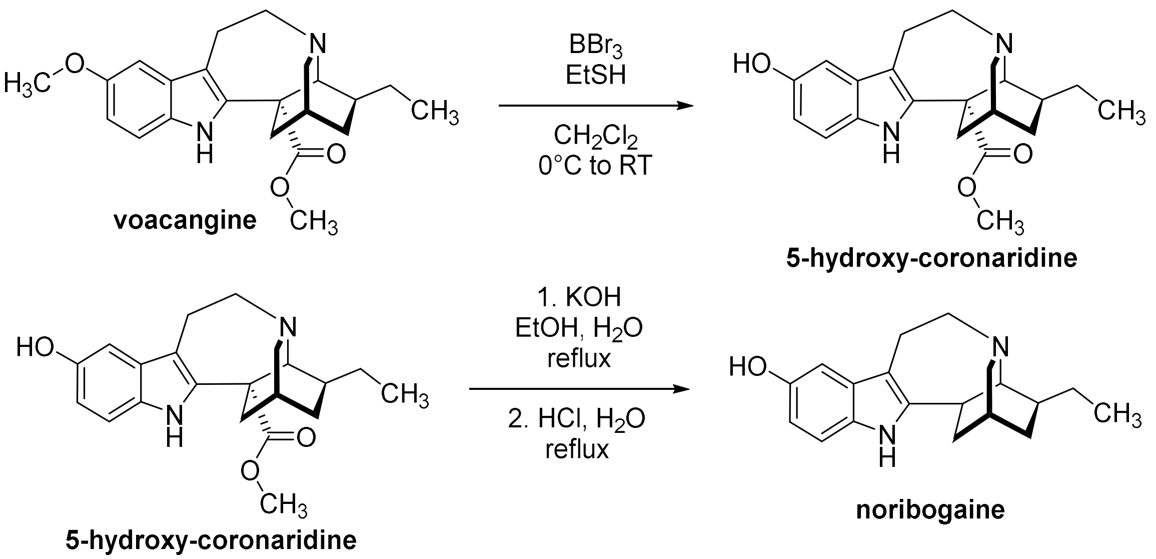


All novel iboga analogs were prepared starting from voacangine alkaloid isolated from *Voacanga africana* according to published procedure.[10] Noribogaine was prepared according to literature procedure from 5-hydroxy-coronaridine to avoid working with ibogaine (DEA schedule I substance in the United States).[11]

**Noribogaine hydrochloride: $^{1}$H NMR (500 MHz, $D_2O$)** δ 7.33 (dd, *J* = 8.7, 0.6 Hz, 1H), 7.06 – 6.95 (m, 1H), 6.84 (ddd, *J* = 8.6, 2.4, 0.5 Hz, 1H), 3.60 (dt, *J* = 13.5, 4.3 Hz, 1H), 3.54 – 3.44 (m, 2H), 3.39 (dt, *J* = 12.4, 2.8 Hz, 1H), 3.32 (ddd, *J* = 12.2, 5.0, 1.8 Hz, 1H), 3.21 (dt, *J* = 12.4, 2.4 Hz, 1H), 3.14 – 3.09 (m, 2H), 2.33 – 2.24 (m, 1H), 2.18 – 2.08 (m, 2H), 2.05 – 1.96 (m, 1H), 1.61 – 1.49 (m, 3H), 1.35 (ddt, *J* = 10.8, 5.1, 2.7 Hz, 1H), 1.00 (t, *J* = 7.3 Hz, 3H).

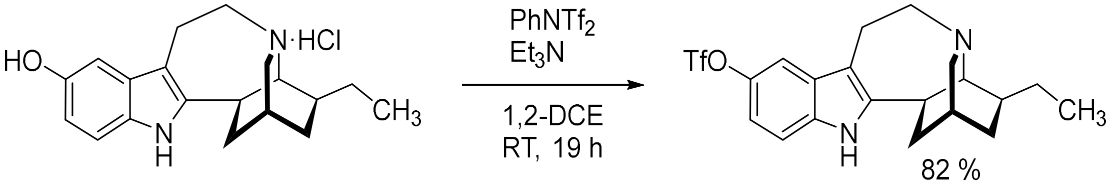


**Ibogamine-5-triflate**

Noribogaine hydrochloride (100 mg, 0.3 mmol) and bis(trifluoromethanesulfonyl)aniline (118 mg, 0.33 mmol) were suspended in 1,2-dichloroethane (anhydrous, 2.2 mL) and $Et_3N$ (125 µL, 0.9 mmol) was added. After stirring at RT for 19 h, reaction was quenched by pouring into $H_2O$ (15 mL) and the mixture was extracted with $CH_2Cl_2$ (3 × 10 mL). Combined extracts were dried over $Na_2SO_4$, filtered, and concentrated under reduced pressure. Crude material was purified by column chromatography (gradient of 10 to 20% AcOEt in hexanes + 2% $Et_3N$). Ibogamine-5-triflate was obtained as an off-white amorphous solid (106 mg, 82%).

**$^{1}$H NMR (500 MHz, $CDCl_3$)** δ 7.85 (s, 1H), 7.33 (d, *J* = 2.4 Hz, 1H), 7.22 (d, *J* = 8.7 Hz, 1H), 6.98 (dd, *J* = 8.7, 2.4 Hz, 1H), 3.42 – 3.30 (m, 2H), 3.19 – 3.09 (m, 1H), 3.06 (dt, *J* = 9.4, 2.4 Hz, 1H), 3.00 (dt, *J* = 9.4, 3.0 Hz, 1H), 2.93 (ddd, *J* = 11.6, 4.3, 1.8 Hz, 1H), 2.85 (t, *J* = 1.7 Hz, 1H), 2.58 (ddd, *J* = 16.2, 5.8, 3.0 Hz, 1H), 2.06 (ddt, *J* = 14.1, 11.9, 2.7 Hz, 1H), 1.90 – 1.77 (m, 2H), 1.64 (dq, *J* = 13.2, 3.4 Hz, 1H), 1.62 – 1.51 (m, 2H), 1.47 (pd, *J* = 8.9, 2.3 Hz, 1H), 1.22 (ddt, *J* = 12.9, 5.2, 2.6 Hz, 1H), 0.91 (t, *J* = 7.1 Hz, 3H). **$^{19}$F NMR (471 MHz, $CDCl_3$)** δ -71.9. **$^{13}$C NMR (126 MHz, $CDCl_3$)** δ 144.9, 143.6, 133.6, 130.3, 122.8, 120.3, 117.7, 115.2, 113.9, 111.0, 110.5, 110.4, 57.4, 54.1, 50.1, 42.1, 41.7, 34.2, 32.1, 28.0, 26.6, 20.7, 12.0. **LRMS (APCI+)** calcd. for $C_{20}H_{24}F_3N_2O_3S^+$ $[M+H]^+$ 429.1, found 429.0.

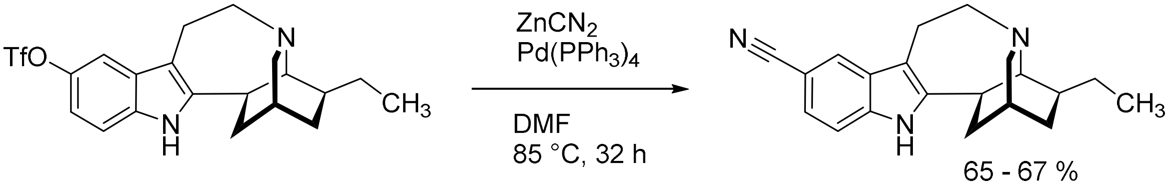


**5-cyano-ibogamine**

Ibogamine-5-triflate (128 mg, 0.3 mmol), $ZnCN_2$ (140 mg, 1.2 mmol), $Pd(PPh_3)_4$ (35 mg, 0.03 mmol) were balanced into a reaction vial and dried under vacuum. DMF (anhydrous, 1.5 mL) was added, vial was flushed with argon, closed with a Teflon lined solid screw cap and the reaction mixture was heated to 85°C. After 15 h, TLC indicated only partial conversion. Additional $ZnCN_2$ (140 mg, 1.2 mmol) and $Pd(PPh_3)_4$ (35 mg, 0.03 mmol) were added and heating was continued for an additional 23 h, until TLC indicated that no more starting material remained. The reaction mixture was filtered through a plug of silica and eluted with AcOEt + 2% $Et_3N$. Crude material was purified by column chromatography (gradient of 20 to 30% of AcOEt in hexanes + 2% $Et_3N$). Product was isolated as a white amorphous solid (59 mg, 65%).

**$^1$H NMR (500 MHz, $CDCl_3$)** δ 8.15 (s, 1H), 7.79 (s, 1H), 7.33 (d, *J* = 8.2 Hz, 1H), 7.29 (d, *J* = 8.3 Hz, 1H), 3.43 – 3.30 (m, 2H), 3.19 – 3.09 (m, 1H), 3.09 – 2.93 (m, 3H), 2.85 (s, 1H), 2.66 – 2.57 (m, 1H), 2.13 – 2.02 (m, 1H), 1.90 – 1.84 (m, 1H), 1.84 – 1.76 (m, 1H), 1.64 (dq, *J* = 13.9, 3.4 Hz, 1H), 1.60 – 1.50 (m, 2H), 1.50 – 1.41 (m, 1H), 1.25 – 1.16 (m, 1H), 0.89 (t, *J* = 7.0 Hz, 3H). **$^{13}$C NMR (126 MHz, $CDCl_3$)** δ 144.4, 136.6, 129.8, 124.1, 123.5, 121.3, 111.0, 110.3, 102.0, 57.5, 54.0, 50.1, 42.0, 41.5, 34.2, 32.1, 27.9, 26.5, 20.6, 12.0. **HRMS (ESI+)** calcd. for $C_{20}H_{24}N_3^+$ $[M+H]^+$ 306.1970, found 306.1980.

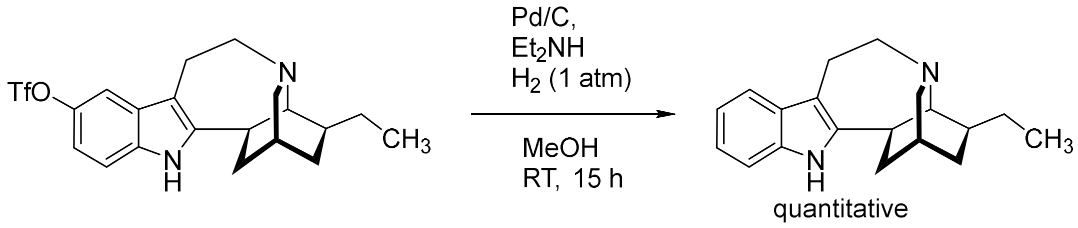


#### Ibogamine

Ibogamine-5-triflate (20 mg, 0.047 mmol) was dissolved in MeOH (not dry, 1.4 mL) Pd/C (10%, moistened, 2 mg) was added and the reaction mixture was purged with $H_2$ (sparging using a needle and balloon). Diethylamine (6 µL, 0.056 mmol) was added and the reaction mixture was further stirred under $H_2$ atmosphere (1 atm, balloon). After four hours, TLC indicated that most starting material was consumed. After 15 h, the RM was filtered and the solids rinsed with MeOH. Filtrate was concentrated, re-dissolved in $CH_2Cl_2$ and washed with 2 M aq. $Na_2CO_3$ solution. The $CH_2Cl_2$ solution was dried over $Na_2SO_4$, filtered, and concentrated to yield ibogamine as a pale yellow amorphous solid (13 mg, quantitative). Spectral characteristics were in agreement with reported literature data[12]. For *in vivo* experiments, ibogamine was transformed into the hydrochloride salt form. The free base was dissolved in MeOH, acidified with aq. HCl (12.1 M) to pH 1–2 and evaporated to dryness.

**$^1$H NMR (500 MHz, $CDCl_3$)** δ 7.63 (s, 1H), 7.48 (dd, *J* = 7.2, 1.6 Hz, 1H), 7.28 – 7.24 (m, 1H), 7.15 – 7.07 (m, 2H), 3.42 – 3.33 (m, 2H), 3.20 – 3.11 (m, 1H), 3.08 (dt, *J* = 9.3, 2.3 Hz, 1H), 2.98 (dt, *J* = 9.3, 3.0 Hz, 1H), 2.96 – 2.90 (m, 1H), 2.89 – 2.85 (m, 1H), 2.72 – 2.66 (m, 1H), 2.09 – 2.01 (m, 1H), 1.88 – 1.83 (m, 1H), 1.83 – 1.78 (m, 1H), 1.69 – 1.63 (m, 1H), 1.59 – 1.53 (m, 2H), 1.51 – 1.43 (m, 1H), 1.26 – 1.19 (m, 1H), 0.91 (t, *J* = 7.1 Hz, 3H). **$^{13}$C NMR (126 MHz, $CDCl_3$)** δ 141.97, 134.77, 129.86, 121.08, 119.23, 118.04, 110.21, 109.37, 57.71, 54.30, 50.07, 42.11, 41.63, 34.34, 32.27, 27.95, 26.66, 20.80, 12.07. **Hydrochloride: $^1$H NMR (400 MHz, $D_2O$)** δ 7.61 – 7.54 (m, 1H), 7.46 (dq, *J* = 8.0, 0.9 Hz, 1H), 7.31 – 7.24 (m, 1H), 7.24 – 7.18 (m, 1H), 3.54 (dt, *J* = 13.7, 4.4 Hz, 1H), 3.45 – 3.23 (m, 4H), 3.20 – 2.96 (m, 3H), 2.32 – 2.19 (m, 1H), 2.19 – 2.04 (m, 2H), 2.04 – 1.90 (m, 1H), 1.56 (p, *J* = 7.4 Hz, 2H), 1.46 – 1.37 (m, 1H), 1.37 – 1.27 (m, 1H), 1.01 (t, *J* = 7.3 Hz, 3H). **LRMS (APCI+)** calcd. for $C_{19}H_{25}N_2^+$ $[M+H]^+$ 281.2, found 281.5.

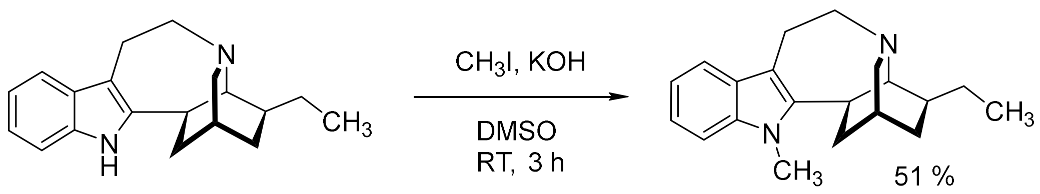


##### *N*-methyl-ibogamine

Ibogamine (50 mg, 0.18 mmol) and KOH (20 mg, 0.36 mmol) were mixed in DMSO (anhydrous, 0.3 mL). $CH_3I$ (17 µL, 0.27 mmol) was added into the suspension and reaction mixture was stirred at RT for three hours. Reaction was quenched by adding $H_2O$ (15 mL) and the mixture was extracted with $CH_2Cl_2$ (3 × 5 mL). Combined extracts were dried over $Na_2SO_4$, filtered, and concentrated under reduced pressure. Crude material was purified by PTLC (1:60 AcOEt in hexanes + 2% $Et_3N$, developed twice). The product was isolated as a yellow waxy solid (27 mg, 51%).

For *in vivo* experiments, *N*-methyl-ibogamine was transformed to the hydrochloride salt form. The free base was dissolved in MeOH, acidified with aq. HCl (12.1 M) to pH 1–2 and evaporated to dryness.

**$^1$H NMR (500 MHz, $CDCl_3$)** δ 7.52 (d, *J* = 7.8 Hz, 1H), 7.30 – 7.25 (m, 1H), 7.24 – 7.17 (m, 1H), 7.15 – 7.07 (m, 1H), 3.68 (s, 3H), 3.48 – 3.34 (m, 2H), 3.27 – 3.20 (m, 1H), 3.20 – 3.14 (m, 1H), 3.12 – 3.02 (m, 2H), 2.93 (s, 1H), 2.85 – 2.78 (m, 1H), 2.16 – 2.07 (m, 1H), 1.94 – 1.84 (m, 2H), 1.71 – 1.46 (m, 4H), 1.38 – 1.24 (m, 1H), 0.97 (t, *J* = 7.2 Hz, 3H). **$^{13}$C NMR (126 MHz, $CDCl_3$)** δ 143.2, 136.2, 128.3, 120.8, 118.8, 118.1, 108.9, 108.8, 57.5, 54.4, 50.4, 42.5, 38.6, 33.7, 32.2, 29.8, 27.9, 26.6, 21.1, 12.1. **Hydrochloride: $^1$H NMR (400 MHz, MeOD)** δ 7.47 (d, *J* = 7.9 Hz, 1H), 7.32 (d, *J* = 8.2 Hz, 1H), 7.17 (ddd, *J* = 8.3, 7.2, 1.1 Hz, 1H), 7.06 (ddd, *J* = 7.9, 7.1, 0.9 Hz, 1H), 3.78 – 3.65 (m, 6H), 3.65 – 3.59 (m, 1H), 3.49 (dt, *J* = 12.3, 2.6 Hz, 1H), 3.41 (d, *J* = 12.0 Hz, 1H), 3.29 – 3.25 (m, 1H), 2.43 (t, *J* = 12.8 Hz, 1H), 2.25 – 2.09 (m, 3H), 1.76 – 1.62 (m, 3H), 1.50 – 1.41 (m, 1H), 1.07 (t, *J* = 7.3 Hz, 3H). **HRMS (ESI+)** calcd. for $C_{20}H_{27}N_2^+$ $[M+H]^+$ 295.2174, found 295.2194.

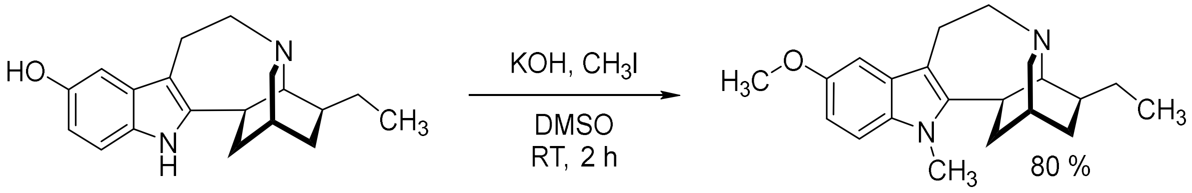


##### *N*-methyl-ibogaine

Noribogaine (200 mg, 0.67 mmol) and KOH (150 mg, 2.7 mmol) were combined in DMSO (anhydrous, 2.0 mL). After stirring for one hour at RT, $CH_3I$ (125 µL, 2.0 mmol) was added in one portion and the stirring continued. Reaction mixture was poured into $H_2O$ after one hour and extracted with $Et_2O$ (3 × 15 mL). Combined extracts were washed with $H_2O$ and brine, dried over $Na_2SO_4$, filtered, and concentrated under reduced pressure. Crude material was purified by column chromatography (1:20, AcOEt in hexanes + 2% $Et_3N$). The product was isolated as a pale-yellow viscous oil (174 mg, 80%). Spectral characteristics were in agreement with reported literature data[13].

**$^1$H NMR (500 MHz, $CDCl_3$)** δ 7.11 (d, *J* = 8.7 Hz, 1H), 6.94 (d, *J* = 2.4 Hz, 1H), 6.82 (dd, *J* = 8.7, 2.4 Hz, 1H), 3.86 (s, 3H), 3.61 (s, 3H), 3.39 – 3.28 (m, 2H), 3.23 – 3.12 (m, 1H), 3.12 – 3.02 (m, 2H), 2.98

(dt, $J$ = 9.4, 2.9 Hz, 1H), 2.85 (t, $J$ = 1.6 Hz, 1H), 2.76 – 2.64 (m, 1H), 2.11 – 1.99 (m, 1H), 1.91 – 1.79 (m, 2H), 1.65 – 1.52 (m, 3H), 1.52 – 1.42 (m, 1H), 1.29 – 1.19 (m, 1H), 0.91 (t, $J$ = 7.2 Hz, 3H). **$^{13}$C NMR (126 MHz, $CDCl_3$)** δ 153.9, 144.1, 131.6, 128.5, 110.6, 109.5, 108.6, 100.5, 57.5, 56.3, 54.4, 50.5, 42.5, 38.8, 33.8, 32.2, 29.9, 28.0, 26.6, 21.2, 12.1. **LRMS (APCI+)** calcd. for $C_{21}H_{29}N_2O^+$ $[M+H]^+$ 325.2, found 325.2.

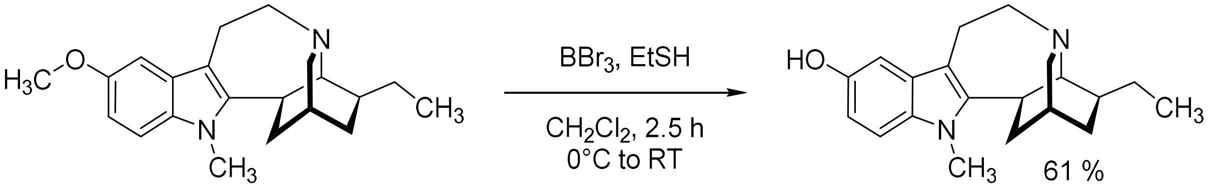


##### *N*-methyl-noribogaine

*N*-methyl-ibogaine (80 mg, 0.25 mmol) was dissolved in $CH_2Cl_2$ (anhydrous, 1.6 mL) and cooled in ice bath. EtSH (0.079 mL, 1.1 mmol) and $BBr_3$ (1 M in $CH_2Cl_2$, 0.37 mL) were added, and the reaction mixture was further stirred at RT for 1.5 h – significant amount of starting material remained. Additional EtSH (0.027 mL, 1.1 mmol) and $BBr_3$ (1 M in $CH_2Cl_2$, 0.12 mL) were added, reaction was stirred for one hour more and quenched by pouring into sat. aq. $NaHCO_3$ solution (20 mL). The mixture was extracted with $CH_2Cl_2$ (3 × 10 mL), combined extracts were dried over $Na_2SO_4$, filtered, and concentrated under reduced pressure. Crude material was pre-purified by column chromatography (gradient of 1:4 to 1:2 AcOEt in hexanes + 2% $Et_3N$), isolated fraction was evaporated from a mixture of MeOH/$H_2O$ to remove excess $Et_3N$. The free base was further dissolved in MeOH, acidified with aq. HCl (12.1 M) to pH 1–2, evaporated to dryness and repeatedly washed with $CH_3CN$ (3 × 1 mL, solid was sedimented by centrifugation and solvent decanted). Product was isolated as a pale yellow amorphous solid (52 mg, 61%) as a hydrochloride salt.

**$^1$H NMR (500 MHz, $CDCl_3$)** δ 7.06 (dd, $J$ = 8.6, 0.5 Hz, 1H), 6.88 (dd, $J$ = 2.5, 0.5 Hz, 1H), 6.72 (dd, $J$ = 8.6, 2.4 Hz, 1H), 4.44 (s, 1H), 3.60 (s, 3H), 3.38 – 3.25 (m, 2H), 3.18 – 3.10 (m, 1H), 3.10 – 3.05 (m, 1H), 3.03 (dt, $J$ = 9.3, 2.3 Hz, 1H), 2.98 (dt, $J$ = 9.4, 2.9 Hz, 1H), 2.85 (t, $J$ = 1.7 Hz, 1H), 2.68 – 2.57 (m, 1H), 2.09 – 2.01 (m, 1H), 1.88 – 1.79 (m, 2H), 1.65 – 1.51 (m, 4H), 1.51 – 1.41 (m, 1H), 1.27 – 1.19 (m, 1H), 0.91 (t, $J$ = 7.3 Hz, 3H). **$^{13}$C NMR (126 MHz, $CDCl_3$)** δ 149.3, 144.1, 131.5, 128.7, 110.2, 109.3, 108.2, 102.8, 57.4, 54.4, 50.3, 42.4, 38.5, 33.5, 31.9, 29.8, 27.8, 26.4, 20.9, 12.0. **Hydrochloride: $^1$H NMR (500 MHz, MeOD)** δ 7.15 (d, $J$ = 8.7 Hz, 1H), 6.84 (d, $J$ = 2.3 Hz, 1H), 6.72 (dd, $J$ = 8.7, 2.3 Hz, 1H), 3.74 – 3.67 (m, 2H), 3.67 – 3.60 (m, 4H), 3.59 – 3.53 (m, 1H), 3.50 – 3.37 (m, 2H), 3.28 – 3.11 (m, 2H), 2.44 – 2.36 (m, 1H), 2.24 – 2.19 (m, 1H), 2.19 – 2.08 (m, 2H), 1.75 – 1.58 (m, 3H), 1.47 – 1.40 (m, 1H), 1.07 (t, $J$ = 7.3 Hz, 3H). **$^{13}$C NMR (126 MHz, MeOD)** δ 152.0, 141.2, 132.9, 129.1, 112.6, 110.7, 107.6, 103.1, 60.8, 58.4, 53.1, 40.3, 34.2, 31.8, 29.9, 29.9, 27.5, 25.4, 19.8, 12.0. **HRMS (ESI+)** calcd. for $C_{20}H_{27}N_2O^+$ $[M+H]^+$ 311.2123, found 311.2124.

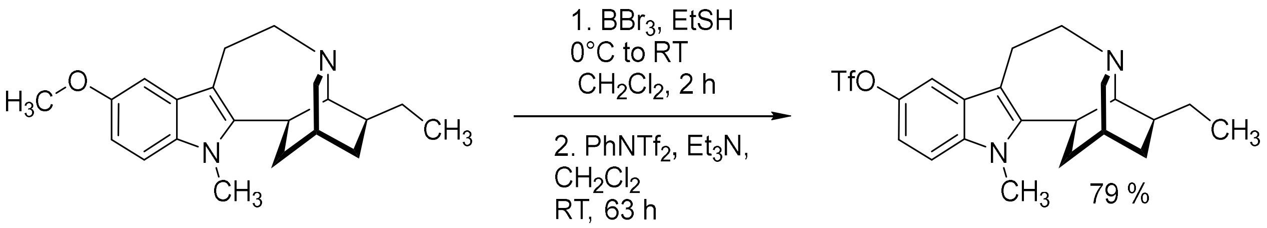


##### ***N*-methyl-ibogamine-5-triflate**

*N*-Methyl-ibogaine (312 mg, 0.96 mmol) was dissolved in $CH_2Cl_2$ (anhydrous, 6.5 mL) and cooled in ice bath. EtSH (0.32 mL, 4.4 mmol) and $BBr_3$ (1 M in $CH_2Cl_2$, 1.4 mL) were added, the reaction mixture was further stirred at RT for two hours and quenched by pouring into sat. aq. $NaHCO_3$ solution (30 mL). The mixture was extracted with $CH_2Cl_2$ (3 × 10 mL), combined extracts were dried over $Na_2SO_4$, filtered, and concentrated under reduced pressure. Crude material was dried on high vacuum for 4 h and used for the next step without further purification.

Crude *N*-methyl-noribogaine and bis(trifluoromethanesulfonyl)aniline (343 mg, 0.96 mmol) were dissolved in $CH_2Cl_2$ (anhydrous, 5 mL) and $Et_3N$ (0.2 mL, 1.4 mmol) was added. After stirring at RT for 63 h, reaction was quenched by pouring into 2 M aq. $Na_2CO_3$ solution (20 mL) and the mixture was extracted with $CH_2Cl_2$ (3 × 10 mL). Combined extracts were dried over $Na_2SO_4$, filtered, and concentrated under reduced pressure. Crude material was purified by column chromatography (basic alumina, hexanes, and hexanes + 2% AcOEt). The product was isolated as a pale-yellow viscous oil (334 mg, 79% over two steps).

**$^1$H NMR (500 MHz, $CDCl_3$)** δ 7.33 (d, *J* = 2.4 Hz, 1H), 7.20 (d, *J* = 8.8 Hz, 1H), 7.03 (dd, *J* = 8.8, 2.5 Hz, 1H), 3.65 (s, 3H), 3.39 – 3.30 (m, 2H), 3.22 – 3.12 (m, 1H), 3.12 – 3.07 (m, 1H), 3.05 – 2.98 (m, 2H), 2.88 – 2.83 (m, 1H), 2.69 – 2.63 (m, 1H), 2.13 – 2.04 (m, 1H), 1.92 – 1.86 (m, 1H), 1.86 – 1.80 (m, 1H), 1.66 – 1.53 (m, 3H), 1.52 – 1.42 (m, 1H), 1.28 – 1.21 (m, 1H), 0.92 (t, *J* = 7.2 Hz, 3H). **$^{19}$F NMR (471 MHz, $CDCl_3$)** δ -72.8. **$^{13}$C NMR (126 MHz, $CDCl_3$)** δ 146.1, 143.4, 135.1, 128.6, 122.9, 120.3, 117.8, 115.2, 113.7, 110.6, 110.0, 109.6, 57.3, 54.1, 50.5, 42.5, 38.9, 33.6, 32.1, 30.1, 28.0, 26.6, 21.0, 12.1. **LRMS** (APCI+) calcd. for $C_{21}H_{26}F_3N_2O_3S^+$ $[M+H]^+$ 443.2, found 443.1.

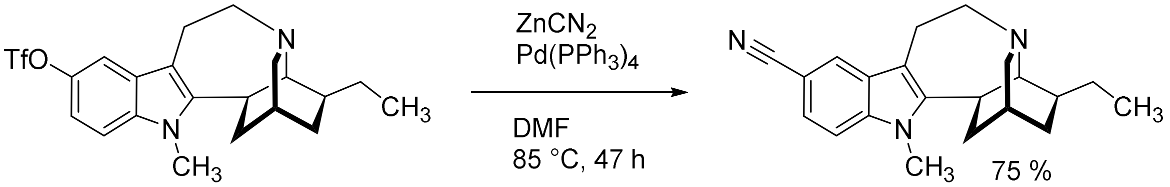


##### ***N*-methyl-5-cyano-ibogamine**

*N*-methyl-ibogamine-5-triflate (44 mg, 0.1 mmol), $ZnCN_2$ (24 mg, 0.2 mmol), $Pd(PPh_3)_4$ (5.8 mg, 0.005 mmol) were balanced into a reaction vial and dried under vacuum. DMF (anhydrous, 0.5 mL) was added, vial was flushed with argon, closed with a Teflon lined solid screw cap and the RM heated to 85 °C. After 24 h, TLC indicated only partial conversion. Additional $ZnCN_2$ (47 mg, 0.4 mmol) and $Pd(PPh_3)_4$ (12 mg, 0.01 mmol) were added and heating was continued for an additional 23 h, until TLC indicated that no more SM remained. RM was diluted with $H_2O$ (10 mL) and sat. aq. $NaHCO_3$ solution (2 mL), mixture was extracted with $Et_2O$ (3 × 5 mL) and AcOEt (3 × 5 mL), extraction is complicated by the presence of insoluble solid precipitates. Combined extracts were washed with $H_2O$, brine, dried over $Na_2SO_4$, filtered, and concentrated. Crude material was purified by column chromatography (gradient of 0%, 5% and 10% of AcOEt in hexanes + 2% $Et_3N$), followed by PTLC (10% of AcOEt in hexanes + 2% $Et_3N$). The product was isolated as a white foamy solid (24 mg, 75%).

**$^1$H NMR (500 MHz, $CDCl_3$)** δ 7.78 (d, *J* = 1.5 Hz, 1H), 7.36 (dd, *J* = 8.5, 1.6 Hz, 1H), 7.23 (d, *J* = 8.5 Hz, 1H), 3.66 (s, 3H), 3.39 – 3.29 (m, 2H), 3.20 – 3.07 (m, 2H), 3.03 – 2.98 (m, 2H), 2.85 (t, *J* = 1.6 Hz, 1H), 2.73 – 2.66 (m, 1H), 2.12 – 2.05 (m, 1H), 1.91 – 1.80 (m, 2H), 1.64 – 1.51 (m, 3H), 1.51 – 1.44 (m, 1H), 1.27 – 1.21 (m, 1H), 0.91 (t, *J* = 7.2 Hz, 3H). **$^{13}$C NMR (126 MHz, $CDCl_3$)** δ 145.8, 137.8, 128.3, 123.8, 123.5, 121.4, 110.1, 109.5, 101.5, 57.2, 53.9, 50.3, 42.4, 38.7, 33.6, 32.1, 30.1, 27.9, 26.5, 20.9, 12.0. **HRMS (ESI+)** calcd. for $C_{21}H_{26}N_3^+$ [M+H]$^+$ 320.2127, found 320.2136.

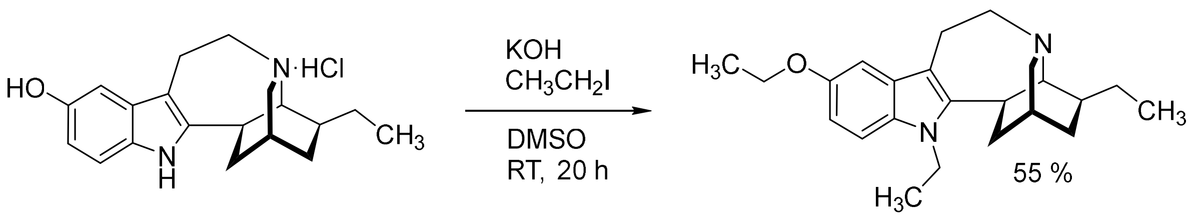


##### *N*-ethyl-5-ethoxy-ibogamine

Noribogaine hydrochloride (50 mg, 0.15 mmol) and KOH (53 mg, 0.94 mmol) were mixed in DMSO (anhydrous, 0.3 mL). $CH_3CH_2I$ (36 μL, 0.45 mmol) was added into the suspension and the resulting mixture was stirred at RT for 20 h. Reaction was quenched by adding $H_2O$ (15 mL) and the mixture was extracted with $CH_2Cl_2$ (3 × 5 mL). Combined extracts were dried over $Na_2SO_4$, filtered, and concentrated under reduced pressure. Crude material was purified by PTLC (1:60, AcOEt in hexanes + 2% $Et_3N$, developed twice). The product was isolated as a yellow waxy solid (29 mg, 55%).

**$^1$H NMR (500 MHz, $CDCl_3$)** δ 7.13 (d, *J* = 8.7 Hz, 1H), 6.96 (d, *J* = 2.4 Hz, 1H), 6.82 (dd, *J* = 8.7, 2.4 Hz, 1H), 4.28 – 3.90 (m, 4H), 3.45 – 3.28 (m, 2H), 3.21 – 3.12 (m, 1H), 3.11 – 3.07 (m, 1H), 3.05 – 2.98 (m, 2H), 2.86 (s, 1H), 2.69 – 2.62 (m, 1H), 2.13 – 2.05 (m, 1H), 1.94 – 1.78 (m, 2H), 1.66 – 1.55 (m, 3H), 1.53 – 1.42 (m, 4H), 1.31 (t, *J* = 7.2 Hz, 3H), 1.28 – 1.22 (m, 1H), 0.93 (t, *J* = 7.2 Hz, 3H). **$^{13}$C NMR (126 MHz, $CDCl_3$)** δ 153.1, 143.5, 130.3, 128.7, 111.2, 109.5, 108.9, 101.9, 64.6, 57.4, 54.5, 50.5, 42.8, 38.6, 37.9, 34.3, 32.1, 28.2, 26.7, 21.0, 16.1, 15.3, 12.1. **LRMS (APCI+)** calcd. for $C_{23}H_{33}N_2O^+$ [M+H]$^+$ 353.3, found 353.8.

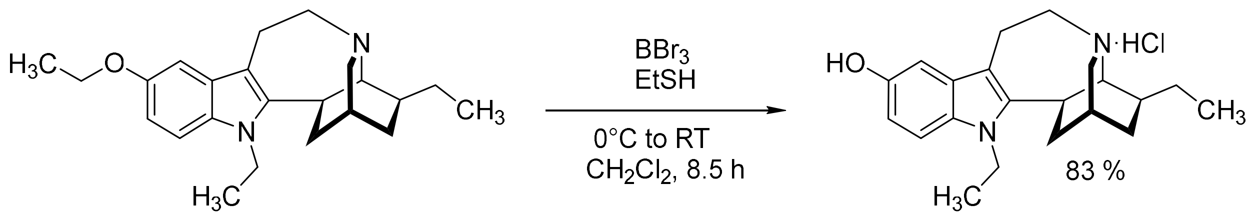

#### *N*-ethyl-noribogaine

*N*-ethyl-5-ethoxy-ibogamine (68 mg, 0.19 mmol) was dissolved in $CH_2Cl_2$ (anhydrous, 1.5 mL) and cooled in an ice bath. EtSH (0.1 mL, 1.4 mmol) and $BBr_3$ (1 M in $CH_2Cl_2$, 0.48 mL) were added, the reaction mixture was further stirred at RT for 8.5 h and quenched by pouring into sat. aq. $NaHCO_3$ solution (10 mL). The mixture was extracted with $CH_2Cl_2$ (3 × 5 mL), combined extracts were dried over $Na_2SO_4$, filtered, and concentrated under reduced pressure. Crude material was purified by column chromatography (gradient of 1:4, 1:3 to 1:2 AcOEt in hexanes + 2% $Et_3N$), excess $Et_3N$ was removed by repeated evaporation from MeOH/$H_2O$ mixture. Due to the continued presence of an impurity, the free base was transformed to hydrochloride salt by treating its MeOH solution with HCl (36 % aq.). The hydrochloride salt was purified by PTLC (5% MeOH in $CH_2Cl_2$ + 0.1% HCl (36% aq.), developed twice). The product was isolated as a yellow amorphous solid (57 mg, 83%).

**Hydrochloride: $^1$H NMR (500 MHz, MeOD)** δ 7.06 (d, *J* = 8.6 Hz, 1H), 6.78 (d, *J* = 2.0 Hz, 1H), 6.64 (dd, *J* = 8.6, 2.0 Hz, 1H), 4.15 – 3.96 (m, 2H), 3.59 – 3.47 (m, 3H), 3.42 – 3.36 (m, 2H), 3.22 (d, *J* = 12.3 Hz, 1H), 3.12 – 2.98 (m, 2H), 2.31 (t, *J* = 12.7 Hz, 1H), 2.11 – 2.02 (m, 3H), 1.68 – 1.57 (m, 1H), 1.57 – 1.51 (m, 1H), 1.51 – 1.44 (m, 1H), 1.36 – 1.27 (m, 1H), 1.19 (t, *J* = 7.0 Hz, 3H), 0.97 (t, *J* = 7.2 Hz, 3H). **$^{13}$C NMR (126 MHz, MeOD)** δ 151.6, 140.5, 131.3, 129.1, 112.2, 110.8, 107.7, 103.0, 60.7, 58.2, 52.9, 40.1, 38.6, 33.7, 32.1, 29.6, 27.5, 25.0, 19.4, 16.1, 11.8. **HRMS** (ESI+) calcd. for $C_{21}H_{29}N_2O^+$ $[M+H]^+$ 325.2280, found 325.2285.

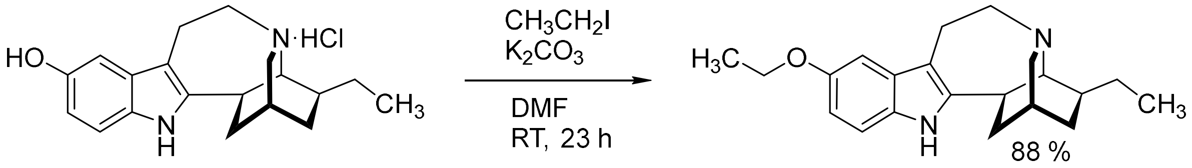

#### 5-ethoxy-ibogamine

Noribogaine hydrochloride (166 mg, 0.5 mmol) and $K_2CO_3$ (276 mg, 2.0 mmol) were combined in DMF (anhydrous, 2.0 mL) and $CH_3CH_2I$ (80 µL, 1.0 mmol) was added. The reaction mixture was further stirred 23 h under argon atmosphere and quenched by pouring into $H_2O$ (20 mL). Aqueous mixture was extracted with $Et_2O$ (3 × 10 mL), combined extracts were washed with $H_2O$ (10 mL), brine (10 mL) and dried over $Na_2SO_4$. Crude material was purified by column chromatography (gradient of 0%, 10%, 20% and 25% $Et_2O$ in hexanes + 2% $Et_3N$). 10-Ethoxy-ibogamine was obtained as a thick colorless oil (143 mg, 88%). For biological studies, this compound was transformed to hydrochloride salt using 2M HCl in $Et_2O$, precipitated solid was collected by filtration, washed with $Et_2O$, hexanes and dried under reduced pressure.

**$^1$H NMR (500 MHz, $CDCl_3$)** δ 7.55 (s, 1H), 7.13 (d, *J* = 8.6 Hz, 1H), 6.95 (d, *J* = 2.4 Hz, 1H), 6.78 (dd, *J* = 8.6, 2.4 Hz, 1H), 4.09 (q, *J* = 7.0 Hz, 2H), 3.42 – 3.28 (m, 2H), 3.20 – 3.04 (m, 2H), 2.98 (dt, *J* =

9.4, 3.0 Hz, 1H), 2.94 – 2.81 (m, 2H), 2.66 – 2.55 (m, 1H), 2.09 – 1.98 (m, 1H), 1.90 – 1.74 (m, 2H), 1.69 – 1.61 (m, 1H), 1.61 – 1.51 (m, 2H), 1.45 (q, *J* = 8.4, 7.0 Hz, 4H), 1.25 – 1.17 (m, 1H), 0.91 (t, *J* = 7.0 Hz, 3H). **$^{13}$C NMR (126 MHz, $CDCl_3$)** δ 153.3, 143.0, 130.3, 129.9, 111.4, 110.9, 109.2, 101.8, 64.5, 57.6, 54.3, 50.1, 42.1, 41.7, 34.3, 32.2, 28.0, 26.6, 20.8, 15.3, 12.1. **Hydrochloride: $^1$H NMR (500 MHz, MeOD)** δ 7.17 (d, *J* = 8.7 Hz, 1H), 6.95 (d, *J* = 2.3 Hz, 1H), 6.75 (dd, *J* = 8.7, 2.3 Hz, 1H), 4.05 (q, *J* = 7.0 Hz, 2H), 3.71 (dt, *J* = 13.4, 4.2 Hz, 1H), 3.66 – 3.56 (m, 2H), 3.42 (s, 2H), 3.38 (ddd, *J* = 12.0, 4.7, 1.9 Hz, 1H), 3.29 – 3.15 (m, 2H), 2.32 (t, *J* = 12.7 Hz, 1H), 2.21 – 2.12 (m, 2H), 2.06 – 1.98 (m, 1H), 1.77 – 1.63 (m, 3H), 1.39 (t, *J* = 7.0 Hz, 4H), 1.05 (t, *J* = 7.3 Hz, 3H). **$^{13}$C NMR (126 MHz, MeOD)** δ 154.5, 140.4, 131.9, 129.9, 113.2, 112.5, 107.3, 102.2, 65.3, 61.6, 57.4, 51.9, 40.3, 36.4, 32.6, 30.1, 27.3, 25.2, 19.4, 15.4, 11.9.

**HRMS** (ESI+) calcd. for $C_{21}H_{29}N_2O^+$ $[M+H]^+$ 325.2280, found 325.2280**.**

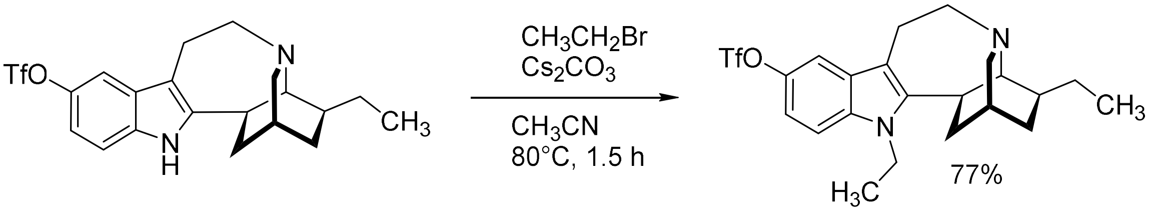


##### *N*-ethyl-ibogamine-5-triflate

Ibogamine-10-triflate (67 mg, 0.156 mmol) and $Cs_2CO_3$ (5 equiv., 254 mg, 0.78 mmol) were placed in a reaction vial, $CH_3CN$ (3.1 mL, 0.05M) and $CH_3CH_2Br$ (4 equiv., 47 µL, 0.62 mmol) were sequentially added. Vial was closed with solid screw cap and the content was stirred while heating to 80°C for 1.5h. After cooling to RT, mixture was poured into $H_2O$, extracted with $CH_2Cl_2$ (3×), combined extracts were dried over $Na_2SO_4$, filtered and concentrated. Crude material was purified by column chromatography (gradient of 0% to 5%, AcOEt in hexanes + 2% $Et_3N$). 10-Ethoxy-ibogamine was obtained as a yellow foamy solid (55 mg, 77%).

**$^1$H NMR (500 MHz, $CDCl_3$)** δ 7.34 (d, *J* = 2.4 Hz, 1H), 7.21 (d, *J* = 8.8 Hz, 1H), 7.02 (dd, *J* = 8.8, 2.5 Hz, 1H), 4.19 – 4.03 (m, *J* = 7.4 Hz, 2H), 3.43 – 3.31 (m, 2H), 3.16 (ddd, *J* = 14.0, 12.7, 3.5 Hz, 1H), 3.10 – 2.99 (m, 3H), 2.86 (s, 1H), 2.64 (dt, *J* = 16.9, 3.1 Hz, 1H), 2.12 (ddt, *J* = 14.3, 11.5, 2.8 Hz, 1H), 1.94 – 1.82 (m, 2H), 1.69 – 1.55 (m, 3H), 1.55 – 1.42 (m, 1H), 1.33 (t, *J* = 7.2 Hz, 3H), 1.29 – 1.21 (m, 1H), 0.92 (t, *J* = 7.3 Hz, 3H**). $^{13}$C NMR (126 MHz, $CDCl_3$)** δ 145.6, 143.4, 133.8, 128.8, 122.9, 120.3, 117.8, 115.2, 113.5, 110.5, 110.3, 109.6, 57.3, 54.3, 50.5, 42.7, 38.6, 38.2, 34.2, 32.0, 28.2, 26.6, 20.9, 15.9, 12.0. **$^{19}$F NMR (470 MHz, $CDCl_3$)** δ -72.82. **LRMS (APCI+)** calcd. for $C_{22}H_{28}N_2O_3S^+$ $[M+H]^+$ 457.2, found 457.2.

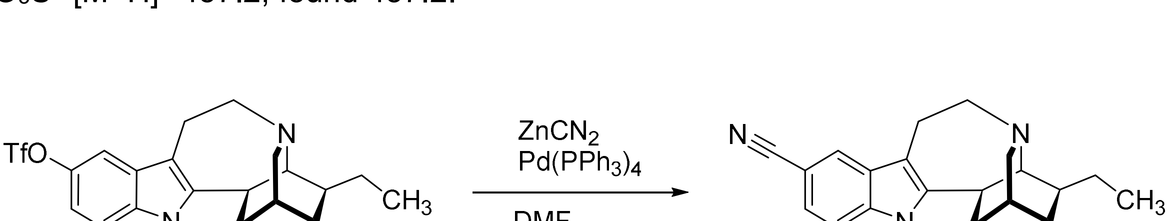


##### *N*-ethyl-5-cyano-ibogamine

*N*-ethyl-ibogamine-5-triflate (54 mg, 0.118 mmol), $ZnCN_2$ (28 mg, 0.235 mmol), $Pd(PPh_3)_4$ (6.8 mg, 0.006 mmol) were added into a reaction vial and dried under vacuum. DMF (anhydrous, 0.6 mL) was added, vial was flushed with argon, closed with a Teflon lined solid screw cap and the RM heated to 85 °C. After 19 h, TLC indicated only partial conversion. Additional $ZnCN_2$ (28 mg, 0.235 mmol) and $Pd(PPh_3)_4$ (6.8 mg, 0.006 mmol) were added and heating was continued for an additional 26 h. TLC still indicated only partial conversion, additional $ZnCN_2$ (55 mg, 0.47 mmol) and $Pd(PPh_3)_4$ (13.6 mg, 0.012 mmol) were added and heating was continued for a total of 72 h, with no improvement in conversion observed. After cooling to RT, mixture was filtered through plug of celite (eluted using AcOEt + 2% $Et_3N$) and concentrated under reduced pressure. Crude material was purified by preparative TLC (using 5% of AcOEt in hexanes + 2% $Et_3N$). The product was isolated as an off-white foamy solid (10 mg, 25%).

**$^1$H NMR (500 MHz, $CDCl_3$)** δ 7.79 (d, *J* = 1.7 Hz, 1H), 7.36 (dd, *J* = 8.4, 1.4 Hz, 1H), 7.25 (d, *J* = 8.5 Hz, 1H), 4.12 (ddt, *J* = 19.1, 15.0, 7.5 Hz, 2H), 3.36 (td, *J* = 15.7, 4.7 Hz, 2H), 3.17 (d, *J* = 15.3 Hz, 1H), 3.05 (dt, *J* = 9.5, 2.5 Hz, 3H), 2.92 – 2.78 (m, 1H), 2.67 (d, *J* = 13.4 Hz, 1H), 2.20 – 2.07 (m, 1H), 1.90 (s, 2H), 1.68 – 1.54 (m, 3H), 1.48 (q, *J* = 7.8 Hz, 1H), 1.33 (t, *J* = 7.3 Hz, 3H), 1.29 – 1.22 (m, 1H), 0.91 (t, *J* = 7.3 Hz, 3H). **$^{13}$C NMR (126 MHz, $CDCl_3$)** δ 145.3, 136.7, 128.5, 123.8, 123.7, 121.4, 110.6, 109.5, 101.6, 57.3, 54.2, 50.5, 42.7, 38.5, 38.2, 34.2, 32.0, 28.1, 26.6, 20.7, 15.9, 12.1. **LRMS (APCI+)** calcd. for $C_{22}H_{28}N_3^+$ $[M+H]^+$ 334.2, found 334.3.

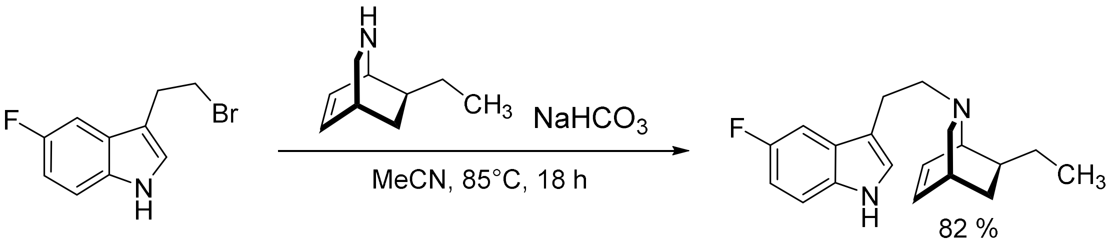


**(1S,4R,7S)-7-ethyl-2-(2-(5-fluoro-1H-indol-3-yl)ethyl)-2-azabicyclo[2.2.2]oct-5-ene**

The isoquinuclidine (1S,4R,7S)-7-ethyl-2-azabicyclo[2.2.2]oct-5-ene (283 mg, 2.07 mmol. 1 eq.) was treated with 3-(2-bromoethyl)-5-fluoro-1H-indole[14] (500 mg, 2.07 mmol, 1 eq.) and $NaHCO_3$ (694 mg, 8.26 mmol, 4 eq.) and refluxed in $CH_3CN$ (10 mL, 0.2 M) until the starting material was consumed (usually around 18 h). After quenching with water, the mixture was extracted with AcOEt (30 mL × 3). The combined organic layers were washed with water, brine, dried ($Na_2SO_4$), and concentrated to afford the crude product which was purified by flash column chromatography (using 10% of AcOEt in hexanes + 2% $Et_3N$). The product was isolated as a light-yellow oil (505 mg, 82%).

**$^1$H NMR (500 MHz, $CDCl_3$)** δ 8.10 (s, 1H), 7.26 (td, *J* = 9.5, 3.5 Hz, 2H), 7.08 (d, *J* = 2.4 Hz, 1H), 6.96 (td, *J* = 9.1, 2.6 Hz, 1H), 6.43 – 6.32 (m, 2H), 3.33 (dt, *J* = 5.3, 1.9 Hz, 1H), 3.17 (dd, *J* = 9.2, 2.4 Hz, 1H), 2.94 – 2.76 (m, 3H), 2.60 – 2.46 (m, 2H), 2.02 (dt, *J* = 9.2, 2.6 Hz, 1H), 1.72 – 1.48 (m, 3H), 1.42 – 1.31 (m, 1H), 1.06 – 0.90 (m, 4H). **$^{13}$C NMR (126 MHz, $CDCl_3$)** δ 158.66, 156.80, 133.26, 132.77, 132.51, 128.16, 128.09, 123.64, 114.99, 111.78, 111.70, 110.24, 110.03, 103.91, 103.72, 58.87, 56.25, 56.06, 41.10, 31.56, 29.70, 27.29, 24.22, 12.63. **$^{19}$F NMR (470 MHz, $CDCl_3$)** δ -125.17. **LRMS (EI)** calcd. for $C_{19}H_{23}FN_2$ $[M^+]$ 298.04, found 298.17.

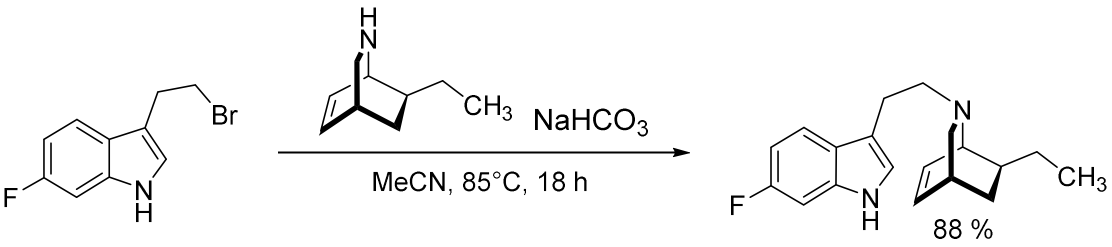

**(1S,4R,7S)-7-ethyl-2-(2-(6-fluoro-1H-indol-3-yl)ethyl)-2-azabicyclo[2.2.2]oct-5-ene**

The isoquinuclidine (1S,4R,7S)-7-ethyl-2-azabicyclo[2.2.2]oct-5-ene (283 mg, 2.07 mmol. 1 eq.) was treated with 3-(2-bromoethyl)-6-fluoro-1H-indole[15] (500 mg, 2.07 mmol, 1 eq.) and $NaHCO_3$ (694 mg, 8.26 mmol, 4 eq.) and refluxed in $CH_3CN$ (10 mL, 0.2 M) until the starting material was consumed (usually around 18 h). After quenching with water, the mixture was extracted with AcOEt (30 mL × 3). The combined organic layers were washed with water, brine, dried ($Na_2SO_4$), and concentrated to afford the crude product which was purified by flash column chromatography (using 10% of AcOEt in hexanes + 2% $Et_3N$). The product was isolated as a light-yellow oil (545 mg, 88**%).**

**$^1$H NMR (500 MHz, $CDCl_3$)** δ 8.09 (s, 1H), 7.50 (dd, *J* = 8.7, 5.3 Hz, 1H), 7.05 – 6.96 (m, 2H), 6.90 (ddd, *J* = 9.7, 8.7, 2.3 Hz, 1H), 6.43 – 6.31 (m, 2H), 3.33 (dt, *J* = 5.4, 1.8 Hz, 1H), 3.16 (dd, *J* = 9.2, 2.3 Hz, 1H), 2.92 – 2.76 (m, 3H), 2.62 – 2.45 (m, 2H), 2.01 (dt, *J* = 9.2, 2.6 Hz, 1H), 1.72 – 1.49 (m, 3H), 1.41 – 1.28 (m, 1H), 1.03 – 0.91 (m, 4H). **$^{13}$C NMR (126 MHz, $CDCl_3$)** δ 160.22, 158.34, 135.43, 135.33, 132.28, 132.01, 123.70, 121.15, 121.12, 118.94, 118.86, 114.44, 107.18, 106.99, 96.74, 96.53, 58.26, 55.54, 55.29, 40.49, 30.95, 29.11, 26.59, 23.61, 11.90. **$^{19}$F NMR (470 MHz, $CDCl_3$)** δ -121.76. **LRMS (EI)** calcd. for $C_{19}H_{23}FN_2$ [$M^+$] 298.04, found 298.05.

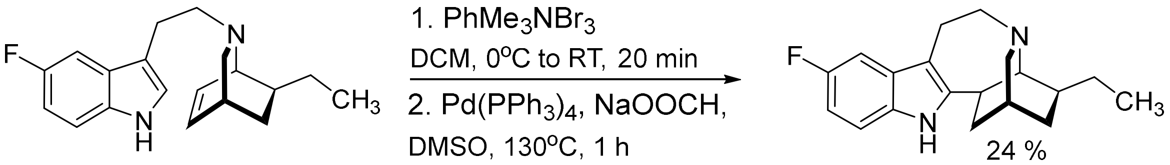

**5-fluoro-ibogamine**

Synthesized utilizing procedures found in our prior publication.[14] To a solution of the substrate (930 mg, 3.12 mmol, 1 eq.) in anhydrous $CH_2Cl_2$ (31 mL, 0.1 M) was added a solution of trimethylphenylammonium tribromide (1.29 g, 3.43 mmol, 1.1 eq.) in anhydrous $CH_2Cl_2$ (17 mL, 0.2 M) dropwise at 0°C over 20 min. The resulting dark-red solution was stirred at room temperature until TLC showed no starting material (~10-20 min) and then quenched with $H_2O$ (5.0 mL) and basified with aqueous saturated $NH_4OH$ (1 mL), and the aqueous layer was removed. The remaining organic layer was then washed with $H_2O$ (20.0 mL) and concentrated *in vacuo* to provide the crude bromide as a foamy brown solid. To the crude bromide, tetrakis(triphenylphosphine)palladium(0) (360 mg, 0.31 mmol, 10 mol%) and sodium formate (848 mg, 12.5 mmol, 4 eq., powdered and dried with gentle heating under vacuum before use) were added followed by anhydrous DMSO (12 mL, 0.25 M). The resulting mixture was heated to 130°C for 1 h (gas evolution occurs). The reaction was then diluted with $H_2O$ (20 mL) and extracted with $CH_2Cl_2$ (3 × 40 mL). The crude product was purified by flash column chromatography (using 20% of AcOEt in hexanes + 2% $Et_3N$). The product was isolated as a light yellow solid (226 mg, 24% over 2 steps).

**$^1$H NMR (500 MHz, MeOD)** δ 7.15 (dd, *J* = 8.7, 4.4 Hz, 1H), 7.02 (dd, *J* = 10.0, 2.5 Hz, 1H), 6.75 (ddd, *J* = 9.6, 8.6, 2.5 Hz, 1H), 3.13 – 3.00 (m, 3H), 2.94 (dt, *J* = 9.5, 3.0 Hz, 1H), 2.86 – 2.83 (m, 1H), 2.58 (dt, *J* = 16.3, 3.5 Hz, 1H), 2.12 (ddt, *J* = 14.0, 11.7, 2.7 Hz, 1H), 1.94 – 1.81 (m, 3H), 1.67 – 1.54 (m, 4H), 1.49 (dt, *J* = 13.0, 7.1 Hz, 1H), 1.22 (ddt, *J* = 12.6, 5.4, 2.6 Hz, 1H), 0.94 (t, *J* = 7.2 Hz, 3H). **$^{19}$F NMR (471 MHz, MeOD)** δ -127.48. **LRMS (EI)** calcd. for $C_{19}H_{23}FN_2$ [$M^+$] 298.41, found 298.25.

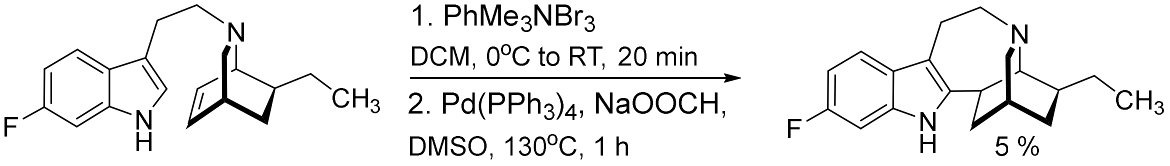


**6-fluoro-ibogamine**

Synthesized utilizing procedures found in our prior publication.[14] To a solution of the substrate (1.05 g, 3.50 mmol, 1 eq.) in anhydrous $CH_2Cl_2$ (35 mL, 0.1 M),a solution of trimethylphenylammonium tribromide (1.46 g, 3.87 mmol, 1.1 eq.) in anhydrous $CH_2Cl_2$ (19 mL, 0.2 M) was dropwise added at 0°C over 20 min. The resulting dark-red solution was stirred at room temperature until TLC showed no starting material (~10-20 min) and then quenched with $H_2O$ (5.0 mL) and basified with aqueous saturated $NH_4OH$ (1 mL), and the aqueous layer was removed. The remaining organic layer was then washed with $H_2O$ (20.0 mL) and concentrated *in vacuo* to provide the crude bromide as a foamy brown solid. To the crude bromide, tetrakis(triphenylphosphine)palladium(0) (407 mg, 0.35 mmol, 10 mol%) and sodium formate (957 mg, 14.1 mmol, 4 eq., powdered and dried with gentle heating under vacuum before use) was added followed by anhydrous DMSO (14 mL, 0.25 M). The resulting mixture was heated to 130°C for 1 h (gas evolution occurs). The reaction was then diluted with $H_2O$ (20 mL) and extracted with $CH_2Cl_2$ (3 × 40 mL). The crude product was purified by flash column chromatography (using 20% of AcOEt in hexanes + 2% $Et_3N$). The product was isolated as a light brown solid (51 mg, 5% over 2 steps).

**$^1$H NMR (500 MHz, $CDCl_3$)** δ 7.66 (s, 1H), 7.35 (dd, *J* = 8.6, 5.3 Hz, 1H), 6.93 (dd, *J* = 9.6, 2.3 Hz, 1H), 6.84 (ddd, *J* = 9.7, 8.6, 2.3 Hz, 1H), 3.44 – 3.28 (m, 2H), 3.15 (ddd, *J* = 13.5, 12.0, 3.7 Hz, 1H), 3.09 – 2.99 (m, 2H), 2.92 (ddd, *J* = 11.6, 4.0, 2.0 Hz, 1H), 2.87 (s, 1H), 2.69 – 2.61 (m, 1H), 2.05 (ddd, *J* = 13.9, 9.9, 2.4 Hz, 1H), 1.90 – 1.77 (m, 2H), 1.64 (dq, *J* = 13.2, 3.5 Hz, 1H), 1.60 – 1.46 (m, 3H), 1.23 (ddd, *J* = 15.3, 5.4, 2.5 Hz, 1H), 0.91 (t, *J* = 7.1 Hz, 3H). **$^{13}$C NMR (126 MHz, $CDCl_3$)** δ 160.59, 158.72, 142.07, 134.74, 134.64, 126.38, 118.68, 118.60, 109.17, 107.74, 107.55, 96.84, 96.63, 57.86, 54.24, 50.02, 41.97, 34.21, 32.11, 27.81, 26.52, 20.75, 14.33, 12.03. **$^{19}$F NMR (470 MHz, $CDCl_3$)** δ -122.77. **LRMS (EI)** calcd. for $C_{19}H_{23}FN_2$ [$M^+$] 298.41, found 298.22.

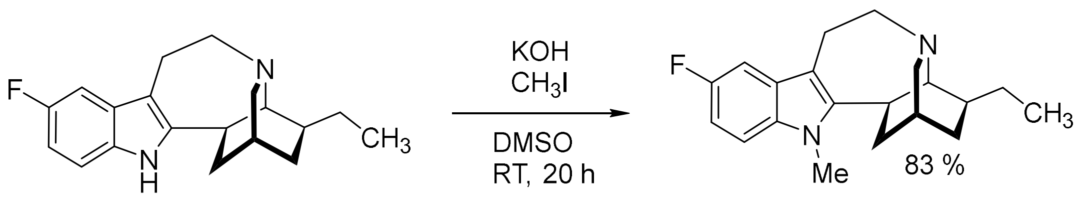


##### *N*-methyl-5-fluoro-ibogamine

5-fluoro-ibogamine (15 mg, 0.05 mmol) and KOH (4.2 mg, 0.075 mmol, 1.5 eq.) were mixed in DMSO (anhydrous, 0.1 mL, 0.5 M). $CH_3I$ (4.7 µL, 0.075 mmol, 1.5 eq.) was added into the suspension and the resulting mixture was stirred at RT for 20h. The reaction was quenched by adding $H_2O$ (5 mL) and the mixture was extracted with $CH_2Cl_2$ (3 × 5 mL). Combined extracts were dried over $Na_2SO_4$, filtered, and concentrated under reduced pressure. Crude material was purified by PTLC (98% hexanes + 2% $Et_3N$). The product was isolated as a light brown solid (13 mg, 83%).

**$^1$H NMR (500 MHz, MeOD)** δ 7.18 (dd, *J* = 8.8, 4.3 Hz, 1H), 7.04 (dt, *J* = 12.6, 5.1 Hz, 1H), 6.83 (ddd, *J* = 9.5, 8.8, 2.5 Hz, 1H), 3.63 (s, 2H), 3.28 – 3.25 (m, 2H), 3.20 (ddd, *J* = 11.7, 4.6, 1.9 Hz, 1H), 3.16 – 3.00 (m, 2H), 2.95 (dt, *J* = 9.5, 3.0 Hz, 1H), 2.85 (t, *J* = 1.8 Hz, 1H), 2.69 – 2.61 (m, 1H), 2.21 – 2.11 (m, 1H), 1.91 – 1.83 (m, 2H), 1.70 – 1.66 (m, 1H), 1.63 – 1.46 (m, 3H), 1.27 – 1.19 (m, 1H), 0.95 (t, *J* = 7.4 Hz, 3H). **$^{13}$C NMR (126 MHz, MeOD)** δ 158.56, 156.72, 144.75, 133.00, 128.55, 128.47, 108.93, 108.85, 107.88, 107.67, 102.08, 101.89, 57.35, 54.07, 49.85, 42.25, 38.32, 33.11, 31.60, 28.59, 27.41, 26.48, 20.37, 10.88. **$^{19}$F NMR (471 MHz, MeOD)** δ -127.44. **LRMS (APCI+)** calcd. for $C_{20}H_{26}FN_2^+$ $[M+H]^+$ 313.4, found 313.2.

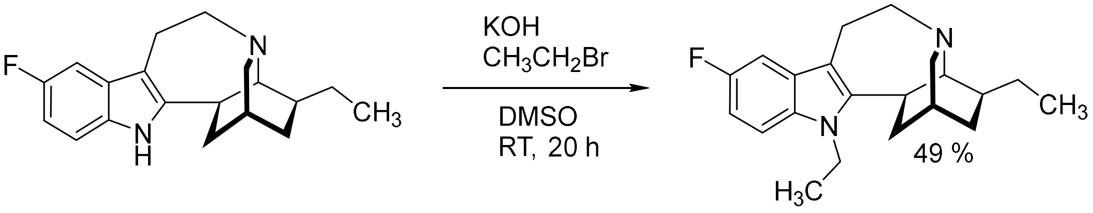


##### *N*-ethyl-5-fluoro-ibogamine

5-fluoro-ibogamine (15 mg, 0.05 mmol) and KOH (4.2 mg, 0.075 mmol, 1.5 eq.) were mixed in DMSO (anhydrous, 0.1 mL, 0.5 M). $CH_3CH_2Br$ (5.6 µL, 0.075 mmol, 1.5 eq.) was added into the suspension, and the resulting mixture was stirred at RT for 20 h. The reaction was quenched by adding $H_2O$ (5 mL), and the mixture was extracted with $CH_2Cl_2$ (3 × 5 mL). Combined extracts were dried over $Na_2SO_4$, filtered, and concentrated under reduced pressure. Crude material was purified by PTLC (98% hexanes + 2% $Et_3N$). The product was isolated as a light brown solid (8 mg, 49%).

**$^1$H NMR (500 MHz, MeOD)** δ 7.19 (dd, *J* = 8.8, 4.3 Hz, 1H), 7.04 (dd, *J* = 9.9, 2.6 Hz, 1H), 6.80 (ddd, *J* = 8.9, 6.0, 2.5 Hz, 1H), 4.20 – 4.04 (m, 3H), 3.16 – 3.03 (m, 4H), 2.99 – 2.92 (m, 1H), 2.85 – 2.77 (m, 1H), 2.60 (dt, *J* = 17.0, 3.3 Hz, 1H), 2.22 – 2.10 (m, 2H), 1.95 – 1.80 (m, 2H), 1.72 – 1.65 (m, 1H), 1.64 – 1.53 (m, 2H), 1.50 – 1.42 (m, 1H), 1.26 – 1.17 (m, 5H), 0.96 – 0.87 (m, 4H). **$^{13}$C NMR (126 MHz, MeOD)** δ 158.58, 144.18, 109.04, 108.96, 107.83, 107.63, 102.11, 101.93, 57.30, 54.25, 49.92, 42.53, 38.13, 37.26, 33.67, 31.53, 27.65, 26.52, 20.18, 14.70, 10.90. **$^{19}$F NMR (471 MHz, MeOD)** δ -127.37. **LRMS (APCI+)** calcd. for $C_{21}H_{28}FN_2^+$ $[M+H]^+$ 327.5, found 237.3.

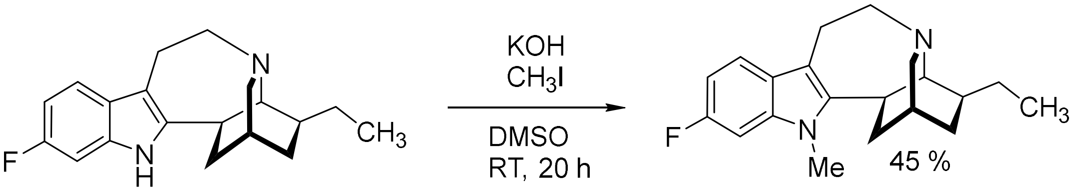


##### ***N*-methyl-6-fluoro-ibogamine**

6-fluoro-ibogamine (15 mg, 0.05 mmol) and KOH (4.2 mg, 0.075 mmol, 1.5 eq.) were mixed in DMSO (anhydrous, 0.1 mL, 0.5 M). $CH_3I$ (4.7 µL, 0.075 mmol, 1.5 eq.) was added into the suspension, and the resulting mixture was stirred at RT for 20h. The reaction was quenched by adding $H_2O$ (5 mL), and the mixture was extracted with $CH_2Cl_2$ (3 × 5 mL). Combined extracts were dried over $Na_2SO_4$, filtered, and concentrated under reduced pressure. Crude material was purified by PTLC (98% hexanes + 2% $Et_3N$). The product was isolated as a light brown solid (7 mg, 45%).

**$^1$H NMR (500 MHz, MeOD)** δ 7.31 (dd, *J* = 8.5, 5.3 Hz, 1H), 6.95 (dt, *J* = 10.8, 3.7 Hz, 1H), 6.78 – 6.68 (m, 1H), 3.58 (s, 3H), 3.30 – 3.25 (m, 2H), 3.19 (ddd, *J* = 11.5, 4.4, 2.0 Hz, 1H), 3.14 – 3.06 (m, 1H), 3.03 (td, *J* = 6.7, 3.0 Hz, 1H), 2.92 (dq, *J* = 10.2, 3.5 Hz, 1H), 2.84 (d, *J* = 2.1 Hz, 1H), 2.72 – 2.66 (m, 1H), 2.15 – 2.09 (m, 1H), 1.94 – 1.82 (m, 2H), 1.72 – 1.62 (m, 1H), 1.61 – 1.44 (m, 4H), 1.22 (ddt, *J* = 12.7, 6.1, 2.5 Hz, 1H), 0.94 (t, *J* = 7.4 Hz, 3H). **$^{13}$C NMR (126 MHz, MeOD)** δ 160.45, 143.22, 136.42, 124.84, 118.02, 117.94, 108.29, 106.15, 105.95, 94.69, 94.48, 57.56, 54.00, 49.74, 42.21, 38.19, 33.20, 31.62, 28.61, 27.30, 26.48, 20.38, 10.87. **$^{19}$F NMR (471 MHz, MeOD)** δ -124.06. **LRMS (APCI+)** calcd. for $C_{20}H_{26}FN_2^+$ $[M+H]^+$ 313.4, found 313.2.

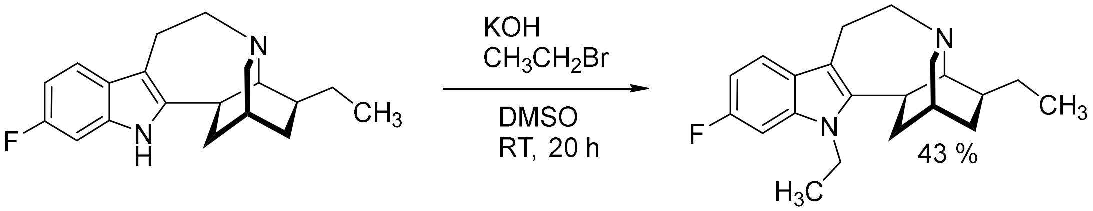


##### ***N*-ethyl-6-fluoro-ibogamine**

6-fluoro-ibogamine (15 mg, 0.05 mmol) and KOH (4.2 mg, 0.075 mmol, 1.5 eq.) were mixed in DMSO (anhydrous, 0.1 mL, 0.5 M). $CH_3CH_2Br$ (5.6 µL, 0.075 mmol, 1.5 eq.) was added into the suspension, and the resulting mixture was stirred at RT for 20 h. The reaction was quenched by adding $H_2O$ (5 mL), and the mixture was extracted with $CH_2Cl_2$ (3 × 5 mL). Combined extracts were dried over $Na_2SO_4$, filtered, and concentrated under reduced pressure. Crude material was purified by PTLC (98% hexanes + 2% $Et_3N$). The product was isolated as a light brown solid (7 mg, 43%).

**$^1$H NMR (500 MHz, MeOD)** δ 7.33 (dd, *J* = 8.6, 5.3 Hz, 1H), 6.99 (dd, *J* = 10.3, 2.3 Hz, 1H), 6.76 (ddd, *J* = 9.7, 8.6, 2.3 Hz, 1H), 4.18 – 4.02 (m, 2H), 3.18 – 3.06 (m, 4H), 2.96 (dt, *J* = 9.6, 3.0 Hz, 1H), 2.85 (d, *J* = 1.9 Hz, 1H), 2.68 (dt, *J* = 17.2, 3.4 Hz, 1H), 2.18 (d, *J* = 8.7 Hz, 1H), 1.97 – 1.84 (m, 3H), 1.74 – 1.67 (m, 1H), 1.59 – 1.44 (m, 5H), 1.27 (t, *J* = 7.2 Hz, 3H), 0.95 (t, *J* = 7.4 Hz, 3H). **$^{13}$C NMR (126 MHz, MeOD)** δ 160.39, 142.66, 125.05, 118.08, 118.00, 108.81, 106.22, 106.03, 94.76, 94.55, 57.50, 54.17,

49.82, 42.49, 38.01, 37.28, 33.80, 31.55, 27.54, 26.52, 20.18, 14.47, 10.89. **$^{19}$F NMR (471 MHz, MeOD)** δ -124.14. **LRMS (APCI+)** calcd. for $C_{21}H_{28}FN_2^+$ $[M+H]^+$ 327.5, found 237.3.

#### C. NMR and MS Spectra.

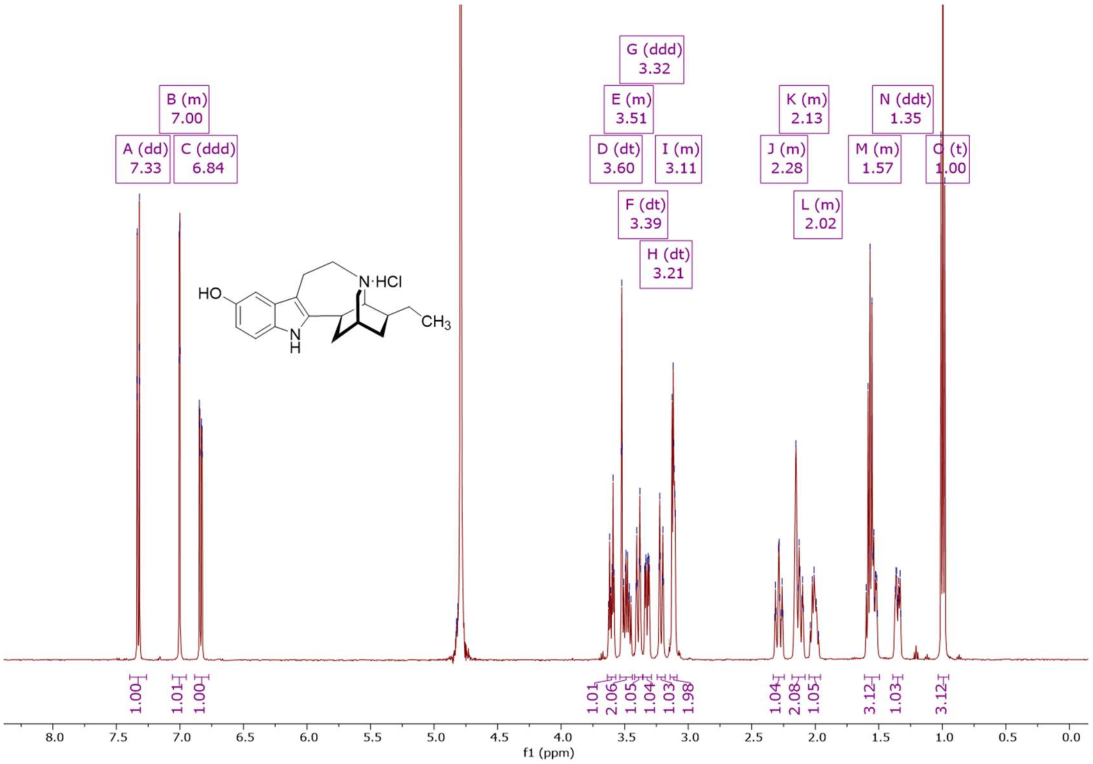


**[1]H NMR (500 MHz, $D_2O$)** spectrum of noribogaine hydrochloride.

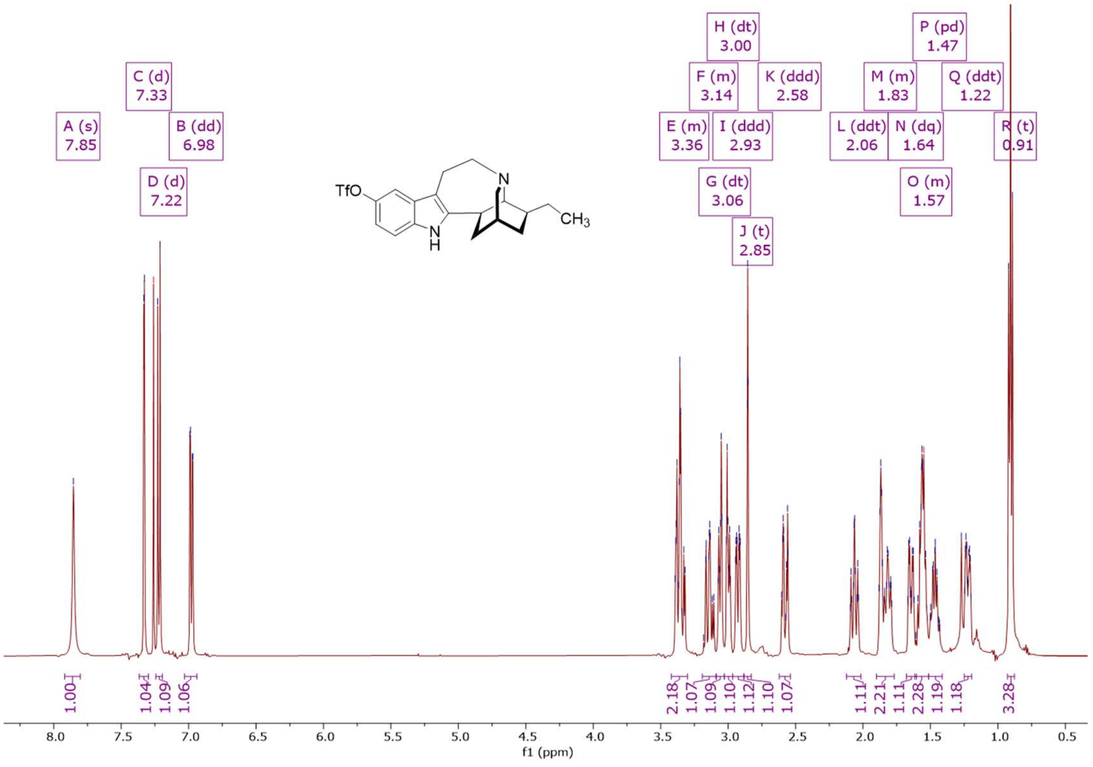


**[1]H NMR (500 MHz, $CDCl_3$)** spectrum of 5-triflate-ibogamine.

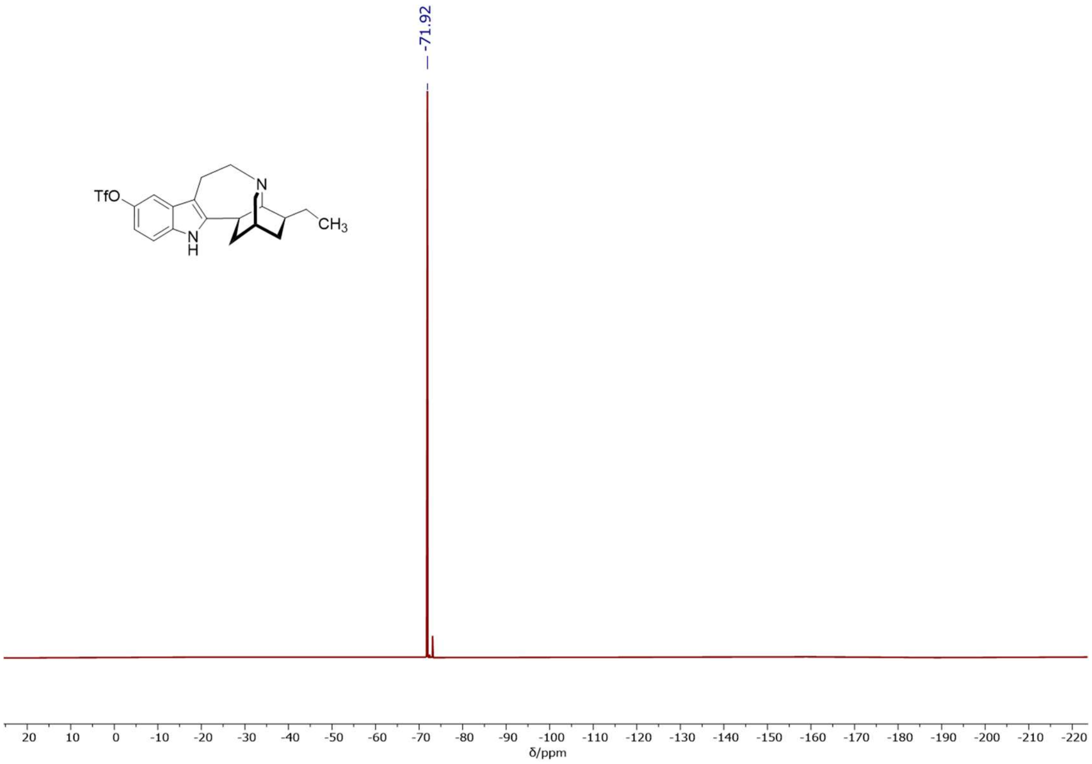


**$^{19}$F NMR (471 MHz, $CDCl_3$)** spectrum of 5-triflate-ibogamine.

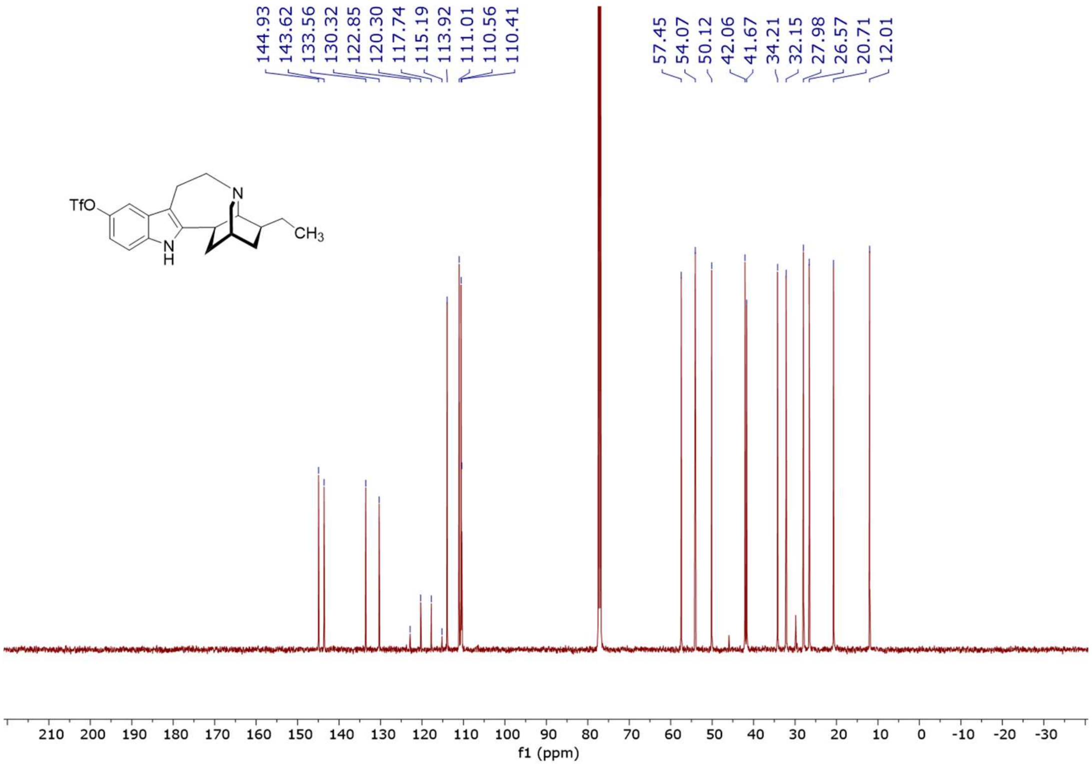


**$^{13}$C NMR (126 MHz, $CDCl_3$)** spectrum of 5-triflate-ibogamine-.

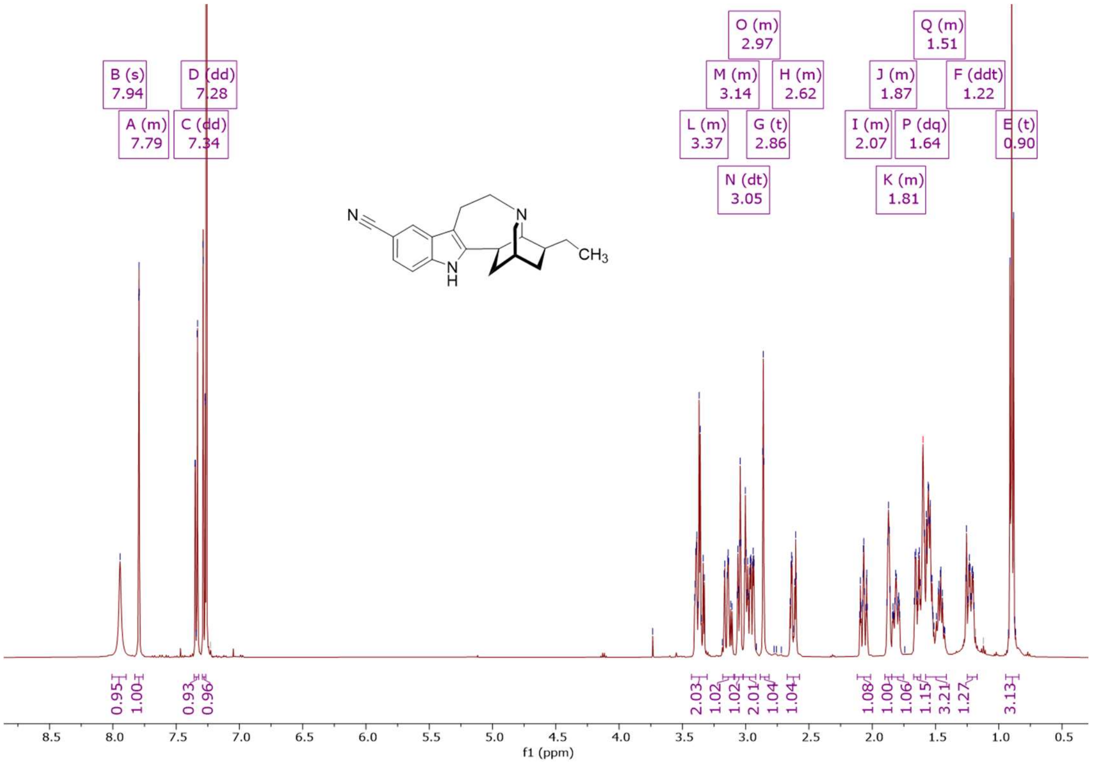


**¹H NMR (500 MHz, CDCl₃)** spectrum of 5-cyano-ibogamine.

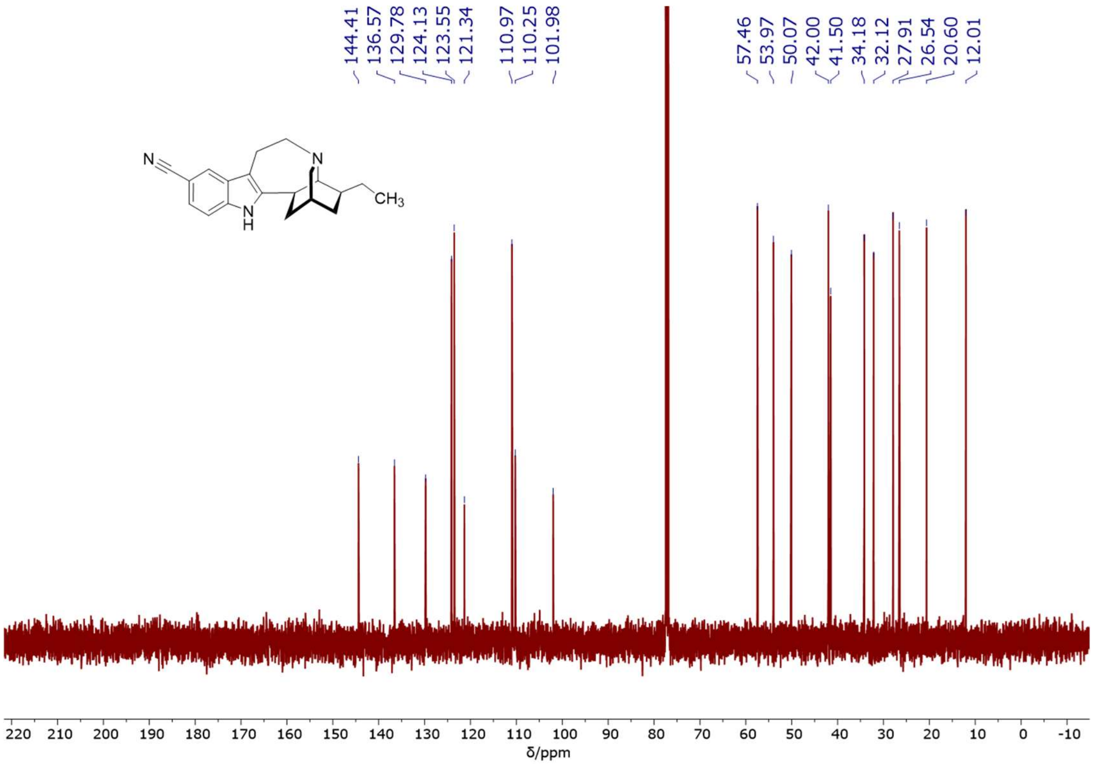


**¹³C NMR (126 MHz, CDCl₃)** spectrum of 5-cyano-ibogamine.

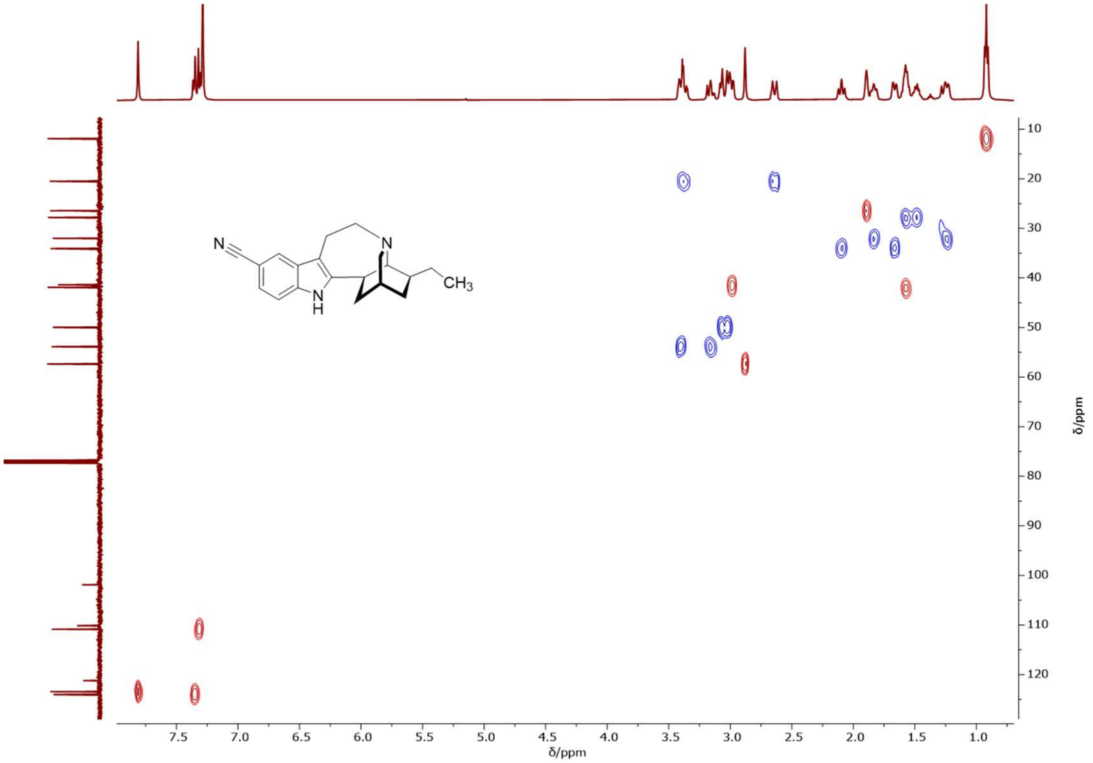


**$^{1}$H-$^{13}$C 2D NMR (500/126 MHz, $CDCl_3$) HSQC** spectrum of 5-cyano-ibogamine.

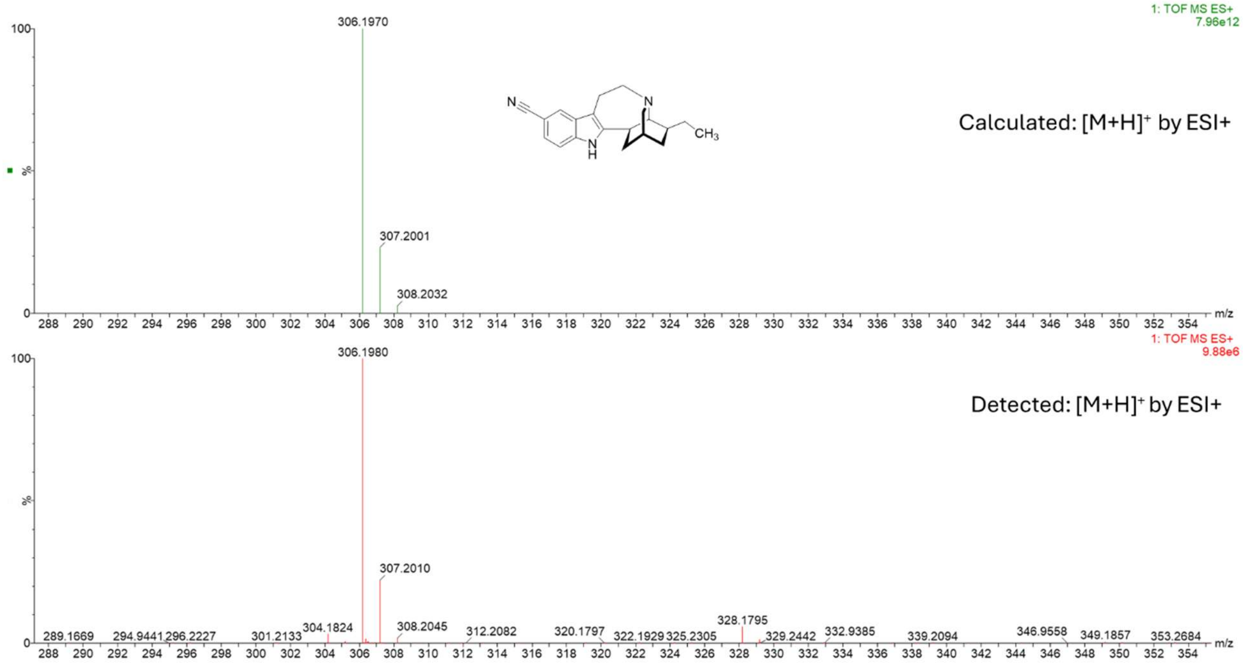


**HRMS (ESI+)** spectrum of 5-cyano-ibogamine.

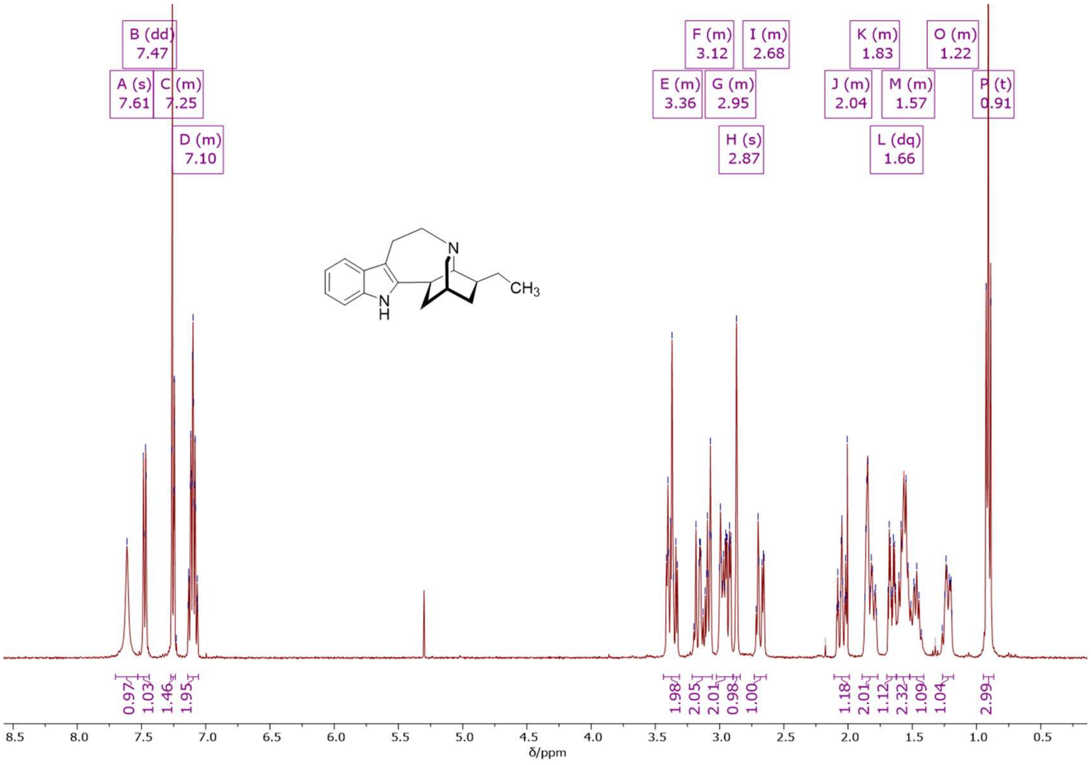


**[1]H NMR (500 MHz, $CDCl_3$)** spectrum of ibogamine.

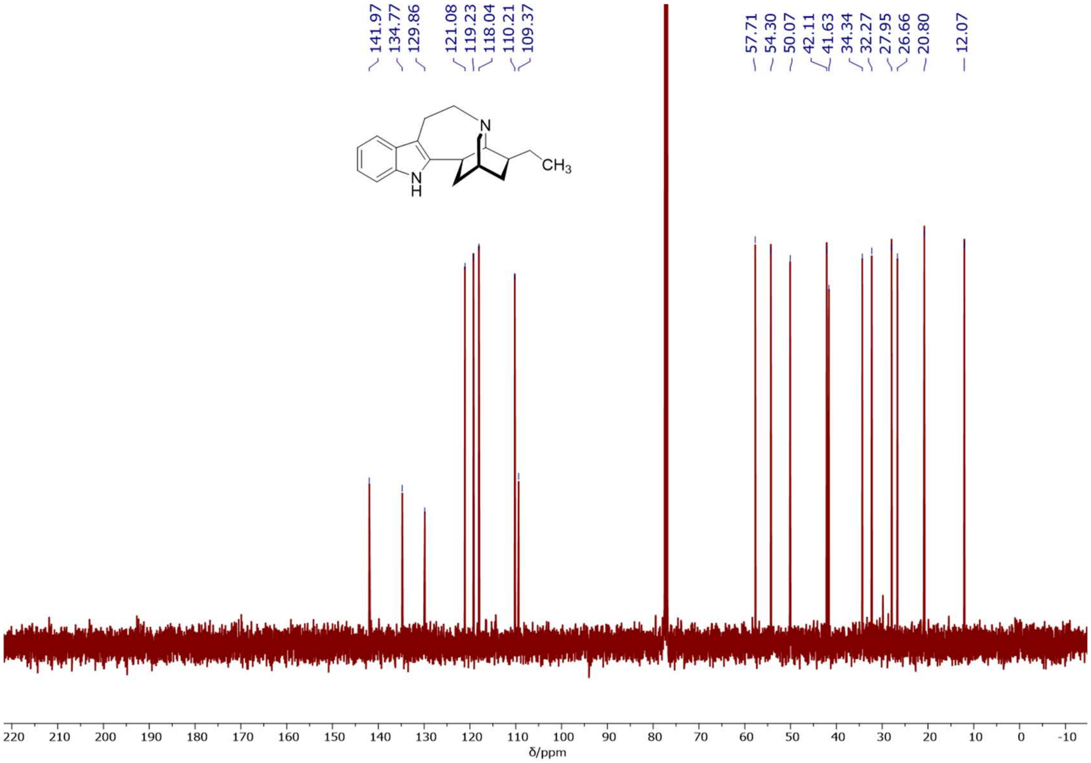


**[13]C NMR (126 MHz, $CDCl_3$)** spectrum of ibogamine.

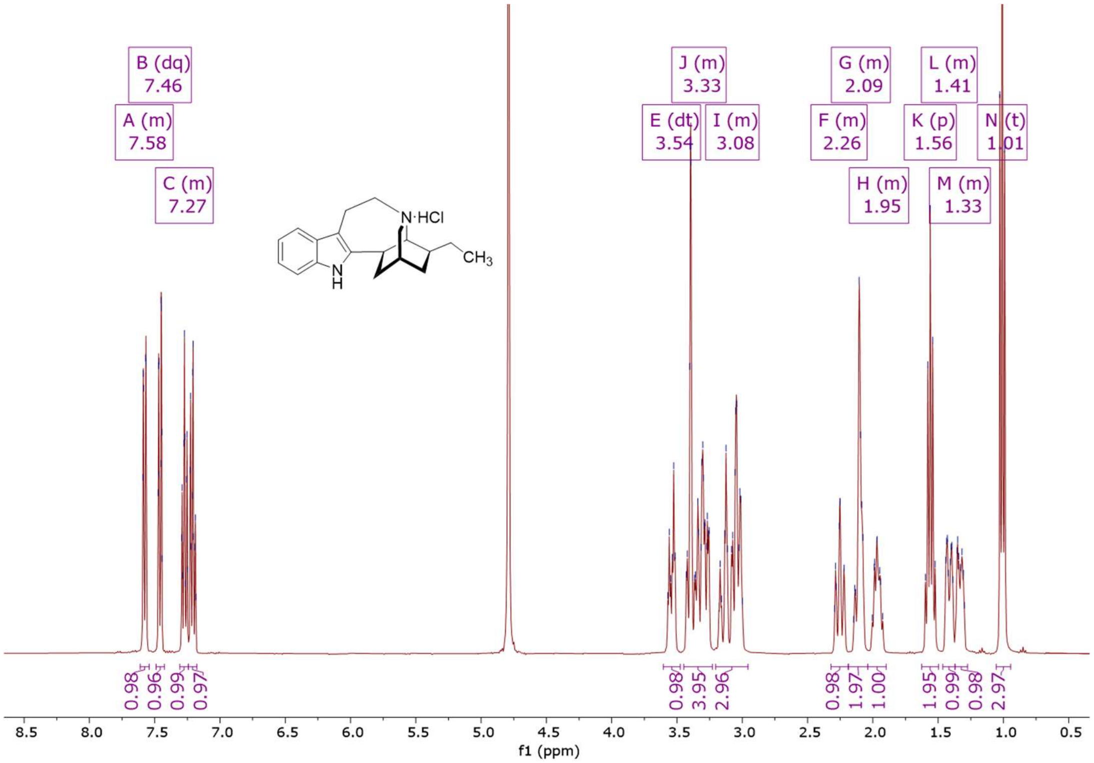


**$^1$H NMR (400 MHz, $D_2O$)** spectrum of ibogamine hydrochloride.

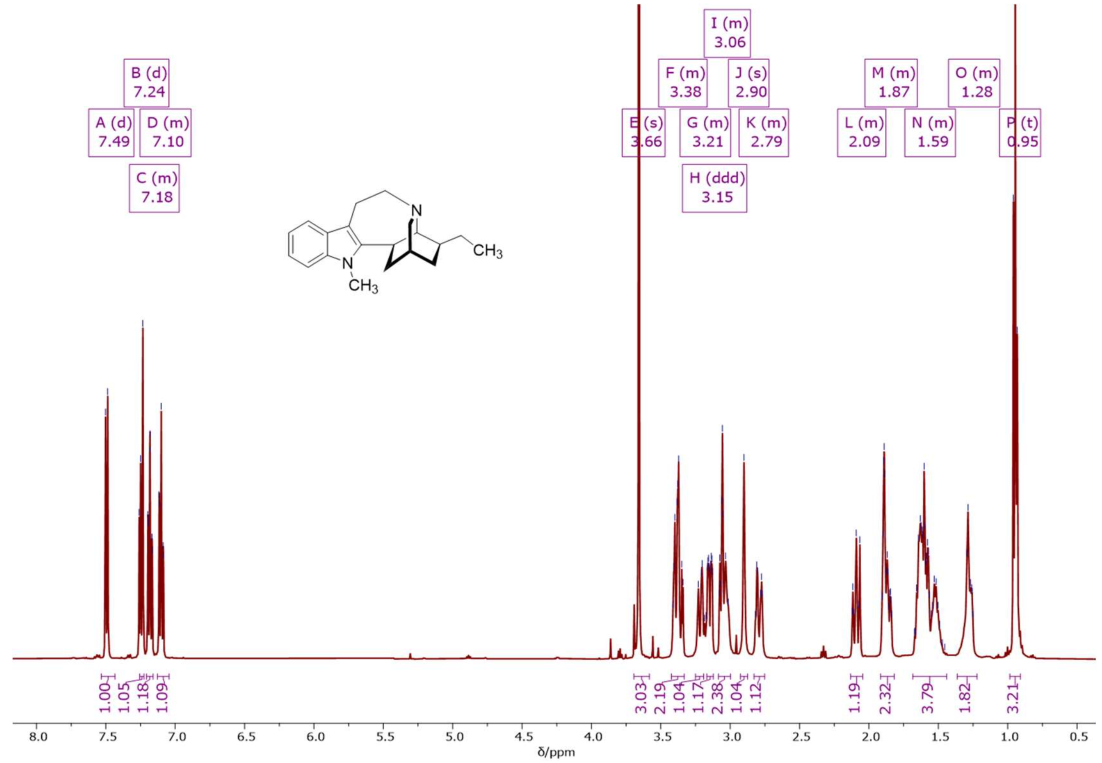


**$^1$H NMR (500 MHz, $CDCl_3$)** spectrum of *N*-methyl-ibogamine.

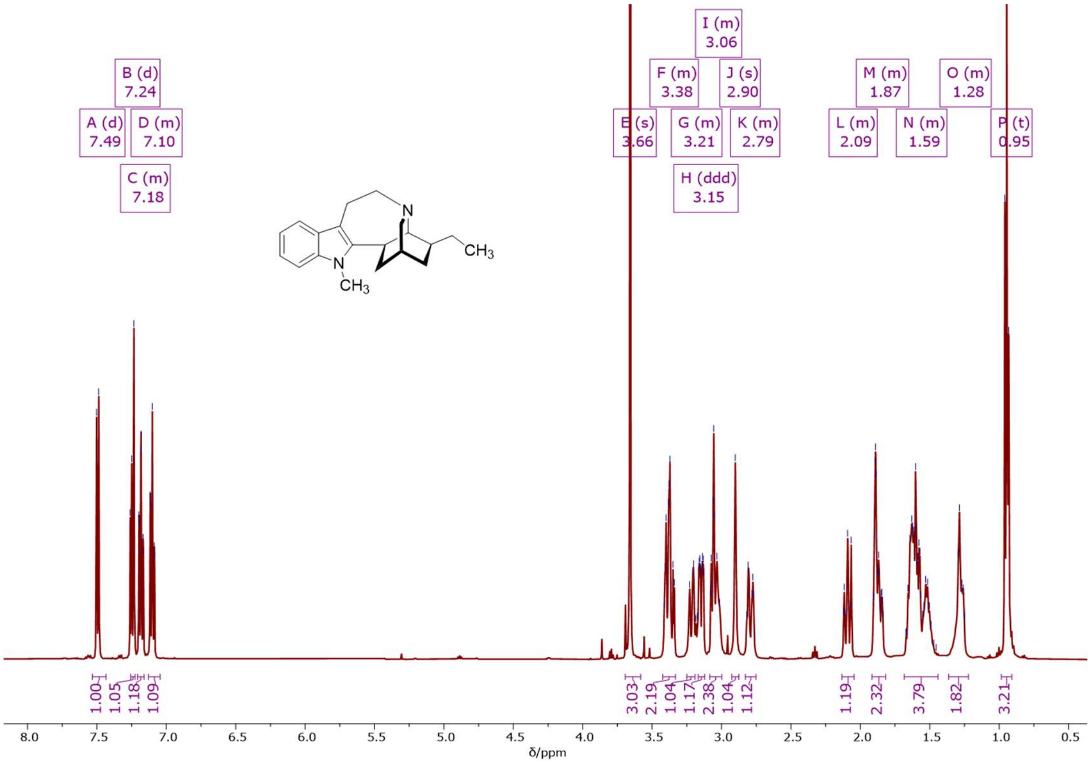


**$^1$H NMR (500 MHz, MeOD)** spectrum of *N*-methyl-ibogamine.

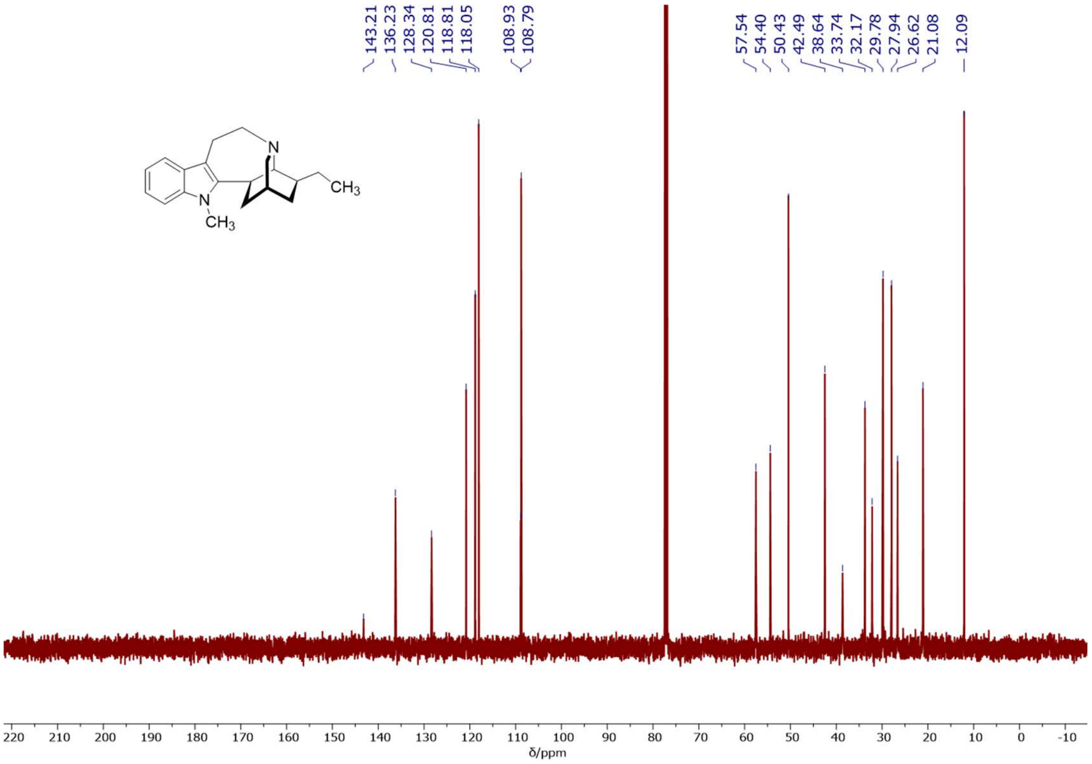


**$^{13}$C NMR (126 MHz, MeOD)** spectrum of *N*-methyl-ibogamine.

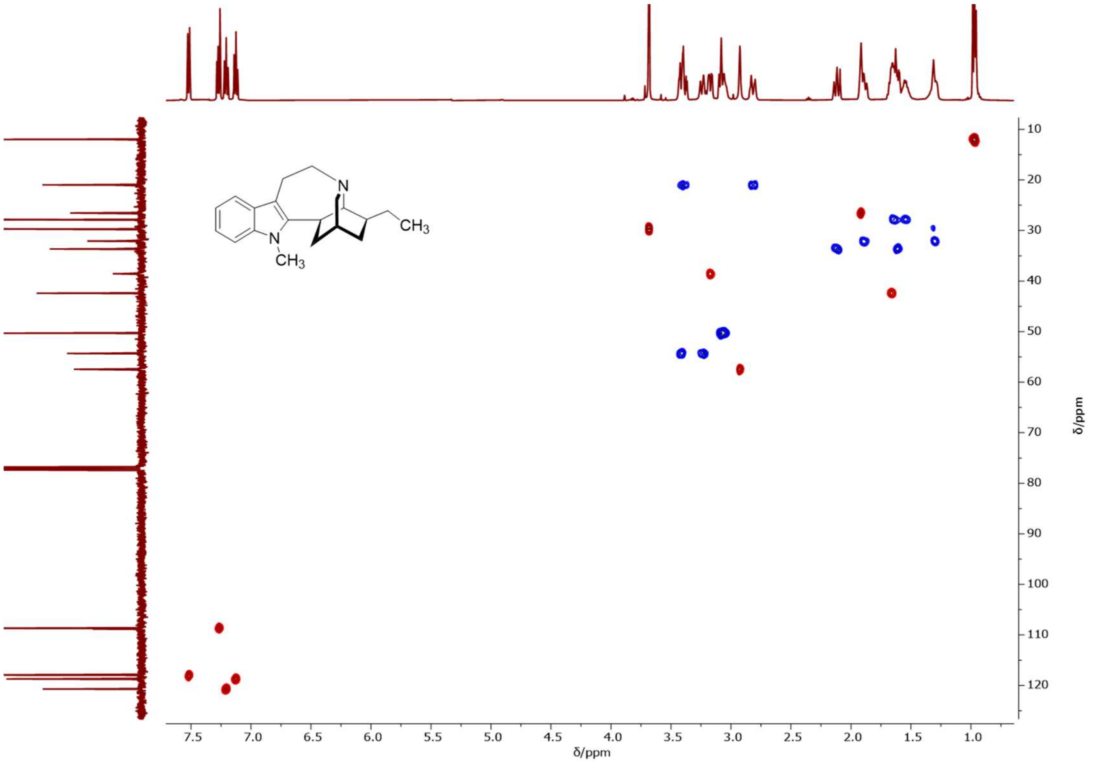

**$^1$H-$^{13}$C 2D NMR (500/126 MHz, $CDCl_3$) HSQC** spectrum of *N*-methyl-ibogamine.

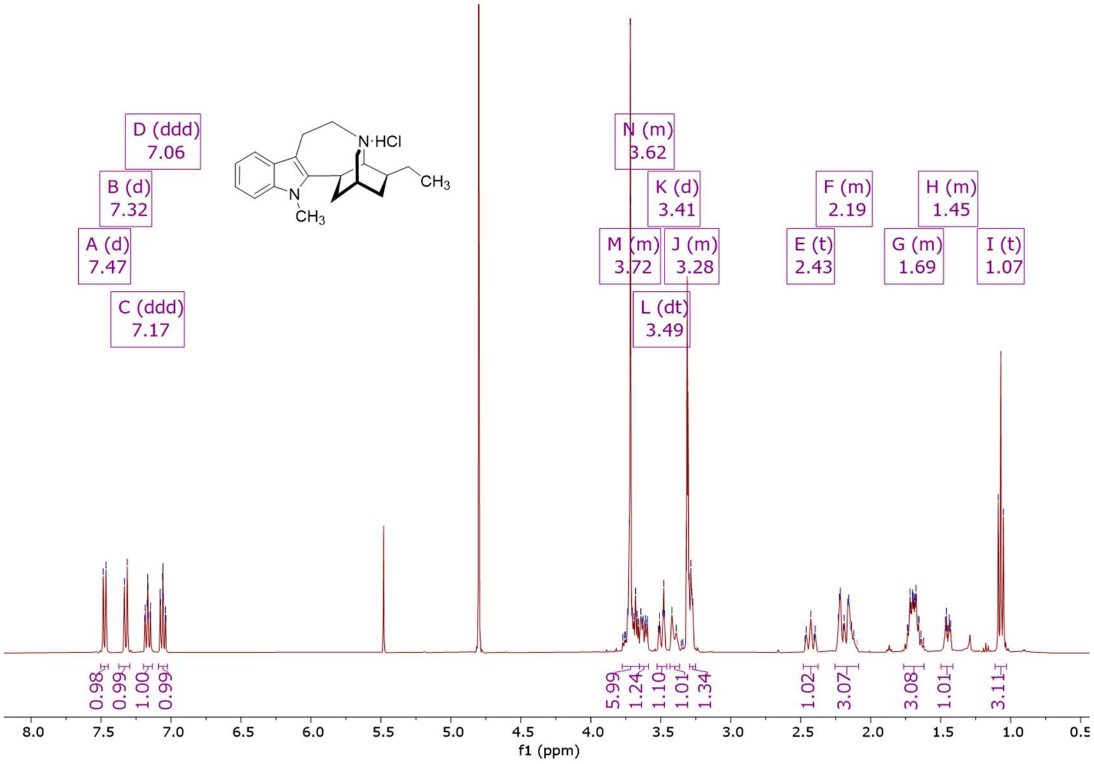


**$^1$H NMR (500 MHz, MeOD)** spectrum of *N*-methyl-ibogamine hydrochloride.

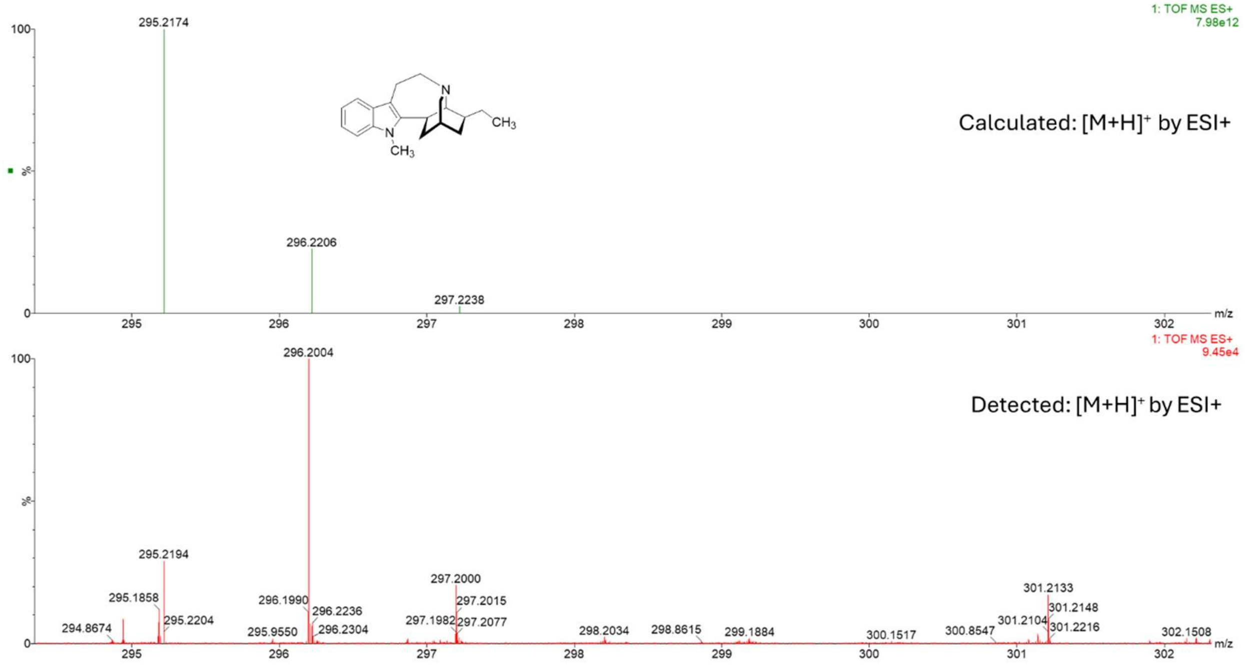


**HRMS (ESI+)** spectrum of *N*-methyl-ibogamine.

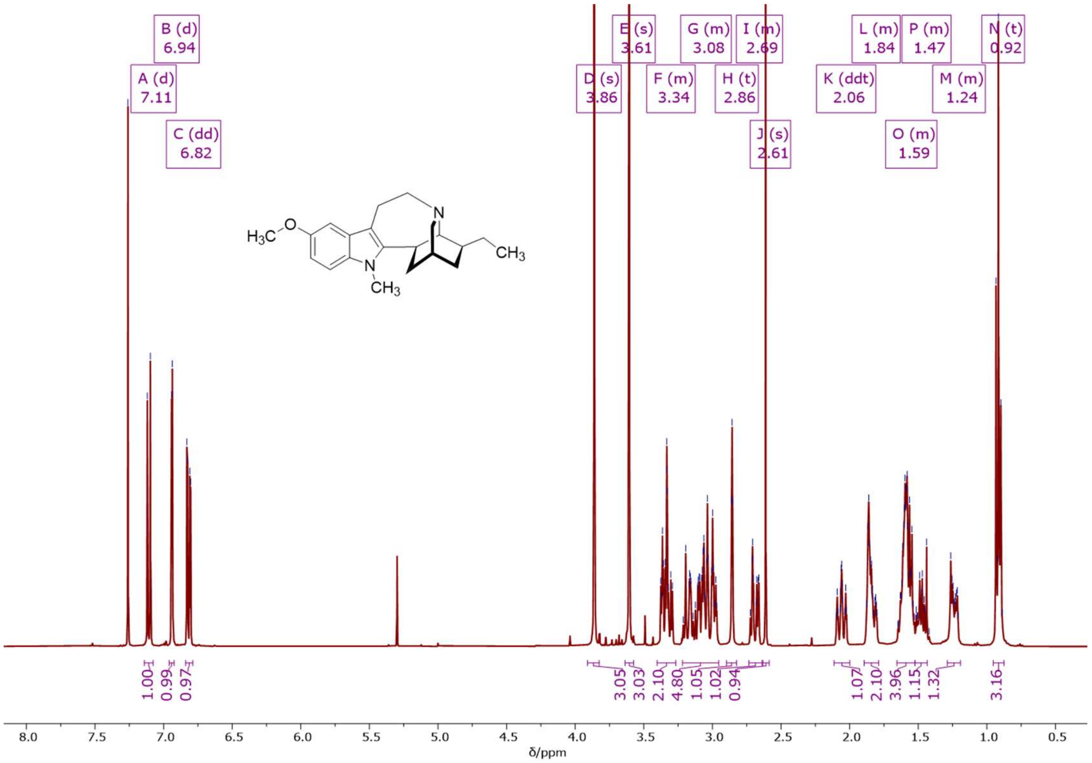


**[1]H NMR (500 MHz, CDCl$_3$)** spectrum of *N*-methyl-ibogaine.

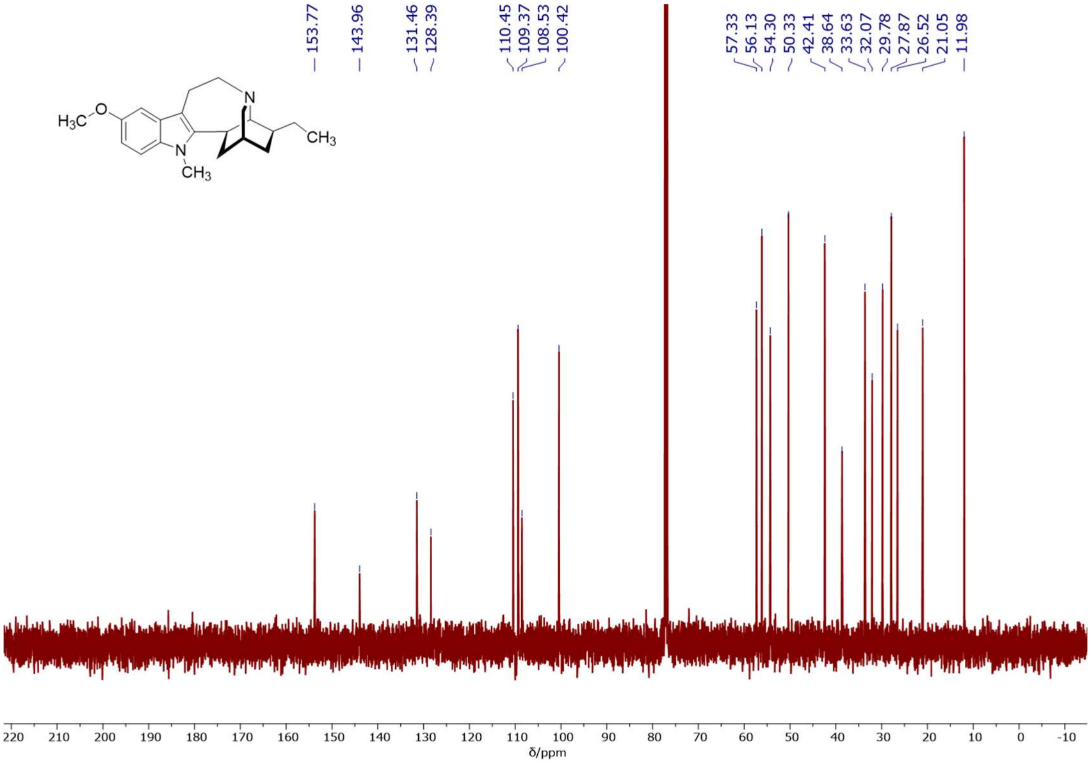

**$^{13}$C NMR (126 MHz, $CDCl_3$)** spectrum of *N*-methyl-ibogaine.

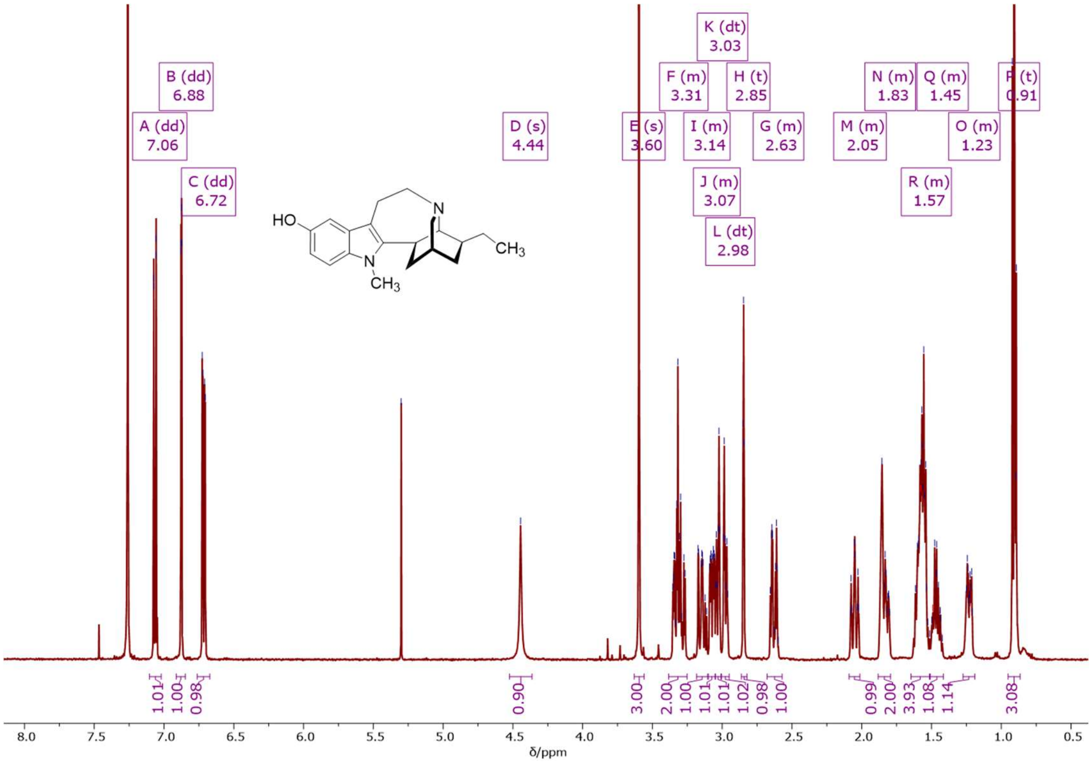

**$^{1}$H NMR (500 MHz, $CDCl_3$)** spectrum of *N*-methyl-noribogaine.

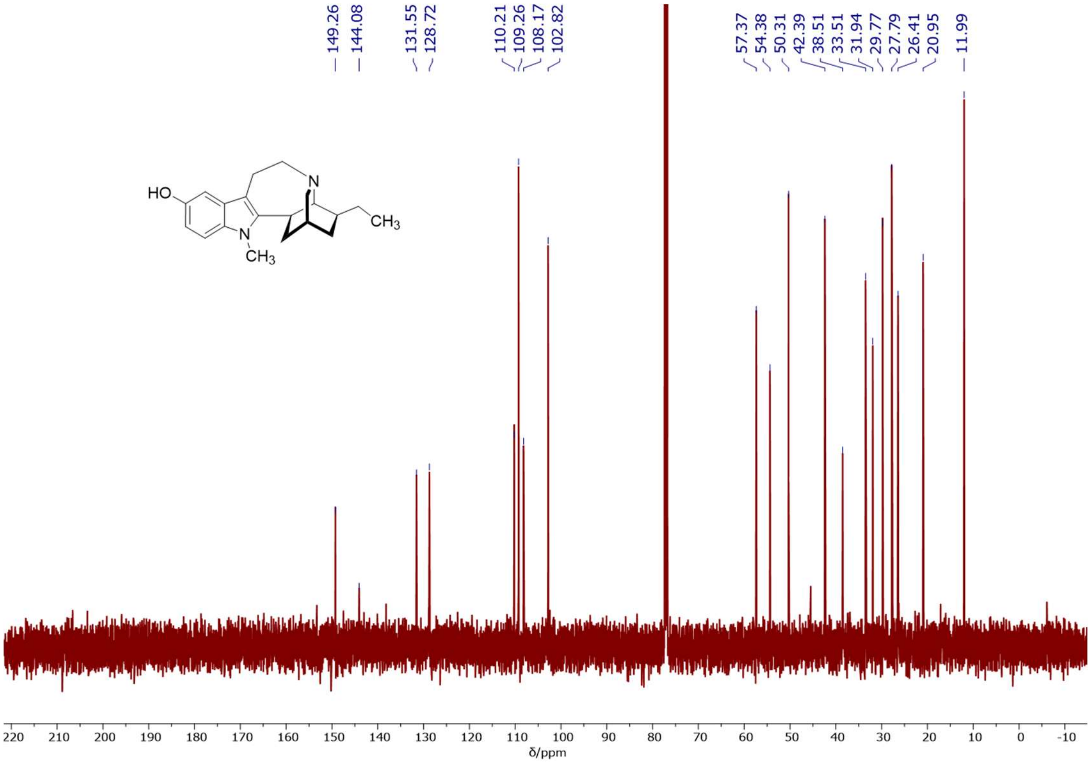


**$^{13}$C NMR (126 MHz, $CDCl_3$)** spectrum of *N*-methyl-noribogaine.

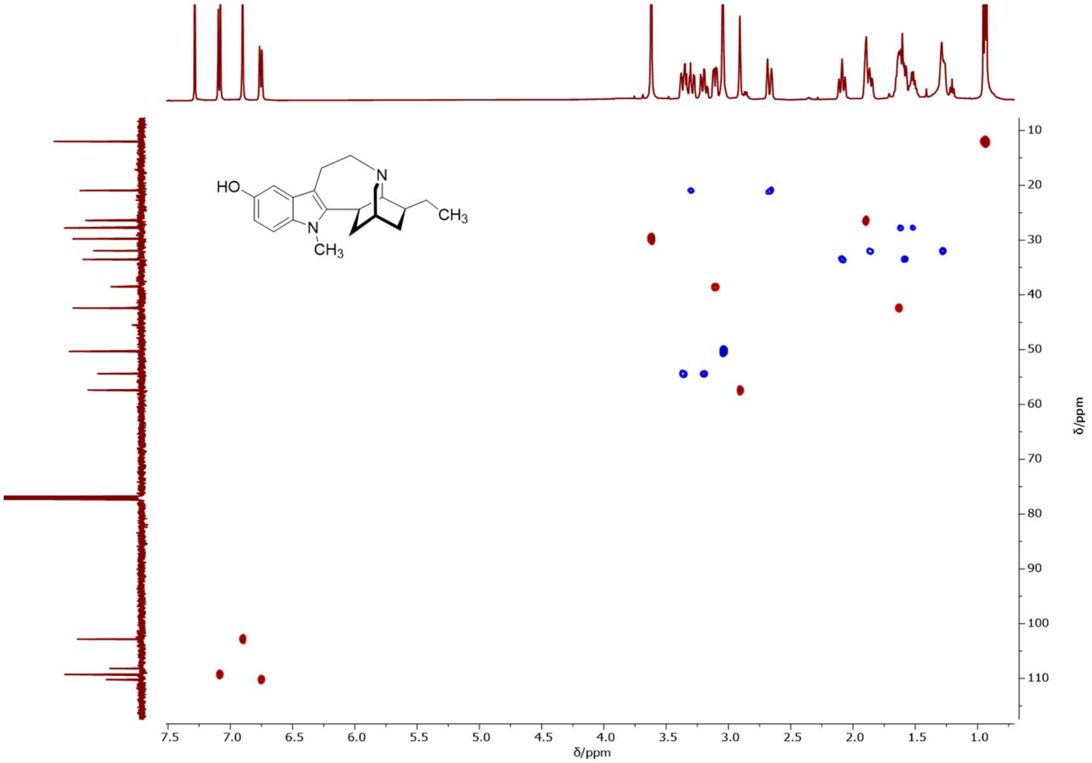


**$^{1}$H-$^{13}$C 2D NMR (500/126 MHz, $CDCl_3$) HSQC** spectrum of *N*-methyl-noribogaine.

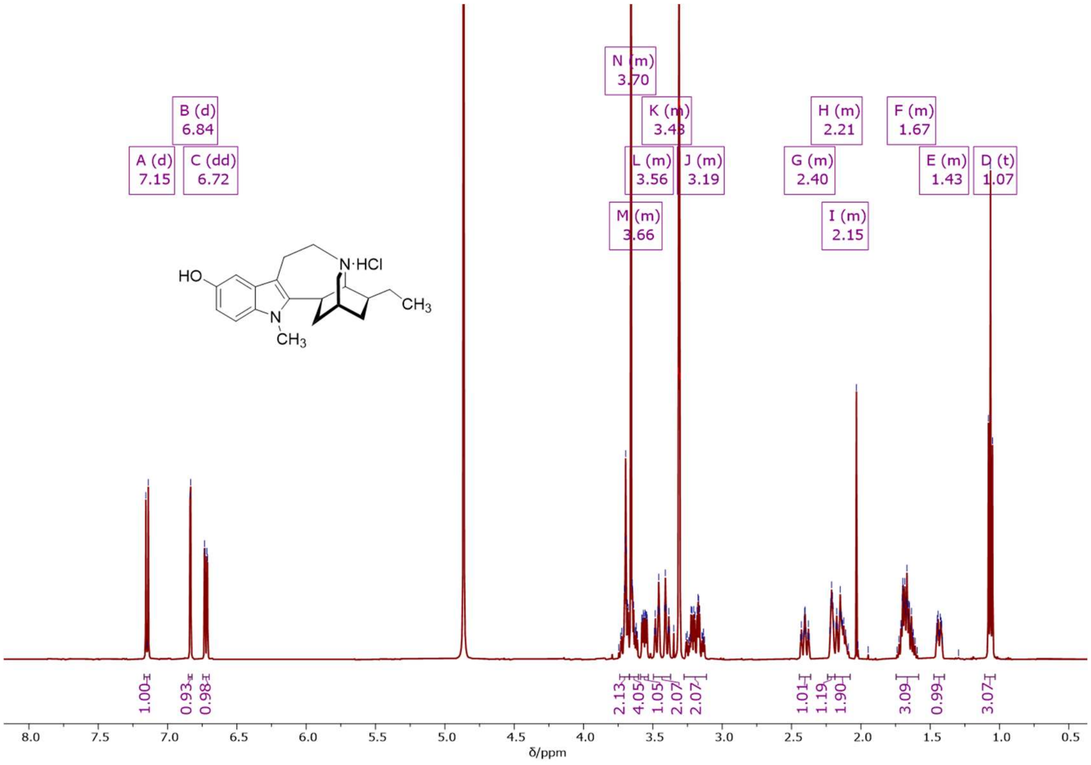


**[1]H NMR (500 MHz, MeOD)** spectrum of *N*-methyl-noribogaine hydrochloride.

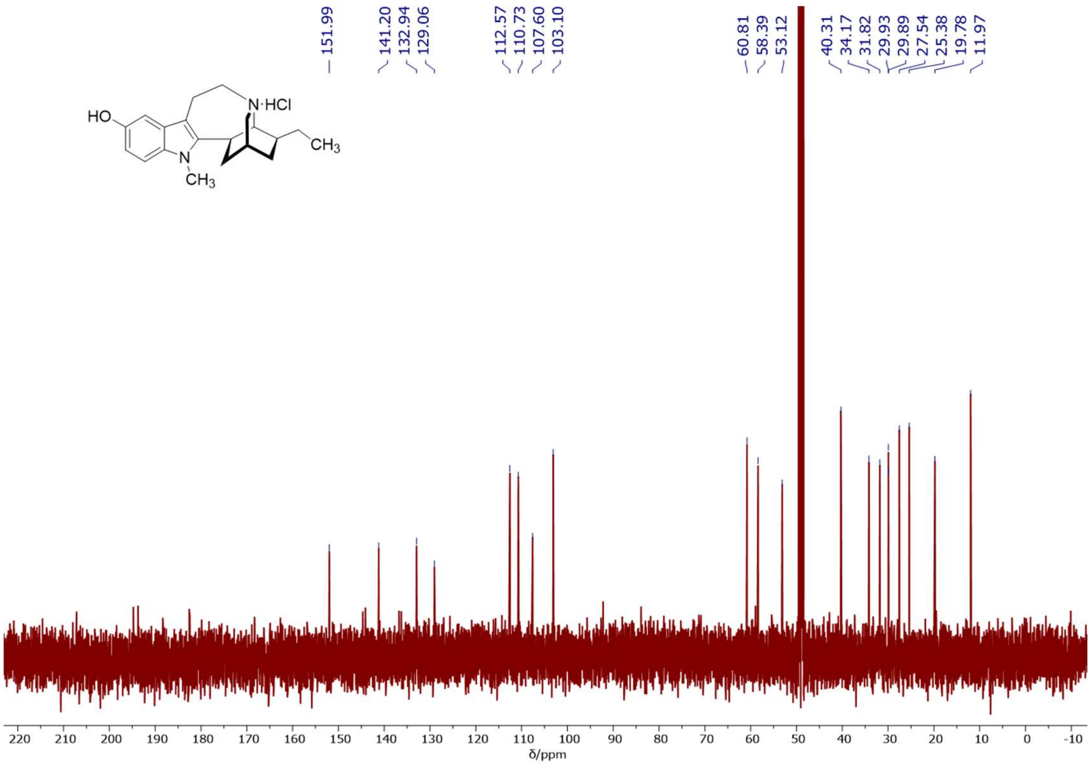


**[13]C NMR (126 MHz, MeOD)** spectrum of *N*-methyl-noribogaine hydrochloride.

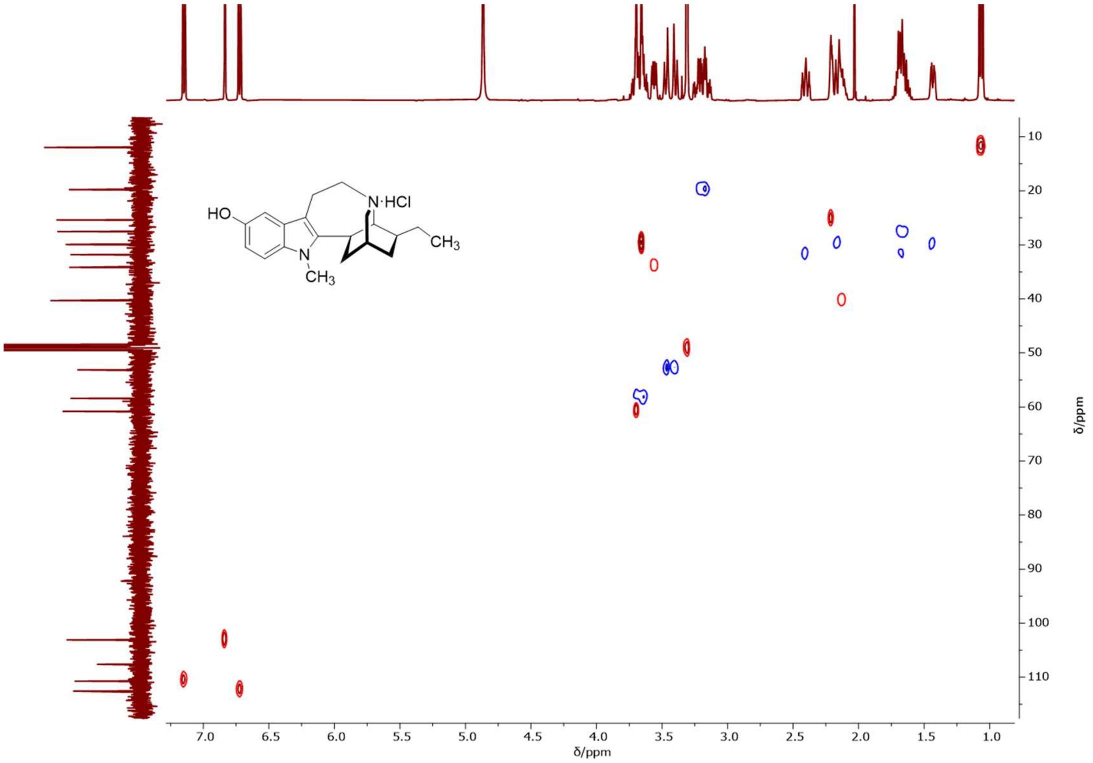


**¹H-¹³C 2D NMR (500/126 MHz, MeOD) HSQC** spectrum of *N*-methyl-noribogaine hydrochloride.

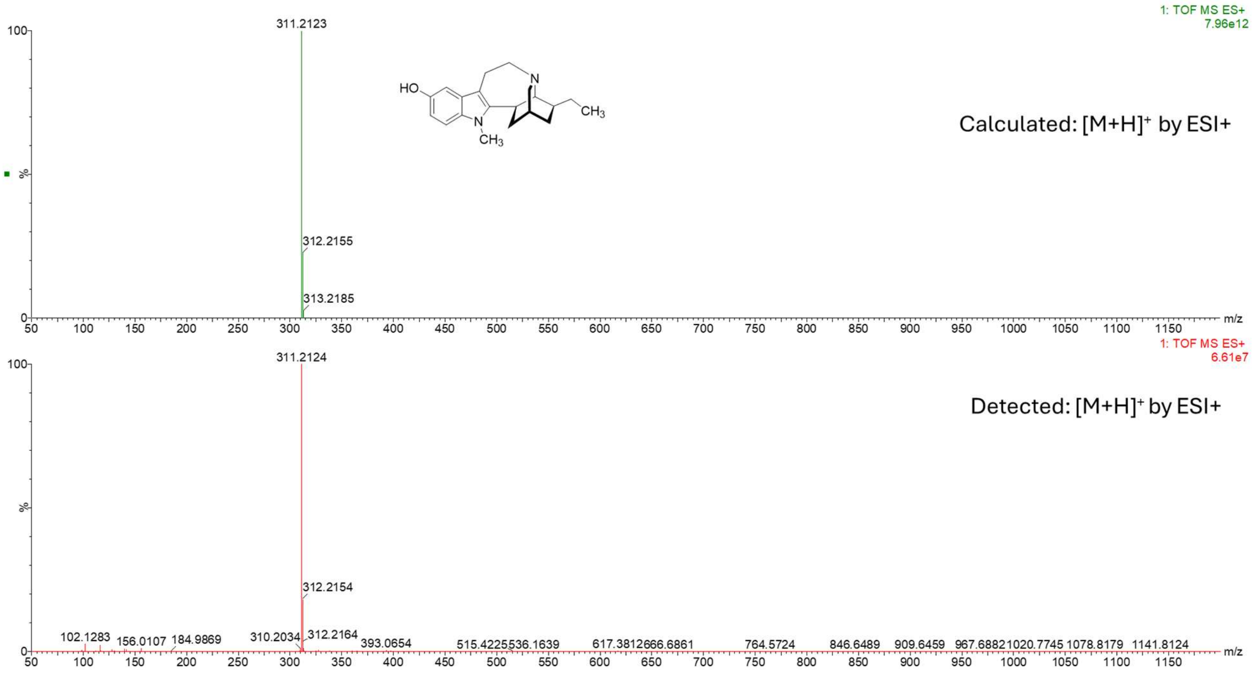


**HRMS (ESI+)** spectrum of *N*-methyl-noribogaine.

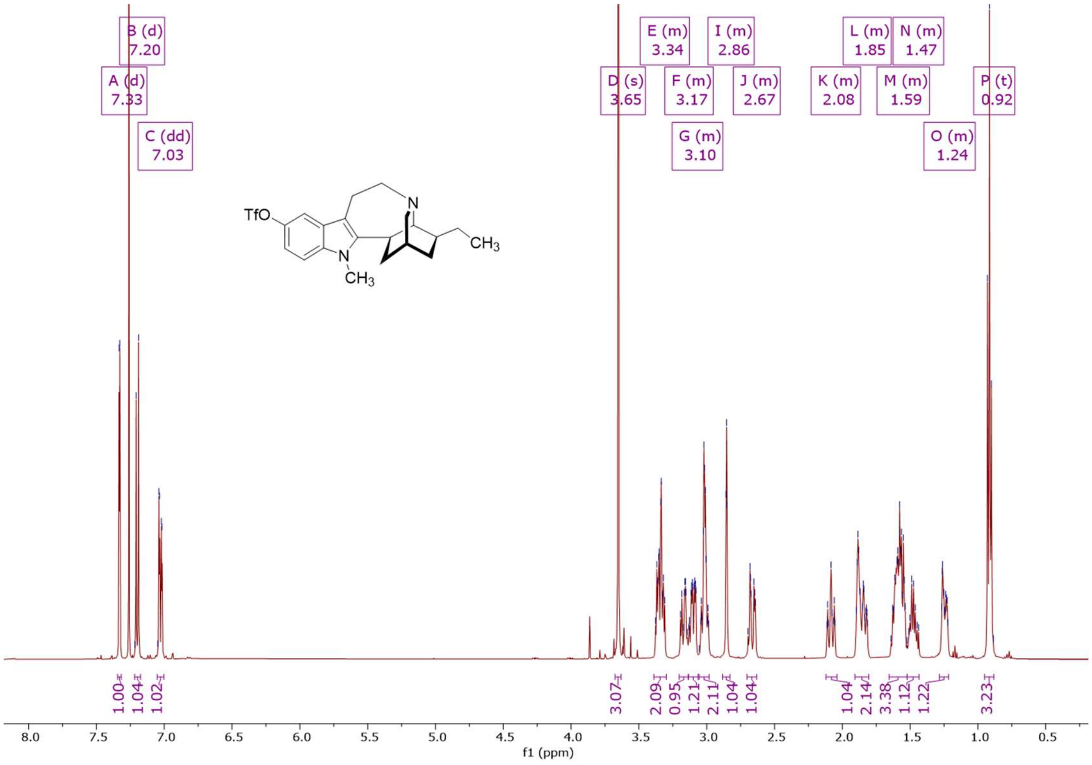


**$^1$H NMR (500 MHz, CDCl$_3$)** spectrum of *N*-methyl-5-triflate-ibogamine.

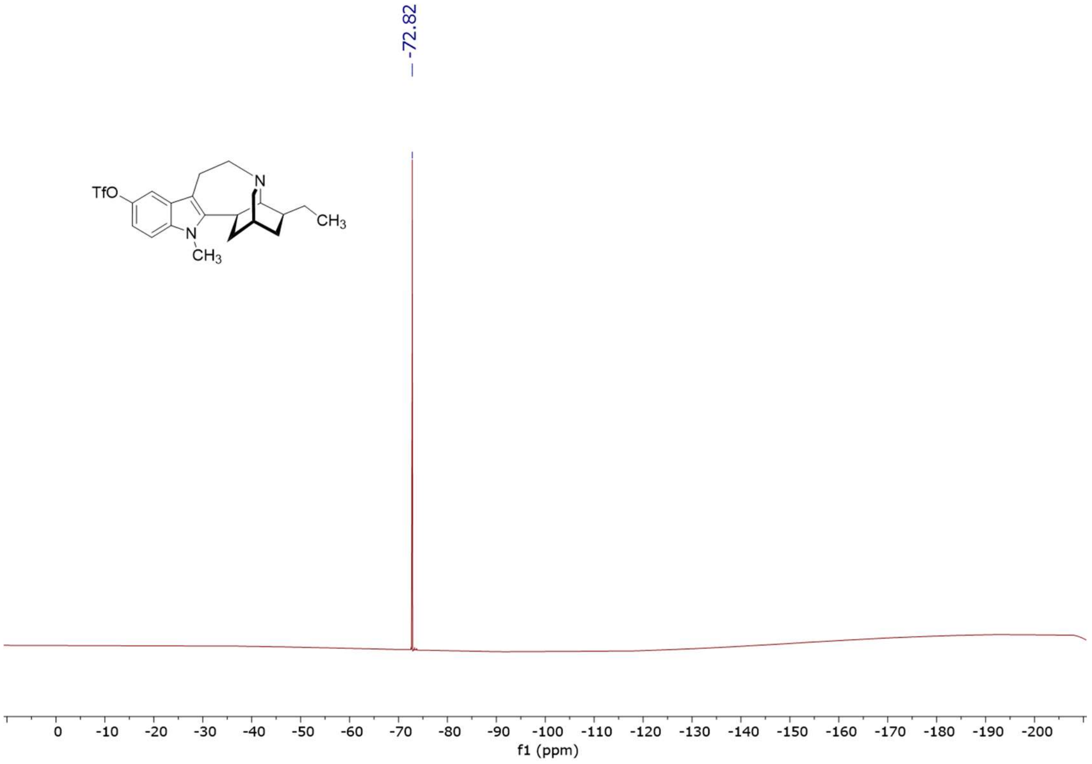


**$^{19}$F NMR (471 MHz, CDCl$_3$)** spectrum of *N*-methyl-5-triflate-ibogamine.

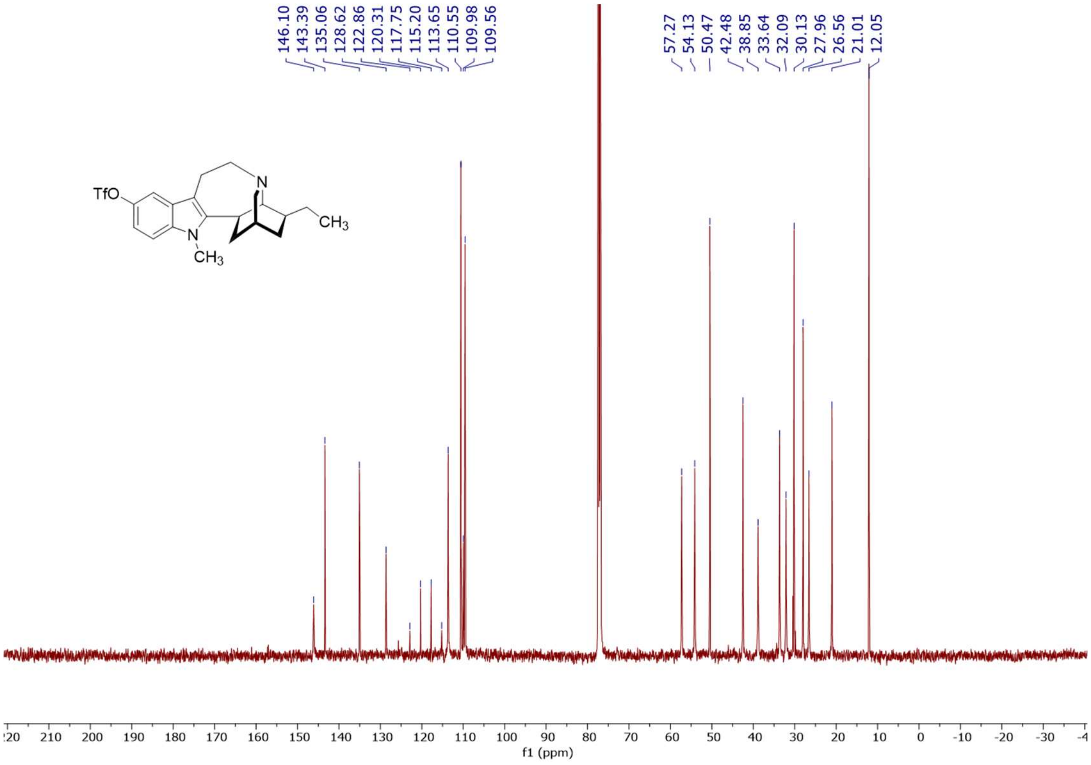


**$^{13}$C NMR (126 MHz, $CDCl_3$)** spectrum of *N*-methyl-5-triflate-ibogamine.

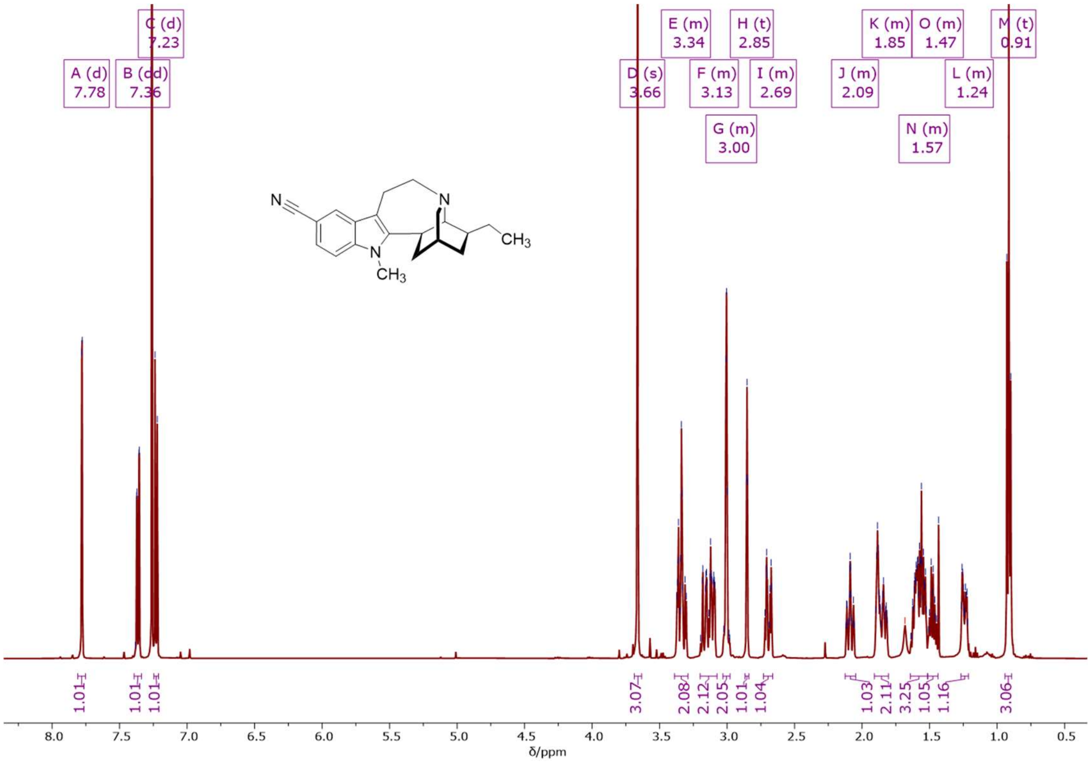


**$^{1}$H NMR (500 MHz, $CDCl_3$)** spectrum of *N*-methyl-5-cyano-ibogamine.

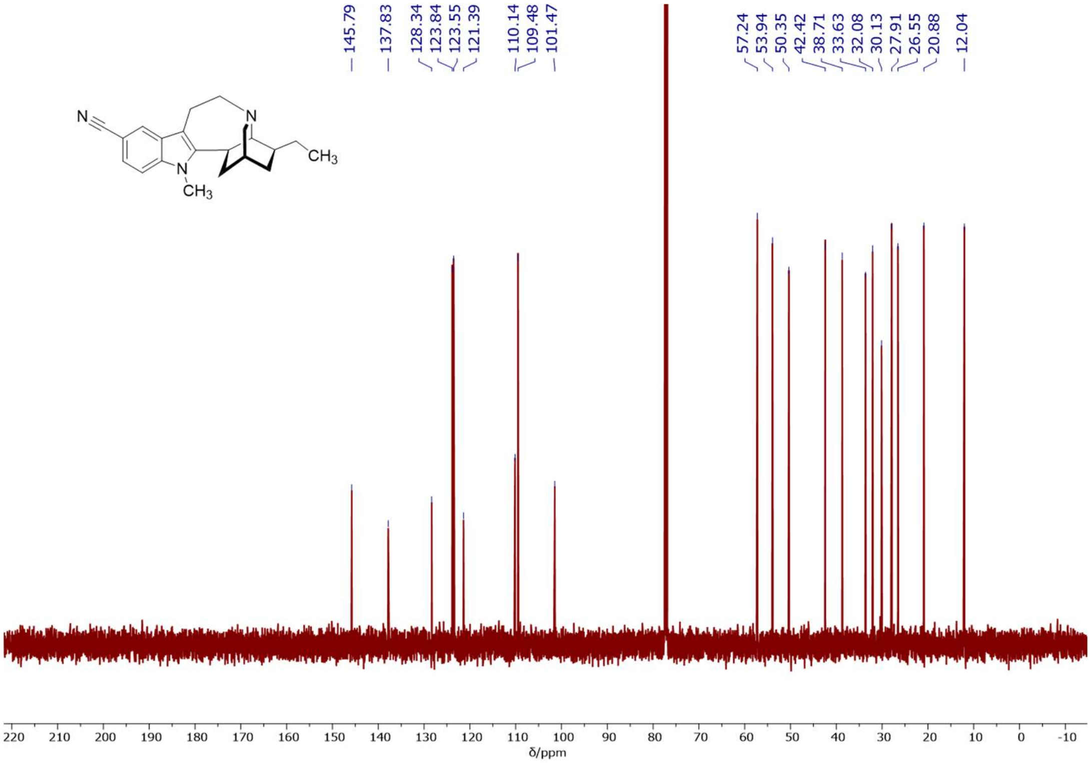


**$^{13}$C NMR (126 MHz, CDCl$_3$)** spectrum of *N*-methyl-5-cyano-ibogamine.

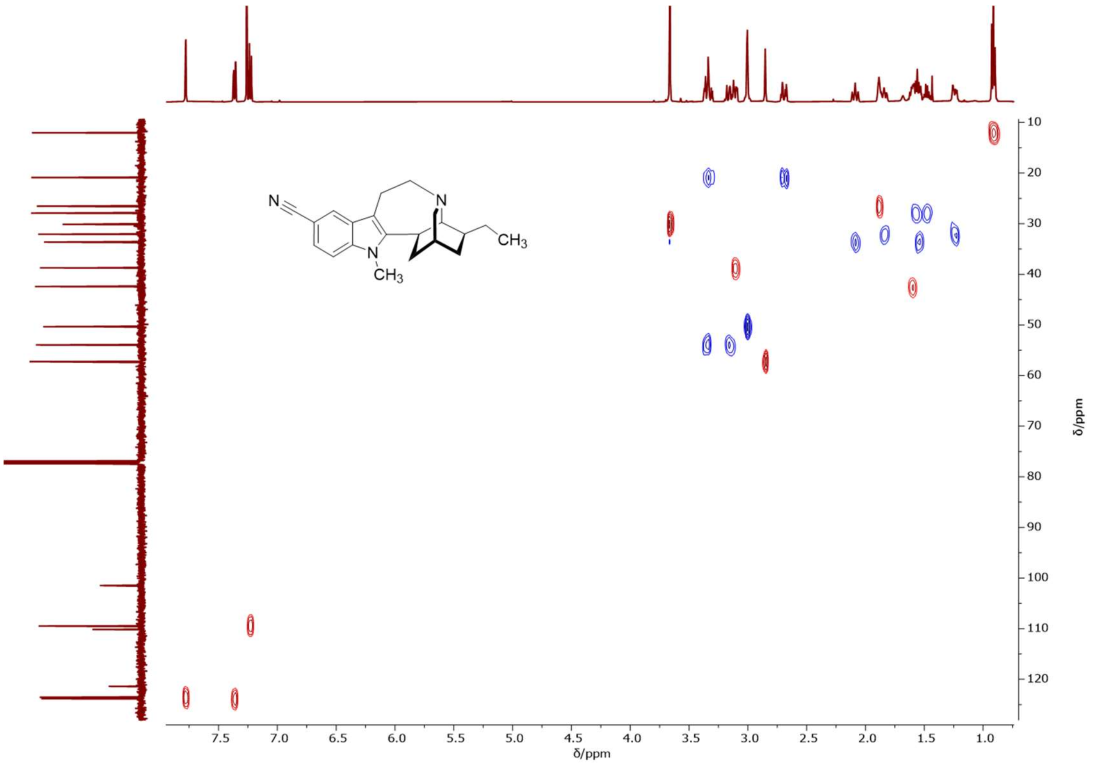


**$^{1}$H-$^{13}$C 2D NMR (500/126 MHz, MeOD) HSQC** spectrum of *N*-methyl-5-cyano-ibogamine.

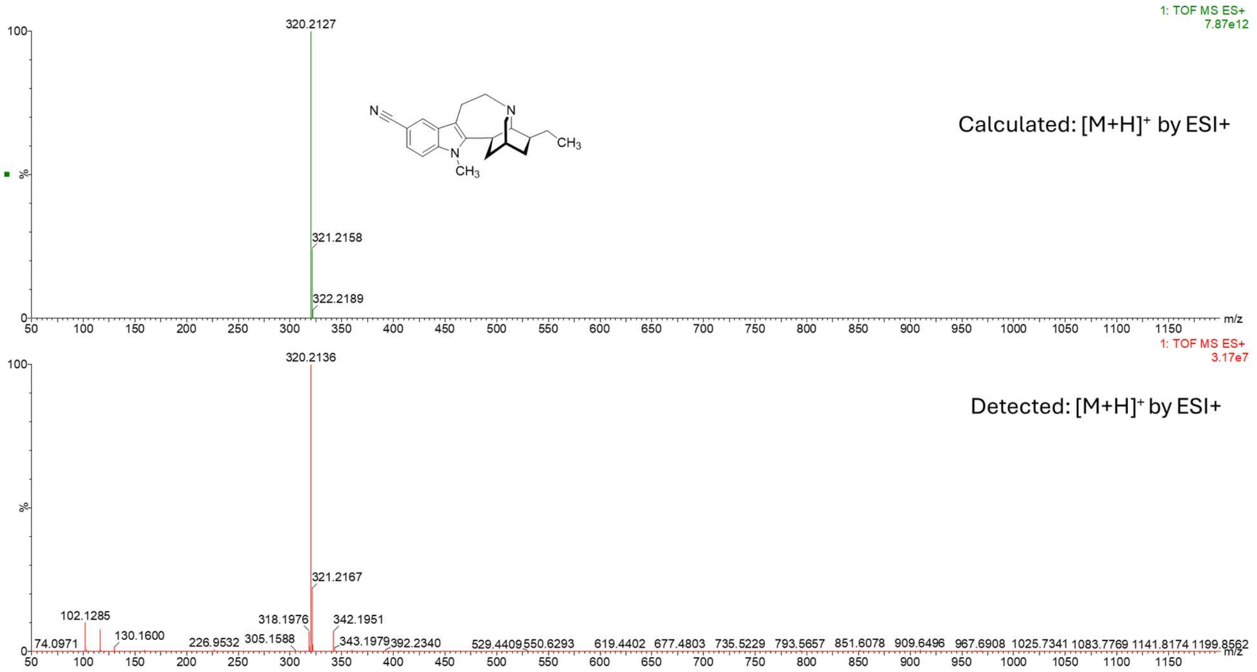


**HRMS (ESI+)** spectrum of *N*-methyl-5-cyano-ibogamine.

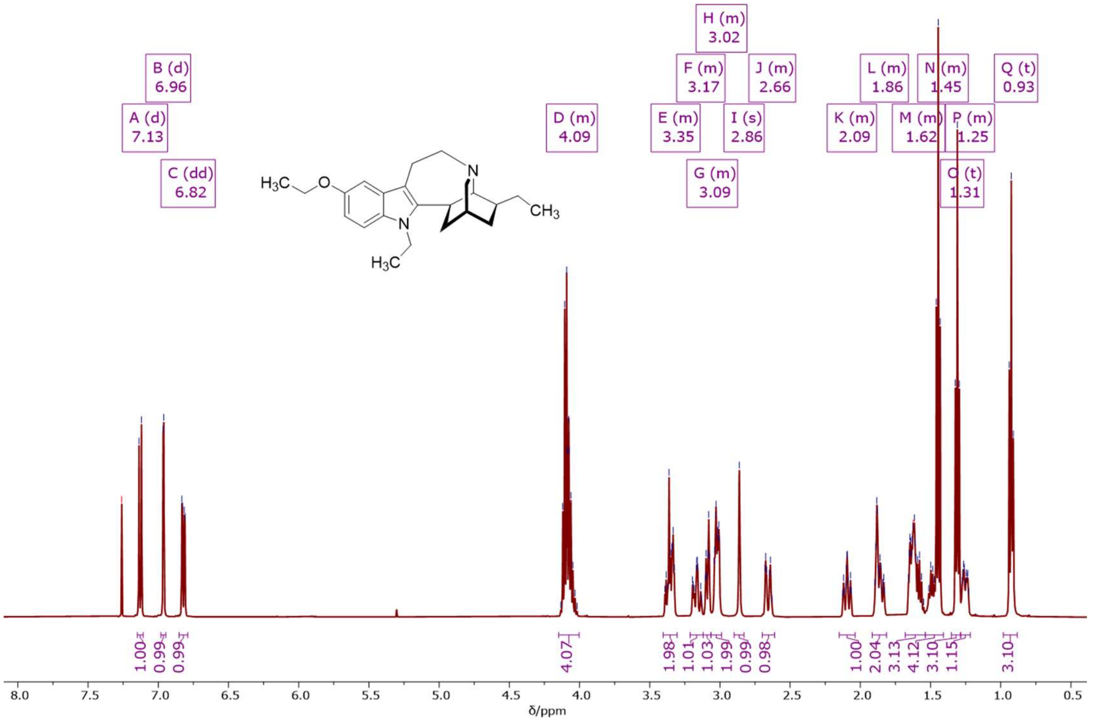


**[1]H NMR (500 MHz, $CDCl_3$)** spectrum of *N*-ethyl-5-ethoxy-ibogamine.

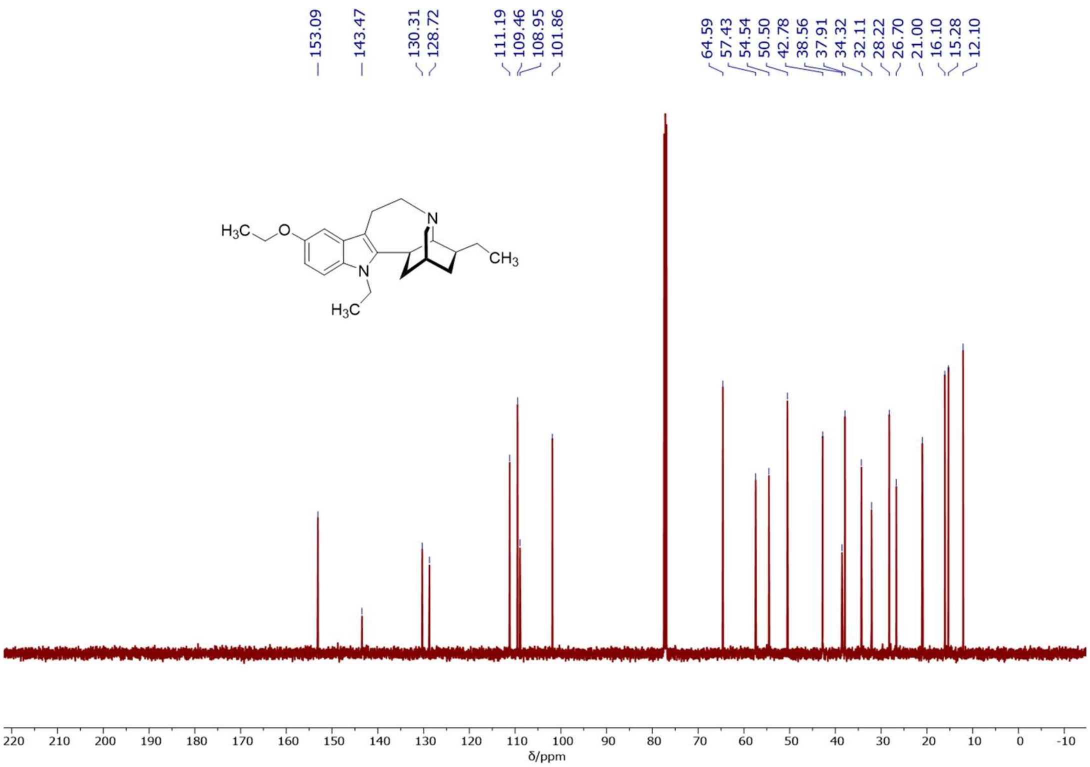


**$^{13}C$ NMR (126 MHz, $CDCl_3$)** spectrum of *N*-ethyl-5-ethoxy-ibogamine.

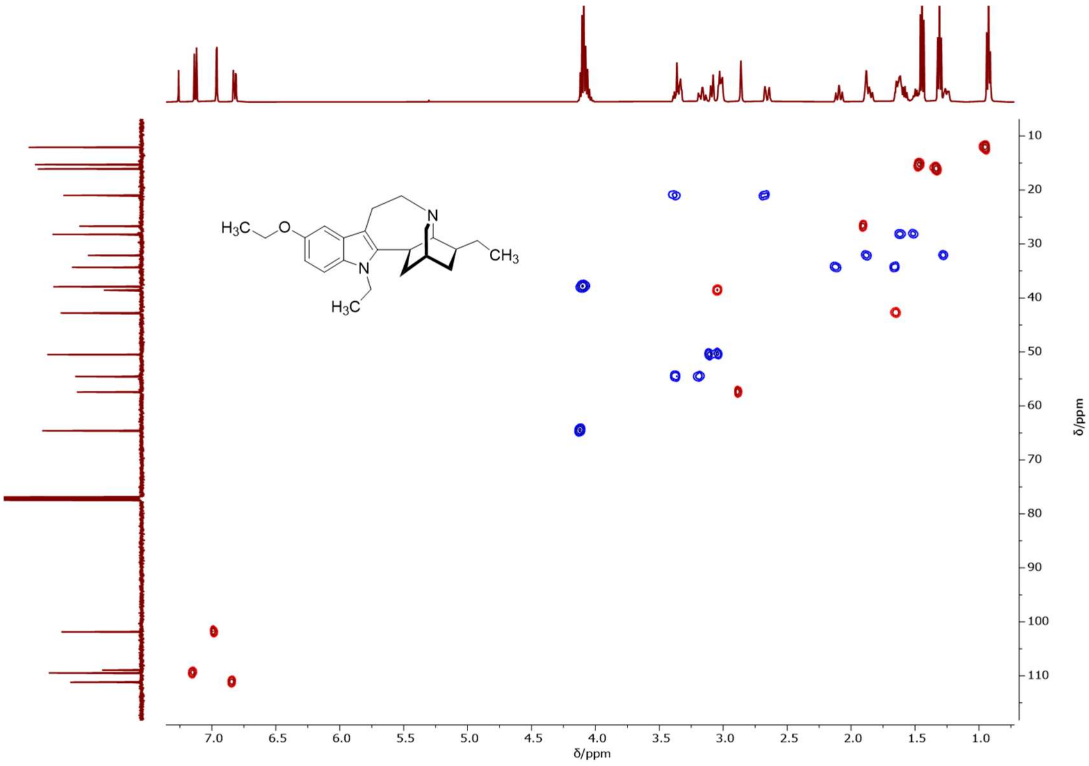


**$^{1}H$-$^{13}C$ 2D NMR (500/126 MHz, $CDCl_3$) HSQC** spectrum of *N*-ethyl-5-ethoxy-ibogamine.

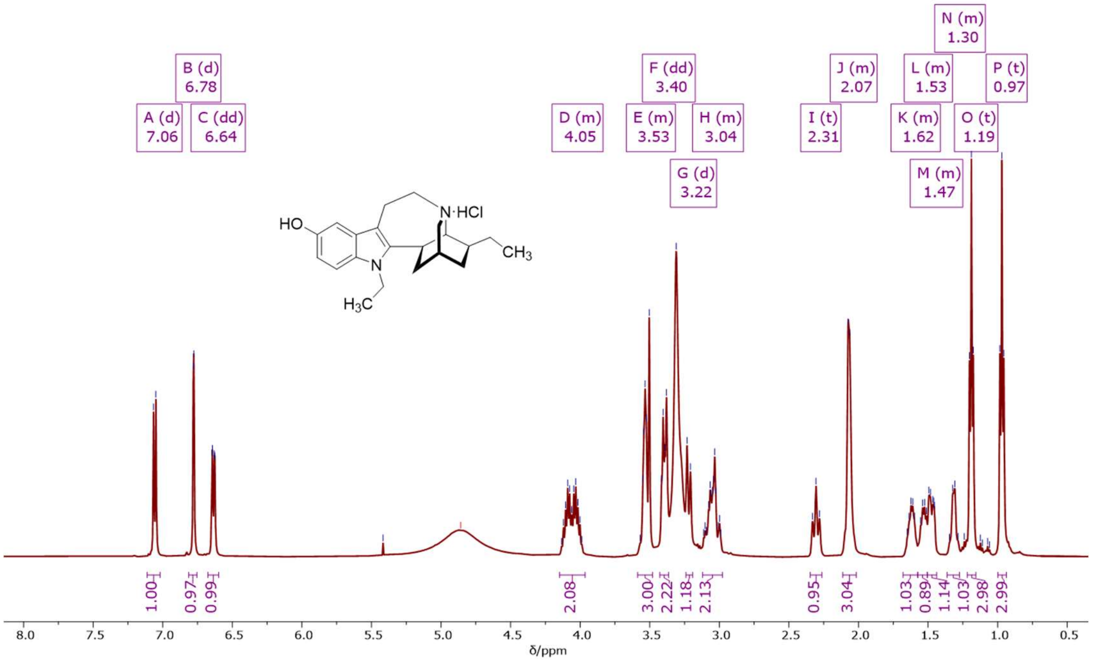


**[1]H NMR (500 MHz, MeOD)** spectrum of *N*-ethyl-noribogaine hydrochloride.

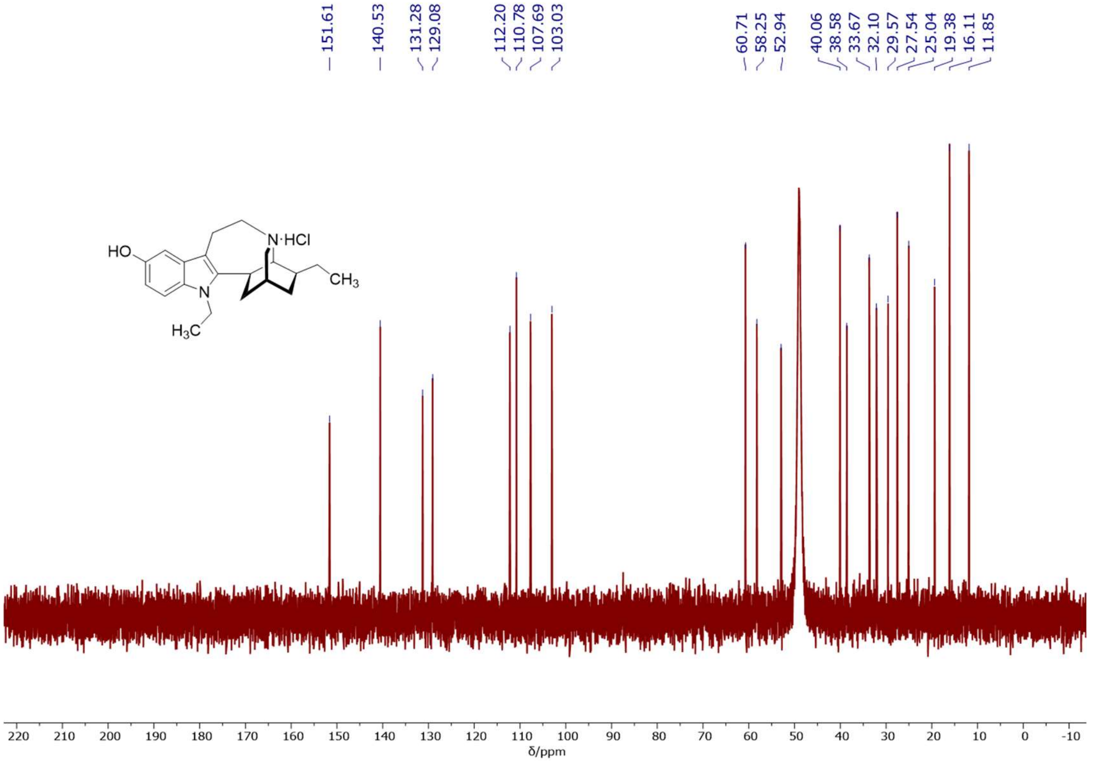


**[13]C NMR (126 MHz, MeOD)** spectrum of *N*-ethyl-noribogaine hydrochloride.

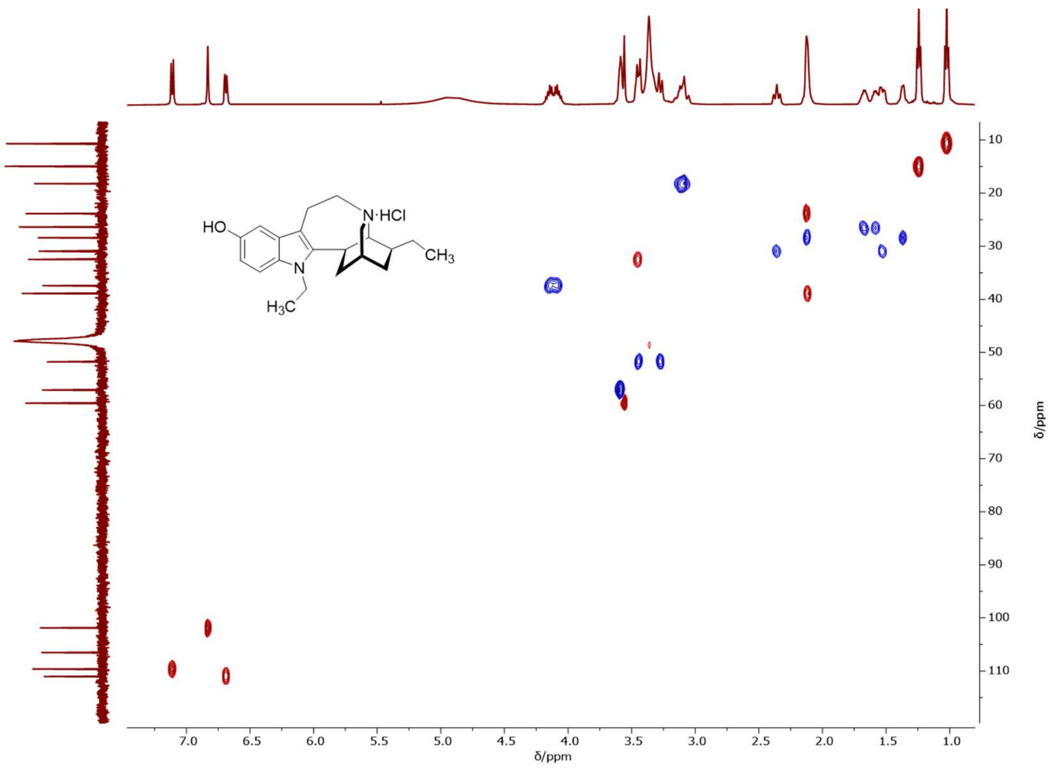

$^{1}H$-$^{13}C$ **2D NMR (500/126 MHz, MeOD) HSQC** spectrum of *N*-ethyl-noribogaine hydrochloride.

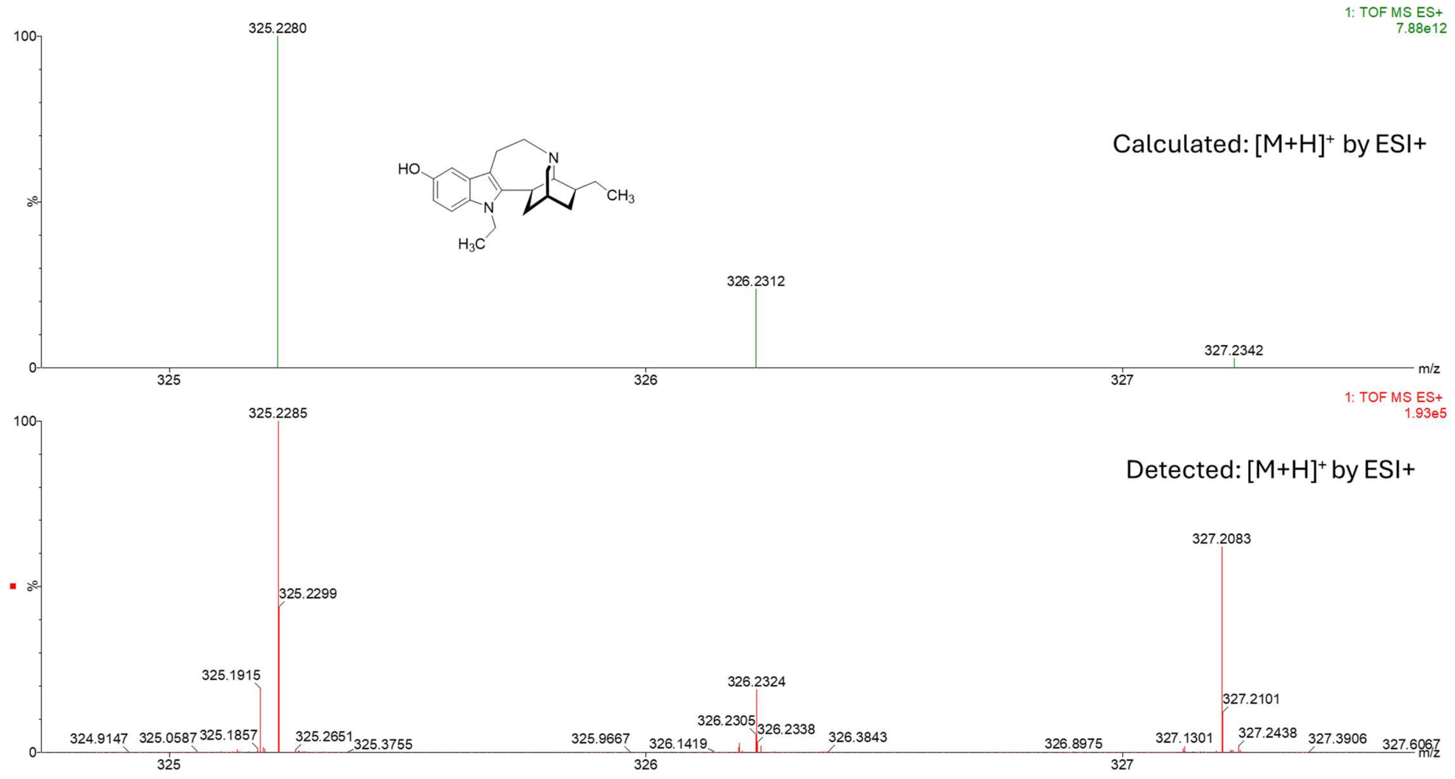


**HRMS (ESI+)** spectrum of *N*-ethyl-noribogaine.

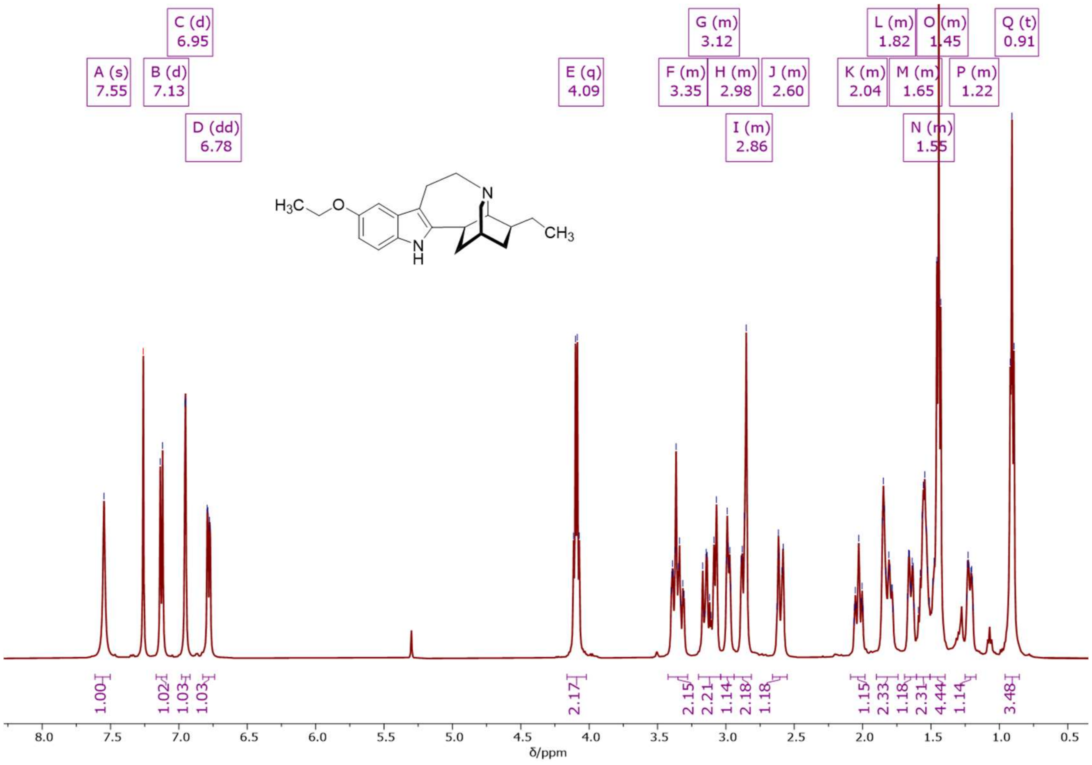


**[1]H NMR (500 MHz, CDCl$_3$)** spectrum of 5-ethoxy-ibogamine.

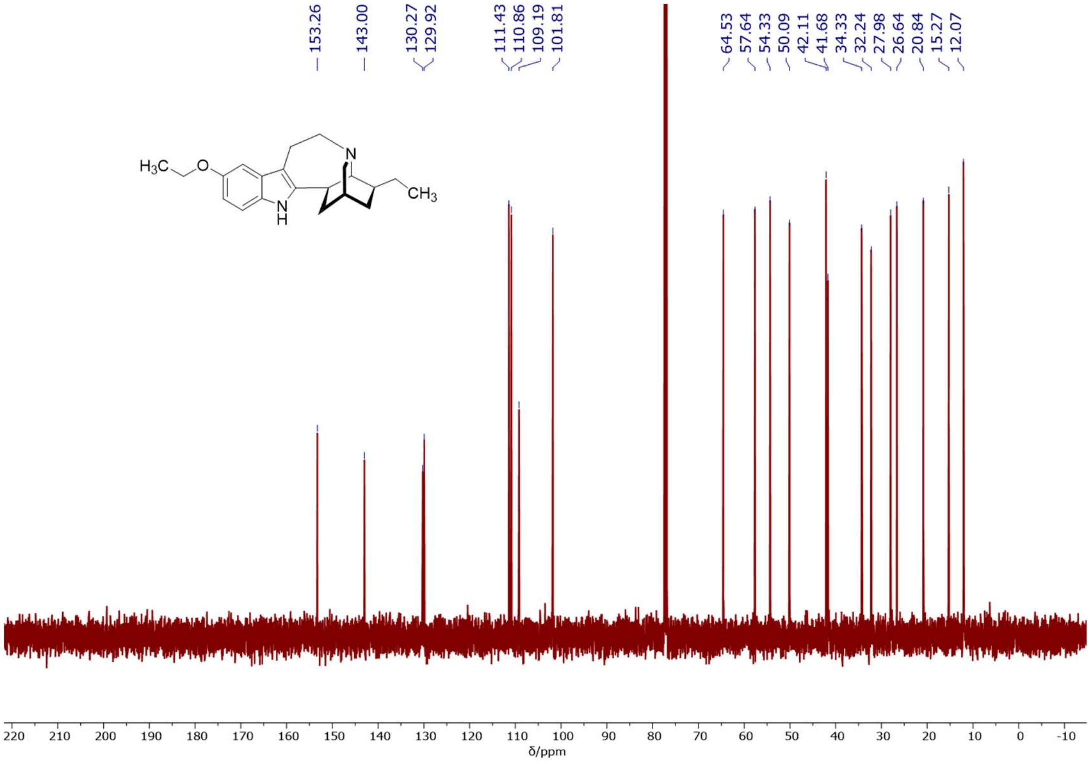


**[13]C NMR (126 MHz, CDCl$_3$)** spectrum of 5-ethoxy-ibogamine.

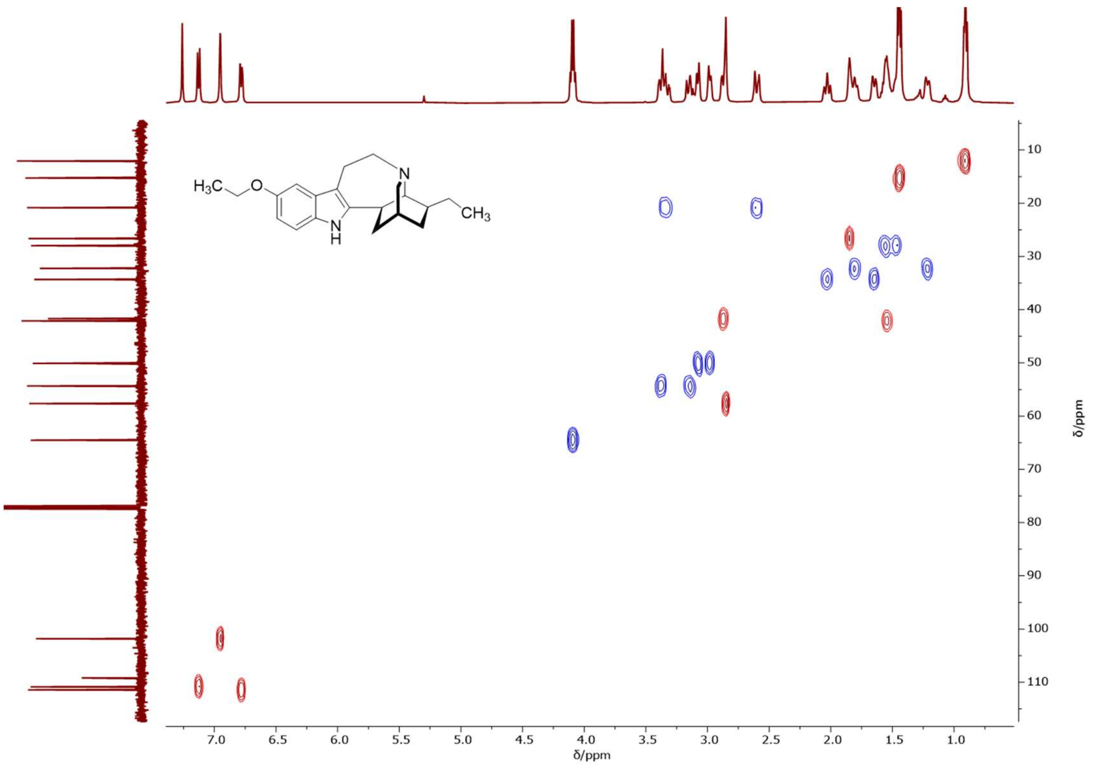


**¹H-¹³C 2D NMR (500/126 MHz, CDCl₃) HSQC** spectrum of 5-ethoxy-ibogamine.

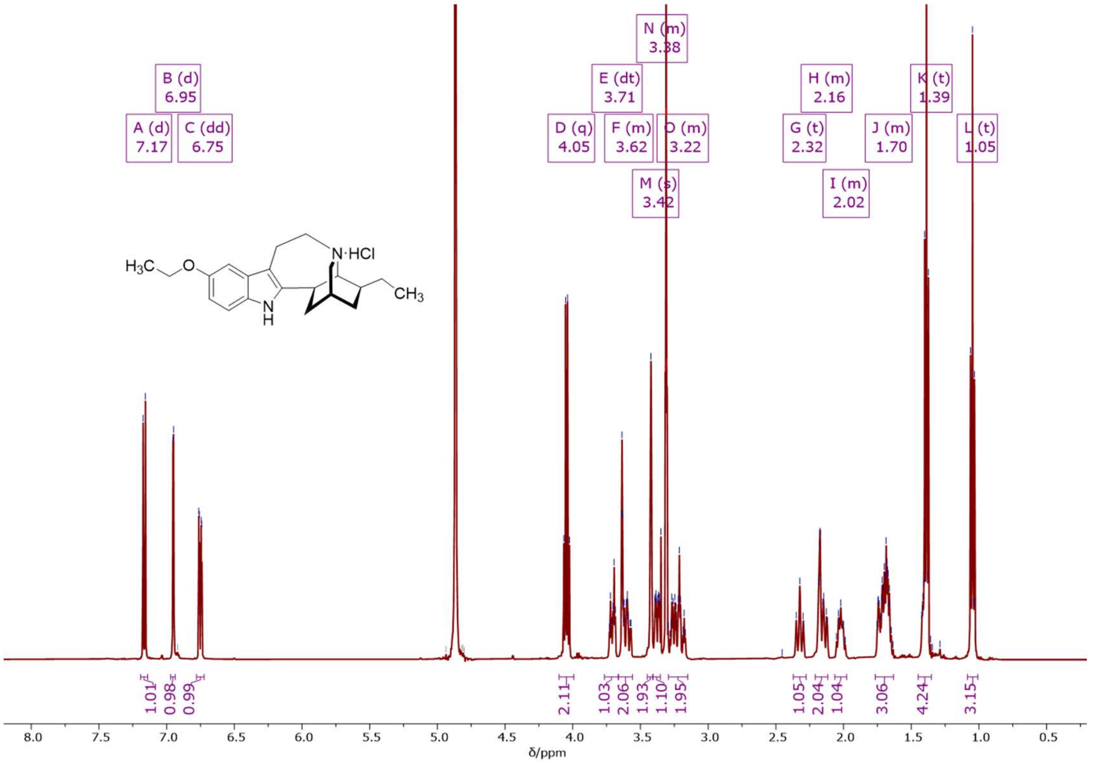


**¹H NMR (500 MHz, MeOD)** spectrum of 5-ethoxy-ibogamine hydrochloride.

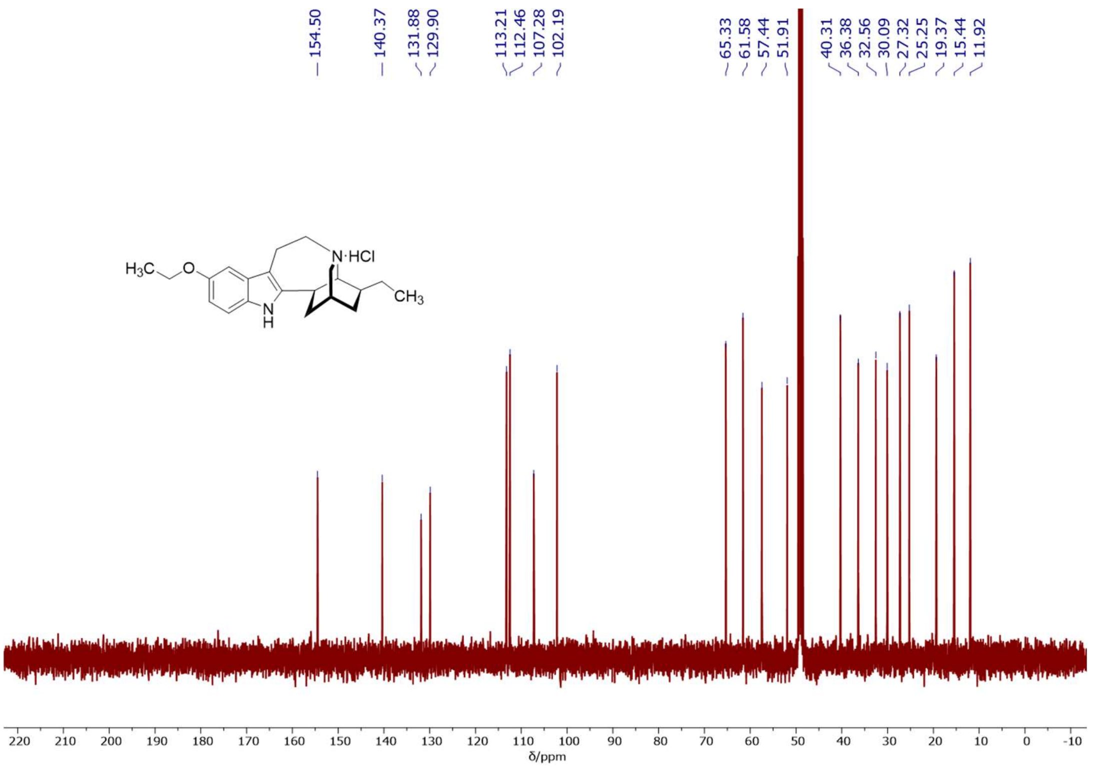


**$^{13}C$ NMR (126 MHz, MeOD)** spectrum of 5-ethoxy-ibogamine hydrochloride.

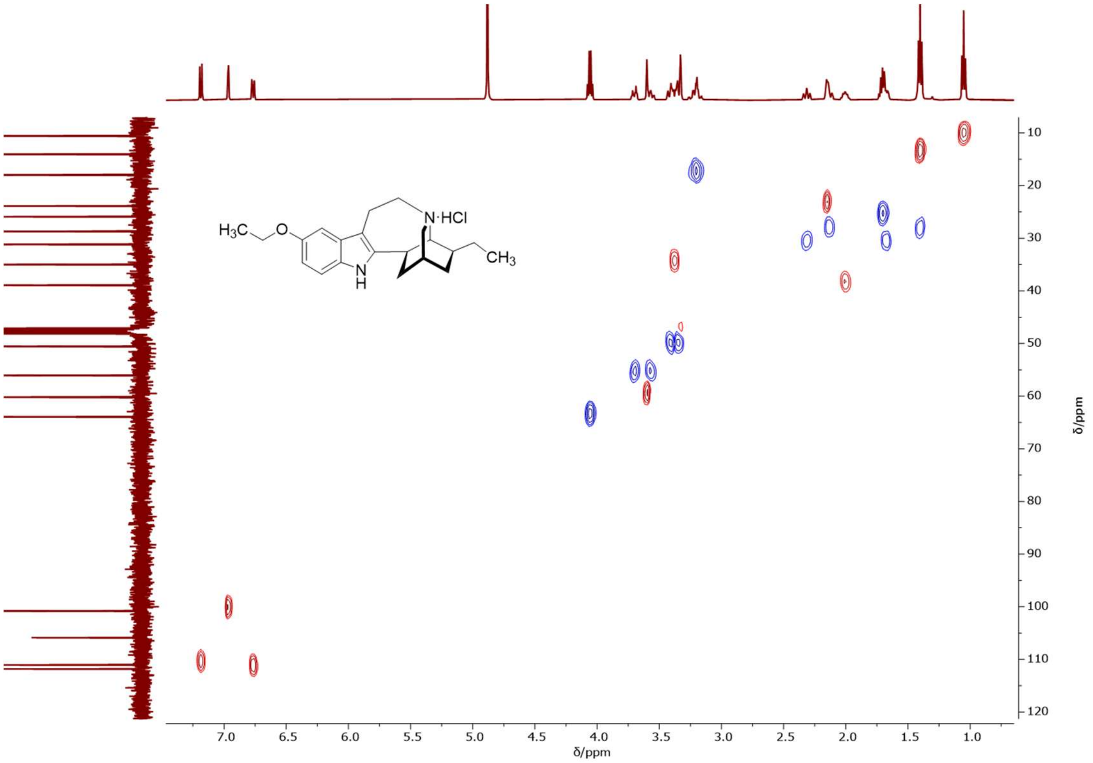


**$^{1}H$-$^{13}C$ 2D NMR (500/126 MHz, MeOD) HSQC** spectrum of 5-ethoxy-ibogamine hydrochloride.

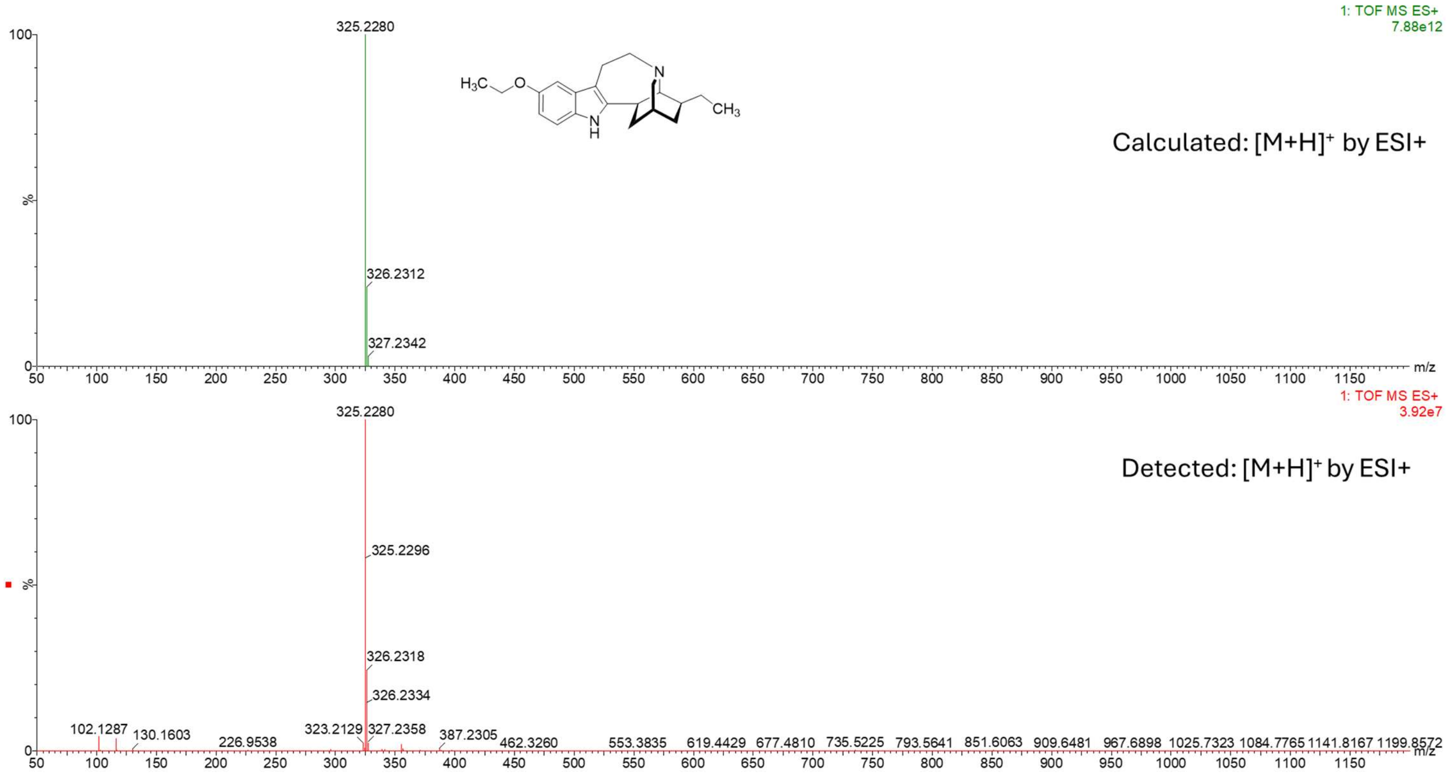


**HRMS (ESI+)** spectrum of 5-ethoxy-ibogamine.

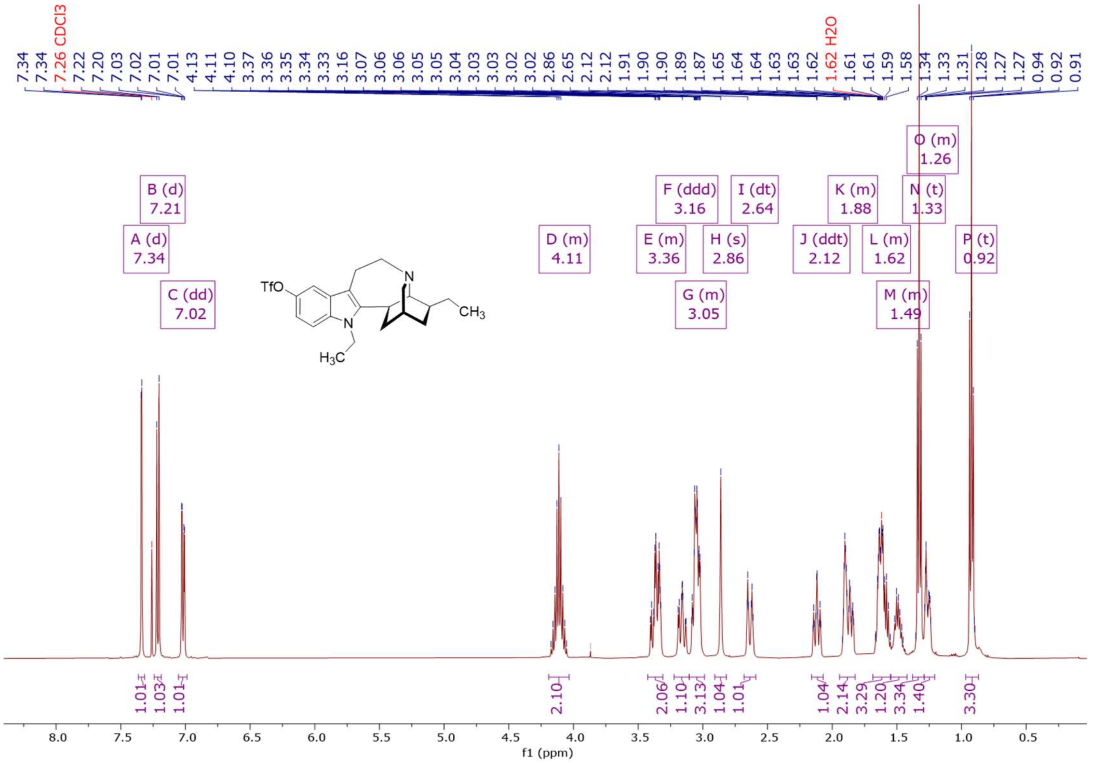


**$^1$H NMR (500 MHz, $CDCl_3$)** spectrum of *N*-ethyl-5-triflate-ibogamine.

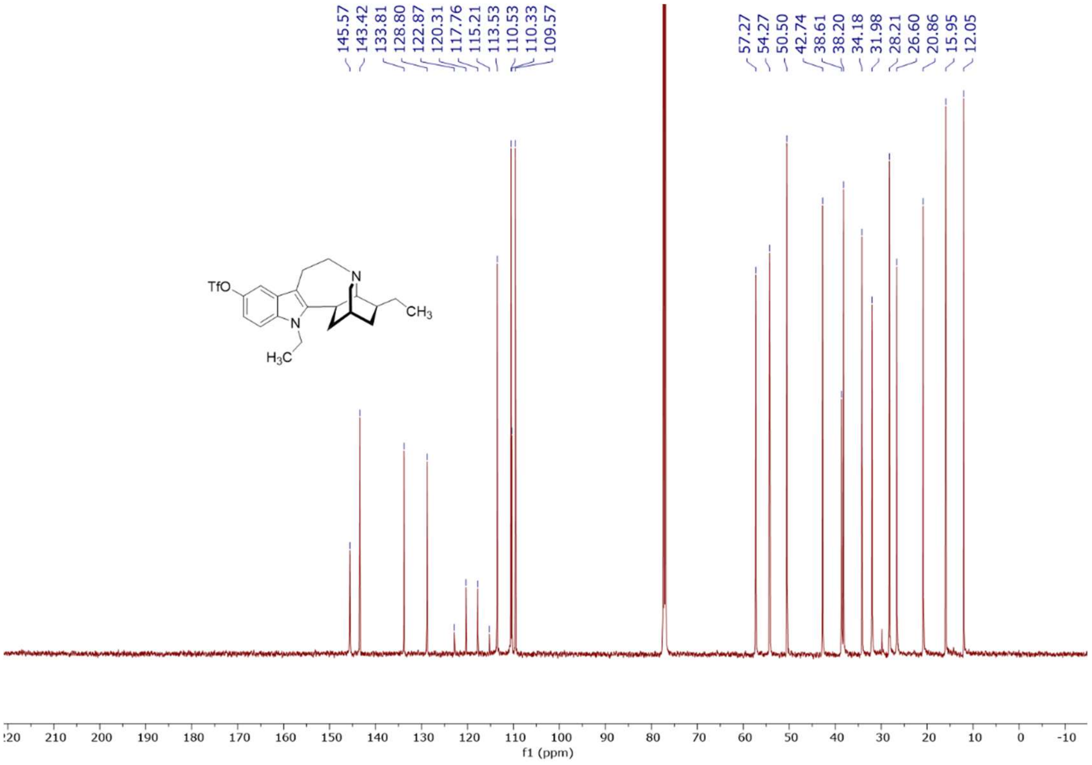


**$^{13}C$ NMR (126 MHz, $CDCl_3$)** spectrum of *N*-ethyl-5-triflate-ibogamine.

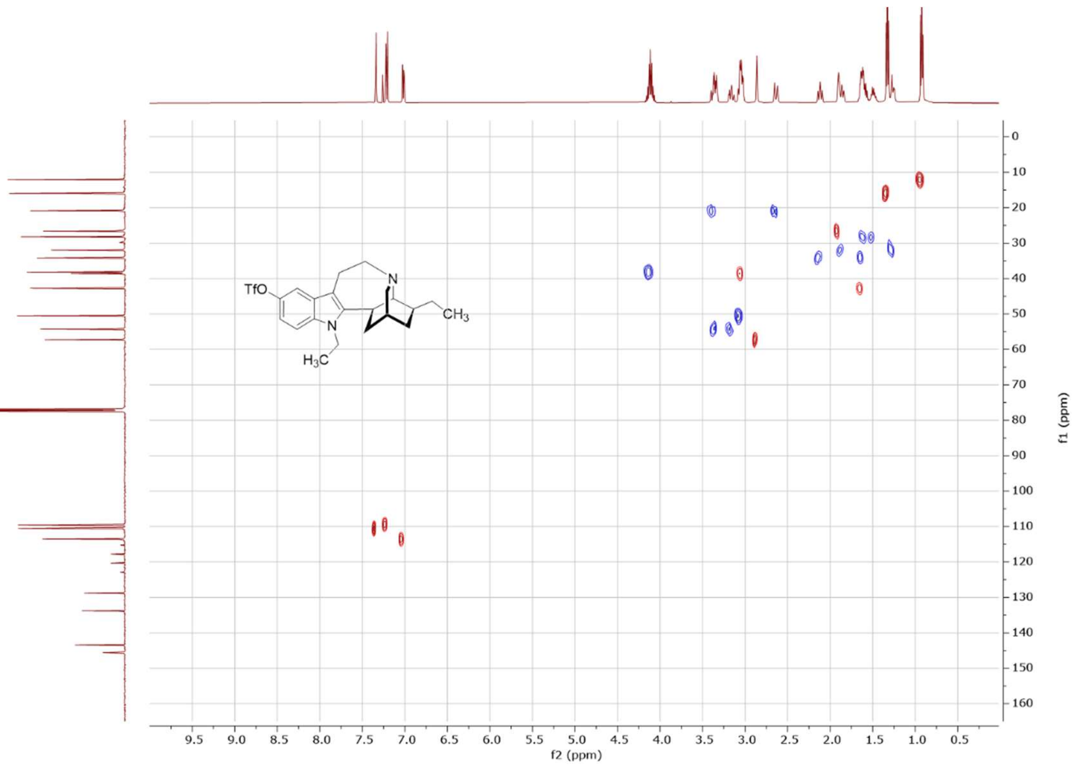


**$^{1}H$-$^{13}C$ 2D NMR (500/126 MHz, $CDCl_3$) HSQC** spectrum of *N*-ethyl-5-triflate-ibogamine.

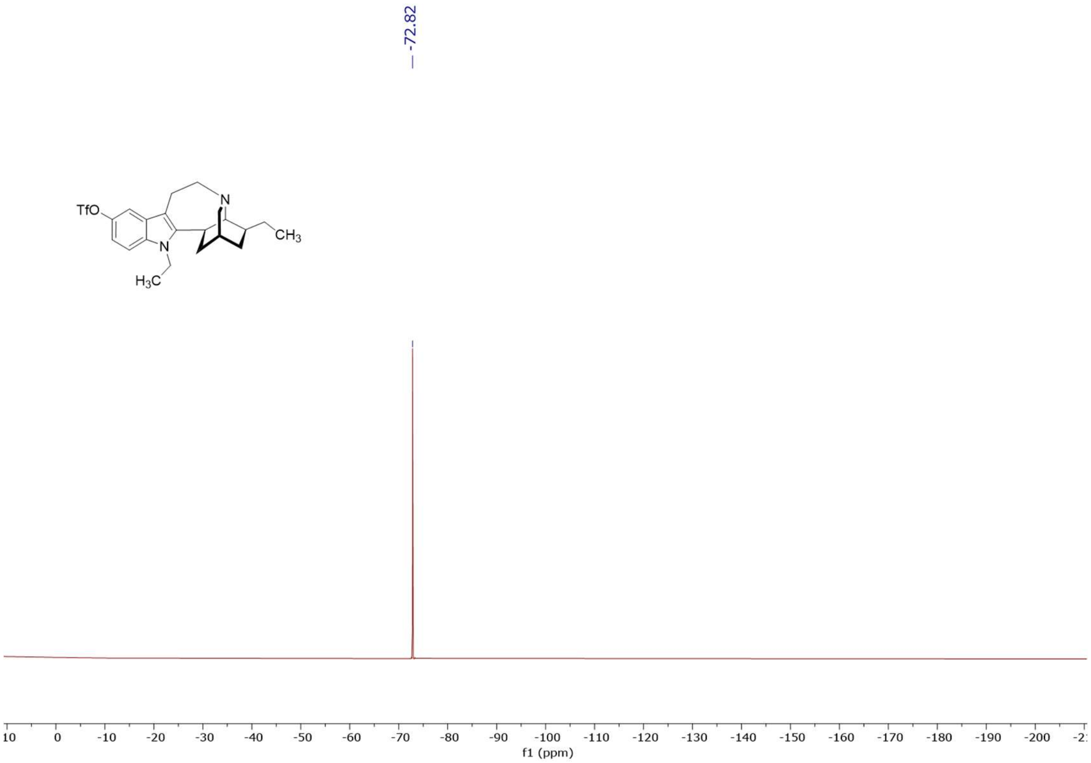


**[19]F NMR (470 MHz, CDCl₃)** spectrum of *N*-ethyl-5-triflate-ibogamine.

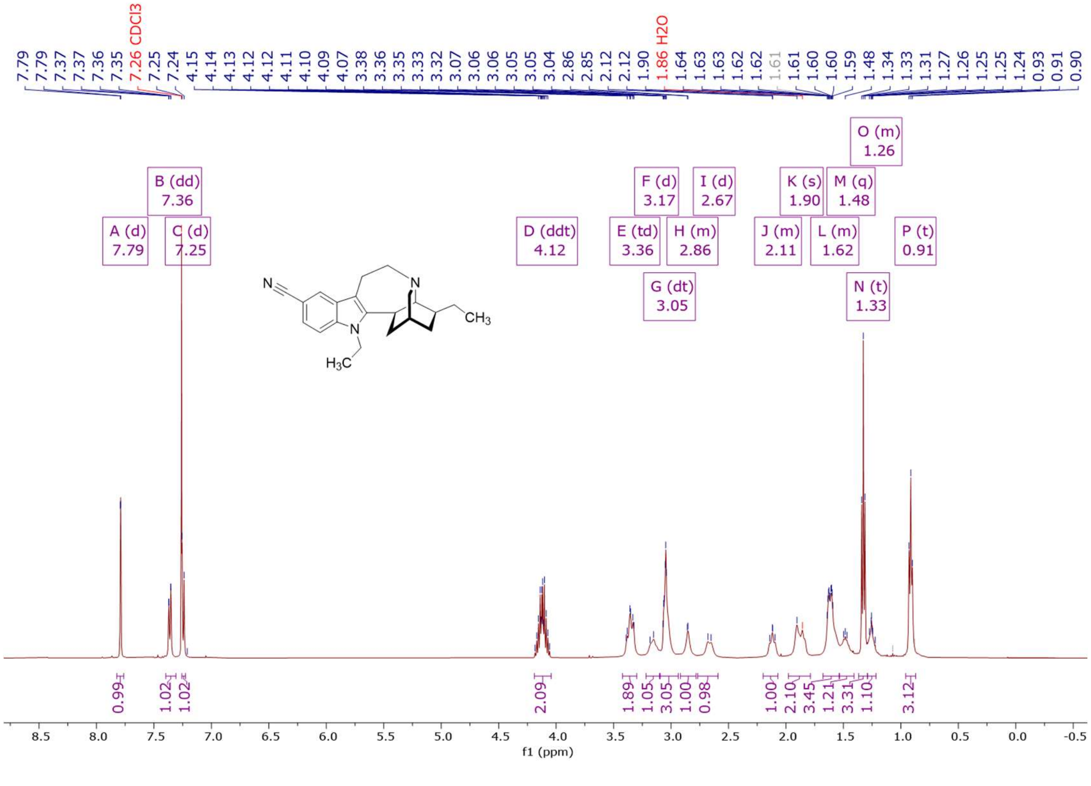


**[1]H NMR (500 MHz, CDCl₃)** spectrum of *N*-ethyl-5-cyano-ibogamine.

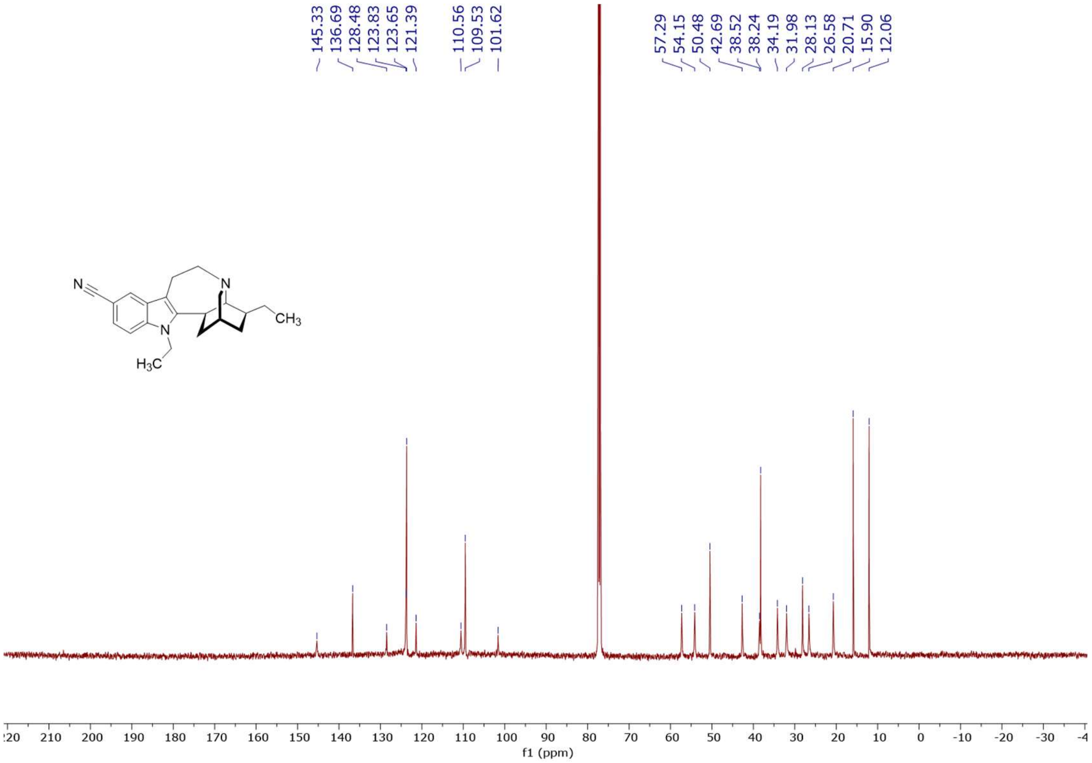


**$^{13}C$ NMR (126 MHz, $CDCl_3$)** spectrum of *N*-ethyl-5-cyano-ibogamine.

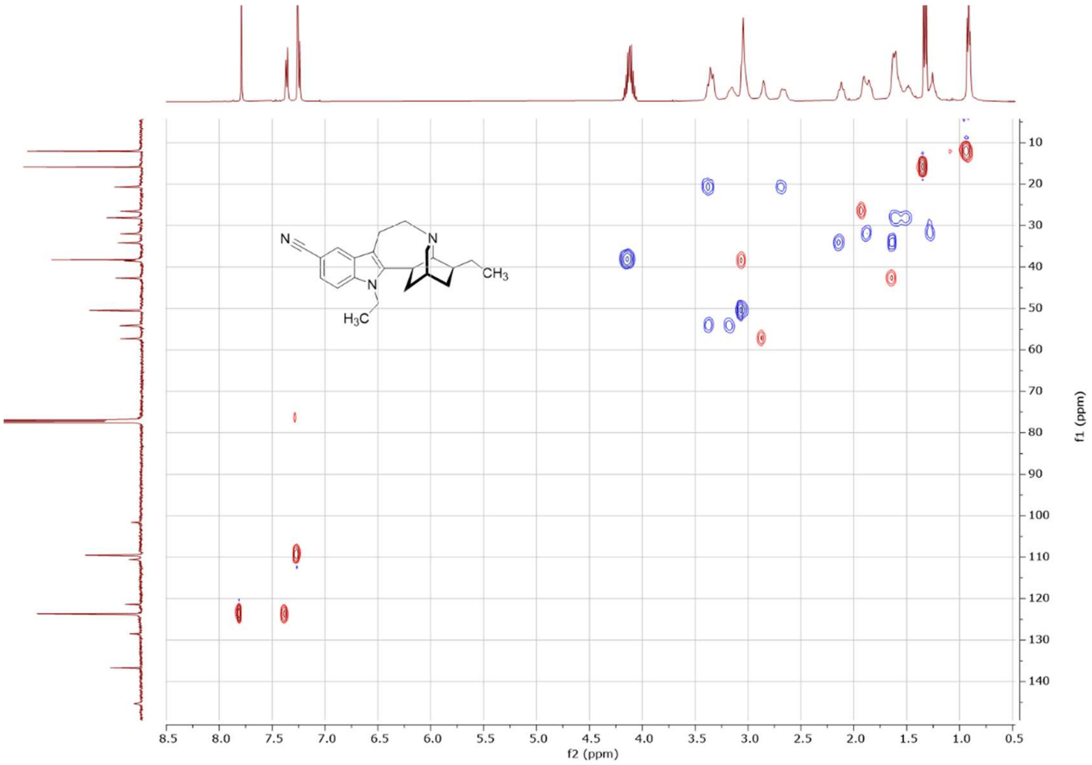


**$^{1}H$-$^{13}C$ 2D NMR (500/126 MHz, $CDCl_3$) HSQC** spectrum of *N*-ethyl-5-cyano-ibogamine.

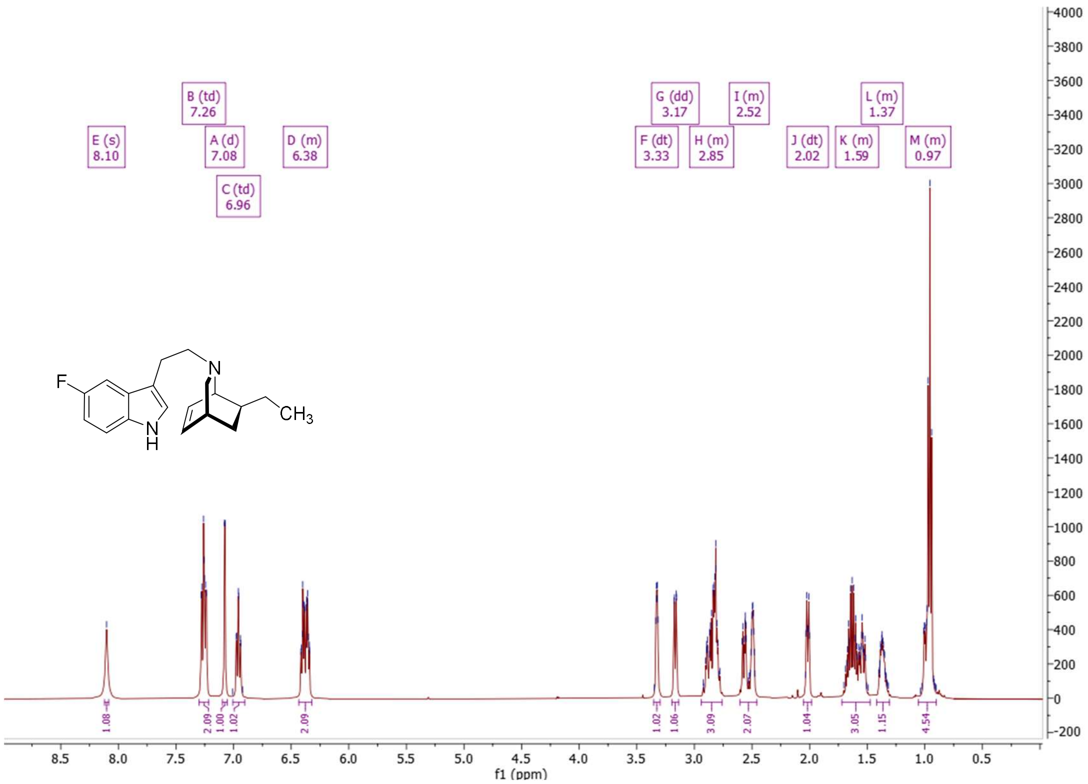


**$^{1}H$ NMR (500 MHz, $CDCl_3$)** spectrum of (1S,4R,7S)-7-ethyl-2-(2-(5-fluoro-1H-indol-3-yl)ethyl)-2-azabicyclo[2.2.2]oct-5-ene.

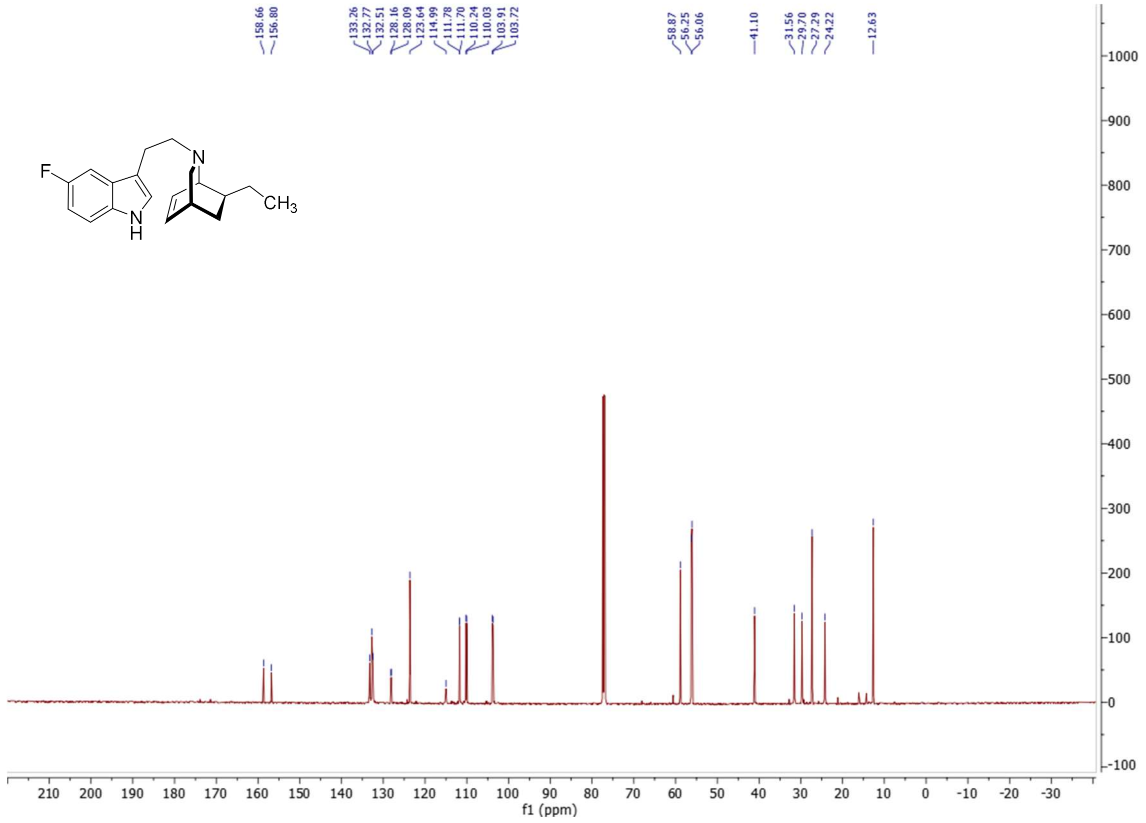


**$^{13}$C NMR (126 MHz, $CDCl_3$)** spectrum of (1S,4R,7S)-7-ethyl-2-(2-(5-fluoro-1H-indol-3-yl)ethyl)-2-azabicyclo[2.2.2]oct-5-ene.

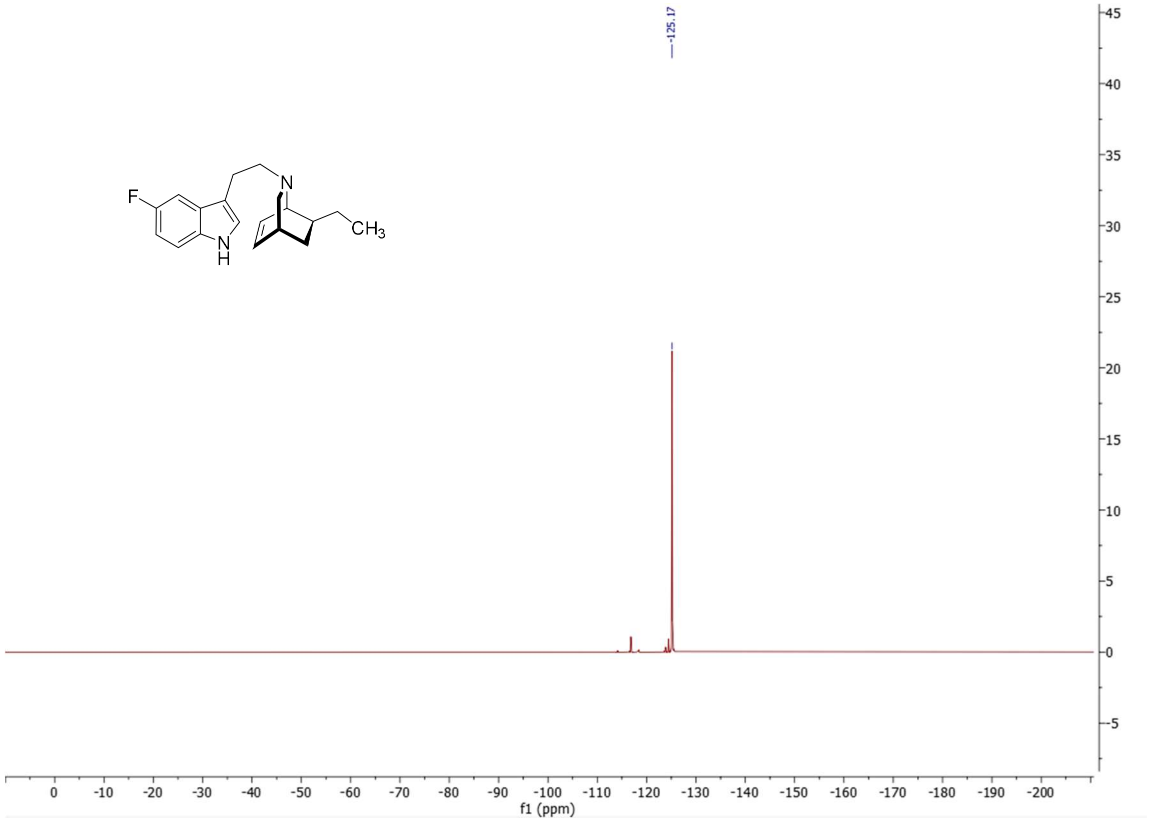


**$^{19}$F NMR (471 MHz, $CDCl_3$)** spectrum of (1S,4R,7S)-7-ethyl-2-(2-(5-fluoro-1H-indol-3-yl)ethyl)-2-azabicyclo[2.2.2]oct-5-ene.

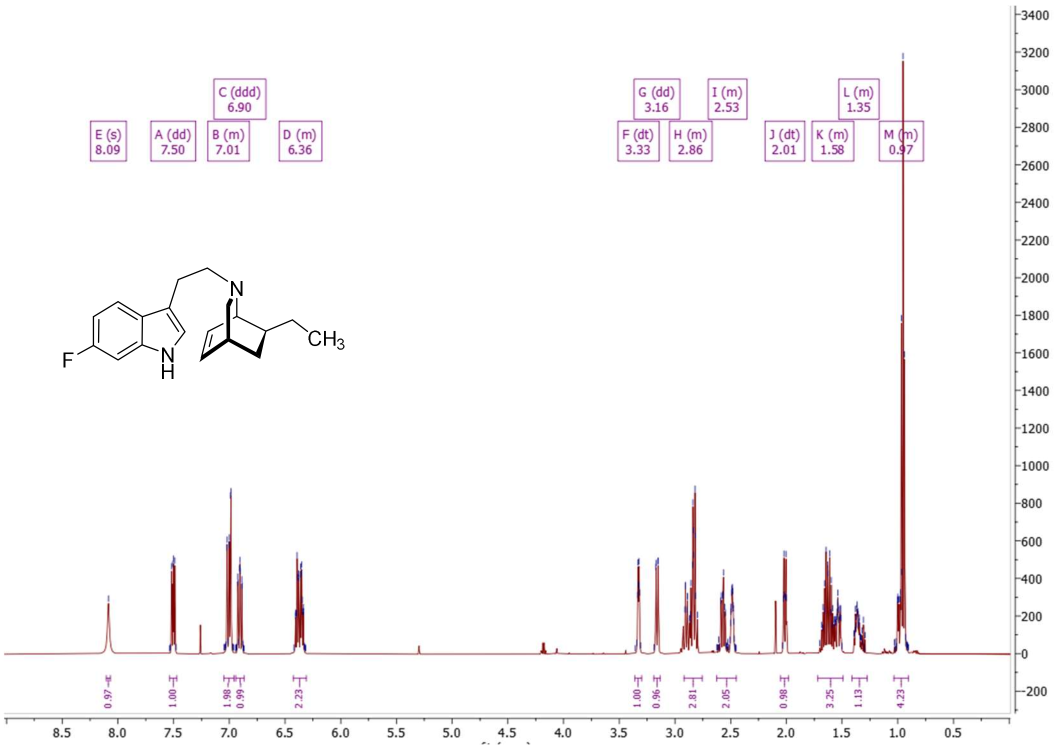


**$^1H$ NMR (500 MHz, $CDCl_3$)** spectrum of (1S,4R,7S)-7-ethyl-2-(2-(6-fluoro-1H-indol-3-yl)ethyl)-2-azabicyclo[2.2.2]oct-5-ene.

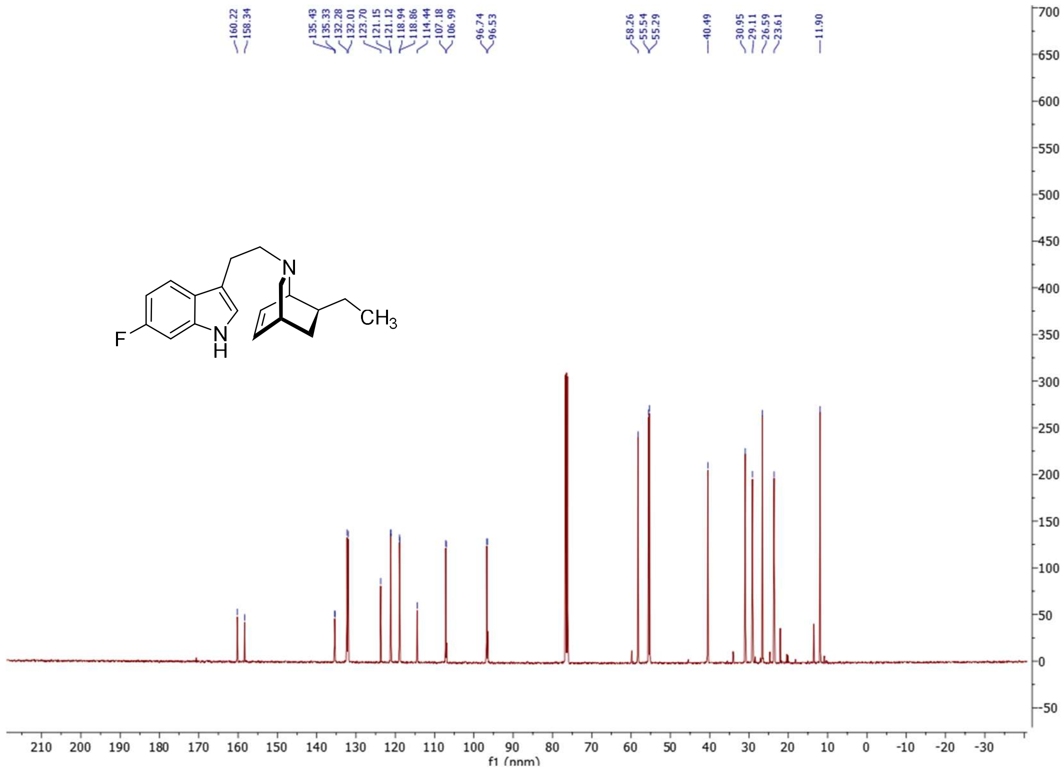


**$^{13}C$ NMR (126 MHz, $CDCl_3$)** spectrum of (1S,4R,7S)-7-ethyl-2-(2-(6-fluoro-1H-indol-3-yl)ethyl)-2-azabicyclo[2.2.2]oct-5-ene.

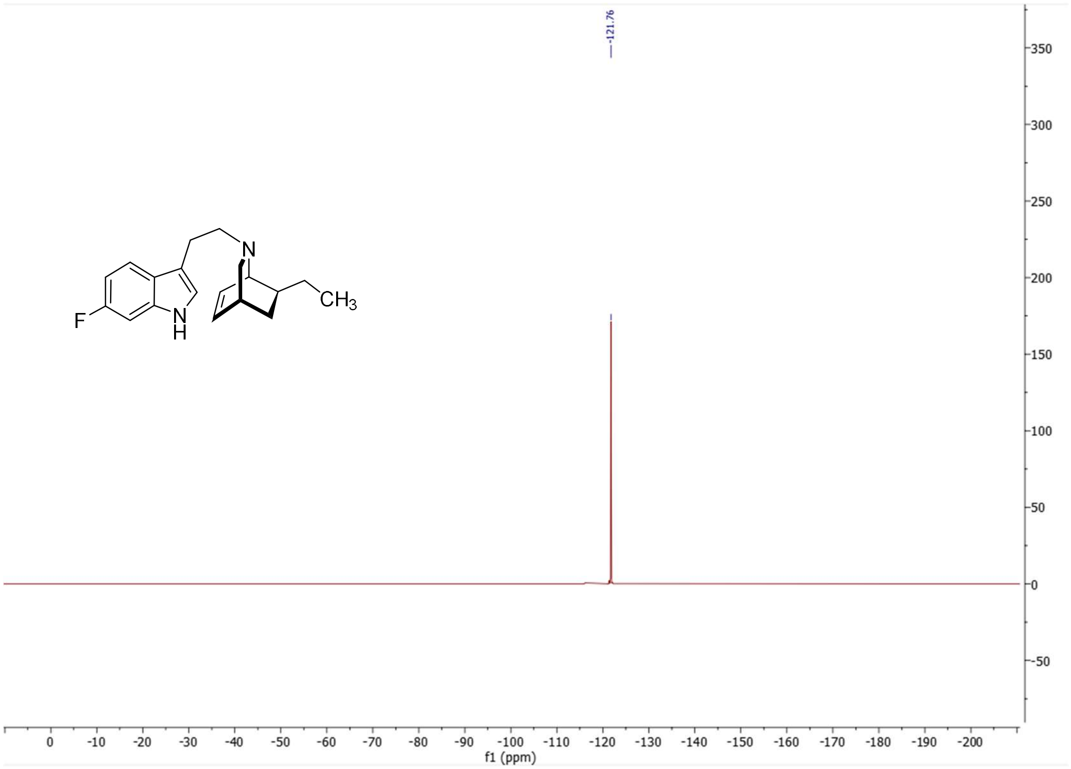


**$^{19}$F NMR (471 MHz, CDCl$_3$)** spectrum of (1S,4R,7S)-7-ethyl-2-(2-(6-fluoro-1H-indol-3-yl)ethyl)-2-azabicyclo[2.2.2]oct-5-ene.

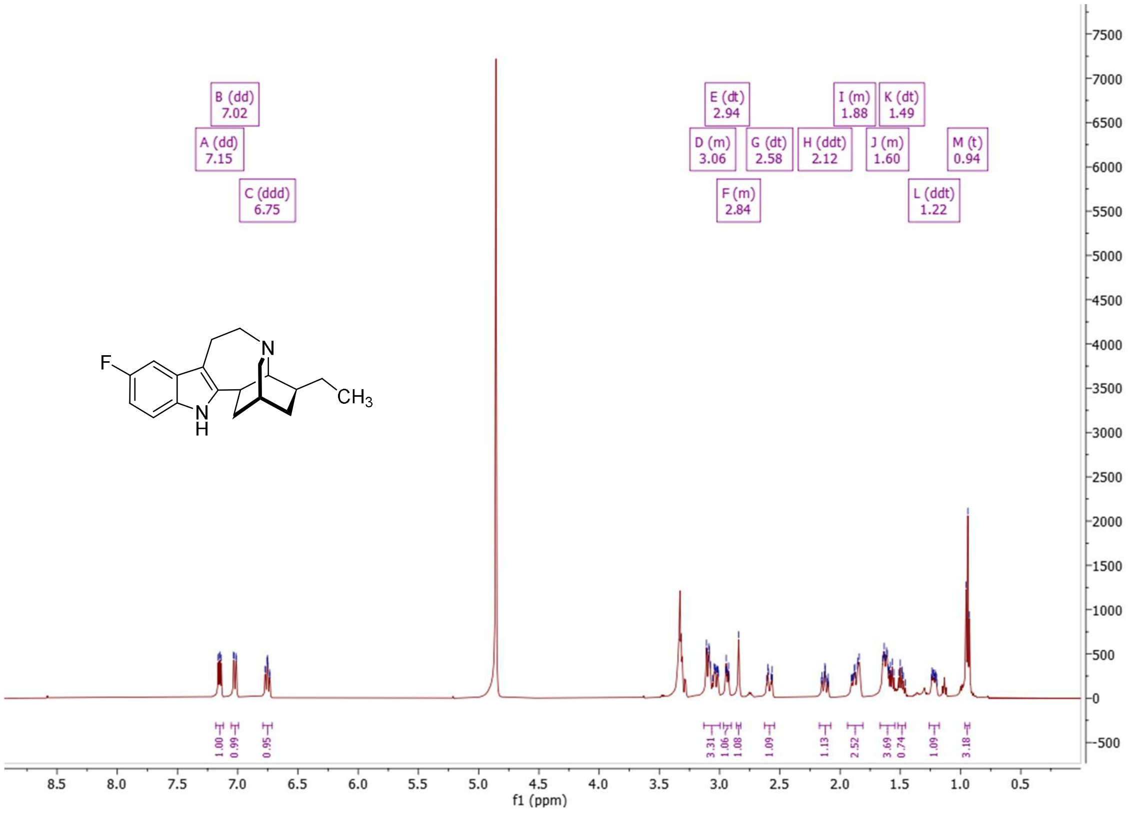


**$^{1}$H NMR (500 MHz, MeOD)** spectrum of 5-fluoro-ibogamine.

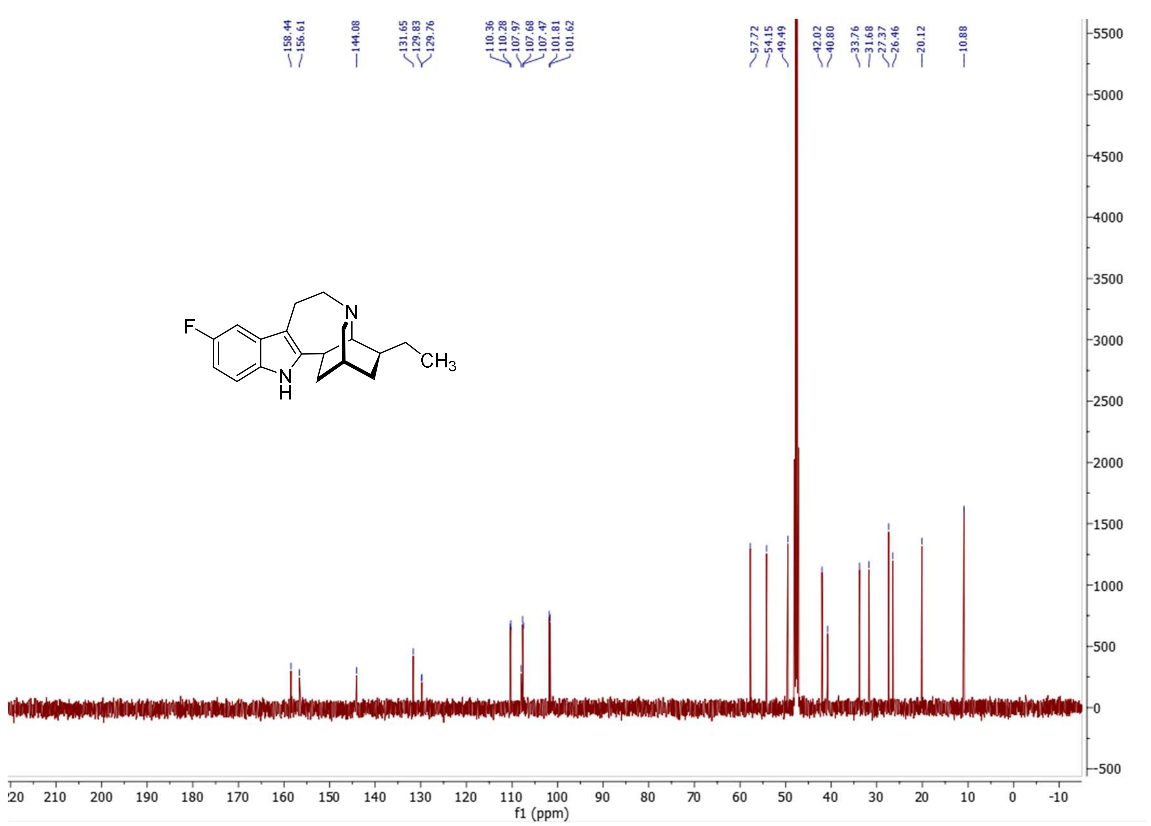


**$^{13}C$ NMR (126 MHz, MeOD)** spectrum of 5-fluoro-ibogamine.

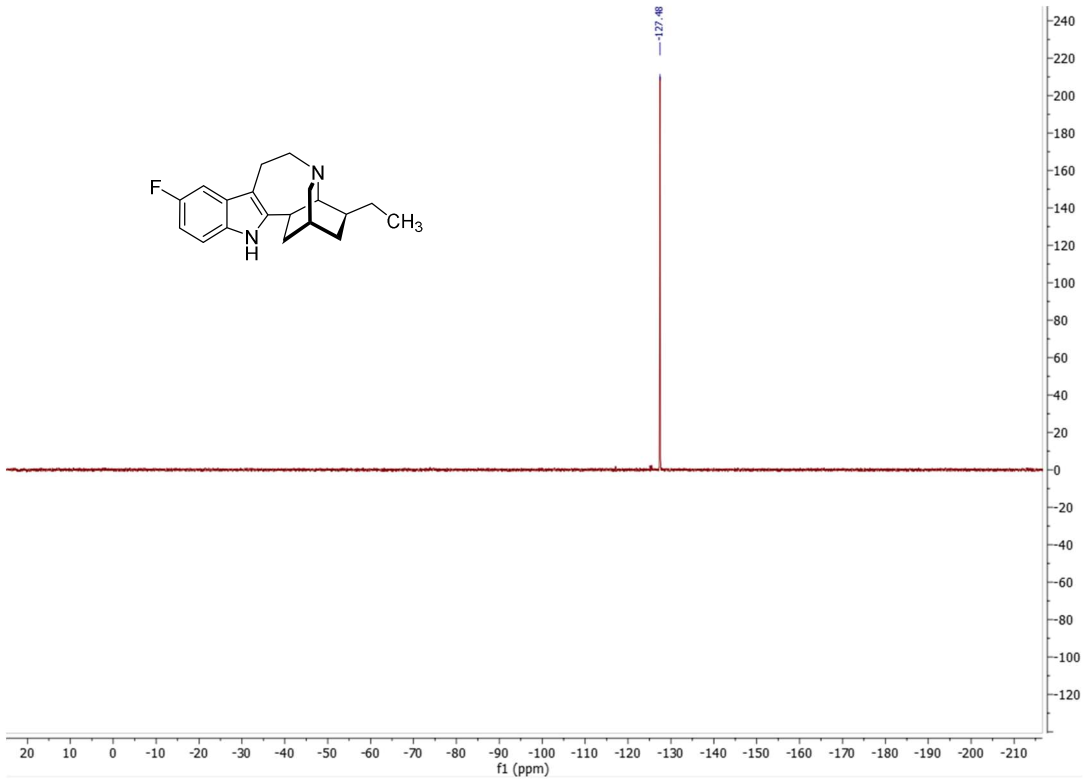


**$^{19}F$ NMR (471 MHz, MeOD)** spectrum of 5-fluoro-ibogamine.

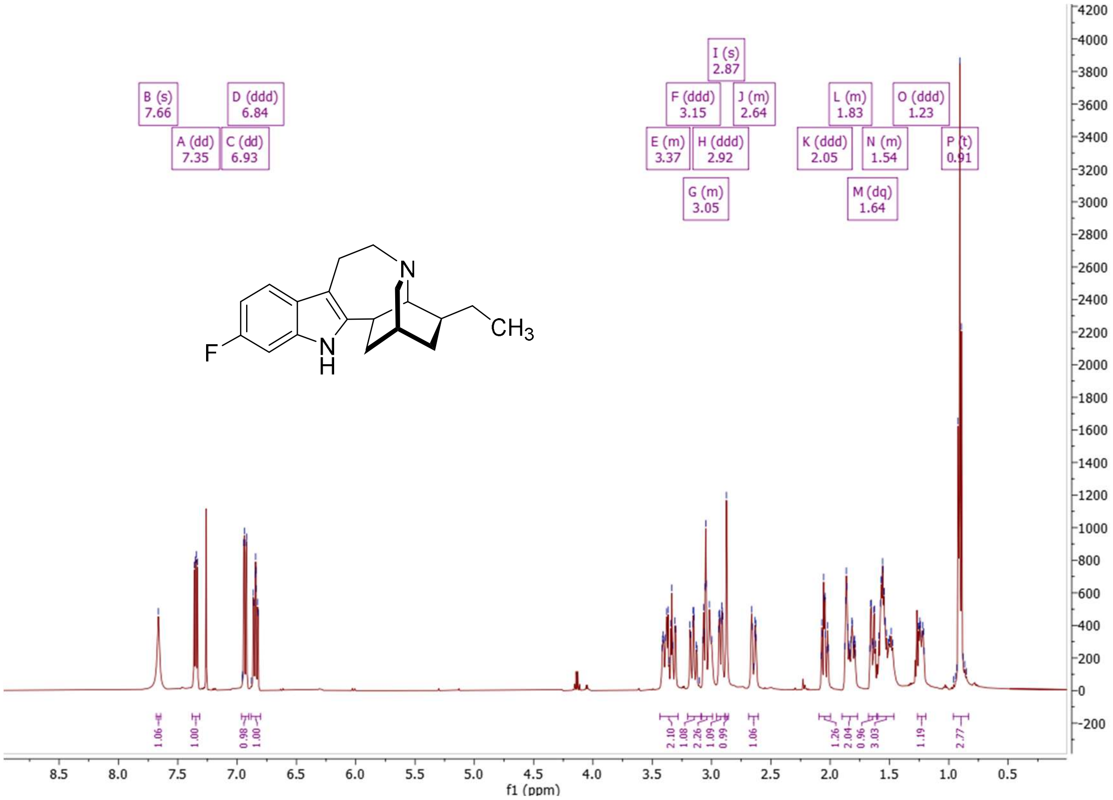


**$^1$H NMR (500 MHz, CDCl$_3$)** spectrum of 6-fluoro-ibogamine.

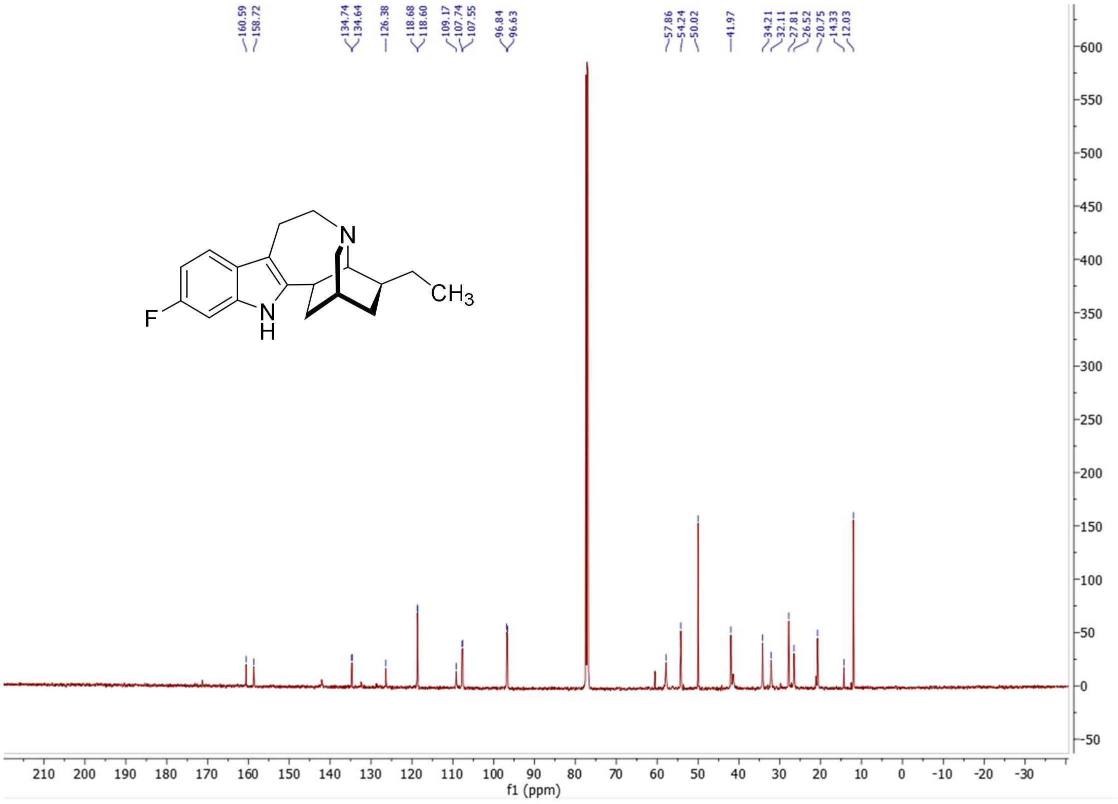


**$^{13}$C NMR (126 MHz, CDCl$_3$)** spectrum of 6-fluoro-ibogamine.

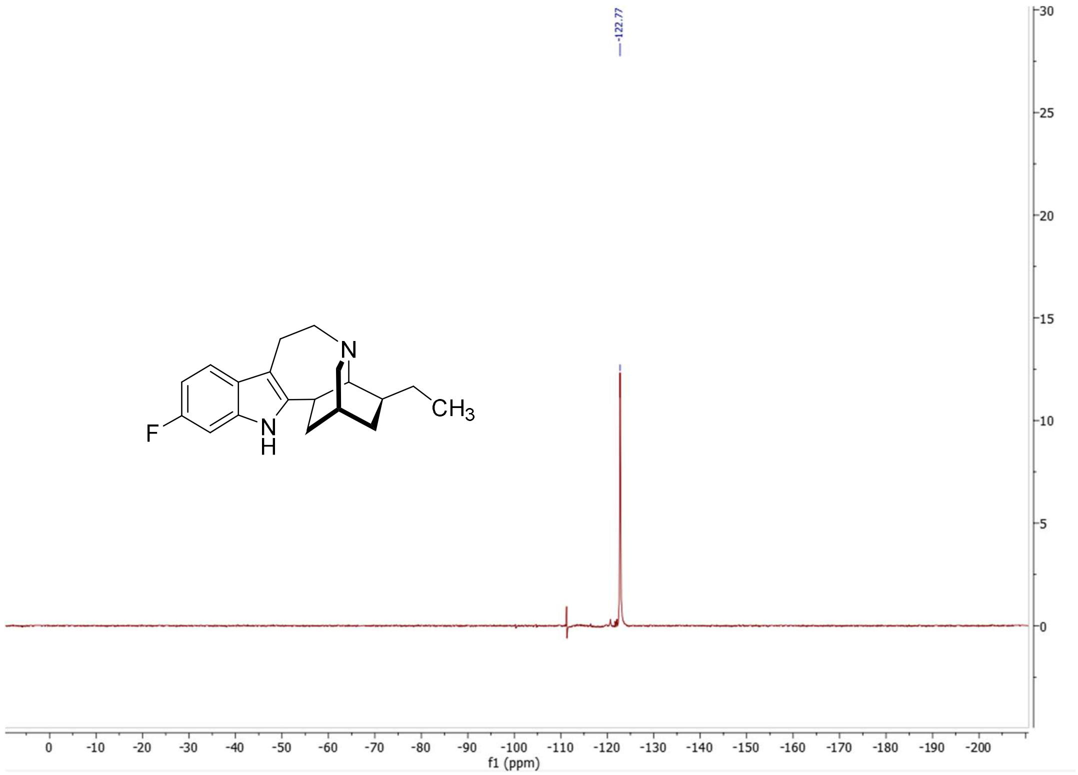


**$^{19}$F NMR (471 MHz, CDCl$_3$)** spectrum of 6-fluoro-ibogamine.

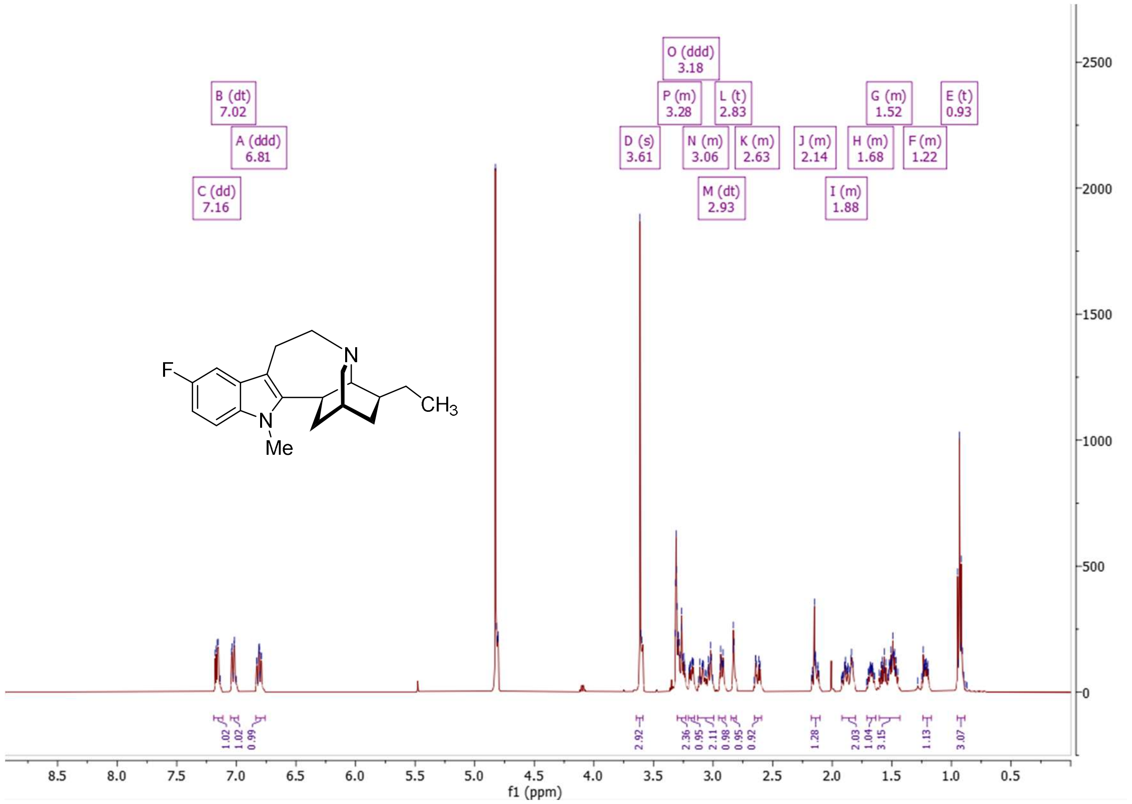


**$^{1}$H NMR (500 MHz, MeOD)** spectrum of *N*-methyl-5-fluoro-ibogamine.

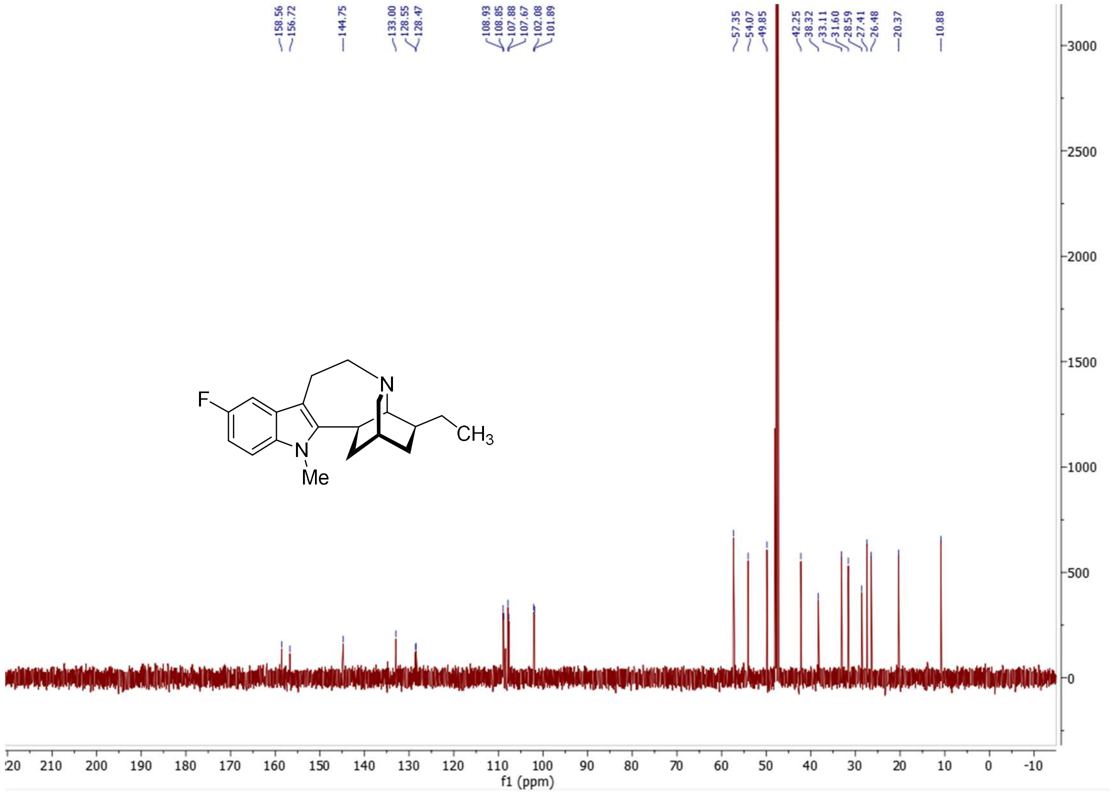


**$^{13}$C NMR (126 MHz, MeOD)** spectrum of *N*-methyl-5-fluoro-ibogamine.

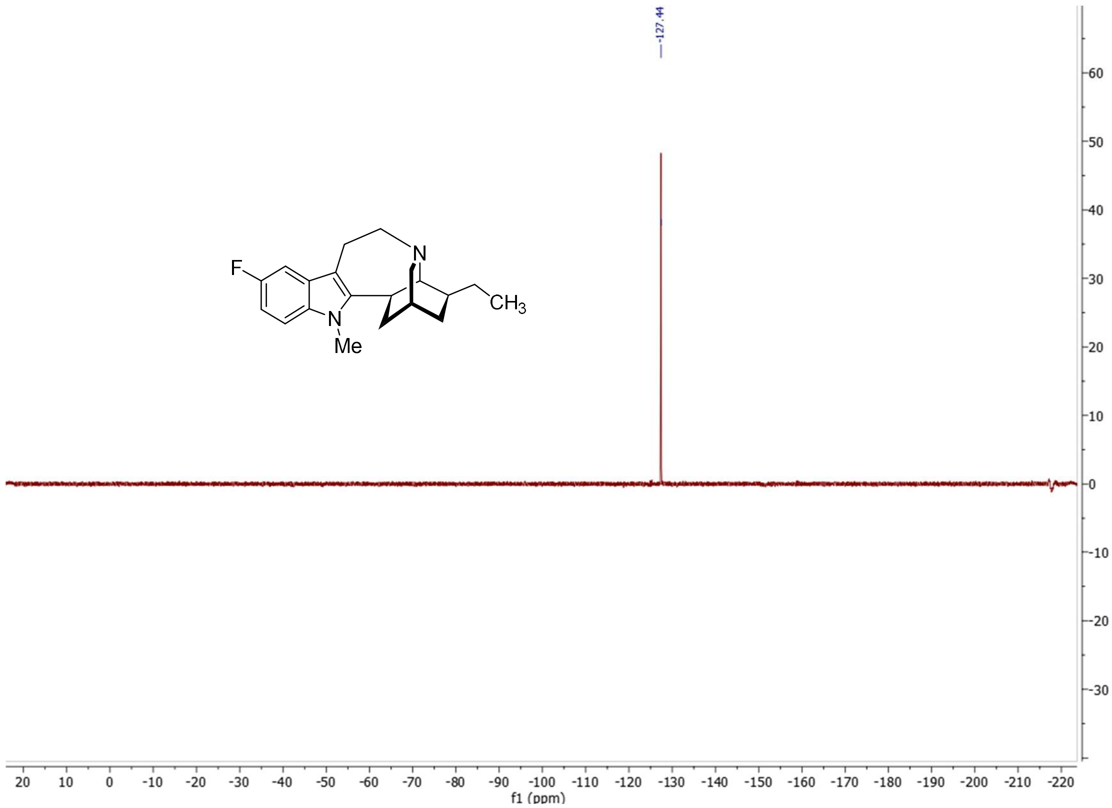


**$^{19}$F NMR (471 MHz, MeOD)** spectrum of *N*-methyl-5-fluoro-ibogamine.

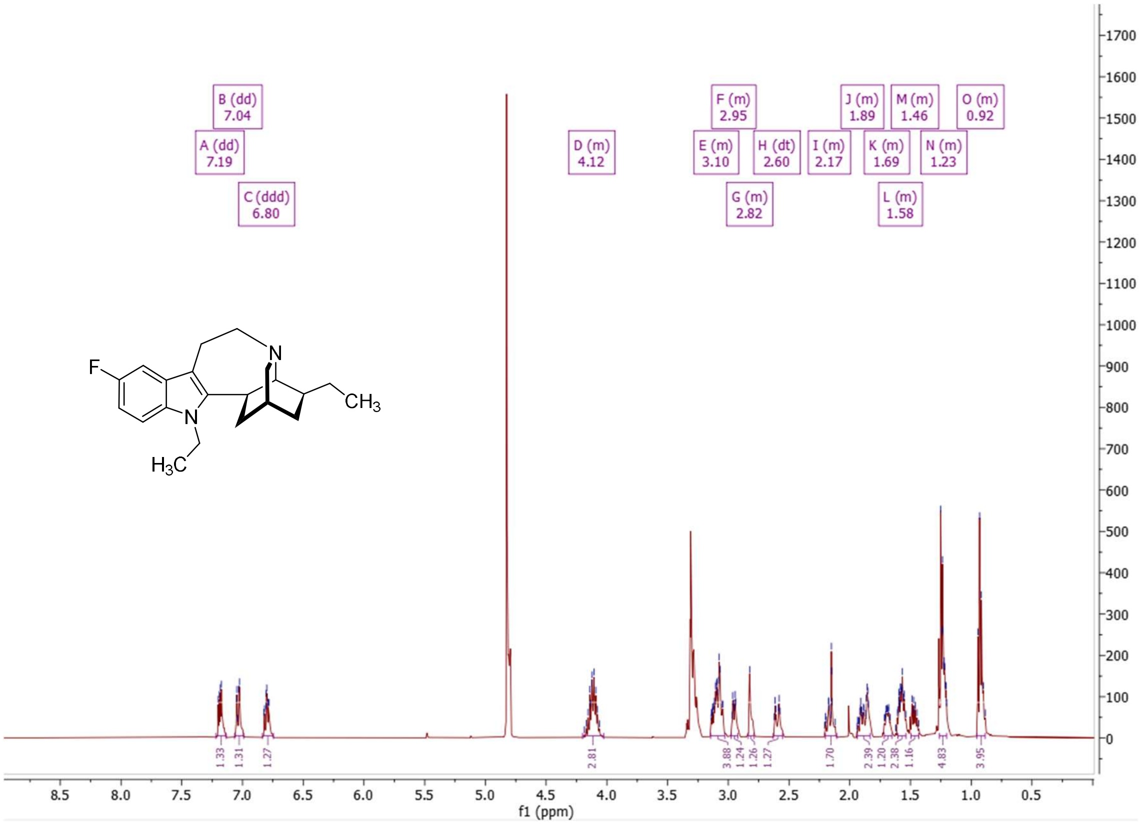


**$^{1}H$ NMR (500 MHz, MeOH)** spectrum of *N*-ethyl-5-fluoro-ibogamine.

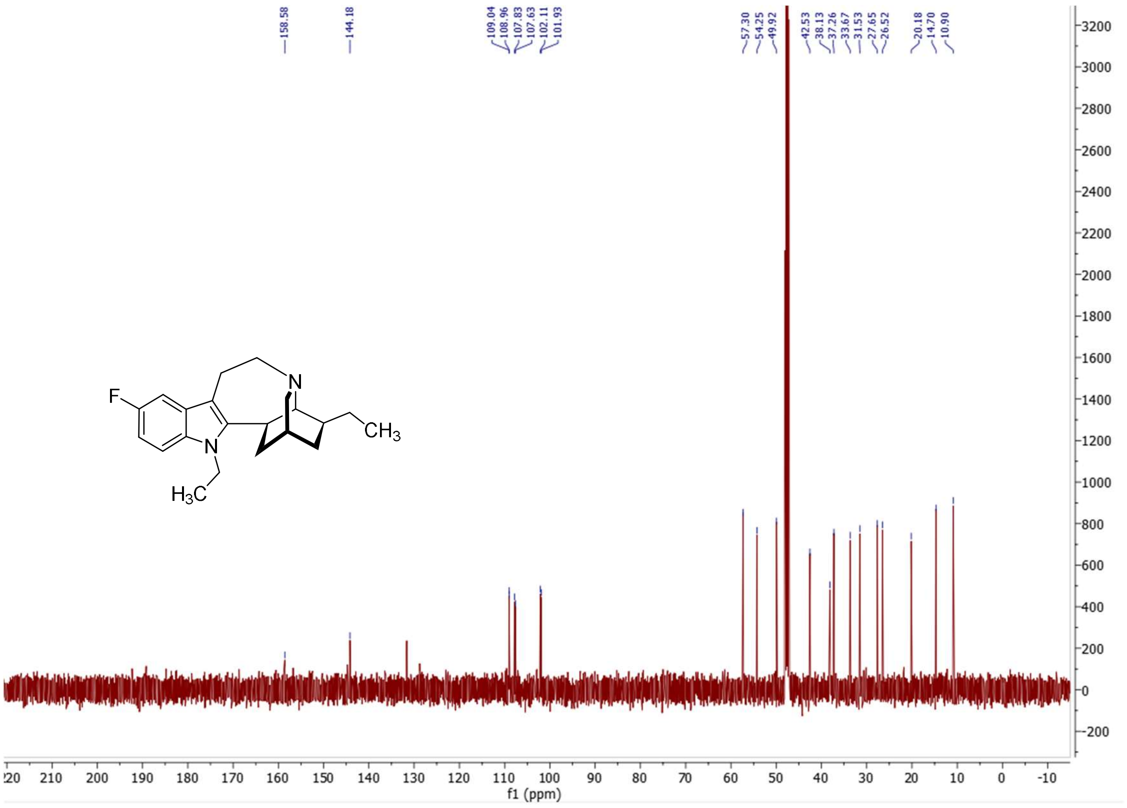


**$^{13}C$ NMR (126 MHz, MeOH)** spectrum of *N*-ethyl-5-fluoro-ibogamine.

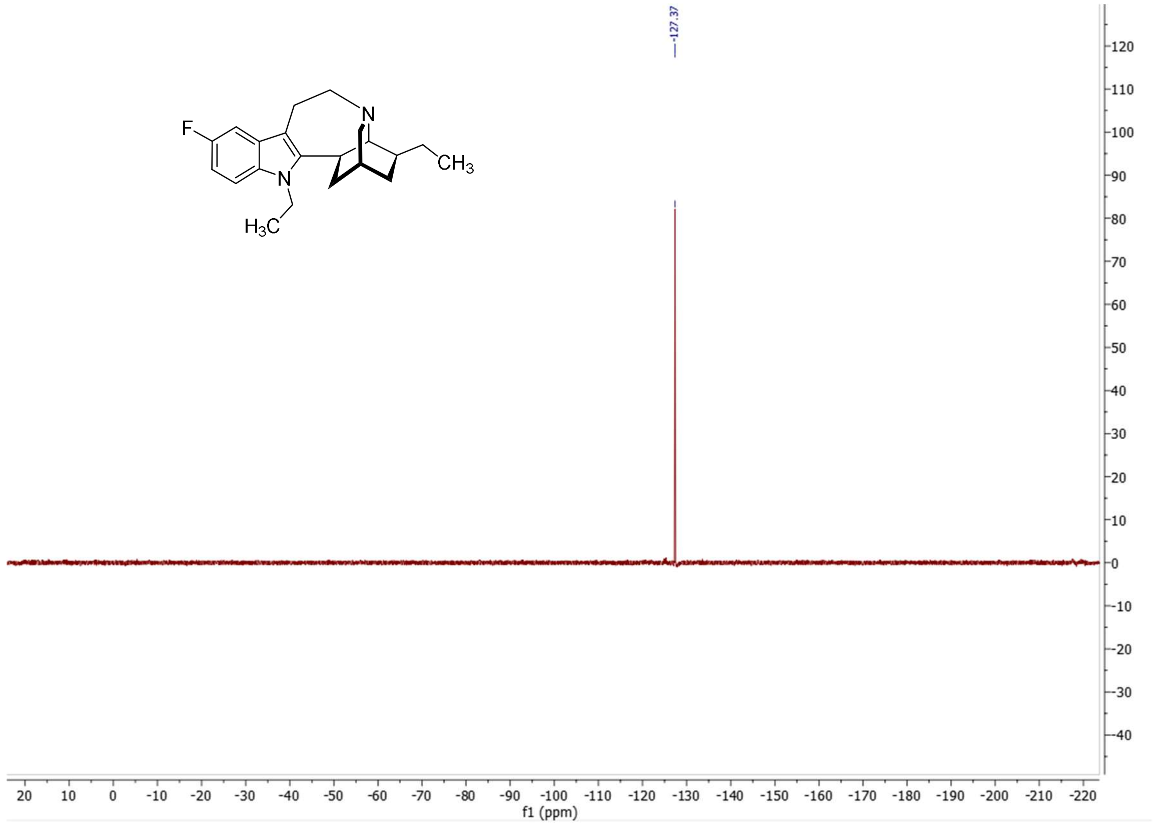


**$^{19}$F NMR (471 MHz, MeOH)** spectrum of *N*-ethyl-5-fluoro-ibogamine.

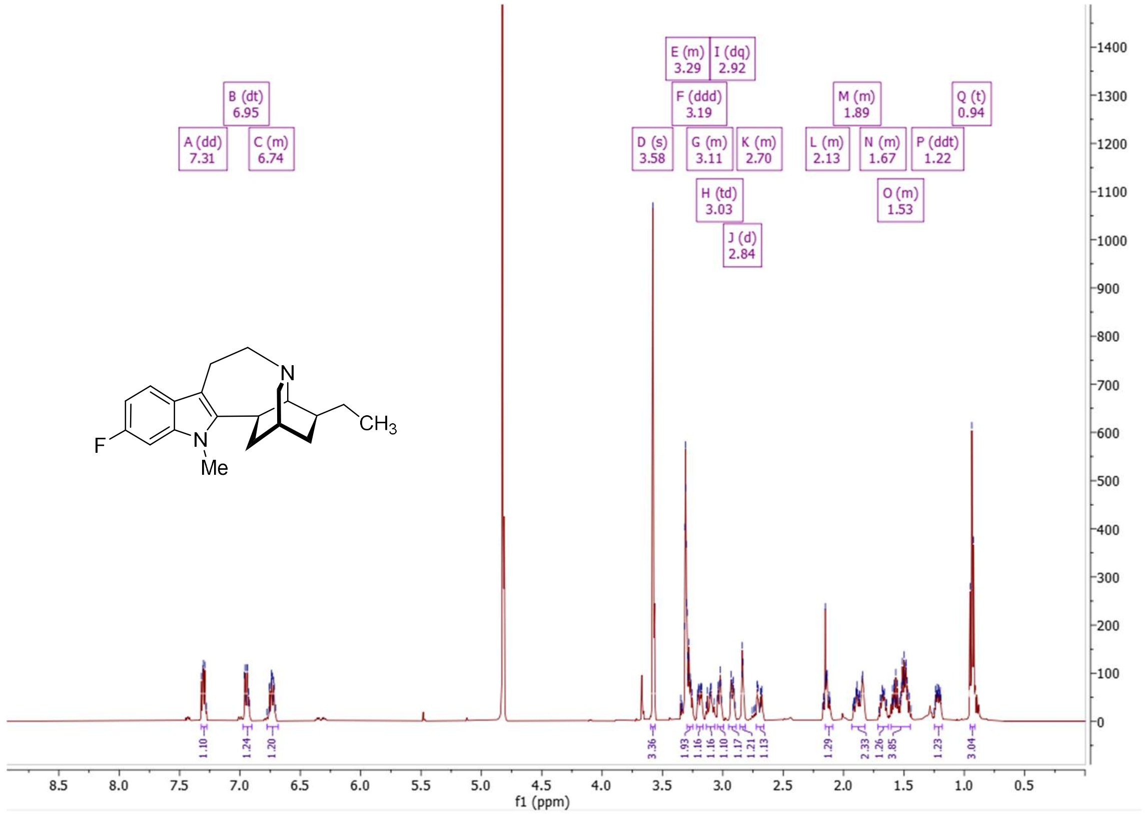


**$^{1}$H NMR (500 MHz, MeOH)** spectrum of *N*-methyl-6-fluoro-ibogamine.

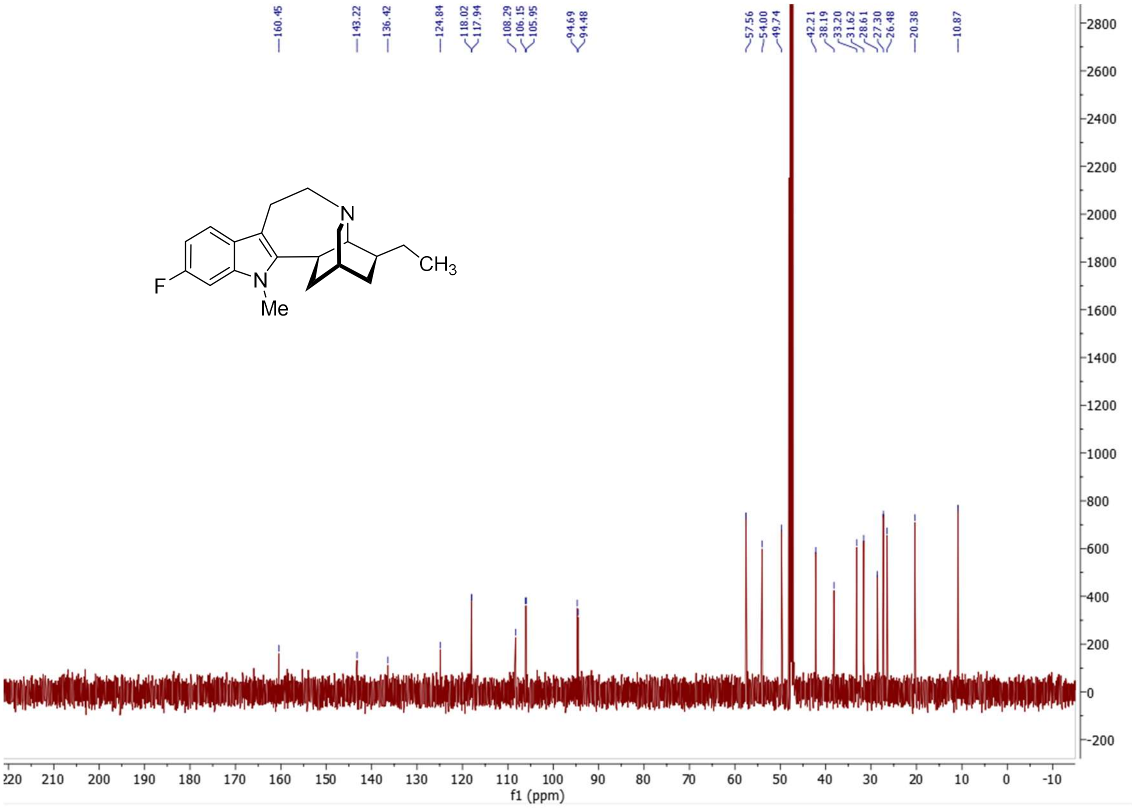


**$^{13}$C NMR (126 MHz, MeOH)** spectrum of *N*-methyl-6-fluoro-ibogamine.

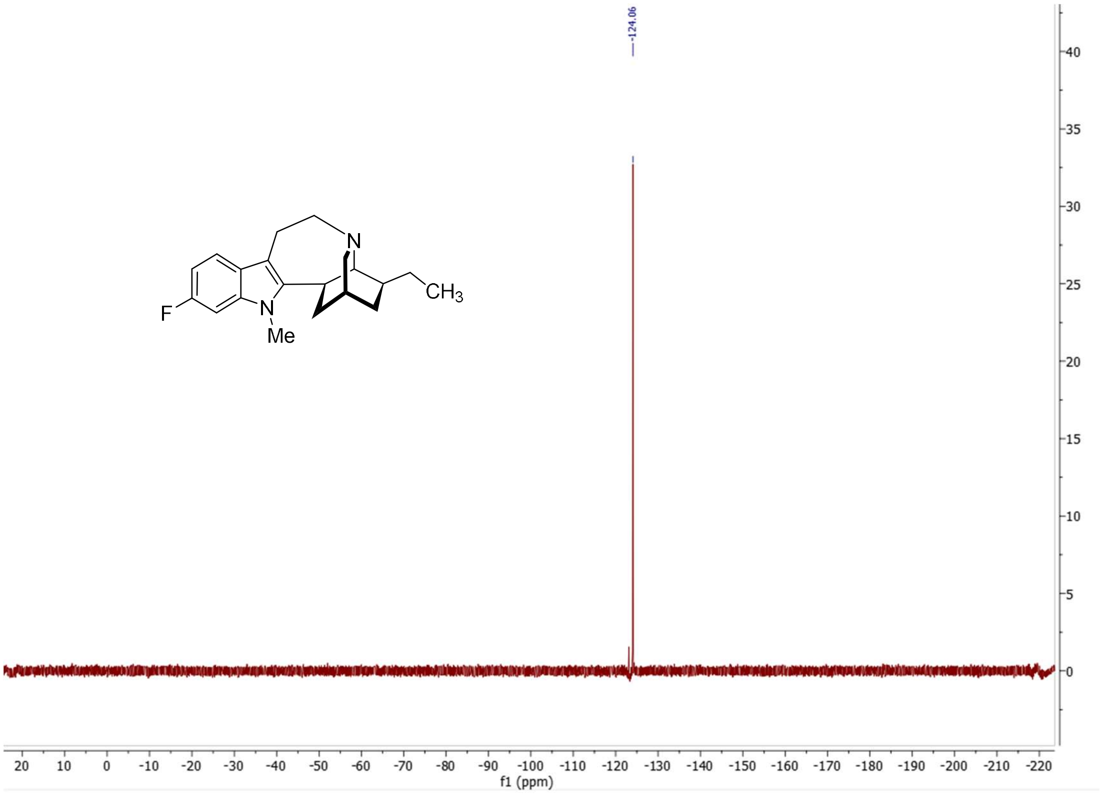


**$^{19}$F NMR (471 MHz, MeOH)** spectrum of *N*-methyl-6-fluoro-ibogamine.

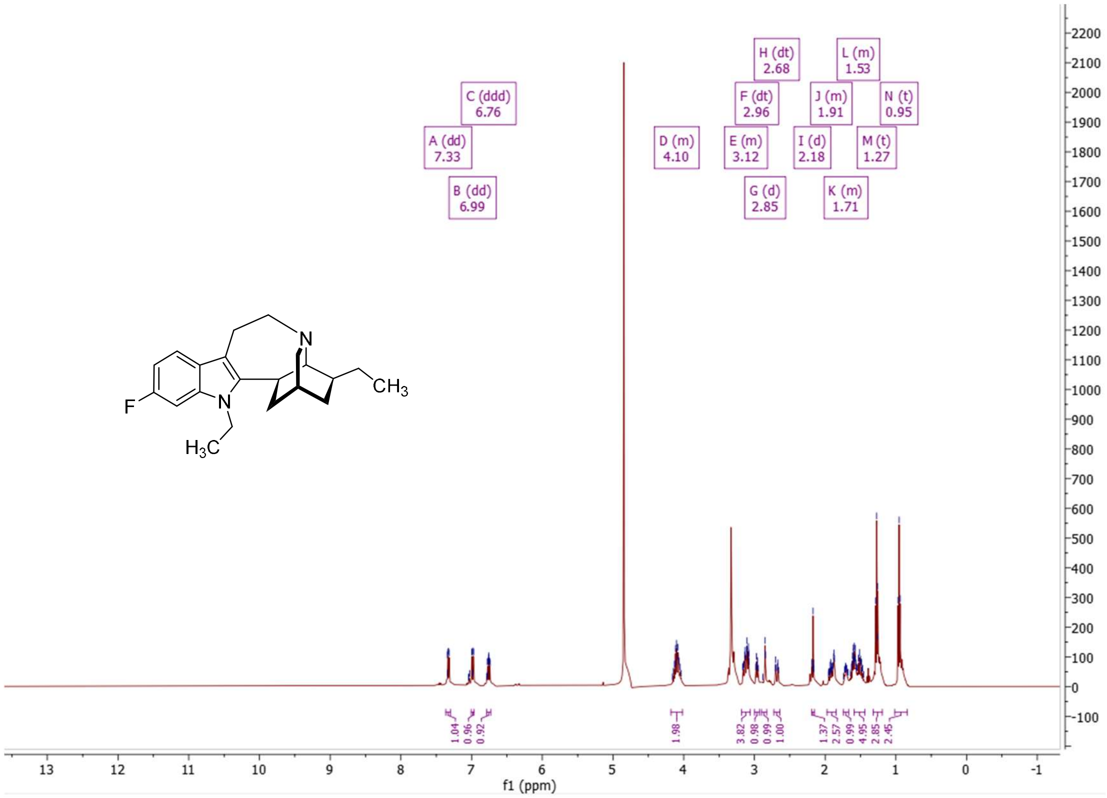


**[1]H NMR (500 MHz, MeOH)** spectrum of *N*-ethyl-6-fluoro-ibogamine.

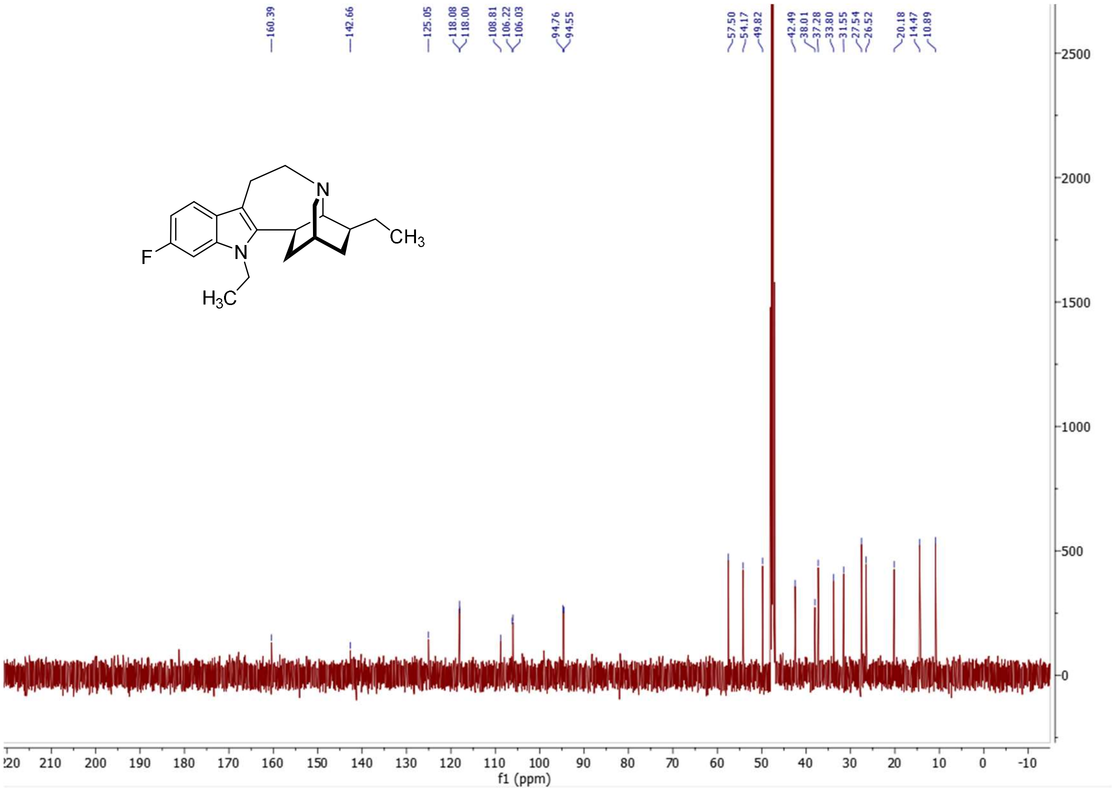


**[13]C NMR (126 MHz, MeOH)** spectrum of *N*-ethyl-6-fluoro-ibogamine.

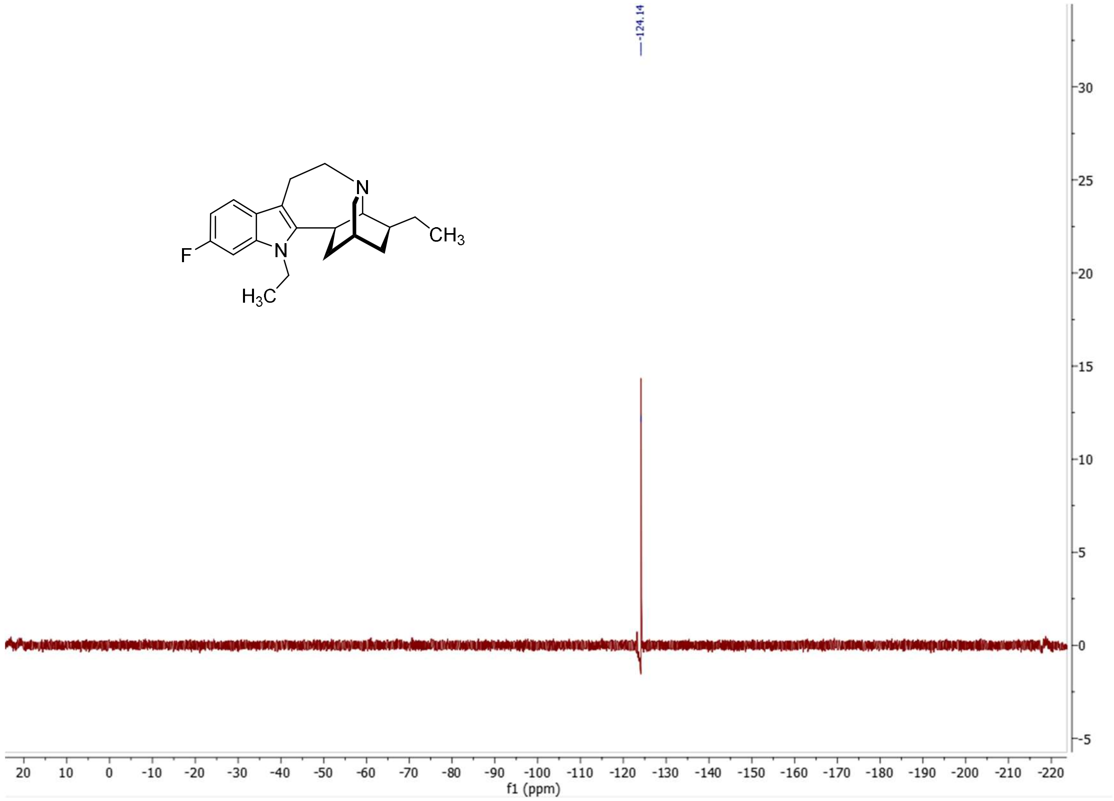


**$^{19}$F NMR (471 MHz, MeOH)** spectrum of *N*-ethyl-6-fluoro-ibogamine.

### D. Materials and Methods.

#### 1. Cell Culture Maintenance and Experimental Preparations.

Human Embryonic Kidney 293 cells stably transfected with either the human dopamine[2] (hDAT; kindly provided by Dr. Jonathan Javitch and Dr. Mark Sonders (Columbia University Irving Medical Center, Department of Psychiatry, New York, NY 10032)), norepinephrine (hNET), serotonin transporter (hSERT), or vesicular monoamine transporter 2[16] (hVMAT2; gifted by Dr. Gary W. Miller and Mr. Joshua M. Bradner (Columbia University Mailman School of Public Health, Department of Environmental Health Sciences, New York, NY 10032)) were maintained in Dulbecco's Minimal Essential Medium (DMEM) with GlutaMAX (Gibco-Thermo Fisher Scientific; Waltham, MA) supplemented with 10% (v/v) fetal bovine serum (FBS; Atlanta Biologicals; Flowery Branch, GA), 100 U/mL penicillin (Invitrogen; Waltham, MA), and 10 μg/mL streptomycin (Invitrogen; Waltham, MA). For hNET and hSERT cultures, 500 μg/mL G418 (Gibco-Thermo Fisher Scientific; Waltham, MA) was added to maintain the transporter transgene. Alternatively, for hVMAT2 cultures, 100 μg/mL zeocin (InvivoGen; San Diego, CA) was supplemented.[16] All cell cultures were plated on 10 cm polystyrene culture plates (Corning Falcon; Corning, NY) and grown in a humidified environment of 37 degrees Celsius and five percent carbon dioxide. Subculturing of fully confluent cultures occurred every three to four days. Growth medium was aspirated and replaced by experimental medium for conducting assays. For hDAT, hNET, and hSERT HEK cell cultures, the utilized experimental medium consisted of DMEM minus phenol red (Gibco-Thermo Fisher Scientific; Waltham, MA), 1% (v/v) FBS, 100 U/mL penicillin, and 10 μg/mL streptomycin. For hVMAT2 cells, 1% (v/v) L-glutamine (Gibco-Thermo Fisher Scientific; Waltham, MA) was additionally included in the above solution.

Furthermore, Flp-In T-REx Human Embryonic Kidney 293 cells (Thermo Fisher Scientific; Waltham, MA) singly transfected with either tetracycline-inducible organic cation transporter 1 (hOCT1), 2 (hOCT2), 3 (hOCT3), or plasma membrane monoamine transporter (hPMAT) were maintained in a solution consisting of Dulbecco's Minimal Essential Medium (DMEM) with GlutaMAX supplemented with 10% (v/v) fetal bovine serum, 100 U/mL penicillin, and 10 μg/mL streptomycin and grown in a humidified environment of 37 degrees Celsius and five percent carbon dioxide. Subculturing was conducted after every three to four days when the cell cultures reached approximately 85 percent confluency in 10 cm polystyrene culture plates. After every two successive passages, 200 μg/mL Hygromycin B (Corning; Corning, NY) and 15 μg/mL Blasticidin (Sigma-Aldrich; St. Louis, MO) were added for 24 hours to maintain the transgene. On experimental assay days, growth medium was aspirated, and cells were bathed in experimental medium with 1× Hanks' Balanced Salt Solution (HBSS; Gibco-Thermo Fisher Scientific; Waltham, MA), 20 mM HEPES (Gibco-Thermo Fisher Scientific; Waltham, MA), 0.1% (m/v) Bovine Serum Albumin (BSA; Sigma-Aldrich; Milwaukee, WI), and 0.01% (m/v) ascorbic acid (Sigma-Aldrich ; St. Louis, MO). These cell lines were a gift provided by Dr. Daniel Wacker and Ms. Audrey Warren (Icahn School of Medicine at Mount Sinai, New York, NY 10029).

#### 2. Determination of *$IC_{50}$* Metrics of Ibogaine Derivatives at hDAT, hNET, and hSERT Through APP+ Uptake Blockage.

Stably transfected cell cultures were seeded in white solid-bottom 96-well plates (Corning; Corning, NY) at a density of $1.00 \times 10^6$ cells/well and allowed to proliferate in a humidified environment of 37 degrees Celsius and five percent carbon dioxide to full confluency in approximately 48 hours. Upon commencement of the experiment, growth medium was aspirated, and the cell culture monolayer was rinsed twice with 120 µL of 1× PBS. Experimental medium solutions consisting of tiered concentrations (ranging from 100 µM to 0.1 µM) of either an ibogaine derivative or DMSO (vehicle, 0.02% v/v, Sigma-Aldrich; St. Louis, MO) were added gently to the cell cultures and consequently, pre-incubated for approximately one hour. A standard was utilized for each experiment depending on the specific transporter being evaluated (indatraline for hDAT[5], reboxetine for hNET[6], and imipramine for hSERT[1]; Sigma-Aldrich; St. Louis, MO). Subsequently, an equivalent amount of experimental medium solutions containing both tiered concentrations (ranging from 100 µM to 0.1 µM) of either an ibogaine derivative or DMSO (vehicle, 0.02% v/v, Sigma-Aldrich; St. Louis, MO) and 2× 4-(4-dimethylamino)phenyl-1-methylpyridinium (APP+; final concentration: 1.1 µM; Sigma-Aldrich; St. Louis, MO)[2] was added to each well and then incubated for an additional 30 minutes to encourage fluorescent probe uptake. Afterwards, the solution contained within each well was aspirated and cells were rinsed with two successive washes with 120 µL 1× PBS washes. A final addition of 120 µL of 1× PBS in each well was necessary for fluorescent uptake bottom mode readout by BioTek Synergy Neo2 Hybrid Multi-Mode Reader (Agilent; Santa Clara, CA), using 3 × 3 area scan and bottom-read mode, at the excitation and emission wavelengths of 436 and 500 nm, respectively. For *$IC_{50}$* data analysis, the average fluorescence of vehicle wells was subtracted by that of wells containing each ibogaine derivative to quantify the respective fluorescence uptake (in mean fluorescence units). These numerics were normalized to the maximum and minimum fluorescence uptake difference values correlated to the utilized standard inhibitor and then fit to a nonlinear curve model ([inhibitor] versus response (three parameters)) as provided by Graphpad Prism 8 software (Graphpad Prism Inc.; San Diego, CA). Outputted *$IC_{50}$* (± SEM) quantities for each ibogaine derivative can be converted into *$K_i$* (± SEM) values using the Cheng-Prusoff equation.[17]

#### 3. Determination of *$IC_{50}$* Metrics of Ibogaine Derivatives at hVMAT2 Through FFN206 Uptake Blockage.

A similar procedure was employed in the section titled "Determination of $IC_{50}$ Metrics of Ibogaine Derivatives at hDAT, hNET, and hSERT Through APP+ Blockage". However, there are a few differences in the protocol. The fluorescent substrate utilized was FFN206[18] (final concentration: 0.75 µM) and the standard inhibitor used was reserpine[19] (Sigma-Aldrich; St. Louis, MO). Furthermore, the pre-incubation period lasted 30 minutes and the probe incubation period was longer (60 minutes). Fluorescence uptake measurements (at the excitation and emission wavelengths of 370 nm and 464 nm, respectively) and *$IC_{50}$* metric determinations were conducted exactly as detailed in the previous section.

#### 4. Determination of $IC_{50}$ Metrics of Ibogaine Derivatives at hOCT1 and hPMAT Through APP+ Uptake Blockage.

A similar procedure was employed in the section titled "Determination of $IC_{50}$ Metrics of Ibogaine Derivatives at hDAT, hNET, and hSERT Through APP+ Blockage". However, there are notable alterations in the protocol. Such stably transfected cell cultures were seeded in white solid-bottom 96-well plates (Corning; Corning, NY) at a density of 0.40-0.60 $\times$ $10^6$ cells/well in growth medium containing 2 μg/mL tetracycline (Sigma-Aldrich; St. Louis, MO) and permitted to grow in a humidified environment of 37 degrees Celsius and five percent carbon dioxide for approximately 48 hours to both reach full confluency and induce transporter expression. The final concentration of APP+ utilized is 12.5 μM. The standard inhibitor used was decynium-22 (Sigma-Aldrich; St. Louis, MO).[7,9] Furthermore, the pre-incubation period once again lasted 30 minutes and the probe incubation period was 60 minutes. Fluorescence uptake measurements (at the excitation and emission wavelengths of 436 nm and 500 nm, respectively) and $IC_{50}$ metric determinations were conducted exactly as detailed in the mentioned section.

#### 5. Determination of $IC_{50}$ Metrics of Ibogaine Derivatives at hOCT2 and hOCT3 Through ASP+ Uptake Blockage.

A similar procedure was employed in the section titled "Determination of $IC_{50}$ Metrics of Ibogaine Derivatives at hDAT, hNET, and hSERT Through APP+ Blockage". However, there are notable alterations in the protocol. Such stably transfected cell cultures were seeded in white solid-bottom 96-well plates (Corning; Corning, NY) at a density of 0.50-0.60 $\times$ $10^6$ cells/well in growth medium containing 2 μg/mL tetracycline (Sigma-Aldrich; St. Louis, MO) and permitted to proliferate in a humidified environment of 37 degrees Celsius and five percent carbon dioxide for approximately 48 hours to both reach full confluency and induce transporter expression. The final concentration of 4-[4-(dimethylamino)styryl]-*N*-methylpyridium iodide (ASP+; Sigma-Aldrich; St. Louis, MO) utilized is 5 μM.[7] The standard inhibitor used was decynium-22 (Sigma-Aldrich; St. Louis, MO).[7,9] The pre-incubation period lasted 30 minutes while the probe incubation period was 30 minutes for experiments using hOCT2 and 60 minutes for experiments using hOCT3. Fluorescence uptake measurements (at the excitation and emission wavelengths of 447 nm and 614 nm, respectively) and $IC_{50}$ metric determinations were conducted exactly as detailed in the mentioned section.

#### 6. Determination of $IC_{50}$ Metrics of Ibogaine Derivatives at hDAT, hNET, and hSERT Through Tritiated Monoamine Uptake Blockage Using Rat Brain Synaptosomes.

Procurement of rat brain synaptosomes, conduction of the tritiated monoamine uptake inhibition assay, and the quantification of $IC_{50}$ metrics of ibogaine and selected derivatives followed this previously published procedure.[20]

#### 7. Determination of $EC_{50}$ Metrics of Ibogaine Derivatives at hDAT, hNET, and hSERT Through Tritiated 5-HT Release By SERT Using Rat Brain Synaptosomes.

Procurement of rat brain synaptosomes, conduction of the tritiated 5-HT release assay at SERT, and the quantification of $EC_{50}$ metrics of ibogaine and selected derivatives followed this previously published procedure.[20]

#### 8. One-Photon Epifluorescence Microscopy Imaging of Respective Uptake in hSERT-HEK and hVMAT2-HEK Cell Cultures.

hSERT or hVMAT2 cell cultures were plated onto previously poly-D-lysine (Alamanda Polymers, Huntsville, AL) coated (one hour) clear bottom μ-Slide 8-well $^{high}$ibiTreat chambered coverslips (ibidi; Gräfelfing, Germany) at a density of $0.06 \times 10^6$ cells per well. The cultures proliferated at 37 degrees Celsius and in a humidified five percent carbon dioxide environment. At the start of this imaging experiment, cell culture medium was aspirated, and the cells were rinsed twice with 200 μL of 1× PBS. 180 μL experimental medium containing either a standard inhibitor (imipramine for hSERT[1] and tetrabenazine for hVMAT2[21]; final concentration: 2 μM; Sigma-Aldrich; St. Louis, MO), ibogaine derivative (final concentration: 2 μM), or DMSO (vehicle, 0.02% v/v) was added to each well before a resulting one hour incubation period. Subsequently, a 20 μL solution consisting of either standard inhibitor (final concentration: 2 μM), ibogaine derivative (final concentration: 2 μM), or DMSO (vehicle, 0.02% v/v) and a fluorescent substrate (APP+ for hSERT[2], final concentration: 2 μM; FFN206 for hVMAT2[18], final concentration: 20 μM) was added and then incubated for a final time period (30 minutes for hSERT and 60 minutes for hVMAT2) for respective probe uptake. Cellular solutions were then aspirated, and cells were consequently washed twice with 200 μL of 1× PBS and maintained in 200 μL of cell-type specific experimental media for epifluorescence imaging. Fluorescent images (three images per well in triplicate wells) were taken using a Leica DMI4000 epifluorescence microscope (Leica Microsystems; Wetzlar, Germany) equipped with a xenon lamp (Sutter Lambda LS; Sutter Instrument; Novato, CA), Leica objective (40×/0.45 HI Plan; Leica Microsystems; Wetzlar, Germany), a PCO.Panda 4.2 camera (PCO; Kelheim, Germany), and a Chroma custom filter cube (For APP+ uptake imaging: $\lambda_{excitation}$ = 470/40 nm, $\lambda_{emission}$ = 525/50 nm; For FFN206 uptake imaging: $\lambda_{excitation}$ = 350/25 nm, $\lambda_{emission}$ = 460/25 nm; Chroma Technology Corporation; Bellows Falls, VT). Micro-Manager software[22] was utilized to acquire all fluorescence images, which were then normalized, using ImageJ software (National Institutes of Health; Bethesda, MD) to the same contrast and brightness level corresponding to the quantitative fluorescence uptake observed without the inclusion of a standard inhibitor or ibogaine derivative.

#### 9. One-Photon Confocal Microscopy Imaging of Fluorescence Uptake in hSERT-HEK and hVMAT2-HEK Cells.

hSERT-HEK or hVMAT2-HEK cells were plated onto poly-D-lysine (Alamanda Polymers; Huntsville, AL) coated clear bottom μ-Slide 8-well $^{high}$ibiTreat chambered coverslips (ibidi; Gräfelfing, Germany) at a density of $2 \times 10^4$ cells per well for 24 hours in a 37 degrees Celsius and five percent carbon dioxide environment. After 24 hours, cell culture medium was aspirated, and cells were washed once with 200 μL of 1× PBS before addition of 200 μL 1× PBS consisting of either inhibitor (2 μM imipramine for hSERT[1] and 2 μM reserpine for hVMAT2[19]), ibogaine compound (2 μM ibogaine, 2 μM noribogaine, 2 μM 5-cyano-ibogamine, or 2 μM ibogamine) or vehicle (DMSO, 0.02% v/v) and subsequent incubation (60 minutes for hSERT or 30 minutes for hVMAT2). Afterwards, 10 μM SERTlight[23] (for hSERT) or 20 μM FFN206[18] (for hVMAT2) were added, and the cells were incubated (30 minutes for hSERT and 120 minutes for hVMAT2) to permit probe uptake. The media was removed, and the cells washed twice with 200 μL of 1× PBS. Cells were maintained in 200 μL 1× PBS for imaging. Differential Interference Contrast (DIC) and fluorescence images (three images per well in duplicate wells) were acquired using a Zeiss LSM800 confocal microscope (Zeiss; Jena, Germany) equipped with a 63×/1.4 numerical aperture (NA) objective lens with Immersol™ 518 F oil (Zeiss; Oberkochen, Germany). For two-dimensional imaging, images were 1024 × 1024 pixels with a bit depth of 16 and scan speed of 5. For three-dimensional imaging, 50 slices at 0.37 μm intervals were obtained at 1024 × 1024 pixels with a bit depth of 16 and a scan speed of 7. Zeiss ZEN software was used to operate the microscope. Two-dimensional images were uniformly contrasted, and three-dimensional renderings were produced using the 3D Projection function in ImageJ software (National Institutes of Health; Bethesda, MD). Zeiss LSM800 confocal microscope and respective analytical resources were kindly provided by the Department of Biological Sciences, Columbia University, New York, NY 10027.

#### 10. Determination of SERT Inhibition by Ibogaine Derivatives in Acute Murine Brain Slices.

**Animal Protocols.** All animal protocols followed NIH guidelines and were approved by Columbia University's Institutional Animal Care and Use Committee (IACUC). Mice were housed in groups of five or less per cage, in a 12-hour day/12-hour night cycle with *ad libitum* access to food and water. Wildtype C57BL/6 mice were obtained from The Jackson Laboratory (Bar Harbor, ME).

**Acute Murine Brain Slice Preparation.** Animals were sacrificed at the age of 9–18 weeks. All animal protocols were approved by the IACUC of Columbia University. Mice were decapitated and acute 300 μm thick coronal slices were cut on a Leica VT1200 vibratome (Leica Microsystems; Wetzlar, Germany) at four degrees Celsius and then allowed to recover for 30 minutes in oxygenated (95% $O_2$, 5% $CO_2$) artificial cerebrospinal fluid (aCSF) containing (in mM): 125 NaCl, 2.5 KCl, 26 $NaHCO_3$, 0.3 $KH_2PO_4$, 2.4 $CaCl_2$, 1.3 $MgSO_4$, 0.8 $NaH_2PO_4$, 10 Glucose (pH = 7.2–7.4, 292–296 mOsm/L). Slices were then used at room temperature for all imaging experiments.

**Application and Two-Photon Imaging of SERT Inhibition by Ibogaine Derivatives.** For inhibition experiments, slices were first preincubated with 2 μM citalopram (Sigma-Aldrich; St. Louis, MO) or ibogaine derivative compound at the desired concentration for 30 minutes, and then co-incubated with

10 μM SERTlight[23] for a 30-minute loading period. Slices were then transferred to an imaging chamber (QE-1; Warner Instruments; Hamden, CT) and held in place by a platinum wire and nylon string custom made holder and superfused (2 mL/min) with oxygenated aCSF. Slices were washed in the perfusion chamber for 10 min before imaging. Fluorescent structures were visualized at depths of at least 30 μm from the slice surface using a Prairie Ultima Multiphoton Microscopy System (Prairie Technologies; Middleton, WI) with a titanium-sapphire Chameleon Ultra II laser (Coherent; Santa Clara, CA) equipped with a 60 × 0.9 NA water immersion objective. SERTlight was excited at 710 nm and 435–485 nm light was collected. Images were captured in 12-bit 90 × 90 μm field of view at 512 × 512-pixel resolution and a dwell time of 3 μs/pixel using Prairie View software (Prairie Technologies; Middleton, WI).

#### 11. Determination of VMAT2 Inhibition by Ibogaine Derivatives in Acute Murine Brain Slices.

**Animal Protocols.** All animal protocols followed NIH guidelines and were approved by Columbia University's Institutional Animal Care and Use Committee (IACUC). Mice were housed in groups of five or less per cage, in a 12 h day/12 h night cycle with *ad libitum* access to food and water. Wildtype C57BL/6 mice were obtained from The Jackson Laboratory (Bar Harbor, ME).

**Acute Murine Brain Slice Preparation.** Animals were sacrificed at the age of 9–18 weeks. All animal protocols were approved by the IACUC of Columbia University. Mice were decapitated and acute 300 μm thick coronal slices were cut on a Leica VT1200 vibratome (Leica Microsystems; Wetzlar, Germany) at four degrees Celsius and then allowed to recover for 30 minutes in oxygenated (95% $O_2$, 5% $CO_2$) artificial cerebrospinal fluid (aCSF) containing (in mM): 125 NaCl, 2.5 KCl, 26 $NaHCO_3$, 0.3 $KH_2PO_4$, 2.4 $CaCl_2$, 1.3 $MgSO_4$, 0.8 $NaH_2PO_4$, 10 Glucose (pH = 7.2–7.4, 292–296 mOsm/L). Slices were then used at room temperature for all imaging experiments.

**Application and Two-Photon Imaging of VMAT2 Inhibition by Ibogaine Derivatives.** For inhibition experiments, slices were first preincubated with 2 μM dihydrotetrabenazine (dhTBZ; Cayman Chemical; Ann Arbor, MI) or ibogaine derivative at the desired concentration for 30 minutes, and then co-incubated with 10 μM FFN200[24] for a 30-minute loading period. Slices were then transferred to an imaging chamber (QE-1; Warner Instruments; Hamden, CT) and held in place by a platinum wire and nylon string custom made holder and superfused (2 mL/min) with oxygenated aCSF. Slices were washed in the perfusion chamber for 10 min before imaging. Fluorescent structures were visualized at depths of at least 30 μm from the slice surface using a Prairie Ultima Multiphoton Microscopy System (Prairie Technologies; Middleton, WI) with a titanium-sapphire Chameleon Ultra II laser (Coherent; Santa Clara, CA) equipped with a 60 × 0.9 NA water immersion objective. FFFN200 was excited at 740 nm and 435–485 nm light was collected. Images were captured in 12-bit 90 × 90 μm field of view at 512 × 512-pixel resolution and a dwell time of 3 μs/pixel using Prairie View software (Prairie Technologies; Middleton, WI).

**Quantification of Brain Slice FFN200 Puncta Number.** FFN200[24] images used for the calculation of respective puncta density were analyzed using Fiji. Fluorescent puncta were identified by defining a threshold of intensity as well as size constraints. Each thresholded image was processed into a binary mask and the watershed function was applied to delineate distinct puncta in close proximity. The "3D

Objects Counter" plugin was applied for final puncta counting, and all results were manually verified to be within one percent of the calculated results. Images taken before and after treatment were analyzed using identical analysis parameters. The percentage of puncta remaining after treatment was calculated by dividing the number of puncta identified after treatment by the number of puncta identified before treatment and multiplying by 100.

#### 12. Determination of Pharmacokinetic Parameters in Mice (Bienta Enamine Biological Services).

**Study Design.** Study design, animal selection, handling and treatment were all in accordance with the Enamine pharmacokinetics (PK) study protocols and Institutional Animal Care and Use Guidelines (BACUC #CU-PK-06012025). Animal treatment and samples preparation were conducted by the Animal Laboratory personnel at Enamine/Bienta. Male C57BL/6J mice (bred in Bienta's Animal Research Centre) 11 weeks old, body weight ranged from 18.9 g to 24.8 g and average body weight across all groups 21.7 g, standard deviation (SD) = 1.4 g were used in this study. The animals were randomly assigned to the treatment groups before the pharmacokinetic study; all animals were fasted for four hours before dosing. Subcutaneous (SC) route of administration; six sampling time points (5, 15, 30, 60, 180, and 360 minutes) for 10 mg/kg dosing group and seven sampling time points (5, 15, 30, 60, 180, 360, and 600 minutes) for 50 mg/kg dosing group were set for this pharmacokinetic study. Each of the time point treatment groups included three animals. There was also a control group of one animal. Mice were injected IP with 2,2,2-tribromoethanol at the dose of 150 mg/kg prior to drawing the blood. Blood collection was performed from the orbital sinus in microtainers containing $K_3$EDTA. Animals were sacrificed by cervical dislocation after the blood samples collection. Blood samples were centrifuged for 10 minutes at 3000 rpm (+4ºC). Brain samples (left hemisphere) were collected and weighed. The samples were immediately processed, flash-frozen at dry ice, and stored at -70°C until subsequent analysis.

**Sample Formulation.** Solid samples were suspended in a mixture of Tween 80 – 0.9% Saline (5% : 95%, v/v), using ~98% of calculated volume in an 8 mL glass vials closed with a screw cap. The mixture was vortexed (~10 seconds) and sonicated at 85ºC (~five minutes). Process was repeated multiple times as needed, until the initially coarse suspension gradually turned into a fine dispersion, and eventually into a clear solution. Mildly acidic formulation pH (4.0 – 6.0) was neutralized with 1M NaOH to pH 7.0 and volume was adjusted to the final desired concentration using the Saline-Tween solution.

Noribogaine hydrochloride was administered in 10 mL/kg dose volume, using concentration of 1 mg/mL (dose 10 mg/kg) and 5 mg/mL (dose 50 mg/kg). *N*-ethyl-noribogaine hydrochloride was administered in 5 mL/kg dose volume, using concentration of 2 mg/mL (dose 10 mg/kg) and 10 mg/mL (dose 50 mg/kg). Doses and formulations were calculated based on the free base form of compound and corrected for drug purity. Working formulations were prepared five minutes prior to initiation of PK study.

**Bio-samples Processing.** *Plasma samples* (40 µL) were mixed with 200 µL of IS(plasma) solution. After mixing by pipetting and centrifuging for four minutes at 6,000 rpm, 0.5-1.0 µL of each supernatant was injected into LC-MS/MS system. Internal standards IS(plasma) used (noribogaine: Crizotinib 1000 ng/mL; *N*-ethyl-noribogaine Sildenafil 400 ng/mL) in water-methanol mixture 1:9, v/v.

*Brain samples* (weight 95 mg – 157 mg) were homogenized with five volumes of IS(brain) solution (1 w + 5 v, e.g. 100 mg + 500 µL) using glass beads (115 mg ± 5 mg) in The Bullet Blender® homogenizer for 30 seconds at speed 8. After this, the samples were centrifuged for four minutes at 14,000 rpm, and 0.25-1 µL of each supernatant was injected into LC-MS/MS system. Internal standards IS(brain) used (noribogaine: Crizotinib 1000 ng/mL; 5-ethoxy-ibogamine Sildenafil 1600 ng/mL) in water-methanol mixture 1:4, v/v.

**Bio-samples Analysis.** The analyte concentration in samples was determined using high performance liquid chromatography/tandem mass spectrometry (HPLC-MS/MS). Shimadzu HPLC system comprised 2 isocratic pumps LC-20AD, an autosampler SIL-20ACXR, a sub-controller FCV-14H and a degasser DGU-20As. Mass spectrometric analysis was performed using an API 3000 (triple-quadrupole) instrument from AB Sciex (Canada) with an electro-spray (ESI) interface. The data acquisition and system control were performed using Analyst 1.6.3 software from AB Sciex.

*Chromatographic Conditions*:
Column InfinityLab Poroshell 120 EC-C18, 2.1 × 50 mm, 4 µm
Mobile phase A: Acetonitrile : Water : Formic acid = 50 : 950 : 1
Mobile phase B: Acetonitrile : Formic acid = 100 : 0.1
Linear gradient: 4 min 0% B, 1 min 100% B, 1.2 min 100% B, 1.21 min 4% B, 2.7 min stop
Elution rate: 400 µL/min. A divert valve directed the flow to the detector from 1.1 to 1.5 min
Column temperature: 30°C

*MS/MS Detection*:
Scan type: Positive MRM, Ion source: Turbo spray, Ionization mode: ESI
Nebulize gas: 15 L/min, Curtain gas: 8 L/min, Collision gas: 4 L/min
Ionspray voltage: 5000 V, Temperature: 400°C

| *Compound ID* | *Parent, m/z* | *Daughter, m/z* | *Time, ms* | *DP, V* | *FP, V* | *EP, V* | *CE, V* | *CXP, V* |
|---|---|---|---|---|---|---|---|---|
| **Noribogaine** | **297.175** | **122.200** | **90** | **66** | **270** | **11** | **47** | **8** |
| Crizotinib | 450.062 | 260.000 | 90 | 61 | 340 | 11 | 37 | 20 |
| ***N*-ethyl-noribogaine** | **325.201** | **122.3** | **50** | **81** | **320** | **11** | **47** | **10** |
| Sildenafil | 475.055 | 58.1 | 50 | 66 | 360 | 11 | 87 | 10 |

**Analyte Quantification.**
*Calibration solutions:* Stock solution (2 mg/kg) of test compound in DMSO was consecutively diluted with either IS(plasma) or IS(brain) to obtain a series of calibration solutions with final concentrations of 10,000, 4,000, 2,000, 1,000, 400, 200, 100, 40, 20, 10, 4, and 2 ng/mL (for 10 mg/kg dosing group); and 10,000, 4,000, 2,000, 1,000, 400, 200, 100, 40, and 20 ng/mL (for 50 mg/kg dosing group).

*Plasma calibration:* Blank mouse plasma samples (40 µL) were mixed with 200 µL of the corresponding calibration solution. After mixing by pipetting and centrifugation for four minutes at 6000 rpm, 0.5 – 1.0 µL of each supernatant was injected into LC-MS/MS system.

*Brain calibration:* Blank brain samples (weight 100 mg ± 1 mg) were homogenized in 500 µL of corresponding calibration solution using glass beads (115 mg ± 5 mg) in The Bullet Blender® homogenizer for 30 seconds at speed 8. After this, the samples were centrifuged for four minutes at 14,000 rpm, and 0.25 µL of each supernatant was injected into LC-MS/MS system.

**Pharmacokinetic Method Analysis.** The concentrations of the test compound below the lower limits of quantitation (LLOQ = 20 ng/mL for plasma, 10 ng/g and 100 ng/g for brain samples) were designated as zero. The pharmacokinetic data analysis was performed using noncompartmental, bolus injection or extravascular input analysis models in WinNonlin 5.2 (PharSight). Data below LLOQ were presented as missing to improve the validity of $T_{½}$ calculations.

For each treatment condition, the final concentration values obtained at each time point were analyzed for outliers using Grubbs' test with the level of significance set at $p < 0.05$.

**Plasma and Brain Tissue Protein Binding (Equilibrium Dialysis).** Determined plasma / brain concentrations and pharmacokinetic parameters were corrected for non-specific plasma and brain tissue protein binding using data determined according to published procedure.[25,26]

Pooled rat brains (Sprague Dawley, male, *n* = 3) and non-sterile rat (Sprague Dawley) plasma with Li-heparin were used.

#### 13. Animal Behavioral Studies and Usage.

**General Mouse Use.** All experimental procedures involving animals were approved by the Columbia University Institutional Animal Care and Use Committee (IACUC) and adhered to principles described in the National Institutes of Health Guide for the Care and Use of Laboratory Animals. The studies were conducted at AAALAC accredited facilities. Animals received regular veterinary care (weekly by institutional veterinarians) including daily health monitoring (by experimenters) of the animals (observing home cage behaviors, nesting, and body weight). All procedures were designed to minimize any stress/distress. Healthy adult male mice C57BL/6J (10 – 15 weeks old) were purchased from the Jackson Laboratory (Bar Harbor, ME) and housed five mice per cage with food and water available *ad libitum*. Mice were maintained on a 12 h light/dark cycle (lights on 7:00-19:00) and all testing was done in the light cycle. The temperature was kept constant at 22 ± 2 degrees Celsius, and relative humidity was maintained at 50 ± 5 percent.

All pertinent experiments involving ibogaine were conducted in the laboratory of Dr. David Sulzer and in compliance with the rules and regulations as specified by both the New York State Department of Health Bureau of Narcotic Enforcement Class 7 Individual Controlled Substance Activity License and the United States Drug Enforcement Administration Schedule I Controlled Substances Researcher License.

**Drug Preparation and Administration for Animal Studies.** Tetrabenazine (Sigma-Aldrich; St. Louis, MO) was administered in a vehicle of 40% 2-Hydroxypropyl-β-cyclodextrin (Sigma-Aldrich; St. Louis, MO) in USP grade 0.85% saline (Teknova; Hollister, CA) and final pH was adjusted to 7.0 with 1 M of NaOH (Sigma-Aldrich; St. Louis, MO). All other compounds were dissolved in USP grade 0.85% saline with 5% Tween-80 (Sigma-Aldrich; St. Louis, MO). Sonication and gentle heating are applied until complete dissolution. The compounds are subsequently filtered through 0.23 μm filters (Cytiva; Buckinghamshire, UK) into a new glass vial. All compounds were administered at a selected subcutaneous dose at a volume of 10 mL/kg of body weight.

**Behavioral Assays in Mice.**

*Catalepsy Bar Test Protocol.* To assess for catalepsy, C57BL6J mice were administered a drug/dose combination at t = 0 minutes via subcutaneous injection. Mice were tested individually at t = 30, 60, and 90 minutes, 3×. To assess for potentiation, C57BL6J mice are administered tetrabenazine (10 mg/kg) at t = 0 minutes and noribogaine (10, 20, or 30 mg/kg) or citalopram (5 or 10 mg/kg; Sigma-Aldrich; St. Louis, MO) at t = 30 minutes through subcutaneous injection. Mice are tested individually at t = 60 minutes, 3×. Catalepsy is defined by the absence of movement for 30–60 seconds with a cutoff time set to 60 seconds. The automated Maze Engineers Catalepsy Bar test (MazeEningeers; Skokie, IL), featuring sensors to detect forelimb removal from the bar, facilitates automatic recording of forelimb movement duration from the bar to the ground.

*Open Field (OF) Locomotion Protocol.* Mice were allowed to habituate for 30 minutes. Immediately after receiving a subcutaneous injection of the compound solution, mice were then placed gently in a clear Plexiglass arena (27.31 × 27.31 × 20.32 cm, Med Associates ENV-510; Fairfax, VT) lit with dim light (~5 lux) and allowed to ambulate freely for 60 min. The locomotion of the animals was tracked by infrared beams embedded along the X, Y, Z axes of the area and automatically recorded. Data was collected on Activity Monitor by Med Associates (Fairfax, VT).

### E. References.

(6) Wong, E. H. F.; Sonders, M. S.; Amara, S. G.; Tinholt, P. M.; Piercey, M. F. P.; Hoffmann, W. P.; Hyslop, D. K.; Franklin, S.; Porsolt, R. D.; Bonsignori, A.; Carfagna, N.; McArthur, R. A. Reboxetine:

A Pharmacologically Potent, Selective, and Specific Norepinephrine Reuptake Inhibitor. *Biol. Psychiatry* **2000**, *47* (9), 818–829. https://doi.org/10.1016/S0006-3223(99)00291-7.
